# Breathing dysfunction and alveolar damage in a mouse model of Dravet syndrome

**DOI:** 10.1101/2022.05.20.492889

**Authors:** Min-Jee Goh, Cassandra E. Deering-Rice, Jacklyn Nguyen, Evalien Duyvesteyn, Alessandro Venosa, Christopher A. Reilly, Cameron S. Metcalf

**Author notes:** Corresponding author: Cameron S. Metcalf, PhD., Department of Pharmacology & Toxicology, University of Utah; 30 S. 2000 E. Room 201, Salt Lake City, UT 84112.

## Abstract

**Objective:** The incidence of Sudden Unexpected Death in Epilepsy (SUDEP) is especially high in those with Dravet syndrome (DS). Risk factors have been identified, but the mechanism(s) by which death occurs is not fully understood. Evidence supports ventilatory dysfunction in the pathophysiology of SUDEP. Understanding specific respiratory patterns present at baseline and after seizures at different ages, as well as the health of lung tissue, will allow us to better understand how sudden death occurs in this population.

**Methods:** Whole body plethysmography (WBP) was used to monitor respiration before and after electrically induced seizure in the *Scn1a^A1783V/WT^* mouse model of DS weekly for a period of four weeks. Following the four-week WBP study, lungs from surviving animals were collected and stained with hematoxylin and eosin and Weigert’s elastin and the density of tissue and elastin were analyzed.

**Results:** Breathing was diminished in the DS mouse at baseline and following evoked seizures in younger aged mice (P18-P24), consistent with prolonged post-ictal inspiratory time and low respiratory drive compared to the response seen in older animals. In older DS mice, consisting of those that have survived a critical period for mortality, the response to seizure was more robust and included higher respiratory drive, peak inspiratory and expiratory flow rates, tidal and expiratory volumes, and breathing frequency compared to wild-type and relative to baseline. Alveolar damage was also observed in P46-P52 DS mice.

**Significance:** Differences in specific respiratory parameters in younger DS animals, during the time when mortality is greatest, compared to older DS animals (i.e. those that have survived the critical period) may allow us to better understand respiratory differences contributing to SUDEP. Lung tissue damage in DS may also contribute to respiratory dysfunction in SUDEP.

**KEY POINTS:**

- Baseline respiration is diminished in DS mice compared to wild type.
- Electrically induced seizure produced a different respiratory response in younger DS mouse compared to older DS animals.
- Alveolar septal damage is present in DS mice.
- Baseline and post-ictal breathing dysfunction and inefficient oxygenation and CO_2_ clearance likely potentiated by lung damage may serve as a potential mechanism by which SUDEP occurs in DS.

## INTRODUCTION

Sudden Unexpected Death in Epilepsy (SUDEP) is a risk for all patients with epilepsy, but the incidence is especially high in those with Dravet syndrome (DS) [Shmuely, et al. 2016]. DS is a genetic form of epilepsy that typically manifests before the first year of life [Dravet 2011]. Hyperthermia-induced seizures are common early in the disease, but patients experience many seizure types as the condition progresses. These seizures are often refractory to treatment and event frequency positively correlates with risk of SUDEP [Dravet, et al. 2011; Shmuely, et al. 2016; Ricobaraza, et al. 2019]. In addition to seizures, patients suffer from many cognitive and psychomotor comorbidities [Shmuely, et al. 2016; Dravet, et al. 2011]. DS primarily results from a mutation in the *SCN1A* gene which encodes the alpha subunit of the voltage-gated sodium channel Nav1.1. Disinhibition caused by inhibitory interneuron Nav1.1 dysfunction is believed to be the primary mechanism by which hyperexcitability and the clinical manifestations of DS arise [Ricobaraza, et al. 2019, Ogiwara, et al. 2007]. Despite knowledge of factors contributing to DS pathophysiology, we do not understand the mechanisms contributing to sudden death.

While many mechanisms of SUDEP have been proposed, its underlying pathophysiology is still not fully understood. Recent clinical evidence supports respiratory dysfunction following seizures as an important cause of SUDEP [Ryvlin, et al. 2013; Kim, et al. 2018]. Moreover, respiratory distress has been observed in witnessed SUDEP cases [Moseley, et al. 2012]. In a comprehensive survey of epilepsy monitoring units, increased breathing rate, a common ventilatory response to hypoxia, was found to consistently lead the series of post-ictal cardiorespiratory events that resulted in SUDEP and terminal apnea always occurred before the permanent cessation of cardiac activity that ultimately led to death [Ryvlin, et al. 2013]. Furthermore, breathing dysfunction is observed in human patients with DS and in DS animal models [Kim, et al. 2018; Kuo, et al. 2019; Simeone, et al. 2018]. This suggests that seizures may present a recurrent respiratory challenge in DS and inability to adequately regulate breathing responses to such post-ictal respiratory insults may eventually contribute to dysfunction in normal restorative breathing as a basis for SUDEP.

Mouse models with mutations in *SCN1A* have been developed and characterized. A mutation such as the *Scn1a^A1783V/WT^* heterozygous (Het) knock-in mutation has been identified in human DS patients and recapitulates the human disease in mice [Ricobaraza, et al. 2019]. Het mice experience febrile and spontaneous seizures and have a higher mortality rate than wild-type (WT) littermates [Pernici, et al. 2021, Pernici, et al. bioRxiv]. Death most frequently occurs during post-natal day 15 (P15) to P30 [Pernici, et al. 2021]. Therefore, mice with this mutation demonstrate key clinical findings of DS and SUDEP and further study in this model may shed light on potential mechanisms contributing to mortality in this patient population.

Our aim was to provide a detailed analysis of how baseline and post-seizure respiration differs in the *Scn1a^A1783V/WT^* Het mouse model of DS compared to WT and how baseline and post-ictal breathing change with age. Whole body plethysmography (WBP) involves noninvasive monitoring of respiration and provides information on ventilatory patterns [Lomask, et al. 2006, Criee, et al. 2011]. A longitudinal WBP study revealed differences in Het and WT breathing patterns before and after weekly electrically-induced seizures. Further, this seizure type (6 Hz 44 mA) has been shown to activate neurons in the hindbrain [Barton 2001] and therefore may be used as a seizure model that mimics potentially terminal seizures. We also evaluated lung tissue of DS animals to assess another potential contributor to respiratory dysfunction in these animals and found alveolar septal damage in Het mice, and particularly those who had experienced a terminal seizure. Together, these results reveal altered ventilatory patterns at baseline and following induced seizures that change with age and suggest that either failure to develop or loss of proper lung structure may underlie some of the respiratory differences we observe in DS.

## MATERIALS AND METHODS

### Animals

All animal care and experimental procedures were approved by the Institutional Animal Care and Use Committee of the University of Utah. *Scn1a^A1783V/WT^* (Het) animals were generated by crossing a floxed stop male *Scn1a^A1783V/WT^* with a Sox2-Cre female to produce heterozygous and wild-type offspring as previously described [Pernici, et al. 2021]. Mice were housed under a 12h light/12h dark light cycle and had access to food and water *ad libitum* except during plethysmography experiments. Age-matched mice of each genotype were used for experiments. Both male and female mice were included in the studies.

### Whole body plethysmography (WBP)

Whole body plethysmography (Scireq vivoFlow) was used to monitor respiration in Het and WT animals twice weekly for 4 weeks at the following ages: week 1, post-natal day 18-24 (P18-P24); week 2, P25-P31; week 3, P32-P38; week 4, P39-P45. Animals were weighed once weekly prior to WBP monitoring. Twice weekly, animals were placed into individual plethysmography chambers where they were allowed to move freely. Chambers (440 mL) maintained constant air flow (0.5L/min) via a bias flow ventilator. After a 45-minute acclimation period, 15 minutes of baseline respiration was recorded for analysis. Following the baseline recording, a sham stimulation (day 1) or a 6 Hz 44 mA corneal stimulation (day 2) was given. Electrically induced seizure was chosen over hyperthermia-induced seizure due to the speed of seizure induction (minimizing time out of the WBP recording chamber) and concerns of temperature changes affecting the WBP results. Respiration was monitored for 15 minutes immediately following sham and seizure stimulation and plotted with and without baseline normalization as noted in figure legends. Respiratory activity was monitored by Emka IOX2 software. Parameters of interest include time of inspiration and expiration (Ti and Te, msec), peak inspiratory and expiratory flow (PIF and PEF, mL/sec), tidal and expiratory volume (TV and EV, mL), breathing frequency (f, breaths/min, calculated as 1/[time of inspiration + time of expiration]), end inspiratory and expiratory pause (EIP and EEP, msec), enhanced pause (Penh, calculated as (Te/RT – 1)*(PEF/PIF) where RT is relaxation time, the time (msec) it takes to expire a constant percentage of tidal volume), expiratory flow 50% through expiration (EF50, mL/sec) and minute ventilation (MV, mL/min). From these parameters, inspiratory time/expiratory time (Ti/Te), and inspiratory (TV/Ti) and expiratory (TV/Te) drive were calculated.

### 6 Hz seizure induction

Seizures were induced using an A-M Systems stimulator. Following application of 0.5% tetracaine to the eyes, a 3 second 6 Hz 44 mA seizure was induced via corneal stimulation, as previously described [Metcalf, et al 2017; Barton, et al. 2001]. Seizures were characterized by stun and forelimb clonus. For sham seizure stimulations, tetracaine was applied to eyes and corneal electrodes were placed, but electrical stimulation was not delivered.

### Statistical methods

The 15-minute baseline recording was averaged to represent the baseline value for each respiratory parameter. Time courses consisted of one baseline measurement (average measurement of the 15-minute baseline recording, time 0 min) and 7 time points following seizure or sham stimulation: the average of minute 0-1 (time 1 min), average of minute 1-2 (time 2 min), average of minute 2-3 (time 3 min), average of minute 3-4 (time 4 min), average of minute 4-5 (time 5 min), average of minute 5-10 (time 10 min), and average of minute 10-15 (time 15 mine). Time courses were normalized to the averaged baseline value (time 0 min) to illustrate the magnitude of the change in responses. Absolute measurements are provided in the data supplement. Statistical comparisons were conducted using Student’s t-test for baseline values and two-way (repeated measures) ANOVA for time course data. Significance was defined as a p value <0.05. All analyses were conducted with GraphPad Prism 9. Data are represented as the mean + standard error of the mean.

### Histology

One week following the 4-week WBP study (P46-P52), lungs were fixed by tracheal instillation of 10% neutral buffered formalin at a constant pressure (25 cm H_2_O). Following paraffin embedding, 6 *μ*m sections were cut and stained with Hematoxylin & Eosin (H&E) by the Associated Regional and University Pathologists Inc., at the University of Utah.

Unstained lung sections collected from Het and WT mice were deparaffinized in xylene, subjected to serial ethanol washes (100%-50%), and then distilled water. Slides were first stained in Weigert’s Iron Hematoxylin solution for 10 min at room temperature (Polysciences Inc., Warrington, PA). This was followed by 2 min wash in tap water, and incubation in Resorcin Fuchsin solution for 45 min. Slides were then rapidly rinsed in 95% ethanol and washed in tap water. Van Gieson’s solution was added for 1 min and extra stain removed with tap water. Lastly, slides were dehydrated with 95% and 100% alcohol, followed by xylenes. Mounting was performed as described. A minimum of five 400x images were taken for each lung section. Each image was randomly selected from regions within 500 μm from the visceral pleural wall and avoiding blood vessels and large airways, as those express high levels of elastic fibers. Images were then converted to 8-bit using Image J, and elastic fibers quantified as the relative percent of each image, by applying a fixed threshold to all images. Those same images were further analyzed on ImageJ by removing the threshold setting, thus highlighting the percentage of tissue in each image. For both analyses, the average of the 5 images was used to represent staining for each individual mouse.

## RESULTS

### Survival and body weights during 4 weeks of observation

WT and Het mice were evaluated beginning at P18 and continuing through P45. Several mice died during the study, consistent with previous observations [Pernici, et al. 2021]. Deaths of Hets occurred between P26 and P45; one death occurred at each of the following ages: P26, P30, P35, P37, P38, and P45. Male Hets had a higher survival rate; of the Hets that died, only one was male (P45) (**Figures 1A, 1B**). All WT mice survived the 4-week study. Body weight was not significantly different between Het and WT except in week 2 of the study (P25-P31) (**Figure 1C**).

**Figure 1.**
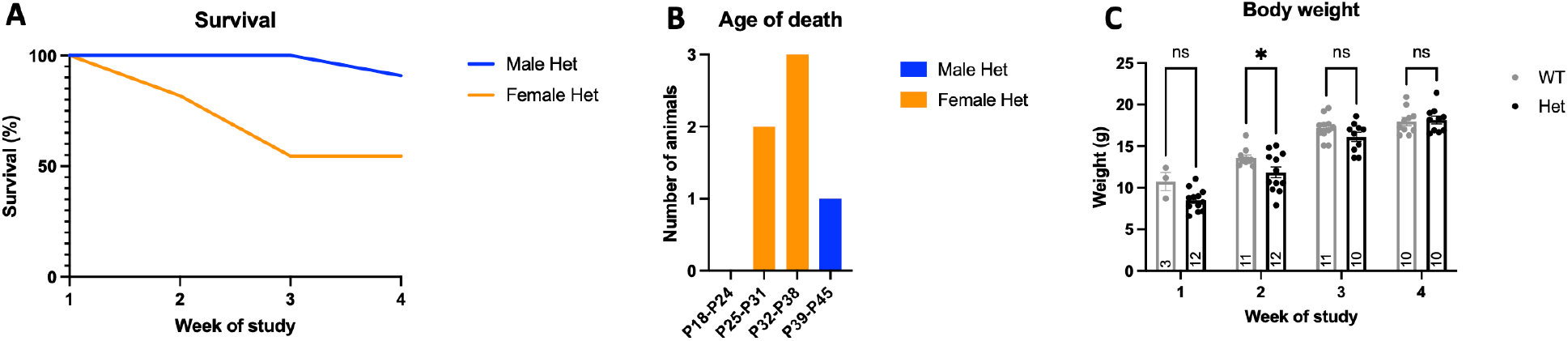
Survival, age of death, and body weight changes for WT and Het mice evaluated for 4 consecutive weeks. WT and Het mice were observed twice weekly for 4 consecutive weeks starting at P18. Deaths were observed during the study, primarily between post-natal day 26 (P26) and P37 (weeks 2-4; (**A**)). A majority of deaths were in females, with only one death observed in males (**B**). Weekly body weights were obtained and were significantly different in P25-P31 Hets and WT (**C**). Two-way ANOVA, *P<0.05.

### Baseline respiration is diminished in Het mice

One-second representative waveforms of WT and Het breathing at week 3 baseline illustrated abnormalities in Hets including significantly increased time of inspiration and reduced breathing frequency. Time of expiration appears to be greater in WT compared to Hets during this time point but this is not significant. Additionally, an abnormal sawtooth pattern was seen during the exhalation phase of Hets (**Figure 2B**) that was not present in the WT waveform (**Figure 2A**).

**Figure 2.**
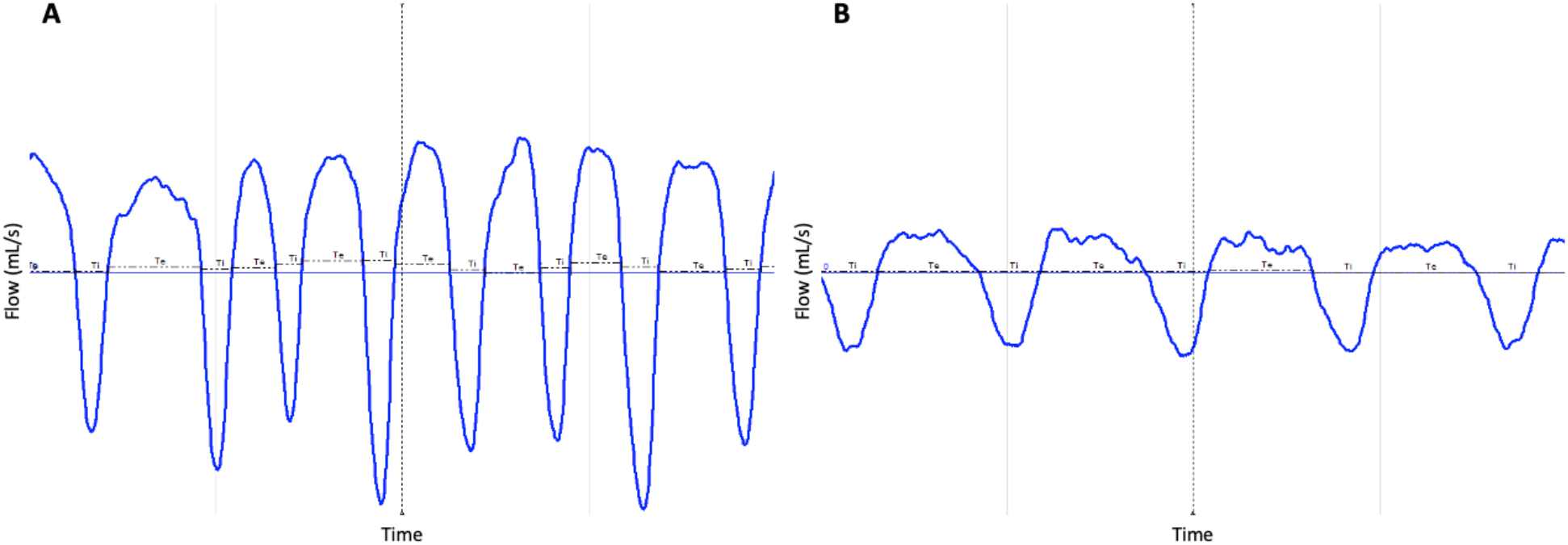
Representative respiratory waveforms of baseline breathing for WT and Het mice. Representative respiratory waveforms (1 second) of P32-P38 WT (**A**) and Het (**B**) show breathing patterns differ at baseline. Time of inspiration (Ti) is greater, breathing frequency is lower, and a sawtooth appearance was observed during the exhalation phase of Hets.

Several differences in baseline respiration were observed between Het and WT mice across the study period (**Table 1**). Specific parameters of concern are inspiratory drive, calculated as tidal volume over time of inspiration (TV/Ti) and minute ventilation (TV*F) which are both decreased in the Het mice after week one, potentially resulting in hypercapnia. Breathing frequency was also significantly reduced for Hets at week 3. Peak inspiratory flow and peak expiratory flow were lower in Hets in weeks 2-4. Expiratory flow 50% through expiration and expiratory volume were reduced in Hets at most points of the study. These results show substantial baseline respiratory dysfunction in Hets even prior to additional insult associated with seizure. There were no differences in baseline end inspiratory pause, end expiratory pause, and enhanced pause between the two groups. See supplementary tables S1 and S2 for all baseline respiratory parameter mean + SEM values.

**Table 1.**
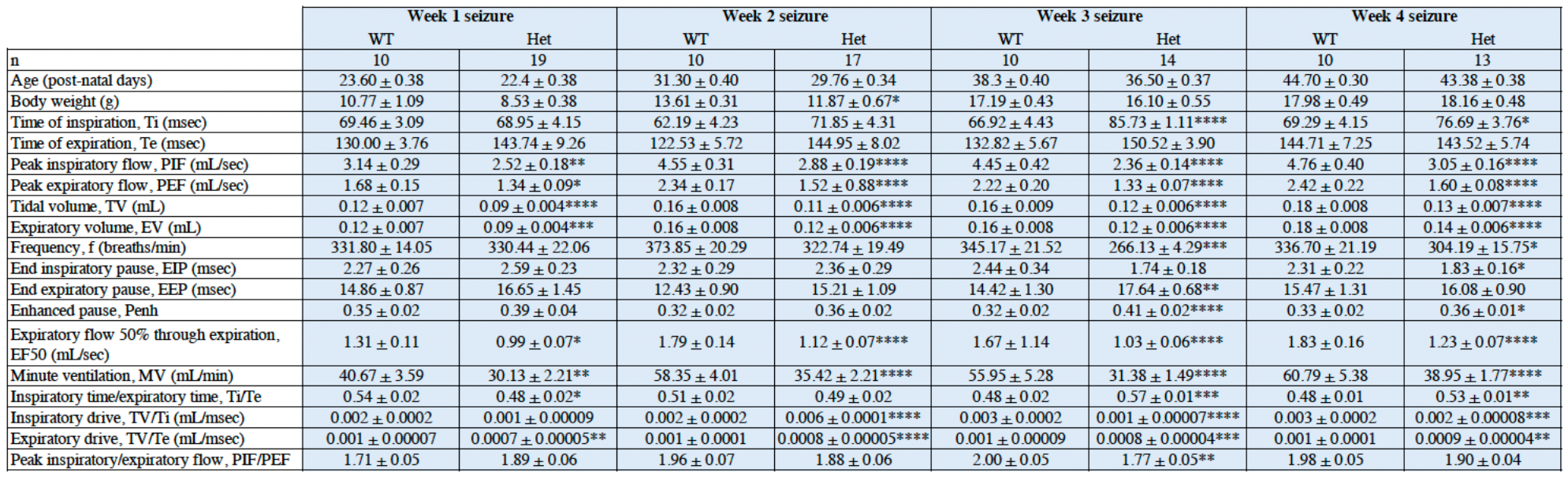
Baseline respiratory parameters in Het and WT prior to seizure induction. Data are presented as mean + SEM. Student’s t-test, *P<0.05, **P<0.01, ***P<0.001, ****P<0.0001.

### Post-seizure respiration is diminished in younger Hets

Following the acclimation period and 15-minute baseline recording, a sham stimulation or a 6 Hz 44 mA seizure was induced and respiration was monitored for an additional 15 minutes. This post-seizure respiration was normalized to the averaged pre-seizure baseline value. At week 1, pre-seizure baseline breathing was not as robust as seen in weeks 2-4 and therefore the normalized values closely mimicked the absolute values (Table 1). Sham stimulation did not result in any major changes between WT and Hets that was not reflected in the baseline results (data not shown). Following an induced seizure in week 1, P18-P24 Hets had greater change in time of inspiration (ΔTi) (**Figure S2**) compared to WT, indicating a slower inhalation rate among DS mice. The change in time of expiration (ΔTe) relative to baseline did not differ in Hets and WT (**Figure S2A**) The Ti/Te ratio peaked post-ictally and this increase was significantly greater in Hets, indicating a potential increase in mean airway pressure (**Figure 3Ai**). The change in inspiratory drive (ΔTV/Ti, **Figure 3Aii**) did not change in Hets or WT following seizure induction, but expiratory drive (ΔTV/Te, **Figure 3Aiii**) was briefly elevated in Hets post-seizure. Similarly, there was no difference in the change in peak inspiratory flow between the two groups (**Figure 3Aiv**) but the change in peak flow rate of exhalation (**Figure 3Av**) was temporarily increased with seizure induction. There were no significant differences in the change in breathing frequency (**Figure 3Avi**), tidal volume (**Figure 3Avii**), and expiratory volume (**Figure 3Aviii**) between week 1 Hets and WT with electrical seizure induction. There were no significant differences in the change in end inspiratory pause, end expiratory pause, expiratory flow 50% through expiration, or minute ventilation throughout the post-seizure period between P18-P24 Hets and WT (**Figure S2A**).

**Figure 3.**
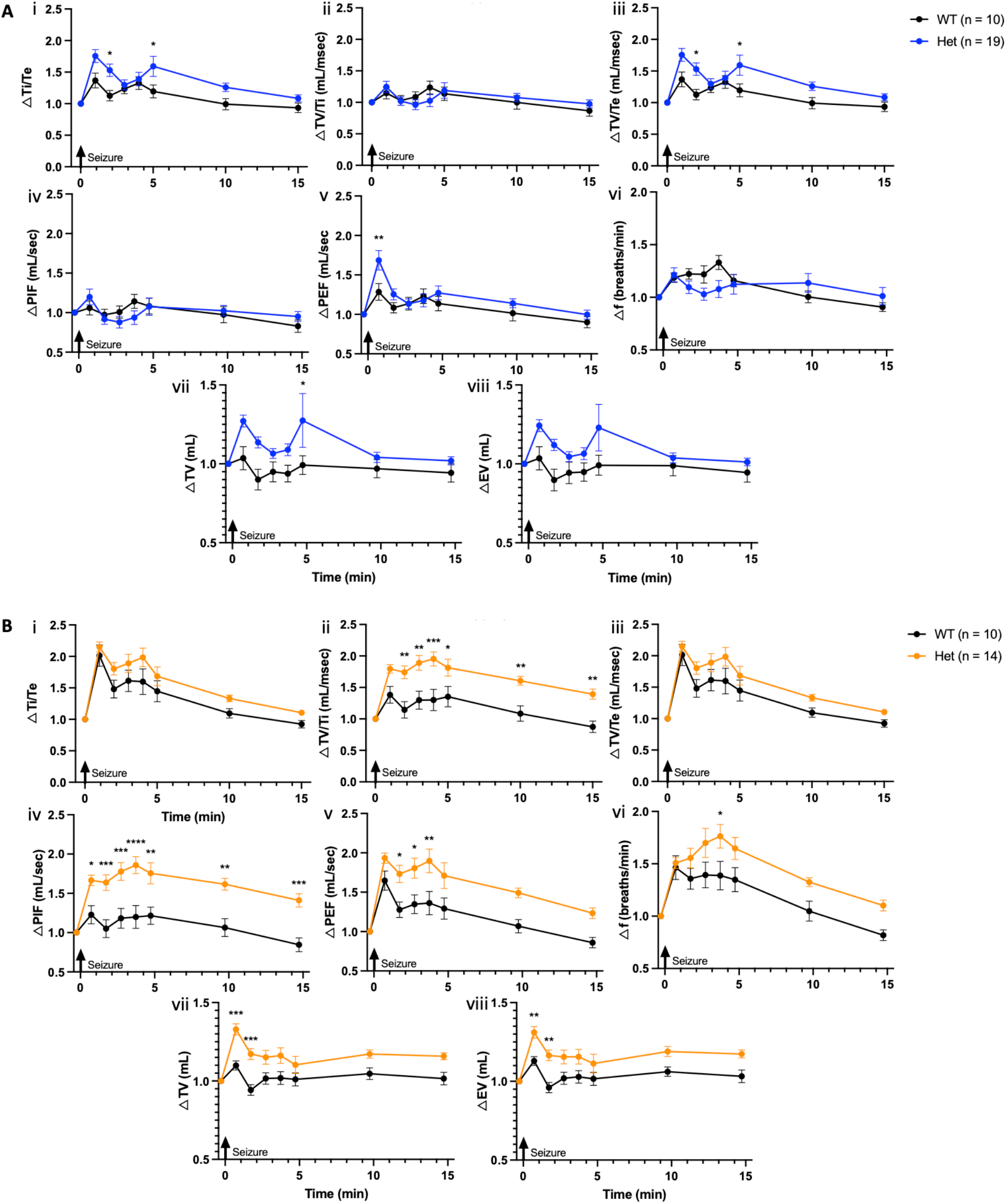
Respiration following electrically induced 6 Hz seizure in WT and Het. Fifteen minutes of post-seizure respiratory monitoring normalized to baseline shown for week 1 (**A**) and week 3 (**B**) WT vs. Het differences suggest age-dependent alteration of breathing following seizure. Change in inspiratory/expiratory ratio (ΔTi/Te; **i**), inspiratory drive (ΔTV/Ti; **ii**), expiratory drive (ΔTV/Te; **iii**), peak inspiratory flow (ΔPIF; **iv**), peak expiratory flow (ΔPEF; **iv**), breathing frequency (Δf; **vi**), tidal volume (ΔTV; **vii**), and expiratory volume (ΔEV; **viii**) derived from waveforms collected. Two-way ANOVA, *P<0.05, **P<0.01, ***P<0.001, ****P<0.0001.

**Figure 4.**
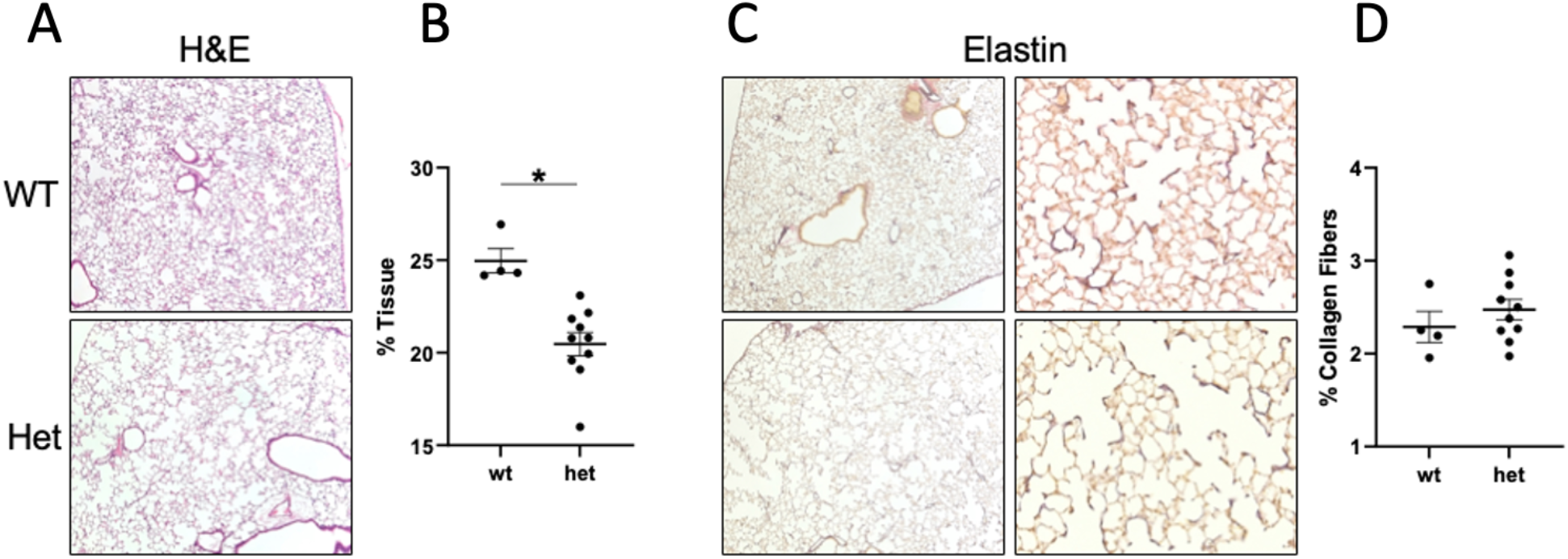
Histological assessment of WT and Het lungs. (**A**) Hematoxylin and Eosin and (**C**) Weigert’s elastin stain of tissue sections were prepared from WT and Het mice age post-natal day P46-P52. Original magnification: 100x or 400x. Representative images of distal lung from at least 3 mice/group are shown. Quantification of (**B**) total tissue area/alveolar density and (**D)** elastin fiber (Weigert’s) in WT and Het mice aged post-natal day 46 to 52. Each point represents the mean of five 100x images of randomly selected regions from each slide. Images were converted to 8-bit on ImageJ. For total tissue quantification, a threshold highlighting all of the tissue was used; for elastin stain, a threshold highlighting only elastic fibers (blue) was selected. The same threshold was applied to all images. Data are presented as mean ± SEM (n = 4-10), analyzed using unpaired t-test. A p<0.05 (*) was considered significant.

In week 2, there were no significant differences from baseline in post-seizure breathing between Het and WT relative to baseline. However, the altered respiratory patterns still persisted (**Figures S2 and S3**). The greatest differences in post-seizure breathing between Het and WT were observed in week 3 animals. Baseline values of time of inspiration in Het mice are significantly higher at week 3 (P32-P38) and therefore the normalized graphs (**Figure 3B**) and the observed responses (**Figure S3B**) are quite different. Unlike week 1, the change in time of inspiration (ΔTi) in week 3 P32-P38 Hets was reduced by a greater magnitude than that of WT after seizure induction and remained lower until the end of the 15-minute post-seizure monitoring period (**Figure S2B**). As in week 1, there was no difference in post-ictal time of expiration, expiratory drive, or Ti/Te relative to baseline between Het and WT (**Figure S2B, 3Biii and 3Bi**). There was a greater post-ictal increase in inspiratory drive relative to baseline (ΔTV/Ti) (**Figure 3Bii**), however the absolute TV/Ti was still lower than WT mice (**Figure S3B**). Post-seizure increases in the magnitude of change in peak inspiratory flow (**Figure 3Biv**), peak expiratory flow (**Figure 3Bv**), breathing frequency (**Figure 3Bvi**), tidal volume (**Figure 3Bvii**), and expiratory volume (**Figure 3Bviii**) was greater in Hets than WT, though absolute values still remained lower than WT mice. Change in end inspiratory and expiratory pause relative to baseline were also observed (**Figure S2B**), however the absolute values indicate that there are no differences in apnea following inhalation and exhalation in Hets compared to WT (**Figure S3B**). There were no differences in the post-ictal change of enhanced pause in the week 3 animals. The increases in expiratory flow 50% through expiration and minute ventilation were greater in Hets than in WT post-seizure (**Figure 3SB**). The trends observed in week 4 animals were similar to those seen in week 3.

### Alveolar damage is present in Het lungs

Lungs from Het and WT animals were collected, sectioned, and stained for hematoxylin and eosin (H&E) and Weigert’s elastin stain solution to visualize tissue morphology and elastin content. H&E staining of lung tissue revealed alveolar septal disruption in regions near the visceral pleural exclusively in Het mice (**Figure 5**). Consistent with this, acquisition of images from these peripheral areas were collected and total tissue density analysis revealed significantly reduced tissue abundance in Het lungs, akin to an emphysematous lung. Further analysis examining disruption in elastin fibers showed no differences between Het and WT animals (**Figure 5**).

## DISCUSSION

Clinical and preclinical evidence supports central respiratory dysfunction as a primary contributor to SUDEP [Ryvlin, et al. 2013; Kim, et al. 2018; Kuo, et al. 2019]. SUDEP is a leading cause of death in DS [Shmuley 2016], but the mechanism(s) by which this sudden death occurs is not understood. We evaluated respiratory function and lung morphology in a mouse model of DS to determine whether breathing abnormalities in this model were consistent with other Dravet mouse models [Kuo, et al. 2019; Kim, et al. 2018; Wenker, et al. 2022] and with clinical SUDEP data [Ryvlin, et al. 2013; Kim, et al. 2018; Villella, et al. 2018]. Specifically, breathing patterns were compared between heterozygous *Scn1a^A1783V/WT^* (Het) DS mice and WT littermate mice during basal and post-ictal periods. Differences in several respiratory parameters including peak flow rates, respiratory volumes, and respiratory drive were observed at baseline. We evaluated which respiratory differences occurred in younger mice during the time when up to 50% of Hets die and found that the change in time of inspiration, Ti/Te ratio, and expiratory drive differed between younger and older DS animals. Potential contributors to respiratory dysfunction occurring outside of the CNS were also explored. Differences in lung tissue structure were observed in this DS mouse model and may contribute to respiratory dysfunction. However, the origins of these differences (i.e., developmental deficiency resulting from *SCN1A* deletion, altered respiration, or seizure burden) remain undefined.

Basal breathing patterns also differed between Hets and WT throughout the four-week WBP study. Inspiratory drive, inspiratory and expiratory volumes, peak inspiratory and expiratory flow rate, and minute ventilation were lower in Hets than in WT throughout the 4 weeks. Diminished breathing in the DS mouse model was especially apparent in week 3, when time to inhale and breathing frequency were also slowed compared to WT. These data illustrating low volume and rate of breathing suggest that respiration is abnormal at baseline compared to WT. These findings are also consistent with baseline breathing reported in another SUDEP mouse model where hypopnea was observed even in the absence of seizures [Simeone, et al. 2018].

In addition to evaluating differences in baseline respiration (inter-ictal), we sought to determine whether breathing was altered following seizures. Using an electrical seizure induction model (6 Hz stimulation), we observed diminished breathing in Hets. The Ti/Te ratio is the inspiratory to expiratory, or I:E ratio, and can be used as an estimate of mean airway pressure [Lessard MR, et al. 1994]. A higher I:E ratio is indicative of elevated mean airway pressure and results in improved oxygenation, while a lower I:E ratio favors CO_2_ clearance. The Ti/Te ratio increased immediately following seizure induction in Hets and WT in all weeks but returned to baseline values by 15 minutes post-seizure. In week 1, the post-seizure increase in Ti/Te relative to baseline was significantly greater in Hets than in WT, likely due to the increase in time of inspiration seen in this group. Blood gas abnormalities are known to occur post-ictally [Farrell, et al. 2016; Simeone, et al. 2018; Kim, et al. 2018] and likely occurred in animals following electrically induced seizure. These data suggest that CO_2_ clearance in P18-P24 Hets may be less efficient after seizures resulting a respiratory insufficiency. There also were robust and long-lasting increases in post-ictal respiratory drive, particularly inspiratory drive (TV/Ti), in week 3 P32-P38 Hets (**Figure 3Bii**) that was not present in the younger week 1 P18-P24 Hets (**Figure 3Aii**). Respiratory drive is a consequence of brainstem respiratory center activity and is responsible for the respiratory muscle activity that facilitates oxygenation and clearance of CO_2_ [Jonkman, et al. 2020]. Increasing inspiration and inspiratory drive, a function of adequate central respiratory output, facilitates efficient oxygenation [Lessard, et al. 1994; Kotani, et al. 2016] and this is observed in the older Hets that have survived the critical period. There also were greater increases in post-seizure peak inspiratory and expiratory flow (**Figure 3Aiv-v, 3Biv-v**), indicative of respiratory effort, and breathing frequency (**Figure 3Avi, 3Bvi**), which would facilitate more effective CO_2_ removal in week 3 Hets compared to week 1 Hets. Respiration involves synchronized activity between central respiratory nuclei, lungs, and muscles involved in breathing. Blood gas abnormalities are detected by peripheral and central chemoreceptors and prompt an increase in respiratory drive to normalize O2 and CO_2_ levels. When CO_2_ levels rise, neurons of the retrotrapezoid nucleus (RTN) are activated via the nucleus of the solitary tract (NTS) and lead to increased respiratory rate, inspiratory amplitude, and active expiration [Zoccal, et al. 2014; Guyenet, et al. 2016]. With continuous or prolonged blood gas imbalance, chemoreceptors can become desensitized, and brainstem respiratory centers may not respond adequately to hypercapnia [Dempsey, et al. 2014]. This *SCN1A* mutation itself may reduce chemoreceptor function in the RTN, implicating this mutation in respiratory control. Accordingly, diminished breathing and reduced respiratory response to CO_2_ has been documented in a similar *SCN1A* DS mouse model [Kuo, et al. 2019]. Blood gas abnormalities and the resulting changes to respiratory drive have also been shown to coincide with the peak age of sudden death, and central inhibition of respiratory drive causes post-ictal death in other mouse models of SUDEP [Simeone, et al. 2018; Schilling, et al. 2019]. These data suggest that younger Hets that are at greatest risk for sudden death may be unable to respond to respiratory demands following a seizure event, while older Hets are able to mediate an adequate respiratory response. This inability to respond to respiratory challenges may predispose DS mice to death via respiratory failure and this may arise through a combination of reduced alveolar structure and central mechanisms. However, further studies are necessary to confirm blood gas abnormalities in Hets. Additionally, the involvement of deficiency in central mechanisms as well as seizure burden in the ability for a subset of younger Hets to regulate blood gases and survive to adulthood is unclear.

Blood gas abnormalities that can be caused by seizures [Farrell, et al. 2016; Kim, et al. 2018] may be exacerbated by the emphysema-like lung phenotype observed in the DS model. To our knowledge, disruption of lung morphology in DS has yet to be investigated as a potential contributing factor to respiratory dysfunction. Histology results show expansion of alveolar airspace, which could be interpreted as a precursor to an emphysematous phenotype in Hets and may present an additional respiratory challenge that Hets are unable to overcome. It is plausible that the structural phenotype observed in these studies precedes pathological development of a chronic pathology such as emphysema, a form of chronic pulmonary disease accompanied by degradation of alveoli that causes breathing difficulty and progressive elastin degradation [Suki, et al. 2017; Sellami, et al. 2016]. This notion is also supported by the absence of elastin fiber loss in the young Het mice, but could also imply a developmental failure at critical stages of lung development arising from *SCN1A* mutation and or respiratory dysfunction. Inadequate gas exchange as well as breathing abnormalities can also result from emphysematous alveolar damage, and this likely contributes to respiratory disruption in this DS mouse model. Evidence of respiratory disruption is supported by WBP studies wherein the anticipated increase in respiratory drive was largely absent in younger Hets that are most vulnerable to SUDEP. This alveolar damage may also explain the irregular expiration waveform observed in Hets and reveals an additional aspect of abnormal respiration that may not be captured by analysis of WBP respiratory parameters alone. Alveolar development in mice occurs between post-natal day 5 and 30 [Warburton, et al. 2010]. It may be possible that lung development is delayed or inhibited in Hets. Such differential lung development may contribute to the differences in postictal respiratory parameters seen in younger Hets compared to WT than is observed in older animals. Future studies will include assessment of lung histopathology in the younger ages of this DS mouse model.

Clinically, lung structure and respiratory function are not commonly assessed in DS patients. Breathing dysfunction occurs in human DS patients [Kim 2018], and patients with DS have more respiratory disruption than in patients experiencing focal seizures. Paradoxical breathing has also been observed in a DS patient [Kim 2018]. Paradoxical breathing is common in COPD patients and this type of breathing is indicative of inadequate gas exchange [Aliverti, et al. 2009; Saraya, et al. 2016]. A case study of an infant with partial deletion of chromosome 2 which includes *SCN1A* described severe seizures as well as pulmonary emphysema [Takatsuki, et al. 2009]. Additionally, respiratory tract infections are a prevalent co-morbidity, affecting almost half of DS patients [Cardenal-Munoz, et al. 2021; Schubert-Bast, et al. 2022]. Though direct respiratory characteristics have yet to be assessed in DS, our results and clinical reports support lung structure and function in this population as a contributor to respiratory dysfunction.

The results presented herein show diminished breathing at baseline in a DS mouse model. Following electrically induced seizure, we observe changes in respiratory patterns, particularly in time of inspiration, respiratory drive, peak flow rates, breathing frequency, and inhalation and exhalation volumes that correspond to diminished breathing in younger Hets and a more robust respiratory response in older Hets that survive the high mortality period. Increased respiratory drive is a homeostatic response to blood gas abnormalities and is mediated by peripheral and central chemoreceptors. We also report alveolar lung damage in the *Scn1a^A1783V/WT^* DS mouse model. This tissue damage may contribute to respiratory insufficiency in DS animals that have diminished breathing even at baseline, ultimately contributing to respiratory decline and sudden death.

## ACKNOWLEDGEMENTS

We would like to thank Kevin Yang for the Python script used for WBP analysis.

