## Supplemental material for "Breathing dysfunction and alveolar damage in a mouse model of Dravet syndrome"

**Table S1.** Values for all respiratory parameters at baseline and following seizure stimulation.

**Table S2.** Values for all respiratory parameters at baseline and following sham stimulation.

### Table S1. Respiratory parameters following seizure stimulation

#### Time of inspiration

Week 1

WT — Time (min) — Animal ID — Week 1

| Time (min) | 83 | 86 | 87 | 88 | 81 | 82 | A | 156 | 158 | 161 | Average | SD | SE |
|---|---|---|---|---|---|---|---|---|---|---|---|---|---|
| 0 | 54.5555556 | 66.7 | 80.0111111 | 58.2444444 | 74.7777778 | 79.5222222 | 59.2111111 | 67.0555556 | 73.2 | 81.3111111 | 69.4588889 | 9.78467777 | 3.09418679 |
| 1 | 66 | 58.3333333 | 66.3333333 | 60.16667 | 65.4 | 66.8333333 | 59.6666667 | 53 | 67.4 | 67.2 | 63.0333337 | 4.92692255 | 1.55802971 |
| 2 | 51 | 63.6666667 | 58.1666667 | 71 | 68.5 | 70.6666667 | 49.6666667 | 48.5 | 63.3333333 | 64 | 60.85 | 8.58776405 | 2.71568944 |
| 3 | 47 | 63.3333333 | 56.8333333 | 85.66667 | 73.8333333 | 71.4166667 | 37.8333333 | 63 | 58.3333333 | 64.3333333 | 62.1583337 | 13.5234399 | 4.2764872 |
| 4 | 51.6666667 | 53.5 | 48.1666667 | 54.1666667 | 71.6666667 | 55.8181818 | 44.3333333 | 42 | 51 | 64.6666667 | 53.6984848 | 8.90991959 | 2.81756397 |
| 5 | 54.1666667 | 56.1666667 | 64.8333333 | 80.5 | 73.8333333 | 55.1666667 | 53.1666667 | 60.5 | 62 | 61 | 62.1333333 | 8.9037376 | 2.81560905 |
| 10 | 81 | 62.4666667 | 79.2333333 | 79.6666667 | 71.2333333 | 71.5 | 61.5666667 | 66.5666667 | 55.7 | 68.9333333 | 69.7866667 | 8.48726279 | 2.68390815 |
| 15 | 83.7666667 | 67 | 81.4333333 | 88.6666667 | 72.9333333 | 81.037037 | 79.3333333 | 67.6666667 | 77.2333333 | 80.7333333 | 77.9803704 | 6.92279083 | 2.18917868 |

Het — Time (min) — Animal ID — Week 1

| Time (min) | B | D | E | F | G | H | 155 | 157 | 159 | 160 | 162 | 190 | 196 | 199 | 201 | 205 | 207 | 208 | 210 | Average | SD | SE |
|---|---|---|---|---|---|---|---|---|---|---|---|---|---|---|---|---|---|---|---|---|---|---|
| 0 | 90.1 | 78.5444444 | 64.8777778 | 89.6888889 | 70.6444444 | 55.4 | 46.2777778 | 50.5111111 | 73.2333333 | 54.2 | 54.6888889 | 45.7444444 | 90.4111111 | 93.6333333 | 62.1111111 | 41.3444444 | 66.1333333 | 89.3666667 | 93.1888889 | 68.9526316 | 18.0974579 | 4.15184158 |
| 1 | 79.3333333 | 59.8 | 67.5 | 86.2 | 66.8 | 65.2 | 60.4 | 66.5 | 70 | 75.4 | 68.2 | 64.5 | 76.1666667 | 71.3333333 | 71.3333333 | 62.8333333 | 74.1666667 | 64.5 | 98 | 70.9561404 | 9.33144857 | 2.14078112 |
| 2 | 116.666667 | 63.6666667 | 99.6666667 | 93.5 | 76 | 49.5 | 59.5 | 60.1666667 | 82.8333333 | 89.1666667 | 78.1666667 | 86.6666667 | 72.3333333 | 76.8333333 | 64.5 | 52.5 | 82.8333333 | 90.1666667 | 90.3333333 | 78.1578947 | 17.1577173 | 3.93625032 |
| 3 | 116.666667 | 53.6666667 | 111.833333 | 103.333333 | 83.8333333 | 61 | 49.8333333 | 59.5 | 82.1666667 | 95.5 | 72 | 73.3333333 | 67.6666667 | 77 | 68.6666667 | 64.1666667 | 76.3333333 | 93 | 101.333333 | 79.5175439 | 19.4771071 | 4.46835481 |
| 4 | 110.833333 | 45.5 | 126 | 127.666667 | 89.5 | 73.5 | 48.5 | 77.1666667 | 82.3333333 | 75.8333333 | 71.1666667 | 63.8333333 | 60.1666667 | 62.5 | 62.1666667 | 59.8333333 | 69.8333333 | 87 | 92.1666667 | 78.1842105 | 23.1973481 | 5.32183664 |
| 5 | 141.166667 | 45.3333333 | 114.333333 | 143.166667 | 78.8333333 | 76 | 55.3333333 | 78.6666667 | 65.5 | 63.8333333 | 71 | 61.1666667 | 54.5 | 56.6666667 | 49.3333333 | 51.6666667 | 70.1666667 | 79 | 95.5 | 76.377193 | 28.5514395 | 6.55014945 |
| 10 | 80.1 | 46.1666667 | 94.9666667 | 103.9 | 71.6 | 64.7 | 52.0666667 | 74.2666667 | 75.1333333 | 57.9333333 | 59.5333333 | 56.1 | 77.2333333 | 42.3666667 | 52.9666667 | 49.5666667 | 70.6666667 | 75.2 | 90.7666667 | 68.1701754 | 17.007282 | 3.90173808 |
| 15 | 79.8666667 | 48.7666667 | 83.8 | 90.0333333 | 66.7333333 | 68.0666667 | 77.9 | 71.3333333 | 78.6 | 61.6666667 | 72.0333333 | 60.8 | 78.8 | 48.8666667 | 68.9666667 | 71.5333333 | 81.8333333 | 83.6333333 | 85 | 72.5385965 | 11.5630705 | 2.65275031 |

Week 2

WT — Time (min) — Week 2

| Time (min) | 83 | 86 | 87 | 88 | 81 | 82 | A | 156 | 158 | 161 | Average | SD | SE |
|---|---|---|---|---|---|---|---|---|---|---|---|---|---|
| 0 | 52.1 | 61.2111111 | 78.6888889 | 51.5888889 | 60.9111111 | 86.9555556 | 55.3444444 | 74.8 | 47.0222222 | 53.2777778 | 62.19 | 13.4068574 | 4.23962057 |
| 1 | 59 | 56.6666667 | 51.5 | 51.1666667 | 54.6666667 | 54.8333333 | 60.1666667 | 98 | 62.1666667 | 58.3333333 | 60.65 | 13.6006649 | 4.30090789 |
| 2 | 50.1666667 | 55.8333333 | 51.8333333 | 52.8333333 | 60.6666667 | 49 | 61.3333333 | 50.6666667 | 57.1666667 | 51.3333333 | 54.0833333 | 4.41605516 | 1.39647926 |
| 3 | 54.6666667 | 49.3333333 | 53.8333333 | 52.5 | 62.1666667 | 54.1666667 | 71.8333333 | 63.3333333 | 56.5 | 54 | 57.2333333 | 6.65053604 | 2.10308415 |
| 4 | 73.1666667 | 50.8333333 | 54.3333333 | 58 | 57.5 | 51.8333333 | 66 | 67.5 | 55 | 40.5 | 57.4666667 | 9.42671539 | 2.98098915 |
| 5 | 50.8333333 | 49.8333333 | 63.8333333 | 52 | 54.5 | 55.5 | 68.8333333 | 85 | 62.5 | 47.8333333 | 59.0666667 | 11.3782335 | 3.59811336 |
| 10 | 63.1 | 53.8333333 | 71.7333333 | 51 | 71.9333333 | 69.5 | 85.5 | 95.2 | 71.2333333 | 60.5666667 | 69.36 | 13.4934808 | 4.2670133 |
| 15 | 61.3333333 | 64.3333333 | 83.4 | 67 | 84.0333333 | 76.6 | 74.3 | 99.4 | 55.3 | 76.5 | 74.22 | 12.9072876 | 4.08164274 |

Het — Time (min) — Week 2

| Time (min) | B | D | E | G | H | 155 | 157 | 160 | 162 | 190 | 196 | 199 | 201 | 205 | 207 | 208 | 210 | Average | SD | SE |
|---|---|---|---|---|---|---|---|---|---|---|---|---|---|---|---|---|---|---|---|---|
| 0 | 63 | 87.6222222 | 75.2888889 | 86.2 | 57.0555556 | 76.3777778 | 56.4333333 | 60.8111111 | 69.0222222 | 85.2 | 102.255556 | 99.9777778 | 82.7666667 | 43.8111111 | 75.8888889 | 42.4444444 | 57.2666667 | 71.848366 | 17.7564342 | 4.30656787 |
| 1 | 66.3333333 | 68.1666667 | 61.8 | 69.6666667 | 46.1666667 | 65.1666667 | 77.8333333 | 68.3333333 | 66.6666667 | 56.8333333 | 62.1666667 | 73.8333333 | 67.5 | 66.5 | 57 | 59.6666667 | 54.6666667 | 64.0176471 | 7.58780196 | 1.84031229 |
| 2 | 80.3333333 | 64.1666667 | 60.6666667 | 73.5 | 50.5 | 64 | 102.166667 | 54 | 55.8333333 | 49.8333333 | 46.5 | 66.6666667 | 49 | 55.1666667 | 47.3333333 | 78.3333333 | 44.5 | 61.3235294 | 15.2737146 | 3.70441991 |
| 3 | 89.5 | 64.3333333 | 56.6666667 | 70 | 54.3333333 | 66.5 | 97.3333333 | 61 | 55.1666667 | 41.6666667 | 39.8333333 | 64.5 | 44.1666667 | 51.6666667 | 41.5 | 93 | 47.6666667 | 61.1078431 | 17.9205365 | 4.34636853 |
| 4 | 53.3333333 | 63 | 49.8333333 | 72.3333333 | 60.1666667 | 69.1666667 | 89.6666667 | 59 | 64.6666667 | 44.5 | 46.8333333 | 50.6666667 | 42.5 | 59.3333333 | 46.8333333 | 60.8333333 | 44.5 | 57.4803922 | 12.2720723 | 2.97641472 |
| 5 | 49 | 54.3333333 | 55.1666667 | 59.1666667 | 68.6666667 | 66.6666667 | 90 | 63.8333333 | 84.6666667 | 45.5 | 46.8333333 | 58.1666667 | 52.1666667 | 60.6666667 | 41 | 51.6666667 | 56.5 | 59.0588235 | 12.9814158 | 3.1484558 |
| 10 | 79.4666667 | 60.3333333 | 58.2666667 | 65.5333333 | 75.2 | 68.8 | 75.2 | 74.3333333 | 89.0333333 | 61.8666667 | 81.7333333 | 58.9666667 | 75.1333333 | 64.1333333 | 56.9 | 56.2666667 | 53.7 | 67.9333333 | 10.3587859 | 2.5123746 |
| 15 | 68.4666667 | 65.6 | 66.7 | 79.8333333 | 81.5 | 59.2333333 | 76.9 | 97.1333333 | 146.033333 | 63.4 | 72.4 | 60.9 | 67.0333333 | 64.5666667 | 85.7 | 62.8666667 | 61.9333333 | 75.3058824 | 20.9115241 | 5.07178956 |

Week 3

WT — Time (min) — Week 3

| Time (min) | 83 | 86 | 87 | 88 | 81 | 82 | A | 156 | 158 | 161 | Average | SD | SE |
|---|---|---|---|---|---|---|---|---|---|---|---|---|---|
| 0 | 51.1333333 | 55.5 | 63.3444444 | 64.4222222 | 87.7333333 | 82.7888889 | 50.6111111 | 76.3888889 | 81.3 | 56.0111111 | 66.9233333 | 14.0043766 | 4.42857272 |
| 1 | 50.3333333 | 51.5 | 45.8333333 | 50.5 | 50 | 47 | 51.1666667 | 62.5 | 69.6666667 | 68.3333333 | 54.6833333 | 8.75681393 | 2.76914771 |
| 2 | 41 | 62.1666667 | 42.1666667 | 48.1666667 | 48.5 | 62.3333333 | 55.8333333 | 76.6666667 | 75.5 | 64.8333333 | 57.7166667 | 12.7717498 | 4.0387819 |
| 3 | 42.3333333 | 63.6666667 | 35.1666667 | 43.1666667 | 47.6666667 | 59.1666667 | 58.1666667 | 77.1666667 | 78.5 | 47.3333333 | 55.2333333 | 14.7269804 | 4.6570801 |
| 4 | 54 | 71.1666667 | 43 | 38 | 44 | 64.6666667 | 47.6666667 | 71.3333333 | 78.1666667 | 46 | 55.8 | 14.3048382 | 4.52358702 |
| 5 | 45.5 | 54 | 39.5 | 46.5 | 42 | 54.5 | 46.6666667 | 82.3333333 | 59.6666667 | 51.1666667 | 52.1833333 | 12.2324326 | 3.86823484 |
| 10 | 47.2 | 77.1666667 | 62.7333333 | 76.0333333 | 48.2666667 | 68.4333333 | 70.8333333 | 86.3333333 | 71.2 | 63.1666667 | 67.1366667 | 12.332517 | 3.8998843 |
| 15 | 50.7 | 87.7 | 95.1333333 | 77.5 | 83.4333333 | 84.9333333 | 91.2 | 73.7 | 79.3666667 | 82.9333333 | 80.66 | 12.2879927 | 3.88580449 |

Het — Time (min) — Week 3

| Time (min) | B | D | E | H | 155 | 157 | 190 | 196 | 199 | 201 | 205 | 207 | 208 | 210 | Average | SD | SE |
|---|---|---|---|---|---|---|---|---|---|---|---|---|---|---|---|---|---|
| 0 | 76.3111111 | 91.2555556 | 83.9 | 85.7222222 | 82.8777778 | 84.6333333 | 85.7777778 | 89.5888889 | 88 | 87.9444444 | 81.0777778 | 85.6888889 | 84.9666667 | 92.4666667 | 85.7293651 | 4.15557294 | 1.11062358 |
| 1 | 60.2 | 61.6666667 | 66.8 | 50.1666667 | 70.6666667 | 72.3333333 | 67.1666667 | 57.8333333 | 67.5 | 79 | 69.1666667 | 59.8333333 | 62.6666667 | 52.5 | 64.1071429 | 7.82600507 | 2.09158783 |
| 2 | 50 | 55.3333333 | 61.8333333 | 46.5 | 74.3333333 | 77 | 48 | 44.1666667 | 68.8333333 | 87.6666667 | 74.1666667 | 51.6666667 | 55.8333333 | 45.5 | 60.0595238 | 13.9716757 | 3.73408741 |
| 3 | 42.8333333 | 41.1666667 | 70.5 | 46 | 79.1666667 | 81 | 44 | 39.5 | 63.1666667 | 49 | 67.3333333 | 41.3333333 | 65.5 | 39.6666667 | 55.0119048 | 15.3644423 | 4.10631993 |
| 4 | 39.8333333 | 44.1666667 | 52.8333333 | 51 | 79.5 | 49 | 47.1666667 | 39 | 53.5 | 47 | 66.8333333 | 50.6666667 | 67.6666667 | 48.8333333 | 52.6428571 | 11.3253851 | 3.02683648 |
| 5 | 61.6666667 | 42.5 | 43.5 | 46.3333333 | 82.1666667 | 48.5 | 54.1666667 | 42.1666667 | 55.5 | 48.5 | 67.3333333 | 42.6666667 | 49.3333333 | 76 | 54.3095238 | 12.9105829 | 3.45049841 |
| 10 | 50.4666667 | 60.5333333 | 60.6666667 | 74.3666667 | 63.7333333 | 67.9333333 | 71.4 | 59.6333333 | 56.5 | 79.1666667 | 54.7 | 65.4666667 | 49.2 | 77.5 | 63.6619048 | 9.51810562 | 2.54382073 |
| 15 | 57.1666667 | 93.0333333 | 75.0666667 | 80.5 | 69.1333333 | 58.4333333 | 82.6 | 64.3666667 | 71.3333333 | 65.9 | 49.7666667 | 82.9 | 84.7 | 95.7666667 | 73.6190476 | 13.7203244 | 3.66691094 |

Week 4

WT — Time (min) — Week 4

| Time (min) | 83 | 86 | 87 | 88 | 81 | 82 | A | 156 | 158 | 161 | Average | SD | SE |
|---|---|---|---|---|---|---|---|---|---|---|---|---|---|
| 0 | 49.6111111 | 78.5777778 | 67.1222222 | 60.5777778 | 93.2333333 | 68.0555556 | 62.0222222 | 69.6111111 | 85.3333333 | 58.7111111 | 69.2855556 | 13.1282375 | 4.1515132 |
| 1 | 55.1666667 | 50.1666667 | 55 | 51.1666667 | 58.5 | 50.8333333 | 46 | 48.3333333 | 57.1666667 | 54.3333333 | 52.6666667 | 3.99768452 | 1.26417884 |
| 2 | 47.3333333 | 56 | 45.8333333 | 48.5 | 59.3333333 | 56.5 | 43.8333333 | 66.6666667 | 82.1666667 | 63.1666667 | 56.9333333 | 11.7152955 | 3.70470172 |
| 3 | 44.8333333 | 51.1666667 | 44.1666667 | 44.1666667 | 57.8333333 | 56.8333333 | 39.5 | 55.3333333 | 71.5 | 50.1666667 | 51.55 | 9.31613826 | 2.94602159 |
| 4 | 44.5 | 51.3333333 | 49 | 48.1666667 | 58.1666667 | 52 | 40.5 | 48.8333333 | 61 | 46.3333333 | 49.9833333 | 6.08400554 | 1.92393148 |
| 5 | 47.3333333 | 50.8333333 | 50.1666667 | 48.6666667 | 49.1666667 | 51.8333333 | 40.6666667 | 49.6666667 | 53 | 46.6666667 | 48.8 | 3.44157587 | 1.08832185 |
| 10 | 61.7 | 64.2333333 | 59.7333333 | 62.7333333 | 56.4 | 68.2 | 70.6666667 | 56.2 | 62.6 | 55.0666667 | 61.7533333 | 5.13297738 | 1.62318997 |
| 15 | 79.8666667 | 90.3333333 | 83.3 | 94.5666667 | 76.2333333 | 87.7666667 | 88.2 | 94.6 | 79.2666667 | 84.7 | 85.8833333 | 6.33573541 | 2.00353546 |

Het — Time (min) — Week 4

| Time (min) | B | E | H | 155 | 157 | 190 | 196 | 199 | 201 | 205 | 207 | 208 | 210 | Average | SD | SE |
|---|---|---|---|---|---|---|---|---|---|---|---|---|---|---|---|---|
| 0 | 79.8111111 | 84.0666667 | 90.5222222 | 83.6444444 | 96.7111111 | 69.3777778 | 95.6777778 | 68.8333333 | 67.2777778 | 81.2 | 53.5111111 | 60.5555556 | 65.9 | 76.6991453 | 13.5504054 | 3.75820627 |
| 1 | 62.6666667 | 63.5 | 52.1666667 | 68.8333333 | 75.3333333 | 63.8333333 | 55.5 | 69.1666667 | 71.5 | 68 | 66 | 72.6666667 | 63 | 65.5512821 | 6.5441466 | 1.8150197 |
| 2 | 52.1666667 | 64.1666667 | 43.5 | 54.1666667 | 60.8333333 | 56 | 43.5 | 72.3333333 | 82.5 | 60.1666667 | 46 | 66 | 51.1666667 | 57.8846154 | 11.4898072 | 3.18669915 |
| 3 | 44 | 53.3333333 | 43.3333333 | 43.8333333 | 51.8333333 | 39.1666667 | 37.1666667 | 65.5 | 75.8333333 | 41.1666667 | 41.3333333 | 58.6666667 | 56 | 50.0897436 | 11.4991174 | 3.18928134 |
| 4 | 44.6666667 | 50.1666667 | 41.1666667 | 49.8333333 | 57.5 | 48.3333333 | 36.5 | 42.3333333 | 47.3333333 | 41 | 41.1666667 | 57.3333333 | 40.8333333 | 46.0128205 | 6.47983394 | 1.79718258 |
| 5 | 45.8333333 | 64.6666667 | 46.6666667 | 69.3333333 | 46.3333333 | 62.8333333 | 46.5 | 43.5 | 45 | 45.6666667 | 44.1666667 | 43 | 52.5 | 50.4615385 | 9.0382812 | 2.50676818 |
| 10 | 58.7333333 | 74.6 | 63.8333333 | 83.0666667 | 67.2 | 61.1333333 | 49.1333333 | 53.7 | 49.0333333 | 60.4666667 | 81.1333333 | 56.4333333 | 87.3333333 | 65.0615385 | 12.8007656 | 3.55029361 |
| 15 | 60.9 | 67.8333333 | 96.1 | 82.3333333 | 92.3666667 | 87.6666667 | 47.2 | 87.2 | 56.5 | 61.2 | 93.7 | 77 | 94.3333333 | 77.2564103 | 16.6504609 | 4.61800697 |

### Time of expiration

Week 1

WT

| Time (min) | Animal ID | | | | | | | | | | | | |
|---|---|---|---|---|---|---|---|---|---|---|---|---|---|
| | **83** | **86** | **87** | **88** | **81** | **82** | **A** | **156** | **158** | **161** | Average | SD | SE |
| 0 | 113.488889 | 117.922222 | 130.155556 | 120.822222 | 130.544444 | 125.977778 | 142.777778 | 125.211111 | 149.455556 | 143.688889 | 130.004444 | 11.8852345 | 3.75844114 |
| 1 | 95.8333333 | 81.3333333 | 98.3333333 | 90.1666667 | 101.6 | 87.3333333 | 135.333333 | 87.2 | 133.2 | 102.4 | 101.273333 | 18.655809 | 5.89948481 |
| 2 | 95.1666667 | 95 | 99.1666667 | 114.166667 | 104.333333 | 103.166667 | 110.833333 | 84 | 143 | 96.1666667 | 104.5 | 16.0316046 | 5.0696385 |
| 3 | 87 | 100.5 | 96.1666667 | 123.5 | 110.5 | 101 | 72.8333333 | 101.833333 | 111.166667 | 95.3333333 | 99.9833333 | 13.8856552 | 4.39102972 |
| 4 | 96.3333333 | 93.1666667 | 85.5 | 97.3333333 | 124.5 | 81.5454545 | 95.8333333 | 65.5 | 100.666667 | 97.3333333 | 93.7712121 | 15.0757676 | 4.76737632 |
| 5 | 104 | 89 | 115.666667 | 118.833333 | 130.666667 | 99.6666667 | 124.666667 | 103.666667 | 108.333333 | 104.5 | 109.9 | 12.5130056 | 3.9569598 |
| 10 | 136.2 | 118.066667 | 133.433333 | 136.333333 | 132.9 | 137.6 | 136.433333 | 117.533333 | 116.866667 | 124.566667 | 128.993333 | 8.74065003 | 2.76403623 |
| 15 | 128.333333 | 107.633333 | 122.6 | 142.4 | 129.433333 | 140.555556 | 166 | 113.233333 | 155.466667 | 133.433333 | 133.908889 | 17.9752061 | 5.68425926 |

Het

| Time (min) | Animal ID | | | | | | | | | | | | | | | | | | | | | |
|---|---|---|---|---|---|---|---|---|---|---|---|---|---|---|---|---|---|---|---|---|---|---|
| | **B** | **D** | **E** | **F** | **G** | **H** | **155** | **157** | **159** | **160** | **162** | **190** | **196** | **199** | **201** | **205** | **207** | **208** | **210** | Average | SD | SE |
| 0 | 170.455556 | 134.644444 | 142.422222 | 252.855556 | 165.866667 | 135.3 | 105.388889 | 100.444444 | 181.888889 | 115.9 | 135.122222 | 84.3111111 | 170.611111 | 172.544444 | 133.3 | 92.7333333 | 154.133333 | 139.4 | 195.088889 | 143.740123 | 40.369264 | 9.26134434 |
| 1 | 116.833333 | 80.2 | 101.666667 | 136.6 | 87.8 | 121 | 86.8 | 80.3333333 | 117.833333 | 133.8 | 94.2 | 109 | 90.1666667 | 106.5 | 101.333333 | 89.5 | 121.333333 | 109.333333 | 141.666667 | 104.67963 | 18.9131889 | 4.33898311 |
| 2 | 125.166667 | 98.3333333 | 131.333333 | 124.5 | 97 | 100.166667 | 107.333333 | 136.5 | 91 | 100.666667 | 116 | 92.5 | 96.8333333 | 134.666667 | 92.1666667 | 109.5 | 101 | 104.166667 | 116.166667 | 108.824074 | 14.9781572 | 3.43622492 |
| 3 | 131.5 | 93 | 162 | 232.5 | 109.5 | 105.166667 | 96.1666667 | 127.5 | 117.5 | 111.333333 | 133.666667 | 86.5 | 105.666667 | 145.333333 | 105 | 111.5 | 102.5 | 120 | 137.333333 | 122.018519 | 32.6819916 | 7.49776308 |
| 4 | 129.833333 | 72.1666667 | 187.833333 | 272.5 | 113 | 103.833333 | 88.6666667 | 138.833333 | 127.833333 | 120 | 130.5 | 76.6666667 | 106.5 | 105.833333 | 100.333333 | 113.166667 | 89 | 113.833333 | 124.666667 | 121.685185 | 44.5416809 | 10.2185624 |
| 5 | 160.666667 | 76.5 | 191.5 | 285.333333 | 106.333333 | 111 | 110.166667 | 155.833333 | 110.833333 | 94.6666667 | 130.5 | 78.3333333 | 97.1666667 | 92.1666667 | 73.3333333 | 97.5 | 94.8333333 | 106.5 | 122.833333 | 120.731481 | 50.0023911 | 11.4713353 |
| 10 | 112.1 | 86.7333333 | 146.366667 | 218.033333 | 116.033333 | 124.066667 | 119.9 | 138.833333 | 121.5 | 99 | 128.5 | 85.0333333 | 163.833333 | 79.3333333 | 96.3666667 | 114.733333 | 123.033333 | 104.566667 | 148.166667 | 120.998148 | 32.2232725 | 7.39252571 |
| 15 | 135.266667 | 98.8333333 | 137.333333 | 148.8 | 120.566667 | 135.366667 | 149.866667 | 140.233333 | 152 | 120.566667 | 183.9 | 100.466667 | 164.066667 | 98.5333333 | 132.366667 | 146.466667 | 150.3 | 136.4 | 153.833333 | 136.185185 | 22.1871649 | 5.09008472 |

Week 2

WT

| Time (min) | Animal ID | | | | | | | | | | | | |
|---|---|---|---|---|---|---|---|---|---|---|---|---|---|
| | **83** | **86** | **87** | **88** | **81** | **82** | **A** | **156** | **158** | **161** | Average | SD | SE |
| 0 | 102.311111 | 106.888889 | 122.466667 | 104.544444 | 119.788889 | 149.677778 | 128.744444 | 154.177778 | 109.688889 | 126.966667 | 122.525556 | 18.0930517 | 5.72152532 |
| 1 | 92.8333333 | 80 | 79.3333333 | 79.6666667 | 79.5 | 80.6666667 | 92.3333333 | 118.833333 | 121.833333 | 92.8333333 | 91.7833333 | 16.1669626 | 5.11244246 |
| 2 | 87.3333333 | 82.1666667 | 82 | 88 | 82.8333333 | 78.6666667 | 89.1666667 | 88 | 130.666667 | 89.8333333 | 89.8666667 | 14.8121572 | 4.68401537 |
| 3 | 92.3333333 | 81.3333333 | 88.3333333 | 95.8333333 | 88.5 | 102 | 96.6666667 | 97.8333333 | 116.5 | 95.6666667 | 95.5 | 9.43561679 | 2.98380402 |
| 4 | 92.8333333 | 84.8333333 | 87 | 122.5 | 88.6666667 | 93.8333333 | 97 | 109.666667 | 109 | 68.3333333 | 95.3666667 | 15.2558024 | 4.8243083 |
| 5 | 85.8333333 | 82.8333333 | 106.166667 | 108.833333 | 97.3333333 | 110.666667 | 120 | 139.166667 | 126.333333 | 90 | 106.716667 | 18.2837798 | 5.78183885 |
| 10 | 115.966667 | 105 | 137.9 | 103.2 | 144.966667 | 170.566667 | 130.7 | 147.466667 | 146.333333 | 123.833333 | 132.593333 | 21.1223058 | 6.67945958 |
| 15 | 122.566667 | 138.833333 | 137.833333 | 125.133333 | 151.633333 | 163.366667 | 129 | 155.7 | 125.433333 | 138.533333 | 138.803333 | 14.0627118 | 4.44701995 |

Het

| Time (min) | Animal ID | | | | | | | | | | | | | | | | | | | |
|---|---|---|---|---|---|---|---|---|---|---|---|---|---|---|---|---|---|---|---|---|
| | **B** | **D** | **E** | **G** | **H** | **155** | **157** | **160** | **162** | **190** | **196** | **199** | **201** | **205** | **207** | **208** | **210** | Average | SD | SE |
| 0 | 115.744444 | 152.566667 | 144.622222 | 129.188889 | 135.9 | 134.955556 | 115.188889 | 139.6 | 214.977778 | 134.933333 | 197.533333 | 198.477778 | 156.577778 | 107.933333 | 148.188889 | 89.3222222 | 148.455556 | 144.95098 | 33.0839432 | 8.02403485 |
| 1 | 88.6666667 | 96 | 81 | 102 | 76 | 88.5 | 126.166667 | 109.666667 | 103.166667 | 82 | 97.6666667 | 116 | 109.5 | 87.6666667 | 86.1666667 | 106.166667 | 78.3333333 | 96.1568627 | 14.3209092 | 3.47333066 |
| 2 | 99.5 | 89.5 | 92.5 | 110.166667 | 91.3333333 | 118.333333 | 115.833333 | 97.1666667 | 104 | 79.1666667 | 87.6666667 | 107.5 | 85.8333333 | 103.166667 | 77.5 | 92.6666667 | 85.5 | 96.3137255 | 12.115171 | 2.93836058 |
| 3 | 121.166667 | 102.166667 | 103 | 105.833333 | 93.1666667 | 132.333333 | 146.166667 | 108.666667 | 115 | 69 | 69.1666667 | 112.833333 | 70 | 95.5 | 68.1666667 | 113 | 89.8333333 | 100.882353 | 22.7602819 | 5.5201792 |
| 4 | 92.1666667 | 100.833333 | 96.8333333 | 101.833333 | 103.833333 | 134.666667 | 169.5 | 108.5 | 152.833333 | 81 | 90.5 | 83.8333333 | 71.8333333 | 102 | 85.5 | 102 | 84 | 103.627451 | 25.9112755 | 6.28440741 |
| 5 | 80 | 87.3333333 | 112.666667 | 100.333333 | 130.333333 | 132.833333 | 169 | 117.333333 | 193.333333 | 87.6666667 | 108.333333 | 103 | 94.6666667 | 108.333333 | 85.5 | 84.6666667 | 108.166667 | 111.970588 | 30.5124767 | 7.40036261 |
| 10 | 148.2 | 100.533333 | 119.333333 | 120.3 | 127.7 | 135.833333 | 138.9 | 179.133333 | 213.266667 | 114.366667 | 190.066667 | 124.2 | 175.5 | 126.7 | 142 | 105.033333 | 117.166667 | 139.896078 | 31.7290348 | 7.69542129 |
| 15 | 127.333333 | 118.833333 | 143.8 | 112.166667 | 132.5 | 133.033333 | 154.166667 | 281.8 | 347.233333 | 129 | 110.6 | 129.066667 | 135.666667 | 156 | 175.266667 | 120.3 | 133.866667 | 155.331373 | 63.1512244 | 15.3164217 |

Week 3

WT

| Time (min) | Animal ID | | | | | | | | | | | | |
|---|---|---|---|---|---|---|---|---|---|---|---|---|---|
| | **83** | **86** | **87** | **88** | **81** | **82** | **A** | **156** | **158** | **161** | Average | SD | SE |
| 0 | 111.911111 | 117.166667 | 153.311111 | 137.6 | 155.3 | 150.044444 | 114.255556 | 149.855556 | 156.877778 | 131.922222 | 137.824444 | 17.9225531 | 5.66760893 |
| 1 | 72.1666667 | 65.6666667 | 64.1666667 | 67.8333333 | 70.5 | 66.8333333 | 73.5 | 83.5 | 131.666667 | 91.6666667 | 78.75 | 20.4764619 | 6.47522582 |
| 2 | 66.5 | 77.5 | 70.5 | 76.3333333 | 80.3333333 | 109.5 | 78.5 | 107.833333 | 176.833333 | 97.6666667 | 94.15 | 32.7041971 | 10.3419752 |
| 3 | 66.6666667 | 95.3333333 | 56.8333333 | 70.1666667 | 80 | 111.333333 | 81.8333333 | 125.333333 | 171.166667 | 88.5 | 94.7166667 | 33.8367586 | 10.7001226 |
| 4 | 94 | 97.6666667 | 73.5 | 61.3333333 | 67.5 | 103.166667 | 74.6666667 | 135.166667 | 175.833333 | 87.6666667 | 97.05 | 34.9592399 | 11.0550823 |
| 5 | 87 | 102.833333 | 81.6666667 | 90.3333333 | 70.5 | 101 | 84.3333333 | 156 | 126.833333 | 118.666667 | 101.916667 | 25.5843135 | 8.09047031 |
| 10 | 87.1 | 147.233333 | 128.833333 | 162.433333 | 106.066667 | 147.966667 | 132.533333 | 153.9 | 143.533333 | 134.833333 | 134.443333 | 22.7864322 | 7.20570256 |
| 15 | 106.2 | 154.7 | 162.8 | 153.833333 | 182.433333 | 164.7 | 152.233333 | 142.833333 | 153.4 | 160.3 | 153.343333 | 19.5769015 | 6.19075984 |

Het

| Time (min) | Animal ID | | | | | | | | | | | | | | | | |
|---|---|---|---|---|---|---|---|---|---|---|---|---|---|---|---|---|---|
| | **B** | **D** | **E** | **H** | **155** | **157** | **190** | **196** | **199** | **201** | **205** | **207** | **208** | **210** | Average | SD | SE |
| 0 | 137.966667 | 167.455556 | 158.511111 | 160.333333 | 146.666667 | 154.688889 | 134.566667 | 145.788889 | 183.677778 | 159.855556 | 127.533333 | 145.355556 | 145.433333 | 140.844444 | 150.619841 | 14.6094775 | 3.90454709 |
| 1 | 80.8 | 88 | 102.8 | 77.8333333 | 95.8333333 | 90.3333333 | 90.8333333 | 85.5 | 98.3333333 | 110.666667 | 108.833333 | 96 | 101.333333 | 82 | 93.5071429 | 10.2397598 | 2.73669091 |
| 2 | 74.3333333 | 87 | 105.666667 | 81.3333333 | 143 | 133.166667 | 77.1666667 | 78.1666667 | 115.833333 | 118.833333 | 110 | 89.5 | 109.666667 | 90.8333333 | 101.035714 | 21.7211523 | 5.80522213 |
| 3 | 67.3333333 | 64.3333333 | 119 | 79.8333333 | 157.5 | 144 | 71.8333333 | 69.5 | 127.166667 | 80 | 122.166667 | 66.8333333 | 144.833333 | 80.3333333 | 99.6190476 | 34.1906063 | 9.1378239 |
| 4 | 64.3333333 | 72.6666667 | 93.3333333 | 72.8333333 | 156.666667 | 81 | 79.3333333 | 74 | 94.1666667 | 76.3333333 | 131.333333 | 80.6666667 | 140.166667 | 100.166667 | 94.0714286 | 28.4847173 | 7.61286093 |
| 5 | 100.333333 | 76 | 80.8333333 | 82.1666667 | 149.833333 | 92 | 97.6666667 | 102.333333 | 107.5 | 89.8333333 | 145 | 89.3333333 | 108 | 122 | 103.059524 | 22.4209539 | 5.992252 |
| 10 | 109.533333 | 132.6 | 165.3 | 144.9 | 127.066667 | 136.2 | 132.5 | 147.9 | 115.6 | 108.833333 | 136.2 | 140.266667 | 122.5 | 150.8 | 133.585714 | 16.1139411 | 4.3066319 |
| 15 | 139.933333 | 196.066667 | 202.566667 | 155.1 | 163.5 | 120.466667 | 153.466667 | 161.7 | 197.066667 | 117.833333 | 140.7 | 157.966667 | 175.133333 | 162.2 | 160.264286 | 26.1968663 | 7.00140703 |

Week 4

WT

| Time (min) | Animal ID | | | | | | | | | | | | |
|---|---|---|---|---|---|---|---|---|---|---|---|---|---|
| | **83** | **86** | **87** | **88** | **81** | **82** | **A** | **156** | **158** | **161** | Average | SD | SE |
| 0 | 105.433333 | 163.533333 | 132.277778 | 118.744444 | 178.677778 | 136.555556 | 144.522222 | 147.933333 | 172.5 | 146.877778 | 144.705556 | 22.9333765 | 7.25217043 |
| 1 | 79.8333333 | 65.8333333 | 77.1666667 | 69.5 | 79.3333333 | 74.5 | 68.1666667 | 66.5 | 113.333333 | 81.8333333 | 77.6 | 13.8597917 | 4.38285098 |
| 2 | 84.6666667 | 87.5 | 75.1666667 | 79.3333333 | 90.8333333 | 94.1666667 | 81.5 | 92.6666667 | 161 | 98 | 94.4833333 | 24.4265654 | 7.72435822 |
| 3 | 78.1666667 | 83.8333333 | 69.5 | 70 | 94.3333333 | 102.5 | 68.8333333 | 90.3333333 | 128.166667 | 80.5 | 86.6166667 | 18.4208433 | 5.82518214 |
| 4 | 78.5 | 78 | 76.8333333 | 78.1666667 | 104.333333 | 101.166667 | 73.1666667 | 86.3333333 | 126.333333 | 86.1666667 | 88.9 | 16.7971191 | 5.31171545 |
| 5 | 87 | 76 | 89.3333333 | 89.5 | 90.6666667 | 98.5 | 80.3333333 | 96.3333333 | 124 | 104 | 93.5666667 | 13.4855204 | 4.26449598 |
| 10 | 136.1 | 122.566667 | 122.033333 | 125.5 | 118.266667 | 137.833333 | 125.833333 | 122.966667 | 143.3 | 124.366667 | 127.876667 | 8.20286084 | 2.59397236 |
| 15 | 160.2 | 152.566667 | 148.066667 | 157.5 | 168 | 152.566667 | 152.033333 | 193.733333 | 154.566667 | 169.5 | 160.873333 | 13.4856009 | 4.26452145 |

Het

| Time (min) | Animal ID | | | | | | | | | | | | | | | |
|---|---|---|---|---|---|---|---|---|---|---|---|---|---|---|---|---|
| | **B** | **E** | **H** | **155** | **157** | **190** | **196** | **199** | **201** | **205** | **207** | **208** | **210** | Average | SD | SE |
| 0 | 157.555556 | 144.322222 | 176.366667 | 142.744444 | 179.977778 | 128.4 | 166.055556 | 128.766667 | 127.933333 | 143.166667 | 115.1 | 124.277778 | 131.111111 | 143.521368 | 20.6870541 | 5.73755648 |
| 1 | 83 | 96 | 82 | 96.1666667 | 90.6666667 | 87.6666667 | 82.5 | 102 | 124.333333 | 98.5 | 105.166667 | 99.5 | 93.6666667 | 95.474359 | 11.5047989 | 3.19085712 |
| 2 | 80.1666667 | 110.333333 | 78.3333333 | 104.5 | 103.333333 | 101.166667 | 75 | 124.333333 | 97.8333333 | 106.5 | 83.5 | 112.166667 | 104 | 98.5512821 | 14.9313839 | 4.14122079 |
| 3 | 61.1666667 | 92.3333333 | 76.6666667 | 91.8333333 | 87.3333333 | 65.5 | 63.1666667 | 139.5 | 101.333333 | 72.1666667 | 72 | 109.5 | 116.666667 | 88.3974359 | 23.4947333 | 6.51626659 |
| 4 | 68 | 97.5 | 76 | 108 | 99 | 82 | 64.6666667 | 69.6666667 | 71.8333333 | 72.5 | 68.5 | 112.333333 | 74 | 81.8461538 | 16.455673 | 4.56398252 |
| 5 | 78.6666667 | 172.333333 | 93.1666667 | 187.333333 | 81 | 115.666667 | 86.3333333 | 78 | 72.6666667 | 98 | 82.6666667 | 84.3333333 | 113.833333 | 103.384615 | 36.4893684 | 10.1203299 |
| 10 | 100.933333 | 163 | 147.833333 | 169.833333 | 145.033333 | 121.4 | 109 | 112.366667 | 91.2 | 148.066667 | 164.233333 | 148.9 | 195.333333 | 139.779487 | 30.6548121 | 8.50211515 |
| 15 | 97.9333333 | 141.666667 | 197.1 | 154.366667 | 206.833333 | 180.8 | 108.166667 | 197.033333 | 108.933333 | 99.6 | 172.6 | 171.758621 | 177.566667 | 154.950663 | 39.5925308 | 10.9809923 |

### Peak inspiratory flow

Week 1

WT — Time (min) / Animal ID

| Time (min) | 83 | 86 | 87 | 88 | 81 | 82 | A | 156 | 158 | 161 | Average | SD | SE |
|---|---|---|---|---|---|---|---|---|---|---|---|---|---|
| 0 | 4.10033333 | 3.46044444 | 2.86511111 | 4.512 | 2.79411111 | 2.54211111 | 3.26533333 | 4.01911111 | 1.92511111 | 1.88255556 | 3.13662222 | 0.90095845 | 0.28490808 |
| 1 | 3.13 | 4.39 | 3.50166667 | 5.34666667 | 3.424 | 3.24083333 | 1.36333333 | 3.892 | 2.036 | 2.266 | 3.25905 | 1.16507584 | 0.36842933 |
| 2 | 3.3 | 4.125 | 3.57666667 | 3.02 | 2.94166667 | 2.87 | 1.96 | 3.63 | 2.05166667 | 2.00666667 | 2.94816667 | 0.74881777 | 0.23679697 |
| 3 | 3.38333333 | 4.025 | 3.63666667 | 2.295 | 2.80833333 | 2.96666667 | 2.64833333 | 3.635 | 2.345 | 2.25 | 2.99933333 | 0.63737318 | 0.2015551 |
| 4 | 3.58166667 | 4.04333333 | 4.575 | 3.9 | 2.73333333 | 3.35545455 | 2.43166667 | 4.54333333 | 2.845 | 2.4 | 3.44087879 | 0.82057765 | 0.25948944 |
| 5 | 3.45833333 | 4.09833333 | 3.77333333 | 2.9 | 2.85833333 | 3.66 | 2.02833333 | 3.99333333 | 2.42666667 | 2.82 | 3.20166667 | 0.69644999 | 0.22023683 |
| 10 | 2.40266667 | 3.61366667 | 2.64466667 | 2.94466667 | 3.09066667 | 3.27133333 | 1.70666667 | 3.45866667 | 2.834 | 2.42766667 | 2.83946667 | 0.57009335 | 0.18027935 |
| 15 | 2.183 | 3.212 | 2.58833333 | 2.495 | 3.14866667 | 2.67111111 | 1.277 | 3.49533333 | 1.832 | 1.862 | 2.47644444 | 0.6991618 | 0.22109437 |

Het — Time (min) / Animal ID

| Time (min) | B | D | E | F | G | H | 155 | 157 | 159 | 160 | 162 | 190 | 196 | 199 | 201 | 205 | 207 | 208 | 210 | Average | SD | SE |
|---|---|---|---|---|---|---|---|---|---|---|---|---|---|---|---|---|---|---|---|---|---|---|
| 0 | 2.06444444 | 2.18555556 | 3.08255556 | 1.46955556 | 2.00355556 | 3.31333333 | 3.28655556 | 3.36855556 | 2.06388889 | 3.42877778 | 2.89233333 | 3.57033333 | 2.06611111 | 2.23 | 3.31677778 | 2.74933333 | 2.28377778 | 1.15122222 | 1.28233333 | 2.51626316 | 0.7694605 | 0.17652634 |
| 1 | 3.34 | 3.728 | 3.43833333 | 1.428 | 1.948 | 2.782 | 3.06 | 2.52166667 | 3.365 | 2.718 | 2.568 | 3.25833333 | 2.88 | 3.68 | 3.41666667 | 1.925 | 2.525 | 2.67333333 | 1.84166667 | 2.79457895 | 0.65996608 | 0.1514066 |
| 2 | 1.97666667 | 3.18 | 2.14333333 | 1.37833333 | 1.55166667 | 2.965 | 2.845 | 2.785 | 1.62166667 | 1.71333333 | 1.96 | 2.095 | 2.76166667 | 3.5 | 3.39833333 | 2.38 | 1.56166667 | 1.105 | 1.32166667 | 2.22333333 | 0.74288726 | 0.17043003 |
| 3 | 1.66166667 | 3.56666667 | 1.56333333 | 1.07666667 | 1.34833333 | 2.73666667 | 3.35333333 | 2.47333333 | 1.605 | 1.475 | 1.93 | 2.55166667 | 3.045 | 3.27833333 | 3.04166667 | 1.97833333 | 1.78333333 | 1.04166667 | 1.17166667 | 2.14114035 | 0.83201953 | 0.19087837 |
| 4 | 1.80333333 | 3.885 | 1.42666667 | 0.975 | 1.36333333 | 1.94166667 | 3.515 | 2.00666667 | 1.56 | 2.145 | 1.89666667 | 3.46833333 | 3.395 | 3.59 | 3.68833333 | 2.285 | 2.03 | 1.08666667 | 1.35166667 | 2.28491228 | 0.9768279 | 0.22409969 |
| 5 | 1.56166667 | 3.76 | 1.57333333 | 3.32833333 | 1.53 | 2.02666667 | 3.12333333 | 1.90166667 | 2.11666667 | 2.73333333 | 1.80666667 | 3.98 | 3.6 | 3.90833333 | 4.24333333 | 2.78666667 | 2.04833333 | 1.26666667 | 1.21666667 | 2.55324561 | 1.01187097 | 0.23213912 |
| 10 | 2.462 | 3.51733333 | 1.83766667 | 0.99533333 | 1.71 | 2.786 | 3.18733333 | 2.257 | 1.93433333 | 3.08566667 | 2.70533333 | 3.78166667 | 2.78933333 | 3.641 | 3.46233333 | 2.71666667 | 2.06166667 | 1.51633333 | 1.27666667 | 2.51177193 | 0.82786802 | 0.18992595 |
| 15 | 2.34433333 | 3.234 | 2.146 | 1.15233333 | 2.10066667 | 2.79733333 | 1.888 | 2.53466667 | 1.803 | 2.95466667 | 2.19766667 | 3.268 | 2.57033333 | 3.57033333 | 3.14033333 | 1.965 | 1.434 | 1.33566667 | 1.33533333 | 2.30377193 | 0.72516568 | 0.16636442 |

Week 2

WT — Time (min) / Animal ID

| Time (min) | 83 | 86 | 87 | 88 | 81 | 82 | A | 156 | 158 | 161 | Average | SD | SE |
|---|---|---|---|---|---|---|---|---|---|---|---|---|---|
| 0 | 5.34077778 | 5.33122222 | 3.64188889 | 6.05922222 | 5.06277778 | 3.01744444 | 5.21055556 | 3.76277778 | 3.83388889 | 4.24866667 | 4.55092222 | 0.97805837 | 0.30928921 |
| 1 | 4.94333333 | 6.345 | 6.15166667 | 7.67333333 | 5.50666667 | 6.13333333 | 4.47 | 3.32833333 | 2.38666667 | 3.4 | 5.03383333 | 1.64248702 | 0.5194 |
| 2 | 4.17666667 | 6.59833333 | 5.87166667 | 8.50166667 | 5.01666667 | 6.545 | 3.39 | 5.31333333 | 2.705 | 3.64833333 | 5.17666667 | 1.76740603 | 0.55890286 |
| 3 | 4.65 | 6.675 | 6.18166667 | 8.17666667 | 4.84833333 | 6.21 | 3.20666667 | 4.015 | 2.91333333 | 3.93666667 | 5.08133333 | 1.68350656 | 0.53237152 |
| 4 | 3.49166667 | 6.775 | 6.53333333 | 7.63333333 | 5.50166667 | 5.84 | 4.04666667 | 3.85 | 3.12333333 | 4.96833333 | 5.17633333 | 1.53158632 | 0.48433012 |
| 5 | 4.88833333 | 6.63 | 5.49333333 | 6.5 | 4.96333333 | 6.03166667 | 3.705 | 3.055 | 3.175 | 4.53833333 | 4.898 | 1.29649951 | 0.40998914 |
| 10 | 4.48866667 | 5.93333333 | 5.008 | 6.50666667 | 4.12233333 | 4.589 | 3.5 | 3.168 | 2.655 | 3.41733333 | 4.33883333 | 1.22839418 | 0.38845235 |
| 15 | 4.63533333 | 4.81333333 | 3.79566667 | 5.866 | 3.338 | 4.022 | 3.83366667 | 2.905 | 3.372 | 2.435 | 3.9016 | 1.00005949 | 0.31624658 |

Het — Time (min) / Animal ID

| Time (min) | B | D | E | G | H | 155 | 157 | 160 | 162 | 190 | 196 | 199 | 201 | 205 | 207 | 208 | 210 | Average | SD | SE |
|---|---|---|---|---|---|---|---|---|---|---|---|---|---|---|---|---|---|---|---|---|
| 0 | 3.96522222 | 2.32777778 | 2.30311111 | 1.14688889 | 4.00155556 | 2.36244444 | 3.37755556 | 2.90911111 | 3.14433333 | 2.35655556 | 2.87244444 | 2.25411111 | 2.94833333 | 3.68044444 | 2.06455556 | 3.93388889 | 3.35844444 | 2.88275163 | 0.78972875 | 0.19153736 |
| 1 | 4.34666667 | 3.845 | 3.262 | 2.07166667 | 2.85833333 | 3.56833333 | 3.42666667 | 2.85 | 2.775 | 4.54333333 | 4.05666667 | 3.635 | 3.835 | 2.14333333 | 3.08666667 | 3.64166667 | 2.87 | 3.34207843 | 0.69981344 | 0.16972969 |
| 2 | 3.21833333 | 4.17166667 | 3.23166667 | 2.24833333 | 3.34166667 | 3.285 | 2.07 | 2.95833333 | 3.095 | 4.36666667 | 4.92166667 | 3.495 | 4.26166667 | 2.44166667 | 3.65166667 | 2.22166667 | 3.825 | 3.34147059 | 0.81204878 | 0.19695076 |
| 3 | 2.54 | 4.005 | 3.37833333 | 2.23666667 | 3.86333333 | 3.14166667 | 1.85666667 | 2.93666667 | 3.10333333 | 4.495 | 5.18333333 | 3.735 | 4.94833333 | 3.01 | 4.13 | 1.67833333 | 3.58166667 | 3.40137255 | 0.99988719 | 0.24250826 |
| 4 | 4.865 | 4.23166667 | 4.17666667 | 2.14666667 | 3.53 | 2.66666667 | 2.08 | 3.24166667 | 2.83833333 | 4.47166667 | 4.795 | 4.54833333 | 4.52 | 2.66666667 | 3.775 | 3.105 | 3.83 | 3.61696078 | 0.91744328 | 0.22251268 |
| 5 | 4.92666667 | 4.79166667 | 3.92833333 | 2.87333333 | 2.99166667 | 2.65333333 | 2.025 | 3.195 | 2.18333333 | 4.2 | 4.44 | 4.43833333 | 4.10333333 | 2.49 | 3.485 | 3.90333333 | 3.06666667 | 3.51147059 | 0.90981089 | 0.22066155 |
| 10 | 2.96666667 | 4.41766667 | 3.58366667 | 2.49233333 | 2.81033333 | 2.51033333 | 2.59266667 | 2.85433333 | 2.20833333 | 3.71833333 | 3.44133333 | 3.82566667 | 3.34333333 | 2.472 | 2.95966667 | 3.37666667 | 3.11033333 | 3.09903922 | 0.5850747 | 0.14190146 |
| 15 | 3.53066667 | 4.025 | 2.873 | 1.55366667 | 2.60833333 | 3.41166667 | 2.80066667 | 2.33033333 | 1.409 | 3.63966667 | 3.57066667 | 3.62166667 | 3.64366667 | 2.63633333 | 1.83133333 | 2.85433333 | 2.724 | 2.88611765 | 0.77834542 | 0.18877649 |

Week 3

WT — Time (min) / Animal ID

| Time (min) | 83 | 86 | 87 | 88 | 81 | 82 | A | 156 | 158 | 161 | Average | SD | SE |
|---|---|---|---|---|---|---|---|---|---|---|---|---|---|
| 0 | 6.334 | 5.63255556 | 5.25855556 | 5.12711111 | 3.26255556 | 3.11611111 | 5.45177778 | 4.17577778 | 2.189 | 3.95022222 | 4.44976667 | 1.3203566 | 0.41753342 |
| 1 | 5.45666667 | 5.89666667 | 7.205 | 6.32333333 | 6.45333333 | 5.06333333 | 5.31166667 | 4.995 | 2.71333333 | 2.965 | 5.23833333 | 1.43918122 | 0.45510906 |
| 2 | 6.02 | 4.91333333 | 6.41166667 | 6.18666667 | 6.19166667 | 3.76833333 | 4.23833333 | 3.08666667 | 2.055 | 2.73833333 | 4.561 | 1.61547707 | 0.51085871 |
| 3 | 6.43166667 | 4.91166667 | 7.15833333 | 7.15166667 | 6.98333333 | 4.05 | 4.92333333 | 3.32166667 | 2.07 | 4.32666667 | 5.13283333 | 1.75961381 | 0.55643875 |
| 4 | 5.58166667 | 4.28666667 | 6.545 | 7.04833333 | 7.73666667 | 3.88 | 5.175 | 3.72833333 | 2.23333333 | 4.93333333 | 5.11483333 | 1.6782399 | 0.53070605 |
| 5 | 5.70166667 | 5.16333333 | 6.33833333 | 6.55666667 | 6.45333333 | 4.83333333 | 5.715 | 3.52833333 | 2.89 | 4.37333333 | 5.15533333 | 1.25405436 | 0.39656681 |
| 10 | 5.39733333 | 4.29733333 | 4.89833333 | 4.22833333 | 5.98766667 | 4.54466667 | 4.108 | 3.995 | 2.937 | 3.65333333 | 4.4047 | 0.86740878 | 0.27429874 |
| 15 | 5.51233333 | 3.36466667 | 2.958 | 4.034 | 3.91366667 | 3.48966667 | 2.712 | 4.52833333 | 2.52066667 | 2.31533333 | 3.53486667 | 0.9914486 | 0.31352358 |

Het — Time (min) / Animal ID

| Time (min) | B | D | E | H | 155 | 157 | 190 | 196 | 199 | 201 | 205 | 207 | 208 | 210 | Average | SD | SE |
|---|---|---|---|---|---|---|---|---|---|---|---|---|---|---|---|---|---|
| 0 | 3.31644444 | 2.68266667 | 1.91788889 | 2.61811111 | 2.51755556 | 2.74377778 | 2.406 | 2.447 | 2.88166667 | 2.80777778 | 1.622 | 1.76266667 | 1.69733333 | 1.64022222 | 2.36150794 | 0.54023107 | 0.14438283 |
| 1 | 5.2 | 4.64166667 | 3.262 | 4.42166667 | 3.80333333 | 3.46666667 | 3.97 | 5.33166667 | 4.40833333 | 4.09833333 | 2.80166667 | 2.995 | 2.50333333 | 3.49833333 | 3.88585714 | 0.85948351 | 0.22970663 |
| 2 | 5.835 | 4.615 | 3.485 | 3.95666667 | 2.96333333 | 3.30833333 | 4.30833333 | 5.57333333 | 4.15 | 2.98666667 | 2.07666667 | 3.30333333 | 3.06666667 | 3.59333333 | 3.80154762 | 1.03236482 | 0.2759111 |
| 3 | 6.275 | 5.475 | 3.3 | 4.735 | 2.63666667 | 2.965 | 4.785 | 6.05 | 4.00666667 | 4.84833333 | 2.59333333 | 4.00833333 | 2.71166667 | 3.7 | 4.14928571 | 1.24858051 | 0.33369718 |
| 4 | 6.34833333 | 5.315 | 5.05333333 | 4.63 | 2.62833333 | 4.28166667 | 4.92 | 5.76 | 4.46333333 | 5.24833333 | 2.54 | 3.535 | 2.55166667 | 3.65333333 | 4.35202381 | 1.21715009 | 0.32529705 |
| 5 | 4.39166667 | 4.87333333 | 6.105 | 4.49 | 2.42166667 | 4.36333333 | 4.25833333 | 4.605 | 4.42 | 4.695 | 2.52666667 | 3.39333333 | 3.68 | 2.42333333 | 4.04619048 | 1.05245997 | 0.28128176 |
| 10 | 5.37366667 | 4.02866667 | 3.777 | 3.39133333 | 3.879 | 3.74833333 | 3.43566667 | 4.17033333 | 4.42966667 | 3.61466667 | 3.37366667 | 2.65466667 | 3.725 | 2.60433333 | 3.729 | 0.69415669 | 0.18552118 |
| 15 | 4.908 | 2.65033333 | 2.894 | 3.036 | 3.62833333 | 4.11933333 | 3.026 | 4.441 | 3.968 | 4.05033333 | 3.62566667 | 2.007 | 2.028 | 1.97433333 | 3.31116667 | 0.9429533 | 0.25201487 |

Week 4

WT — Time (min) / Animal ID

| Time (min) | 83 | 86 | 87 | 88 | 81 | 82 | A | 156 | 158 | 161 | Average | SD | SE |
|---|---|---|---|---|---|---|---|---|---|---|---|---|---|
| 0 | 6.85611111 | 4.24966667 | 4.99122222 | 5.88077778 | 3.14411111 | 4.27555556 | 5.73888889 | 5.60466667 | 2.39577778 | 4.48755556 | 4.76243333 | 1.27062624 | 0.4018073 |
| 1 | 5.94833333 | 6.06833333 | 7.015 | 7.305 | 5.52333333 | 5.00333333 | 6.04833333 | 5.885 | 3.62 | 3.80833333 | 5.6225 | 1.14102786 | 0.36082469 |
| 2 | 5.20833333 | 5.51666667 | 7.325 | 7.07333333 | 5.01666667 | 4.34666667 | 5.865 | 4.48833333 | 1.87333333 | 2.99833333 | 4.97116667 | 1.58827685 | 0.50225724 |
| 3 | 5.785 | 6.02666667 | 8.04833333 | 8.21166667 | 5.025 | 4.67166667 | 6.34833333 | 5.33833333 | 2.40833333 | 4.425 | 5.62883333 | 1.62515709 | 0.5139198 |
| 4 | 6.37 | 7.14 | 7.92833333 | 8.085 | 5.94 | 4.88166667 | 6.315 | 6.08666667 | 3.195 | 5.5 | 6.14416667 | 1.37016002 | 0.43328264 |
| 5 | 6.15 | 6.48666667 | 6.91333333 | 7.26333333 | 6.92833333 | 5.21833333 | 5.73333333 | 6.34 | 3.68 | 4.51 | 5.92233333 | 1.09046911 | 0.34483661 |
| 10 | 5.09966667 | 5.06533333 | 5.547 | 5.39833333 | 5.693 | 4.444 | 4.37333333 | 6.10833333 | 3.591 | 4.20066667 | 4.95206667 | 0.74052805 | 0.23417553 |
| 15 | 3.85166667 | 3.05433333 | 3.972 | 3.024 | 4.51833333 | 3.367 | 3.55033333 | 3.413 | 2.56333333 | 2.61733333 | 3.39313333 | 0.58126435 | 0.18381193 |

Het — Time (min) / Animal ID

| Time (min) | B | E | H | 155 | 157 | 190 | 196 | 199 | 201 | 205 | 207 | 208 | 210 | Average | SD | SE |
|---|---|---|---|---|---|---|---|---|---|---|---|---|---|---|---|---|
| 0 | 3.81144444 | 2.05811111 | 2.669 | 2.83711111 | 2.97722222 | 3.65011111 | 2.58133333 | 3.59377778 | 3.78888889 | 2.15233333 | 3.37833333 | 3.09444444 | 3.04733333 | 3.04918803 | 0.58426845 | 0.16204691 |
| 1 | 6.12166667 | 3.83333333 | 4.49333333 | 4.12333333 | 3.21333333 | 4.445 | 4.83333333 | 4.26333333 | 4.735 | 3.16 | 2.83833333 | 2.57333333 | 2.895 | 3.96371795 | 1.00842611 | 0.27968708 |
| 2 | 5.72666667 | 3.87833333 | 4.89666667 | 3.75 | 3.19166667 | 4 | 5.575 | 3.58 | 3.545 | 3.12833333 | 3.455 | 2.61833333 | 3.52666667 | 3.91320513 | 0.93417013 | 0.25909218 |
| 3 | 7.59 | 5.55666667 | 5.54166667 | 3.77833333 | 3.70666667 | 4.87833333 | 5.85333333 | 3.84833333 | 3.53 | 4.595 | 4.02166667 | 3.27 | 3.47 | 4.58769231 | 1.25670766 | 0.34854799 |
| 4 | 7.395 | 5.07 | 5.10666667 | 4.07 | 3.83166667 | 4.55333333 | 5.93333333 | 5.50833333 | 5.865 | 5.08833333 | 4.43333333 | 3.185 | 4.71166667 | 4.98089744 | 1.07086362 | 0.29700413 |
| 5 | 7.05833333 | 3.52166667 | 4.38 | 3.44333333 | 4.46 | 4.43 | 5.305 | 4.91333333 | 6.04833333 | 4.49333333 | 3.73666667 | 4.20166667 | 3.81666667 | 4.60064103 | 1.03532817 | 0.28714837 |
| 10 | 5.488 | 2.923 | 3.69833333 | 2.84066667 | 4.24666667 | 4.194 | 5.131 | 4.419 | 4.95966667 | 3.61233333 | 2.15833333 | 3.149 | 2.25433333 | 3.77494872 | 1.07774617 | 0.29891301 |
| 15 | 5.39333333 | 3.06633333 | 2.39933333 | 2.877 | 3.11633333 | 3.00933333 | 5.34233333 | 3.456 | 4.34066667 | 4.30433333 | 1.77 | 2.53586207 | 1.981 | 3.35322016 | 1.1711773 | 0.32482614 |

### Peak expiratory flow

Week 1

WT

| Time (min) | 83 | 86 | 87 | 88 | 81 | 82 | A | 156 | 158 | 161 | Average | SD | SE |
|---|---|---|---|---|---|---|---|---|---|---|---|---|---|
| 0 | 2.22122222 | 1.84933333 | 1.68966667 | 2.31277778 | 1.58022222 | 1.56933333 | 1.61266667 | 2.148 | 0.88611111 | 0.94911111 | 1.68184444 | 0.4867444 | 0.1539221 |
| 1 | 2.185 | 2.73166667 | 2.25333333 | 3.495 | 2.216 | 2.32166667 | 0.81166667 | 2.434 | 1.208 | 1.576 | 2.12323333 | 0.76554678 | 0.24208715 |
| 2 | 2.01666667 | 2.41833333 | 2.075 | 1.9 | 1.865 | 1.99583333 | 1.22166667 | 2.11166667 | 0.95833333 | 1.23333333 | 1.77958333 | 0.47310233 | 0.14960809 |
| 3 | 2.015 | 2.24666667 | 2.15833333 | 1.62666667 | 1.84 | 2.04333333 | 1.53 | 2.11333333 | 1.13 | 1.50333333 | 1.82066667 | 0.36035665 | 0.11395478 |
| 4 | 2.12166667 | 2.19166667 | 2.645 | 2.10666667 | 1.61 | 2.05 | 1.33833333 | 2.96333333 | 1.4 | 1.49333333 | 1.992 | 0.53907603 | 0.17047081 |
| 5 | 1.83 | 2.285 | 2.06833333 | 1.91666667 | 1.625 | 2.11833333 | 1.19666667 | 2.32833333 | 1.27166667 | 1.545 | 1.8185 | 0.39966456 | 0.12638503 |
| 10 | 1.399 | 1.848 | 1.528 | 1.76166667 | 1.67733333 | 1.751 | 1.02866667 | 2.24533333 | 1.42833333 | 1.31633333 | 1.59836667 | 0.33578978 | 0.10618605 |
| 15 | 1.431 | 1.90333333 | 1.65466667 | 1.57466667 | 1.75066667 | 1.52962963 | 0.73966667 | 2.28 | 0.88966667 | 1.021 | 1.47742963 | 0.47656266 | 0.15070234 |

Het

| Time (min) | B | D | E | F | G | H | 155 | 157 | 159 | 160 | 162 | 190 | 196 | 199 | 201 | 205 | 207 | 208 | 210 | Average | SD | SE |
|---|---|---|---|---|---|---|---|---|---|---|---|---|---|---|---|---|---|---|---|---|---|---|
| 0 | 1.25744444 | 1.33222222 | 1.585 | 0.92633333 | 0.90588889 | 1.63277778 | 1.678 | 1.91544444 | 0.85033333 | 1.82666667 | 1.23955556 | 2.11288889 | 1.24022222 | 1.29555556 | 1.67088889 | 1.41144444 | 1.095 | 0.71855556 | 0.68322222 | 1.33565497 | 0.39078046 | 0.08965119 |
| 1 | 2.47666667 | 2.46 | 2.80166667 | 1.244 | 1.564 | 1.776 | 2.136 | 1.96 | 2.21333333 | 2.232 | 2.026 | 2.4 | 2.41 | 2.33333333 | 2.3 | 1.455 | 2.42333333 | 2.12333333 | 1.42666667 | 2.09270175 | 0.40082555 | 0.09195569 |
| 2 | 1.84666667 | 2.02333333 | 1.83333333 | 1.16333333 | 1.3 | 1.48 | 1.63333333 | 1.61833333 | 1.49666667 | 1.53666667 | 1.405 | 1.90833333 | 2.065 | 2.01333333 | 2.23833333 | 1.24166667 | 1.3 | 0.99666667 | 1.10166667 | 1.5895614 | 0.35236688 | 0.08083851 |
| 3 | 1.60166667 | 2.09333333 | 1.26 | 0.9 | 1.25666667 | 1.47166667 | 1.80166667 | 1.42166667 | 1.14 | 1.39666667 | 1.06166667 | 2.09166667 | 1.89166667 | 1.765 | 1.965 | 1.235 | 1.37833333 | 0.88333333 | 0.985 | 1.45263158 | 0.38474898 | 0.08826747 |
| 4 | 1.64666667 | 2.41666667 | 1.14 | 0.86166667 | 1.22 | 1.21666667 | 2.02 | 1.18166667 | 0.97333333 | 1.49333333 | 0.97 | 2.71666667 | 1.98666667 | 2.19 | 2.25666667 | 1.23666667 | 1.50333333 | 0.87666667 | 1.05666667 | 1.52438596 | 0.57729552 | 0.13244068 |
| 5 | 1.615 | 2.32333333 | 1.24833333 | 1.77666667 | 1.195 | 1.30833333 | 1.67666667 | 1.14166667 | 1.22333333 | 1.80833333 | 0.945 | 2.91 | 2.05166667 | 2.35666667 | 2.63 | 1.44666667 | 1.46 | 0.95166667 | 0.985 | 1.63438596 | 0.5782981 | 0.13267068 |
| 10 | 1.78666667 | 2.03033333 | 1.35733333 | 0.83633333 | 1.06133333 | 1.514 | 1.64266667 | 1.32133333 | 1.117 | 1.85366667 | 1.24833333 | 2.45933333 | 1.40833333 | 2.11333333 | 2.07466667 | 1.30466667 | 1.24233333 | 1.03 | 0.828 | 1.48577193 | 0.44430187 | 0.10192984 |
| 15 | 1.489 | 1.85633333 | 1.41433333 | 0.947 | 1.20166667 | 1.39 | 1.10733333 | 1.43366667 | 0.86233333 | 1.63833333 | 0.87966667 | 2.05333333 | 1.317 | 2.01133333 | 1.76733333 | 1.001 | 0.74033333 | 0.794 | 0.754 | 1.29778947 | 0.41833808 | 0.09597334 |

Week 2

WT

| Time (min) | 83 | 86 | 87 | 88 | 81 | 82 | A | 156 | 158 | 161 | Average | SD | SE |
|---|---|---|---|---|---|---|---|---|---|---|---|---|---|
| 0 | 2.94888889 | 2.88811111 | 2.32633333 | 3.12144444 | 2.48811111 | 1.68888889 | 2.44533333 | 1.763 | 1.78155556 | 1.92688889 | 2.33785556 | 0.5327492 | 0.16847009 |
| 1 | 3.18666667 | 4.06333333 | 3.72333333 | 4.71333333 | 3.53333333 | 4.01166667 | 2.94166667 | 3.21333333 | 1.45833333 | 2.05166667 | 3.28966667 | 0.96713073 | 0.30583359 |
| 2 | 2.37166667 | 4.03333333 | 3.49333333 | 4.75 | 3.31666667 | 3.83666667 | 2.27166667 | 3.03333333 | 1.51166667 | 1.895 | 3.05133333 | 1.02795038 | 0.32506645 |
| 3 | 2.51833333 | 3.60333333 | 3.425 | 4.25333333 | 2.98666667 | 3.40166667 | 2.345 | 2.43 | 1.505 | 2.075 | 2.85433333 | 0.82606056 | 0.26122329 |
| 4 | 2.49 | 3.61166667 | 3.53 | 3.76166667 | 3.16166667 | 3.20833333 | 2.74666667 | 2.15333333 | 1.63333333 | 2.78666667 | 2.90833333 | 0.6811016 | 0.21538324 |
| 5 | 2.82833333 | 3.50166667 | 3.01666667 | 3.32666667 | 2.75333333 | 3.05166667 | 2.375 | 1.75833333 | 1.535 | 2.47166667 | 2.66183333 | 0.63820903 | 0.20181942 |
| 10 | 2.42 | 2.90333333 | 2.59066667 | 3.4 | 2.08033333 | 2.07366667 | 2.742 | 1.98266667 | 1.21533333 | 1.68533333 | 2.30933333 | 0.63437534 | 0.2006071 |
| 15 | 2.46133333 | 2.27166667 | 2.18666667 | 3.17366667 | 1.81666667 | 1.861 | 2.389 | 1.87733333 | 1.503 | 1.206 | 2.07463333 | 0.55176615 | 0.17448378 |

Het

| Time (min) | B | D | E | G | H | 155 | 157 | 160 | 162 | 190 | 196 | 199 | 201 | 205 | 207 | 208 | 210 | Average | SD | SE |
|---|---|---|---|---|---|---|---|---|---|---|---|---|---|---|---|---|---|---|---|---|
| 0 | 2.34088889 | 1.44844444 | 1.10366667 | 0.828 | 1.827 | 1.362 | 1.77111111 | 1.52788889 | 1.46277778 | 1.51077778 | 1.56777778 | 1.26966667 | 1.61977778 | 1.76288889 | 1.00444444 | 1.97555556 | 1.518 | 1.52356863 | 0.36433876 | 0.08836513 |
| 1 | 3.11833333 | 2.615 | 2.606 | 1.38666667 | 1.79666667 | 2.49 | 2.26666667 | 1.79 | 1.745 | 2.84333333 | 2.415 | 2.18 | 2.37166667 | 1.61 | 2.00166667 | 2.83333333 | 2.07166667 | 2.24358824 | 0.48289139 | 0.11711836 |
| 2 | 2.38 | 2.64 | 1.93333333 | 1.44333333 | 1.72166667 | 1.79 | 1.82 | 1.70666667 | 1.895 | 2.52333333 | 2.74166667 | 1.94166667 | 2.485 | 1.52333333 | 2.20166667 | 1.75 | 2.205 | 2.04127451 | 0.39728616 | 0.09635605 |
| 3 | 1.885 | 2.485 | 1.85 | 1.37833333 | 2.20666667 | 1.7 | 1.31833333 | 1.71 | 1.73333333 | 2.64166667 | 3.11333333 | 2.1 | 2.91166667 | 1.79666667 | 2.51833333 | 1.385 | 2.11666667 | 2.05 | 0.53395602 | 0.12950336 |
| 4 | 2.67666667 | 2.50166667 | 2.27333333 | 1.41333333 | 1.94166667 | 1.47 | 1.28833333 | 1.84 | 1.49833333 | 2.59166667 | 2.69 | 2.66 | 2.66166667 | 1.60333333 | 2.10833333 | 1.85166667 | 2.15333333 | 2.07196078 | 0.50141128 | 0.1216101 |
| 5 | 3.02 | 2.85 | 1.91 | 1.64166667 | 1.52 | 1.49333333 | 1.205 | 1.875 | 1.395 | 2.365 | 2.21666667 | 2.40166667 | 2.31166667 | 1.53333333 | 1.88666667 | 2.33666667 | 1.77 | 1.98421569 | 0.51426293 | 0.12472708 |
| 10 | 1.62866667 | 2.58633333 | 1.77366667 | 1.42766667 | 1.52166667 | 1.381 | 1.52233333 | 1.48966667 | 1.23633333 | 2.01666667 | 1.67866667 | 1.90933333 | 1.72033333 | 1.48066667 | 1.424 | 1.88866667 | 1.72 | 1.67092157 | 0.31479101 | 0.07634804 |
| 15 | 1.93366667 | 2.212 | 1.29333333 | 1.091 | 1.32566667 | 1.65833333 | 1.54733333 | 1.08933333 | 0.696 | 1.89966667 | 2.51533333 | 1.77866667 | 1.91733333 | 1.31933333 | 0.87533333 | 1.58766667 | 1.422 | 1.53894118 | 0.47638382 | 0.11554005 |

Week 3

WT

| Time (min) | 83 | 86 | 87 | 88 | 81 | 82 | A | 156 | 158 | 161 | Average | SD | SE |
|---|---|---|---|---|---|---|---|---|---|---|---|---|---|
| 0 | 3.11711111 | 2.96211111 | 2.42744444 | 2.53988889 | 1.82333333 | 1.68711111 | 2.59211111 | 2.25766667 | 1.04122222 | 1.76666667 | 2.22146667 | 0.63919276 | 0.2021305 |
| 1 | 3.63333333 | 4.04833333 | 4.83666667 | 4.54833333 | 4.26666667 | 3.44833333 | 3.79 | 3.38666667 | 1.665 | 2.13833333 | 3.57616667 | 1.00304353 | 0.31719022 |
| 2 | 3.74166667 | 3.46833333 | 3.75 | 3.88166667 | 3.51166667 | 2.24333333 | 2.70666667 | 2.155 | 1.17166667 | 1.685 | 2.8315 | 0.97363943 | 0.30789182 |
| 3 | 4.08 | 2.955 | 3.89166667 | 4.25166667 | 3.935 | 2.18 | 3.08333333 | 2.03166667 | 1.12166667 | 2.26 | 2.979 | 1.05896852 | 0.33487525 |
| 4 | 3.245 | 2.94833333 | 3.575 | 4.145 | 4.59833333 | 2.215 | 3.00833333 | 2.095 | 1.15 | 2.56166667 | 2.95416667 | 1.01587196 | 0.32124692 |
| 5 | 2.96 | 2.70666667 | 2.93 | 3.38166667 | 4.26666667 | 2.47666667 | 3.05833333 | 1.93666667 | 1.505 | 2.17166667 | 2.73933333 | 0.78118903 | 0.24703366 |
| 10 | 3.164 | 2.18633333 | 2.68466667 | 2.056 | 2.85766667 | 2.24233333 | 2.22633333 | 2.14033333 | 1.384 | 1.737 | 2.26786667 | 0.52258522 | 0.16525596 |
| 15 | 2.96233333 | 1.807 | 1.77933333 | 2.18166667 | 1.88466667 | 1.75733333 | 1.616 | 2.32933333 | 1.17733333 | 1.03566667 | 1.85306667 | 0.55417221 | 0.17524464 |

Het

| Time (min) | B | D | E | H | 155 | 157 | 190 | 196 | 199 | 201 | 205 | 207 | 208 | 210 | Average | SD | SE |
|---|---|---|---|---|---|---|---|---|---|---|---|---|---|---|---|---|---|
| 0 | 1.76233333 | 1.49377778 | 1.104 | 1.20411111 | 1.43355556 | 1.47288889 | 1.54133333 | 1.52155556 | 1.47122222 | 1.575 | 0.94544444 | 0.94966667 | 0.98244444 | 1.16066667 | 1.32985714 | 0.2653197 | 0.07090967 |
| 1 | 3.512 | 2.965 | 2.684 | 2.795 | 2.585 | 2.54166667 | 2.60833333 | 3.31666667 | 2.77166667 | 2.665 | 1.86166667 | 1.83833333 | 1.62 | 2.10166667 | 2.56185714 | 0.54594257 | 0.14590929 |
| 2 | 3.665 | 2.75166667 | 2.97833333 | 2.04166667 | 1.56666667 | 1.88833333 | 2.555 | 3.11166667 | 2.24833333 | 2.20333333 | 1.37333333 | 1.96666667 | 1.73666667 | 1.91166667 | 2.28559524 | 0.64880087 | 0.17339933 |
| 3 | 3.88833333 | 3.18666667 | 2.585 | 2.52 | 1.40666667 | 1.66833333 | 2.75 | 3.40666667 | 2.11166667 | 2.965 | 1.44666667 | 2.40666667 | 1.365 | 1.845 | 2.39654762 | 0.79570306 | 0.21266059 |
| 4 | 3.68166667 | 3.45166667 | 3.53833333 | 2.965 | 1.33 | 2.39166667 | 2.85166667 | 3.09666667 | 2.52 | 3.13833333 | 1.325 | 1.98166667 | 1.295 | 1.79666667 | 2.52595238 | 0.85264839 | 0.22787987 |
| 5 | 2.54333333 | 2.82333333 | 3.91333333 | 2.55333333 | 1.34 | 2.41 | 2.31166667 | 2.13666667 | 2.40166667 | 2.69333333 | 1.285 | 1.69666667 | 1.81166667 | 1.33166667 | 2.2322619 | 0.71424026 | 0.19088874 |
| 10 | 2.68866667 | 2.07666667 | 1.89366667 | 2.117 | 1.986 | 1.917 | 1.911 | 1.93833333 | 2.21633333 | 2.702 | 1.67466667 | 1.31 | 1.78 | 1.289 | 1.96430952 | 0.40965066 | 0.10948375 |
| 15 | 2.31866667 | 1.35933333 | 1.28333333 | 1.665 | 1.689 | 2.067 | 1.63966667 | 2.051 | 1.68466667 | 2.427 | 1.71033333 | 1.03366667 | 0.96433333 | 1.09133333 | 1.6417381 | 0.46179572 | 0.1234201 |

Week 4

WT

| Time (min) | 83 | 86 | 87 | 88 | 81 | 82 | A | 156 | 158 | 161 | Average | SD | SE |
|---|---|---|---|---|---|---|---|---|---|---|---|---|---|
| 0 | 3.49933333 | 2.15133333 | 2.62422222 | 3.182 | 1.80577778 | 2.20533333 | 2.87566667 | 2.80166667 | 1.08866667 | 2.00911111 | 2.42431111 | 0.71314931 | 0.22551761 |
| 1 | 3.81 | 4.18333333 | 4.59833333 | 5.02166667 | 3.83333333 | 3.19833333 | 3.80166667 | 3.84 | 2.06833333 | 2.40833333 | 3.67633333 | 0.90823625 | 0.28720952 |
| 2 | 3.06166667 | 3.55833333 | 4.47166667 | 4.45 | 3.065 | 2.42166667 | 3.15666667 | 2.845 | 1.07666667 | 1.77833333 | 2.9885 | 1.06256836 | 0.33601362 |
| 3 | 3.24 | 3.66333333 | 4.985 | 4.89333333 | 2.83666667 | 2.36833333 | 3.35333333 | 2.95166667 | 1.34 | 2.61166667 | 3.22433333 | 1.1044635 | 0.34926203 |
| 4 | 3.49333333 | 4.65166667 | 4.875 | 4.965 | 3.22333333 | 2.44166667 | 3.365 | 3.27 | 1.535 | 2.845 | 3.4665 | 1.10100948 | 0.34816977 |
| 5 | 3.27166667 | 4.165 | 3.84833333 | 3.955 | 3.87833333 | 2.67166667 | 2.83666667 | 3.255 | 1.62333333 | 2.11666667 | 3.16216667 | 0.84631264 | 0.26762756 |
| 10 | 2.584 | 2.81133333 | 2.98 | 2.98833333 | 3.02333333 | 2.32033333 | 2.401 | 3.064 | 1.63166667 | 1.98966667 | 2.57936667 | 0.48971205 | 0.15486055 |
| 15 | 1.93966667 | 1.81033333 | 2.15766667 | 1.814 | 2.21666667 | 1.88433333 | 2.03766667 | 1.716 | 1.20866667 | 1.183 | 1.7968 | 0.3531228 | 0.11166723 |

Het

| Time (min) | B | E | H | 155 | 157 | 190 | 196 | 199 | 201 | 205 | 207 | 208 | 210 | Average | SD | SE |
|---|---|---|---|---|---|---|---|---|---|---|---|---|---|---|---|---|
| 0 | 1.92077778 | 1.18122222 | 1.227 | 1.64111111 | 1.66122222 | 2.04044444 | 1.51333333 | 1.84422222 | 1.96666667 | 1.20755556 | 1.63788889 | 1.54488889 | 1.42155556 | 1.60060684 | 0.28993605 | 0.08041379 |
| 1 | 4.05 | 3.23666667 | 2.75666667 | 2.72833333 | 2.49666667 | 2.88833333 | 2.99166667 | 2.55 | 3.75833333 | 2.28166667 | 1.815 | 1.94333333 | 1.91666667 | 2.72410256 | 0.68042934 | 0.18871714 |
| 2 | 3.605 | 3.56833333 | 2.69833333 | 2.14333333 | 1.89333333 | 2.22333333 | 2.97 | 1.92666667 | 2.73333333 | 1.85 | 1.84666667 | 1.47 | 1.745 | 2.35948718 | 0.69529083 | 0.19283898 |
| 3 | 5.21166667 | 4.76833333 | 3.07666667 | 2.06 | 2.24 | 2.85666667 | 3.25833333 | 1.83666667 | 2.41333333 | 2.52 | 2.2 | 1.645 | 1.865 | 2.76551282 | 1.10114364 | 0.3054023 |
| 4 | 4.82666667 | 3.70333333 | 2.77333333 | 2.08833333 | 2.31 | 2.68833333 | 3.045 | 3.21666667 | 3.39833333 | 2.82166667 | 2.57 | 1.61666667 | 2.43666667 | 2.88423077 | 0.80458081 | 0.22315057 |
| 5 | 4.03166667 | 2.12 | 2.46166667 | 1.46666667 | 2.59166667 | 2.42 | 2.82333333 | 2.81 | 3.44666667 | 2.29 | 1.91666667 | 2.12 | 1.81166667 | 2.48538462 | 0.68745173 | 0.19066481 |
| 10 | 3.208 | 1.642 | 1.739 | 1.475 | 2.25833333 | 2.28033333 | 2.55333333 | 2.25666667 | 2.668 | 1.89966667 | 1.02633333 | 1.50433333 | 1.05233333 | 1.96641026 | 0.6443317 | 0.17870546 |
| 15 | 3.357 | 1.607 | 1.106 | 1.52433333 | 1.47366667 | 1.52566667 | 2.70933333 | 1.60566667 | 2.31733333 | 2.89066667 | 0.87366667 | 1.16862069 | 0.99766667 | 1.78127851 | 0.78626936 | 0.21807188 |

### Tidal volume

#### Week 1

WT

| Time (min) | 83 | 86 | 87 | 88 | 81 | 82 | A | 156 | 158 | 161 | Average | SD | SE |
|---|---|---|---|---|---|---|---|---|---|---|---|---|---|
| 0 | 0.12866667 | 0.12755556 | 0.13333333 | 0.15333333 | 0.12233333 | 0.12122222 | 0.11366667 | 0.14366667 | 0.08022222 | 0.09111111 | 0.12151111 | 0.02216667 | 0.00700972 |
| 1 | 0.13 | 0.15666667 | 0.145 | 0.19666667 | 0.142 | 0.13416667 | 0.05166667 | 0.13 | 0.086 | 0.096 | 0.12681667 | 0.04037146 | 0.01276658 |
| 2 | 0.105 | 0.16166667 | 0.12666667 | 0.13166667 | 0.12833333 | 0.12416667 | 0.055 | 0.10833333 | 0.07666667 | 0.07666667 | 0.10941667 | 0.03201574 | 0.01012427 |
| 3 | 0.1 | 0.155 | 0.12666667 | 0.12166667 | 0.13 | 0.13166667 | 0.06 | 0.14333333 | 0.08333333 | 0.095 | 0.11466667 | 0.02932155 | 0.00927229 |
| 4 | 0.11333333 | 0.13 | 0.135 | 0.125 | 0.12 | 0.12454545 | 0.065 | 0.12 | 0.09 | 0.10166667 | 0.11245455 | 0.02136164 | 0.00675514 |
| 5 | 0.11166667 | 0.13166667 | 0.15 | 0.14 | 0.12333333 | 0.115 | 0.065 | 0.15 | 0.095 | 0.11166667 | 0.11933333 | 0.02615481 | 0.00827088 |
| 10 | 0.10966667 | 0.129 | 0.123 | 0.13533333 | 0.12733333 | 0.133 | 0.06266667 | 0.14733333 | 0.09666667 | 0.101 | 0.1165 | 0.02465728 | 0.00779732 |
| 15 | 0.10766667 | 0.12966667 | 0.12866667 | 0.13333333 | 0.13533333 | 0.12814815 | 0.05433333 | 0.149 | 0.084 | 0.09266667 | 0.11428148 | 0.02919124 | 0.00923108 |

Het

| Time (min) | B | D | E | F | G | H | 155 | 157 | 159 | 160 | 162 | 190 | 196 | 199 | 201 | 205 | 207 | 208 | 210 | Average | SD | SE |
|---|---|---|---|---|---|---|---|---|---|---|---|---|---|---|---|---|---|---|---|---|---|---|
| 0 | 0.10255556 | 0.10388889 | 0.11533333 | 0.07155556 | 0.08177778 | 0.10766667 | 0.08777778 | 0.09666667 | 0.08411111 | 0.10566667 | 0.08655556 | 0.09544444 | 0.10711111 | 0.12044444 | 0.11355556 | 0.06688889 | 0.08122222 | 0.06655556 | 0.07311111 | 0.09304678 | 0.01687773 | 0.00387202 |
| 1 | 0.15333333 | 0.14 | 0.14 | 0.068 | 0.084 | 0.118 | 0.118 | 0.10666667 | 0.13 | 0.118 | 0.112 | 0.125 | 0.13833333 | 0.16666667 | 0.15 | 0.07666667 | 0.11333333 | 0.09833333 | 0.095 | 0.11849123 | 0.02653843 | 0.00608833 |
| 2 | 0.15 | 0.13 | 0.13166667 | 0.075 | 0.08 | 0.08833333 | 0.1 | 0.10333333 | 0.09166667 | 0.09833333 | 0.10333333 | 0.12166667 | 0.13 | 0.16666667 | 0.13666667 | 0.07666667 | 0.08833333 | 0.07166667 | 0.07833333 | 0.10640351 | 0.02787906 | 0.00639589 |
| 3 | 0.12166667 | 0.12333333 | 0.10333333 | 0.06166667 | 0.08166667 | 0.10333333 | 0.10166667 | 0.09 | 0.08666667 | 0.085 | 0.08833333 | 0.11833333 | 0.12833333 | 0.155 | 0.12833333 | 0.075 | 0.09 | 0.06833333 | 0.08 | 0.09947368 | 0.02423555 | 0.00556002 |
| 4 | 0.12833333 | 0.115 | 0.10333333 | 0.06166667 | 0.08666667 | 0.09166667 | 0.105 | 0.09 | 0.08333333 | 0.09666667 | 0.08666667 | 0.13833333 | 0.12666667 | 0.14 | 0.14166667 | 0.085 | 0.1 | 0.06666667 | 0.08 | 0.10140351 | 0.02422415 | 0.0055574 |
| 5 | 0.135 | 0.11 | 0.105 | 0.305 | 0.085 | 0.09833333 | 0.105 | 0.08666667 | 0.085 | 0.10833333 | 0.08 | 0.15166667 | 0.12 | 0.13833333 | 0.12833333 | 0.09 | 0.09666667 | 0.07 | 0.08 | 0.11464912 | 0.05119431 | 0.01174478 |
| 10 | 0.11966667 | 0.099 | 0.105 | 0.05533333 | 0.079 | 0.10966667 | 0.09766667 | 0.09866667 | 0.091 | 0.106 | 0.09766667 | 0.129 | 0.12466667 | 0.09366667 | 0.106 | 0.07933333 | 0.09233333 | 0.074 | 0.07233333 | 0.09631579 | 0.01865922 | 0.00428072 |
| 15 | 0.11033333 | 0.09166667 | 0.10566667 | 0.05933333 | 0.08833333 | 0.11066667 | 0.085 | 0.10633333 | 0.087 | 0.10533333 | 0.09533333 | 0.116 | 0.11833333 | 0.101 | 0.12533333 | 0.08 | 0.07233333 | 0.071 | 0.069 | 0.09463158 | 0.01863552 | 0.00427528 |

#### Week 2

WT

| Time (min) | 83 | 86 | 87 | 88 | 81 | 82 | A | 156 | 158 | 161 | Average | SD | SE |
|---|---|---|---|---|---|---|---|---|---|---|---|---|---|
| 0 | 0.16144444 | 0.18344444 | 0.17622222 | 0.18655556 | 0.17166667 | 0.15544444 | 0.16622222 | 0.15344444 | 0.10677778 | 0.12966667 | 0.15908889 | 0.02472753 | 0.00781953 |
| 1 | 0.19 | 0.225 | 0.2 | 0.24666667 | 0.2 | 0.205 | 0.17333333 | 0.215 | 0.09 | 0.12666667 | 0.18716667 | 0.0467793 | 0.01479291 |
| 2 | 0.12833333 | 0.23 | 0.19166667 | 0.27833333 | 0.2 | 0.20166667 | 0.12666667 | 0.17166667 | 0.09666667 | 0.115 | 0.174 | 0.05731761 | 0.01812542 |
| 3 | 0.15333333 | 0.20333333 | 0.20666667 | 0.25833333 | 0.18833333 | 0.20166667 | 0.14833333 | 0.155 | 0.10333333 | 0.13666667 | 0.1755 | 0.04450031 | 0.01407223 |
| 4 | 0.16 | 0.21 | 0.22166667 | 0.26333333 | 0.19833333 | 0.18 | 0.175 | 0.155 | 0.10666667 | 0.125 | 0.1795 | 0.04638839 | 0.0146693 |
| 5 | 0.15333333 | 0.20166667 | 0.215 | 0.20666667 | 0.17333333 | 0.19 | 0.16333333 | 0.16 | 0.12166667 | 0.13833333 | 0.17233333 | 0.03071373 | 0.00971253 |
| 10 | 0.164 | 0.18333333 | 0.21266667 | 0.19866667 | 0.17066667 | 0.18466667 | 0.18366667 | 0.18933333 | 0.11666667 | 0.121 | 0.17246667 | 0.03129911 | 0.00989765 |
| 15 | 0.173 | 0.18333333 | 0.193 | 0.22666667 | 0.169 | 0.18466667 | 0.17766667 | 0.18233333 | 0.10833333 | 0.11566667 | 0.17136667 | 0.03510318 | 0.0111006 |

Het

| Time (min) | B | D | E | G | H | 155 | 157 | 160 | 162 | 190 | 196 | 199 | 201 | 205 | 207 | 208 | 210 | Average | SD | SE |
|---|---|---|---|---|---|---|---|---|---|---|---|---|---|---|---|---|---|---|---|---|
| 0 | 0.14288889 | 0.12711111 | 0.10111111 | 0.06466667 | 0.13244444 | 0.10666667 | 0.10744444 | 0.09722222 | 0.12433333 | 0.12111111 | 0.16144444 | 0.12677778 | 0.138 | 0.09744444 | 0.09077778 | 0.10022222 | 0.10555556 | 0.11442484 | 0.02310821 | 0.00560457 |
| 1 | 0.17833333 | 0.17 | 0.136 | 0.09333333 | 0.10666667 | 0.14833333 | 0.16 | 0.12 | 0.12 | 0.16166667 | 0.15166667 | 0.16333333 | 0.165 | 0.09 | 0.115 | 0.13 | 0.10166667 | 0.13594118 | 0.02865 | 0.00694865 |
| 2 | 0.16833333 | 0.17 | 0.12833333 | 0.10333333 | 0.105 | 0.135 | 0.14 | 0.09833333 | 0.11333333 | 0.135 | 0.145 | 0.14 | 0.13 | 0.085 | 0.11 | 0.11 | 0.11 | 0.12509804 | 0.02369123 | 0.00574597 |
| 3 | 0.145 | 0.17 | 0.12333333 | 0.09666667 | 0.135 | 0.13666667 | 0.11166667 | 0.11333333 | 0.11166667 | 0.12166667 | 0.13166667 | 0.15 | 0.13833333 | 0.10166667 | 0.11166667 | 0.10166667 | 0.10833333 | 0.12401961 | 0.01991352 | 0.00482974 |
| 4 | 0.16166667 | 0.17 | 0.13333333 | 0.09833333 | 0.14 | 0.12 | 0.11166667 | 0.11666667 | 0.11 | 0.125 | 0.13833333 | 0.14 | 0.12 | 0.1 | 0.11166667 | 0.12 | 0.11 | 0.12509804 | 0.01992582 | 0.00483272 |
| 5 | 0.15333333 | 0.16666667 | 0.13833333 | 0.10333333 | 0.13166667 | 0.11 | 0.11166667 | 0.12166667 | 0.12 | 0.11666667 | 0.125 | 0.15666667 | 0.12666667 | 0.1 | 0.09166667 | 0.12833333 | 0.10666667 | 0.12401961 | 0.02063286 | 0.0050042 |
| 10 | 0.136 | 0.16766667 | 0.12766667 | 0.09933333 | 0.123 | 0.10766667 | 0.118 | 0.12 | 0.11866667 | 0.13266667 | 0.159 | 0.12966667 | 0.14233333 | 0.09766667 | 0.09833333 | 0.119 | 0.10266667 | 0.1234902 | 0.0201822 | 0.0048949 |
| 15 | 0.13866667 | 0.15966667 | 0.11933333 | 0.08233333 | 0.126 | 0.12 | 0.127 | 0.12433333 | 0.124 | 0.13066667 | 0.15633333 | 0.12566667 | 0.13566667 | 0.101 | 0.098 | 0.11233333 | 0.10033333 | 0.12243137 | 0.01994115 | 0.00483644 |

#### Week 3

WT

| Time (min) | 83 | 86 | 87 | 88 | 81 | 82 | A | 156 | 158 | 161 | Average | SD | SE |
|---|---|---|---|---|---|---|---|---|---|---|---|---|---|
| 0 | 0.18566667 | 0.18011111 | 0.18477778 | 0.18677778 | 0.16577778 | 0.14811111 | 0.16655556 | 0.16611111 | 0.10788889 | 0.12411111 | 0.16158889 | 0.0271367 | 0.00858138 |
| 1 | 0.17666667 | 0.195 | 0.21666667 | 0.205 | 0.21166667 | 0.15166667 | 0.18166667 | 0.19333333 | 0.11833333 | 0.12833333 | 0.17783333 | 0.03434619 | 0.01086122 |
| 2 | 0.155 | 0.19 | 0.17166667 | 0.18666667 | 0.195 | 0.13833333 | 0.14666667 | 0.145 | 0.09666667 | 0.105 | 0.153 | 0.03401888 | 0.01075771 |
| 3 | 0.17666667 | 0.18833333 | 0.16166667 | 0.19833333 | 0.21333333 | 0.14166667 | 0.18 | 0.15833333 | 0.1 | 0.12666667 | 0.1645 | 0.03439109 | 0.01087542 |
| 4 | 0.18666667 | 0.18333333 | 0.17166667 | 0.17166667 | 0.22166667 | 0.14333333 | 0.155 | 0.15833333 | 0.10666667 | 0.14166667 | 0.164 | 0.03101374 | 0.0098074 |
| 5 | 0.15666667 | 0.165 | 0.155 | 0.19 | 0.21166667 | 0.15666667 | 0.17 | 0.17333333 | 0.105 | 0.13833333 | 0.16216667 | 0.0286103 | 0.00904737 |
| 10 | 0.155 | 0.18833333 | 0.17666667 | 0.17466667 | 0.17733333 | 0.17666667 | 0.16866667 | 0.19533333 | 0.12666667 | 0.13266667 | 0.1672 | 0.02251485 | 0.00711982 |
| 15 | 0.166 | 0.175 | 0.16266667 | 0.17733333 | 0.189 | 0.17333333 | 0.14766667 | 0.19833333 | 0.121 | 0.11733333 | 0.16276667 | 0.02681995 | 0.00848121 |

Het

| Time (min) | B | D | E | H | 155 | 157 | 190 | 196 | 199 | 201 | 205 | 207 | 208 | 210 | Average | SD | SE |
|---|---|---|---|---|---|---|---|---|---|---|---|---|---|---|---|---|---|
| 0 | 0.14566667 | 0.14266667 | 0.10022222 | 0.12377778 | 0.12344444 | 0.13333333 | 0.128 | 0.13011111 | 0.13955556 | 0.145 | 0.08311111 | 0.09588889 | 0.09011111 | 0.09888889 | 0.11998413 | 0.02187806 | 0.00584716 |
| 1 | 0.202 | 0.18166667 | 0.148 | 0.14333333 | 0.17166667 | 0.16333333 | 0.17166667 | 0.2 | 0.19 | 0.19 | 0.13 | 0.12 | 0.10333333 | 0.11666667 | 0.15940476 | 0.03272553 | 0.00874627 |
| 2 | 0.185 | 0.16 | 0.14666667 | 0.11333333 | 0.13666667 | 0.15833333 | 0.13 | 0.16 | 0.17833333 | 0.17 | 0.1 | 0.11333333 | 0.10833333 | 0.105 | 0.14035714 | 0.02900176 | 0.00775105 |
| 3 | 0.17333333 | 0.14666667 | 0.15833333 | 0.13833333 | 0.13166667 | 0.15 | 0.13166667 | 0.155 | 0.15666667 | 0.15166667 | 0.11166667 | 0.105 | 0.115 | 0.09 | 0.13678571 | 0.02382687 | 0.006368 |
| 4 | 0.16666667 | 0.15 | 0.17333333 | 0.14333333 | 0.13 | 0.12833333 | 0.14666667 | 0.145 | 0.14833333 | 0.15333333 | 0.11 | 0.11 | 0.11 | 0.10833333 | 0.13738095 | 0.02164905 | 0.00578595 |
| 5 | 0.16666667 | 0.13166667 | 0.16833333 | 0.12666667 | 0.12333333 | 0.13 | 0.13833333 | 0.12 | 0.15166667 | 0.14166667 | 0.10833333 | 0.09 | 0.11333333 | 0.11166667 | 0.13011905 | 0.02206696 | 0.00589764 |
| 10 | 0.16733333 | 0.14833333 | 0.13666667 | 0.15366667 | 0.14933333 | 0.14966667 | 0.14366667 | 0.14566667 | 0.14533333 | 0.169 | 0.111 | 0.10266667 | 0.11233333 | 0.11633333 | 0.13935714 | 0.02087425 | 0.00557888 |
| 15 | 0.16333333 | 0.14633333 | 0.12933333 | 0.14366667 | 0.14666667 | 0.14566667 | 0.144 | 0.16433333 | 0.15766667 | 0.15866667 | 0.10866667 | 0.103 | 0.10466667 | 0.116 | 0.138 | 0.02179665 | 0.0058254 |

#### Week 4

WT

| Time (min) | 83 | 86 | 87 | 88 | 81 | 82 | A | 156 | 158 | 161 | Average | SD | SE |
|---|---|---|---|---|---|---|---|---|---|---|---|---|---|
| 0 | 0.202 | 0.17877778 | 0.181 | 0.20622222 | 0.17633333 | 0.16755556 | 0.19666667 | 0.204 | 0.12188889 | 0.14588889 | 0.17803333 | 0.02728039 | 0.00862682 |
| 1 | 0.21333333 | 0.195 | 0.24666667 | 0.24166667 | 0.21166667 | 0.165 | 0.18 | 0.18333333 | 0.13166667 | 0.12833333 | 0.18966667 | 0.04067942 | 0.01286396 |
| 2 | 0.15333333 | 0.19833333 | 0.21666667 | 0.22166667 | 0.19 | 0.15166667 | 0.165 | 0.18166667 | 0.10166667 | 0.11666667 | 0.16966667 | 0.03991349 | 0.01262175 |
| 3 | 0.16166667 | 0.19833333 | 0.22833333 | 0.23333333 | 0.18166667 | 0.16166667 | 0.155 | 0.17833333 | 0.10666667 | 0.13833333 | 0.17433333 | 0.03889048 | 0.01229825 |
| 4 | 0.18 | 0.23666667 | 0.24666667 | 0.245 | 0.22166667 | 0.16 | 0.16 | 0.18166667 | 0.12 | 0.16 | 0.19116667 | 0.0436813 | 0.01381324 |
| 5 | 0.18 | 0.20833333 | 0.21166667 | 0.22333333 | 0.21833333 | 0.17 | 0.14333333 | 0.19 | 0.11833333 | 0.12666667 | 0.179 | 0.03849162 | 0.01217212 |
| 10 | 0.18666667 | 0.19233333 | 0.203 | 0.20833333 | 0.19633333 | 0.17633333 | 0.18 | 0.201 | 0.13233333 | 0.134 | 0.18103333 | 0.02713544 | 0.00858098 |
| 15 | 0.174 | 0.16866667 | 0.20166667 | 0.17133333 | 0.20533333 | 0.181 | 0.18466667 | 0.188 | 0.12466667 | 0.13266667 | 0.1732 | 0.02640997 | 0.00835157 |

Het

| Time (min) | B | E | H | 155 | 157 | 190 | 196 | 199 | 201 | 205 | 207 | 208 | 210 | Average | SD | SE |
|---|---|---|---|---|---|---|---|---|---|---|---|---|---|---|---|---|
| 0 | 0.171 | 0.11033333 | 0.13644444 | 0.14033333 | 0.17133333 | 0.14522222 | 0.14566667 | 0.138 | 0.14133333 | 0.10777778 | 0.10666667 | 0.10688889 | 0.10611111 | 0.1328547 | 0.02350757 | 0.00651983 |
| 1 | 0.24666667 | 0.16833333 | 0.15166667 | 0.18166667 | 0.15666667 | 0.18 | 0.17 | 0.18833333 | 0.21166667 | 0.145 | 0.12333333 | 0.12666667 | 0.12333333 | 0.16717949 | 0.03589068 | 0.00995428 |
| 2 | 0.19166667 | 0.17666667 | 0.13 | 0.12333333 | 0.11666667 | 0.14 | 0.155 | 0.165 | 0.18666667 | 0.11666667 | 0.1 | 0.11333333 | 0.115 | 0.14076923 | 0.03087171 | 0.00856227 |
| 3 | 0.22166667 | 0.18666667 | 0.14833333 | 0.10333333 | 0.12 | 0.125 | 0.13833333 | 0.155 | 0.16166667 | 0.12 | 0.105 | 0.12166667 | 0.12333333 | 0.14076923 | 0.0339295 | 0.00941035 |
| 4 | 0.21166667 | 0.15666667 | 0.125 | 0.12333333 | 0.13833333 | 0.13833333 | 0.13833333 | 0.15 | 0.17833333 | 0.13333333 | 0.11833333 | 0.10833333 | 0.12166667 | 0.14166667 | 0.02785545 | 0.00772571 |
| 5 | 0.20833333 | 0.14 | 0.12333333 | 0.14166667 | 0.13166667 | 0.17166667 | 0.15333333 | 0.13333333 | 0.17333333 | 0.12833333 | 0.10333333 | 0.11166667 | 0.12333333 | 0.14179487 | 0.02868621 | 0.00795612 |
| 10 | 0.19733333 | 0.13166667 | 0.13433333 | 0.13833333 | 0.17066667 | 0.15066667 | 0.15133333 | 0.141 | 0.14666667 | 0.13066667 | 0.10366667 | 0.10666667 | 0.11866667 | 0.14012821 | 0.02515417 | 0.00697651 |
| 15 | 0.206 | 0.129 | 0.13633333 | 0.14133333 | 0.16733333 | 0.155 | 0.15333333 | 0.16966667 | 0.146 | 0.15366667 | 0.10066667 | 0.11068966 | 0.113 | 0.144771 | 0.02826113 | 0.00783823 |

### Expiratory volume

Week 1

WT

| Time (min) | Animal ID | | | | | | | | | | | | |
|---|---|---|---|---|---|---|---|---|---|---|---|---|---|
| | **83** | **86** | **87** | **88** | **81** | **82** | **A** | **156** | **158** | **161** | Average | SD | SE |
| 0 | 0.131 | 0.12855556 | 0.13422222 | 0.15877778 | 0.12377778 | 0.122 | 0.11755556 | 0.14122222 | 0.08133333 | 0.09233333 | 0.12307778 | 0.02246143 | 0.00710293 |
| 1 | 0.13166667 | 0.15666667 | 0.14833333 | 0.18666667 | 0.146 | 0.13583333 | 0.05 | 0.13 | 0.09 | 0.102 | 0.12771667 | 0.0384173 | 0.01214862 |
| 2 | 0.11 | 0.16166667 | 0.13333333 | 0.13166667 | 0.12666667 | 0.125 | 0.05 | 0.11166667 | 0.07833333 | 0.07666667 | 0.1105 | 0.03316671 | 0.01048824 |
| 3 | 0.105 | 0.15666667 | 0.13666667 | 0.12333333 | 0.12833333 | 0.13333333 | 0.05333333 | 0.14166667 | 0.08333333 | 0.09333333 | 0.1155 | 0.03139798 | 0.00992891 |
| 4 | 0.125 | 0.13166667 | 0.14666667 | 0.12666667 | 0.125 | 0.11272727 | 0.06 | 0.13333333 | 0.09166667 | 0.10166667 | 0.11543939 | 0.02517325 | 0.00796048 |
| 5 | 0.11 | 0.13666667 | 0.15166667 | 0.14166667 | 0.12333333 | 0.12166667 | 0.06333333 | 0.15 | 0.095 | 0.11333333 | 0.12066667 | 0.02716207 | 0.0085894 |
| 10 | 0.11 | 0.13266667 | 0.12433333 | 0.13866667 | 0.13 | 0.13766667 | 0.063 | 0.16066667 | 0.09966667 | 0.10533333 | 0.1202 | 0.02701997 | 0.00854447 |
| 15 | 0.108 | 0.131 | 0.12933333 | 0.13466667 | 0.13533333 | 0.12851852 | 0.05566667 | 0.15766667 | 0.08533333 | 0.09333333 | 0.11588519 | 0.03014046 | 0.00953125 |

Het

| Time (min) | Animal ID | | | | | | | | | | | | | | | | | | | | | |
|---|---|---|---|---|---|---|---|---|---|---|---|---|---|---|---|---|---|---|---|---|---|---|
| | **B** | **D** | **E** | **F** | **G** | **H** | **155** | **157** | **159** | **160** | **162** | **190** | **196** | **199** | **201** | **205** | **207** | **208** | **210** | Average | SD | SE |
| 0 | 0.10277778 | 0.10455556 | 0.11733333 | 0.06888889 | 0.08377778 | 0.11388889 | 0.09255556 | 0.103 | 0.08788889 | 0.106 | 0.09066667 | 0.10233333 | 0.10766667 | 0.12111111 | 0.11977778 | 0.07111111 | 0.08388889 | 0.06566667 | 0.07444444 | 0.09564912 | 0.0175665 | 0.00403003 |
| 1 | 0.14833333 | 0.14 | 0.13833333 | 0.064 | 0.088 | 0.124 | 0.12 | 0.105 | 0.11666667 | 0.12 | 0.114 | 0.13166667 | 0.14666667 | 0.17166667 | 0.155 | 0.075 | 0.115 | 0.10333333 | 0.08833333 | 0.11921053 | 0.02793582 | 0.00640892 |
| 2 | 0.15166667 | 0.13166667 | 0.13166667 | 0.07666667 | 0.085 | 0.08666667 | 0.10166667 | 0.11 | 0.09 | 0.09833333 | 0.10333333 | 0.12166667 | 0.13166667 | 0.16833333 | 0.13833333 | 0.08 | 0.08833333 | 0.07 | 0.08 | 0.10763158 | 0.02806692 | 0.00643899 |
| 3 | 0.12166667 | 0.12333333 | 0.10166667 | 0.05833333 | 0.08166667 | 0.10666667 | 0.10666667 | 0.09333333 | 0.08666667 | 0.085 | 0.09 | 0.12333333 | 0.12833333 | 0.155 | 0.12666667 | 0.07666667 | 0.09166667 | 0.06833333 | 0.08166667 | 0.10035088 | 0.0244481 | 0.00560878 |
| 4 | 0.12833333 | 0.11666667 | 0.10166667 | 0.05166667 | 0.08666667 | 0.09166667 | 0.11166667 | 0.095 | 0.085 | 0.09833333 | 0.08833333 | 0.14 | 0.13 | 0.14166667 | 0.14166667 | 0.08666667 | 0.09666667 | 0.06666667 | 0.08166667 | 0.10210526 | 0.02551229 | 0.00585292 |
| 5 | 0.13166667 | 0.11666667 | 0.10666667 | 0.26166667 | 0.08666667 | 0.1 | 0.10666667 | 0.08666667 | 0.08833333 | 0.10833333 | 0.08 | 0.15 | 0.125 | 0.14 | 0.13166667 | 0.09166667 | 0.09666667 | 0.07 | 0.07833333 | 0.11350877 | 0.04228028 | 0.00969976 |
| 10 | 0.11966667 | 0.10533333 | 0.106 | 0.05033333 | 0.07966667 | 0.116 | 0.104 | 0.10133333 | 0.09233333 | 0.11033333 | 0.1 | 0.134 | 0.12533333 | 0.09733333 | 0.11433333 | 0.08466667 | 0.093 | 0.07433333 | 0.073 | 0.099 | 0.0203224 | 0.00466228 |
| 15 | 0.11033333 | 0.1 | 0.10566667 | 0.057 | 0.08966667 | 0.11633333 | 0.085 | 0.111 | 0.08766667 | 0.10933333 | 0.097 | 0.123 | 0.11766667 | 0.107 | 0.131 | 0.08133333 | 0.07233333 | 0.07166667 | 0.06866667 | 0.09692982 | 0.02044794 | 0.00469108 |

Week 2

WT

| Time (min) | Animal ID | | | | | | | | | | | | |
|---|---|---|---|---|---|---|---|---|---|---|---|---|---|
| | **83** | **86** | **87** | **88** | **81** | **82** | **A** | **156** | **158** | **161** | Average | SD | SE |
| 0 | 0.16333333 | 0.187 | 0.17711111 | 0.19777778 | 0.167 | 0.15677778 | 0.17022222 | 0.15377778 | 0.11244444 | 0.13288889 | 0.16183333 | 0.0250013 | 0.00790611 |
| 1 | 0.19666667 | 0.21833333 | 0.20333333 | 0.25 | 0.20166667 | 0.20833333 | 0.17833333 | 0.21833333 | 0.1 | 0.12833333 | 0.19033333 | 0.044644 | 0.01411767 |
| 2 | 0.13333333 | 0.23166667 | 0.20666667 | 0.27666667 | 0.2 | 0.20166667 | 0.13166667 | 0.175 | 0.10166667 | 0.11833333 | 0.17766667 | 0.05579504 | 0.01764394 |
| 3 | 0.15333333 | 0.20166667 | 0.21 | 0.255 | 0.18833333 | 0.21333333 | 0.15 | 0.16166667 | 0.10666667 | 0.13666667 | 0.17766667 | 0.04387271 | 0.01387377 |
| 4 | 0.15833333 | 0.205 | 0.22 | 0.25166667 | 0.19833333 | 0.18 | 0.175 | 0.16 | 0.10833333 | 0.13 | 0.17866667 | 0.0424395 | 0.01342055 |
| 5 | 0.16333333 | 0.19833333 | 0.21833333 | 0.20166667 | 0.17333333 | 0.17666667 | 0.16666667 | 0.16 | 0.12166667 | 0.14166667 | 0.17216667 | 0.02876093 | 0.00909501 |
| 10 | 0.16933333 | 0.18 | 0.21833333 | 0.20466667 | 0.174 | 0.181 | 0.188 | 0.189 | 0.117 | 0.125 | 0.17463333 | 0.03176241 | 0.01004416 |
| 15 | 0.181 | 0.18833333 | 0.19333333 | 0.23333333 | 0.17066667 | 0.18366667 | 0.184 | 0.18233333 | 0.11166667 | 0.11566667 | 0.1744 | 0.03606982 | 0.01140628 |

Het

| Time (min) | Animal ID | | | | | | | | | | | | | | | | | | | |
|---|---|---|---|---|---|---|---|---|---|---|---|---|---|---|---|---|---|---|---|---|
| | **B** | **D** | **E** | **G** | **H** | **155** | **157** | **160** | **162** | **190** | **196** | **199** | **201** | **205** | **207** | **208** | **210** | Average | SD | SE |
| 0 | 0.14755556 | 0.12822222 | 0.10355556 | 0.06388889 | 0.14077778 | 0.108 | 0.11488889 | 0.10111111 | 0.13755556 | 0.121 | 0.15777778 | 0.12611111 | 0.13966667 | 0.10466667 | 0.093 | 0.10155556 | 0.11033333 | 0.11762745 | 0.02327413 | 0.00564481 |
| 1 | 0.18333333 | 0.17 | 0.138 | 0.09666667 | 0.10333333 | 0.15166667 | 0.165 | 0.12 | 0.11833333 | 0.165 | 0.15666667 | 0.16666667 | 0.16333333 | 0.09166667 | 0.11833333 | 0.14333333 | 0.10166667 | 0.13841176 | 0.02961853 | 0.00718355 |
| 2 | 0.16833333 | 0.17 | 0.12666667 | 0.10333333 | 0.10666667 | 0.135 | 0.13833333 | 0.09833333 | 0.11833333 | 0.13666667 | 0.14666667 | 0.14 | 0.13166667 | 0.08666667 | 0.11333333 | 0.10833333 | 0.11 | 0.12578431 | 0.02334165 | 0.00566118 |
| 3 | 0.14666667 | 0.16833333 | 0.12833333 | 0.09666667 | 0.13666667 | 0.135 | 0.11166667 | 0.11 | 0.11333333 | 0.12166667 | 0.14333333 | 0.15 | 0.13833333 | 0.10333333 | 0.11166667 | 0.10166667 | 0.11333333 | 0.12529412 | 0.02001939 | 0.00485542 |
| 4 | 0.165 | 0.17166667 | 0.14 | 0.1 | 0.13833333 | 0.11666667 | 0.11166667 | 0.12166667 | 0.11666667 | 0.12666667 | 0.14333333 | 0.14666667 | 0.125 | 0.105 | 0.10833333 | 0.12333333 | 0.115 | 0.12794118 | 0.02027185 | 0.00491664 |
| 5 | 0.16166667 | 0.17 | 0.145 | 0.10333333 | 0.135 | 0.11166667 | 0.11 | 0.13333333 | 0.13666667 | 0.12333333 | 0.12833333 | 0.15333333 | 0.13 | 0.10333333 | 0.09666667 | 0.13 | 0.11 | 0.12833333 | 0.02109831 | 0.00511709 |
| 10 | 0.14033333 | 0.17133333 | 0.135 | 0.1 | 0.125 | 0.106 | 0.11933333 | 0.12666667 | 0.13066667 | 0.13266667 | 0.15866667 | 0.13466667 | 0.14466667 | 0.10466667 | 0.10266667 | 0.12566667 | 0.10966667 | 0.1275098 | 0.01976111 | 0.00479277 |
| 15 | 0.143 | 0.16133333 | 0.123 | 0.08266667 | 0.127 | 0.12033333 | 0.12833333 | 0.128 | 0.12666667 | 0.13533333 | 0.15766667 | 0.13133333 | 0.13833333 | 0.106 | 0.09933333 | 0.11733333 | 0.10433333 | 0.12529412 | 0.01997633 | 0.00484497 |

Week 3

WT

| Time (min) | Animal ID | | | | | | | | | | | | |
|---|---|---|---|---|---|---|---|---|---|---|---|---|---|
| | **83** | **86** | **87** | **88** | **81** | **82** | **A** | **156** | **158** | **161** | Average | SD | SE |
| 0 | 0.17488889 | 0.18011111 | 0.184 | 0.18833333 | 0.166 | 0.14766667 | 0.16811111 | 0.16566667 | 0.10911111 | 0.13 | 0.16138889 | 0.02530111 | 0.00800091 |
| 1 | 0.17833333 | 0.19666667 | 0.22333333 | 0.20833333 | 0.21333333 | 0.15666667 | 0.2 | 0.19333333 | 0.125 | 0.13166667 | 0.18266667 | 0.03413119 | 0.01079323 |
| 2 | 0.165 | 0.19166667 | 0.17 | 0.19333333 | 0.19333333 | 0.145 | 0.14833333 | 0.14333333 | 0.10166667 | 0.10666667 | 0.15583333 | 0.03343734 | 0.01057381 |
| 3 | 0.185 | 0.18833333 | 0.15333333 | 0.19833333 | 0.215 | 0.145 | 0.18 | 0.15833333 | 0.1 | 0.12833333 | 0.16516667 | 0.03484932 | 0.01102032 |
| 4 | 0.18666667 | 0.18666667 | 0.16666667 | 0.175 | 0.22166667 | 0.145 | 0.15666667 | 0.16333333 | 0.11 | 0.14333333 | 0.1655 | 0.03021579 | 0.00955507 |
| 5 | 0.15333333 | 0.17 | 0.15 | 0.19166667 | 0.21333333 | 0.15833333 | 0.16666667 | 0.17166667 | 0.10833333 | 0.14666667 | 0.163 | 0.02795278 | 0.00883945 |
| 10 | 0.16433333 | 0.19 | 0.188 | 0.18033333 | 0.17366667 | 0.177 | 0.17 | 0.19566667 | 0.12833333 | 0.13366667 | 0.1701 | 0.02270642 | 0.0071804 |
| 15 | 0.17466667 | 0.175 | 0.16466667 | 0.18566667 | 0.19 | 0.174 | 0.14833333 | 0.204 | 0.12233333 | 0.11833333 | 0.1657 | 0.02812426 | 0.00889367 |

Het

| Time (min) | Animal ID | | | | | | | | | | | | | | | | |
|---|---|---|---|---|---|---|---|---|---|---|---|---|---|---|---|---|---|
| | **B** | **D** | **E** | **H** | **155** | **157** | **190** | **196** | **199** | **201** | **205** | **207** | **208** | **210** | Average | SD | SE |
| 0 | 0.14866667 | 0.14333333 | 0.10122222 | 0.12377778 | 0.12422222 | 0.13377778 | 0.12877778 | 0.13155556 | 0.14344444 | 0.14644444 | 0.08333333 | 0.09588889 | 0.09133333 | 0.1 | 0.12112698 | 0.02241234 | 0.00598995 |
| 1 | 0.198 | 0.18 | 0.15 | 0.14333333 | 0.175 | 0.16333333 | 0.17 | 0.2 | 0.19333333 | 0.17833333 | 0.13 | 0.11833333 | 0.10333333 | 0.11666667 | 0.15854762 | 0.03198627 | 0.00854869 |
| 2 | 0.18833333 | 0.15666667 | 0.14666667 | 0.11166667 | 0.14 | 0.15833333 | 0.13333333 | 0.16 | 0.18 | 0.17 | 0.10166667 | 0.11166667 | 0.10833333 | 0.105 | 0.14083333 | 0.0293938 | 0.00785582 |
| 3 | 0.18 | 0.14333333 | 0.16 | 0.13666667 | 0.13333333 | 0.15 | 0.13166667 | 0.16 | 0.16 | 0.15333333 | 0.11166667 | 0.10666667 | 0.11666667 | 0.095 | 0.13845238 | 0.02427992 | 0.00648908 |
| 4 | 0.16166667 | 0.16666667 | 0.16833333 | 0.14333333 | 0.13 | 0.125 | 0.14833333 | 0.14666667 | 0.14833333 | 0.15833333 | 0.11 | 0.10833333 | 0.11333333 | 0.10666667 | 0.13821429 | 0.02239376 | 0.00598498 |
| 5 | 0.165 | 0.13666667 | 0.17833333 | 0.12666667 | 0.12166667 | 0.13166667 | 0.14333333 | 0.11833333 | 0.15666667 | 0.14333333 | 0.11 | 0.095 | 0.11666667 | 0.11 | 0.13238095 | 0.02322975 | 0.00620841 |
| 10 | 0.17233333 | 0.15466667 | 0.14466667 | 0.16033333 | 0.151 | 0.15066667 | 0.14633333 | 0.14866667 | 0.145 | 0.16833333 | 0.117 | 0.10333333 | 0.11466667 | 0.119 | 0.14257143 | 0.0210077 | 0.00561454 |
| 15 | 0.17066667 | 0.146 | 0.132 | 0.15333333 | 0.151 | 0.15066667 | 0.14333333 | 0.16466667 | 0.16066667 | 0.16233333 | 0.115 | 0.104 | 0.10566667 | 0.11666667 | 0.14114286 | 0.02257361 | 0.00603305 |

Week 4

WT

| Time (min) | Animal ID | | | | | | | | | | | | |
|---|---|---|---|---|---|---|---|---|---|---|---|---|---|
| | **83** | **86** | **87** | **88** | **81** | **82** | **A** | **156** | **158** | **161** | Average | SD | SE |
| 0 | 0.194 | 0.17755556 | 0.18333333 | 0.21 | 0.17655556 | 0.175 | 0.19377778 | 0.19544444 | 0.123 | 0.14711111 | 0.17757778 | 0.02549579 | 0.00806248 |
| 1 | 0.21166667 | 0.19833333 | 0.25 | 0.23833333 | 0.20833333 | 0.17 | 0.18166667 | 0.185 | 0.13666667 | 0.13 | 0.191 | 0.03907258 | 0.01235584 |
| 2 | 0.15833333 | 0.20666667 | 0.22333333 | 0.22666667 | 0.19333333 | 0.15 | 0.165 | 0.17666667 | 0.10333333 | 0.11666667 | 0.172 | 0.04189788 | 0.01324927 |
| 3 | 0.16666667 | 0.19833333 | 0.23 | 0.23 | 0.185 | 0.16 | 0.14833333 | 0.17833333 | 0.10666667 | 0.14333333 | 0.17466667 | 0.03856052 | 0.01219391 |
| 4 | 0.17666667 | 0.23333333 | 0.24333333 | 0.24666667 | 0.225 | 0.16 | 0.16 | 0.19166667 | 0.125 | 0.15666667 | 0.19183333 | 0.04274133 | 0.01351599 |
| 5 | 0.17833333 | 0.22166667 | 0.21833333 | 0.22 | 0.22833333 | 0.17666667 | 0.14666667 | 0.19833333 | 0.12666667 | 0.13166667 | 0.18466667 | 0.03882375 | 0.01227715 |
| 10 | 0.19533333 | 0.20066667 | 0.21033333 | 0.21733333 | 0.20033333 | 0.18633333 | 0.18166667 | 0.21 | 0.14233333 | 0.143 | 0.18873333 | 0.026578 | 0.0084047 |
| 15 | 0.182 | 0.169 | 0.20466667 | 0.174 | 0.204 | 0.181 | 0.18733333 | 0.188 | 0.12466667 | 0.13433333 | 0.1749 | 0.02656511 | 0.00840062 |

Het

| Time (min) | Animal ID | | | | | | | | | | | | | | | |
|---|---|---|---|---|---|---|---|---|---|---|---|---|---|---|---|---|
| | **B** | **E** | **H** | **155** | **157** | **190** | **196** | **199** | **201** | **205** | **207** | **208** | **210** | Average | SD | SE |
| 0 | 0.17277778 | 0.11166667 | 0.13888889 | 0.14255556 | 0.17344444 | 0.15133333 | 0.14677778 | 0.13988889 | 0.14555556 | 0.10955556 | 0.11111111 | 0.11466667 | 0.10844444 | 0.13589744 | 0.02307393 | 0.00639956 |
| 1 | 0.24666667 | 0.16833333 | 0.15166667 | 0.18166667 | 0.155 | 0.18333333 | 0.17 | 0.18833333 | 0.22 | 0.145 | 0.125 | 0.12666667 | 0.12166667 | 0.16794872 | 0.03696942 | 0.01025347 |
| 2 | 0.19333333 | 0.17333333 | 0.12666667 | 0.125 | 0.12 | 0.14166667 | 0.14666667 | 0.165 | 0.185 | 0.12 | 0.10166667 | 0.11166667 | 0.11666667 | 0.14051282 | 0.02981723 | 0.00826981 |
| 3 | 0.22833333 | 0.19 | 0.14666667 | 0.11166667 | 0.125 | 0.13333333 | 0.14666667 | 0.155 | 0.16166667 | 0.12166667 | 0.10166667 | 0.11833333 | 0.12333333 | 0.14333333 | 0.03484091 | 0.00966313 |
| 4 | 0.23166667 | 0.16 | 0.12333333 | 0.12666667 | 0.13833333 | 0.145 | 0.13333333 | 0.15666667 | 0.18 | 0.13333333 | 0.12 | 0.10833333 | 0.125 | 0.14474359 | 0.03238831 | 0.0089829 |
| 5 | 0.215 | 0.145 | 0.13166667 | 0.13833333 | 0.13 | 0.17 | 0.15666667 | 0.14 | 0.17666667 | 0.13 | 0.1 | 0.11 | 0.125 | 0.14371795 | 0.03026239 | 0.00839328 |
| 10 | 0.20166667 | 0.13866667 | 0.145 | 0.14 | 0.17133333 | 0.15466667 | 0.15566667 | 0.14466667 | 0.154 | 0.143 | 0.10466667 | 0.113 | 0.12166667 | 0.14523077 | 0.02492744 | 0.00691363 |
| 15 | 0.20966667 | 0.13433333 | 0.13833333 | 0.141 | 0.17 | 0.15566667 | 0.16033333 | 0.17066667 | 0.15033333 | 0.15633333 | 0.10166667 | 0.11344828 | 0.11566667 | 0.14749602 | 0.0285749 | 0.00792525 |

### Frequency

Week 1

WT

| Time (min) | 83 | 86 | 87 | 88 | 81 | 82 | A | 156 | 158 | 161 | Average | SD | SE |
|---|---|---|---|---|---|---|---|---|---|---|---|---|---|
| 0 | 405.333333 | 352.766667 | 296.433333 | 376.644444 | 306.488889 | 300.911111 | 342.277778 | 374.1 | 288.633333 | 274.422222 | 331.801111 | 44.4197124 | 14.0467464 |
| 1 | 389.166667 | 445.333333 | 373.666667 | 416.333333 | 367.6 | 399.916667 | 332.5 | 487.4 | 324.8 | 370.8 | 390.751667 | 49.4748361 | 15.6453169 |
| 2 | 472.333333 | 384 | 419.5 | 355 | 361.333333 | 364.25 | 456.333333 | 501.666667 | 319.5 | 394.833333 | 402.875 | 58.3098795 | 18.4392029 |
| 3 | 495.833333 | 377.666667 | 426.5 | 288.666667 | 330.5 | 357.5 | 584.666667 | 383.333333 | 374.666667 | 379.333333 | 399.866667 | 84.8953168 | 26.8462564 |
| 4 | 447 | 436.666667 | 498.5 | 441 | 310 | 384 | 489 | 590.666667 | 419.833333 | 374.666667 | 439.133333 | 76.9476253 | 24.3329756 |
| 5 | 423.666667 | 442.833333 | 349.666667 | 311.5 | 298 | 438.5 | 393.333333 | 410.333333 | 361 | 374.333333 | 380.316667 | 50.666024 | 16.0220036 |
| 10 | 290.166667 | 359.766667 | 287.3 | 296.966667 | 309 | 317.633333 | 334.3 | 363.866667 | 386.333333 | 329.7 | 327.503333 | 33.7602932 | 10.6759421 |
| 15 | 291.533333 | 364.8 | 297.566667 | 262.066667 | 308.7 | 274.777778 | 263.7 | 361.7 | 266.3 | 282.866667 | 297.401111 | 37.8795059 | 11.9785515 |

Het

| Time (min) | B | D | E | F | G | H | 155 | 157 | 159 | 160 | 162 | 190 | 196 | 199 | 201 | 205 | 207 | 208 | 210 | Average | SD | SE |
|---|---|---|---|---|---|---|---|---|---|---|---|---|---|---|---|---|---|---|---|---|---|---|
| 0 | 241 | 287.222222 | 317.088889 | 204.288889 | 282.955556 | 368.433333 | 442.122222 | 455.544444 | 262.055556 | 419.644444 | 368.411111 | 504.633333 | 235.744444 | 233.222222 | 348.522222 | 496.966667 | 330.244444 | 266.244444 | 213.955556 | 330.436842 | 96.1385905 | 22.0557053 |
| 1 | 359.166667 | 441.4 | 412.833333 | 303.4 | 392.4 | 348.6 | 424.8 | 428 | 353.166667 | 328 | 388.2 | 378.833333 | 369.5 | 348.166667 | 359.666667 | 416 | 355.166667 | 407.333333 | 330.666667 | 376.068421 | 38.3187758 | 8.79093007 |
| 2 | 248.833333 | 393.666667 | 270.333333 | 289.166667 | 351.833333 | 459.5 | 380.166667 | 371.333333 | 348.833333 | 320.5 | 314 | 338.666667 | 361.333333 | 299.666667 | 399.166667 | 407.166667 | 335.833333 | 313.166667 | 297.333333 | 342.131579 | 52.3587335 | 12.0119173 |
| 3 | 243.833333 | 456.833333 | 224.166667 | 209.333333 | 312.5 | 383 | 449 | 366 | 306.833333 | 296.5 | 298.666667 | 387.166667 | 354.833333 | 288.833333 | 360.666667 | 354.333333 | 348 | 284.333333 | 255.666667 | 325.289474 | 68.7631882 | 15.7753573 |
| 4 | 251.666667 | 551 | 195.333333 | 173.333333 | 302.166667 | 340 | 473.5 | 290.166667 | 291.5 | 337.166667 | 301.166667 | 431.166667 | 379.833333 | 398.333333 | 395.333333 | 370.333333 | 379.5 | 298.833333 | 280.5 | 338.991228 | 91.4979841 | 20.9910772 |
| 5 | 200.333333 | 529.833333 | 204.833333 | 167.5 | 325.666667 | 328.666667 | 390.5 | 266.833333 | 349.666667 | 400.5 | 300.333333 | 438.333333 | 430.833333 | 449.666667 | 519.5 | 426.333333 | 364.666667 | 325 | 275.833333 | 352.359649 | 102.416185 | 23.4958842 |
| 10 | 320.366667 | 507.333333 | 253.166667 | 209.833333 | 329.633333 | 342.833333 | 392.133333 | 290.233333 | 311.766667 | 414.066667 | 346.033333 | 447.666667 | 262.1 | 550.566667 | 455.1 | 411.866667 | 326.266667 | 336.666667 | 254.9 | 355.922807 | 90.5339136 | 20.7699042 |
| 15 | 283.166667 | 463.033333 | 276.2 | 259.4 | 341.766667 | 318.233333 | 269.633333 | 299.666667 | 263.633333 | 365.8 | 246.4 | 393.633333 | 251.4 | 469.433333 | 327.066667 | 294.666667 | 260.733333 | 275.8 | 254.9 | 311.292982 | 67.8800436 | 15.57275 |

Week 2

WT

| Time (min) | 83 | 86 | 87 | 88 | 81 | 82 | A | 156 | 158 | 161 | Average | SD | SE |
|---|---|---|---|---|---|---|---|---|---|---|---|---|---|
| 0 | 447.1 | 400.9 | 306.833333 | 428.5 | 389.088889 | 261.955556 | 382.9 | 292.077778 | 432.5 | 396.633333 | 373.848889 | 64.1657689 | 20.2909978 |
| 1 | 425.333333 | 468.5 | 475.5 | 486.833333 | 460.666667 | 472.166667 | 414.666667 | 279.166667 | 361.666667 | 414.166667 | 425.866667 | 64.3409994 | 20.3464105 |
| 2 | 477.666667 | 465.666667 | 482.666667 | 451.166667 | 436.333333 | 507.5 | 428.666667 | 458.5 | 361.333333 | 458.166667 | 452.766667 | 39.3724068 | 12.4506483 |
| 3 | 435.5 | 492.833333 | 457.5 | 438.333333 | 414.333333 | 422.666667 | 366 | 396.166667 | 370 | 417.666667 | 421.1 | 38.4475111 | 12.1581705 |
| 4 | 373.5 | 480 | 464.333333 | 389.5 | 431.666667 | 447.166667 | 376 | 360.166667 | 392.333333 | 575.5 | 429.016667 | 65.9804708 | 20.8648569 |
| 5 | 481.333333 | 481 | 376 | 434.5 | 434.166667 | 424.166667 | 336.833333 | 271.833333 | 341.666667 | 480 | 406.15 | 71.9065945 | 22.7388617 |
| 10 | 368.2 | 429.166667 | 316.333333 | 435.966667 | 309.666667 | 297.6 | 306.133333 | 252.433333 | 284.033333 | 362.766667 | 336.23 | 61.0617628 | 19.3094248 |
| 15 | 346.5 | 323.833333 | 272.766667 | 348.666667 | 257.8 | 269.433333 | 317.1 | 238.1 | 379.266667 | 281.1 | 303.456667 | 46.188607 | 14.60612 |

Het

| Time (min) | B | D | E | G | H | 155 | 157 | 160 | 162 | 190 | 196 | 199 | 201 | 205 | 207 | 208 | 210 | Average | SD | SE |
|---|---|---|---|---|---|---|---|---|---|---|---|---|---|---|---|---|---|---|---|---|
| 0 | 377.566667 | 256.522222 | 286.355556 | 285.877778 | 355.633333 | 288.4 | 398.211111 | 359.888889 | 271.244444 | 279.788889 | 226.788889 | 216.222222 | 267.066667 | 449.611111 | 285.111111 | 512.777778 | 369.555556 | 322.742484 | 80.3809572 | 19.4952457 |
| 1 | 406.166667 | 377.166667 | 427.2 | 359 | 351.666667 | 397.166667 | 348.833333 | 351.833333 | 381.166667 | 448.833333 | 400.833333 | 327.166667 | 368.166667 | 396.333333 | 429.5 | 400.166667 | 463.5 | 390.276471 | 37.6075417 | 9.12116862 |
| 2 | 342.333333 | 395 | 400.166667 | 333.166667 | 463.333333 | 338.5 | 277 | 442.666667 | 416.833333 | 493.833333 | 497.166667 | 355.666667 | 492.166667 | 396 | 514.666667 | 371 | 495.333333 | 413.22549 | 71.4448588 | 17.3279235 |
| 3 | 295.166667 | 380.166667 | 393.166667 | 348.333333 | 429.833333 | 313.166667 | 251.5 | 380.833333 | 389.166667 | 589.166667 | 577.166667 | 357.166667 | 561.5 | 420 | 579.833333 | 301.166667 | 456.666667 | 413.176471 | 106.923803 | 25.9328313 |
| 4 | 441.333333 | 384.833333 | 437.333333 | 350.166667 | 383.5 | 307.166667 | 238.833333 | 393.333333 | 327.166667 | 532.5 | 487.833333 | 492.5 | 563.666667 | 378.666667 | 499.833333 | 390.5 | 491.5 | 417.686275 | 86.6784023 | 21.0226005 |
| 5 | 499 | 459.333333 | 377.166667 | 405.666667 | 320.333333 | 309 | 236 | 351.666667 | 241 | 512.5 | 438.5 | 411.833333 | 453.333333 | 359.833333 | 530.5 | 467 | 388.333333 | 397.705882 | 87.7916672 | 21.2926069 |
| 10 | 276.366667 | 402.533333 | 365.666667 | 356.333333 | 318.533333 | 302.1 | 292.066667 | 289.6 | 239.033333 | 386.733333 | 267.733333 | 377.233333 | 285.366667 | 327.133333 | 379.5 | 394.066667 | 383.666667 | 331.980392 | 51.8739311 | 12.5812763 |
| 15 | 331.3 | 354.966667 | 299.166667 | 316.666667 | 284.133333 | 337.633333 | 267.933333 | 191.733333 | 125.7 | 364.6 | 359.866667 | 354.166667 | 339.733333 | 300.733333 | 235.433333 | 345.866667 | 336.766667 | 302.729412 | 65.5963878 | 15.9094609 |

Week 3

WT

| Time (min) | 83 | 86 | 87 | 88 | 81 | 82 | A | 156 | 158 | 161 | Average | SD | SE |
|---|---|---|---|---|---|---|---|---|---|---|---|---|---|
| 0 | 443.422222 | 413.466667 | 340.955556 | 347.922222 | 254.7 | 272.633333 | 420.644444 | 320.666667 | 263.011111 | 374.244444 | 345.166667 | 68.0537944 | 21.5204994 |
| 1 | 527 | 525.333333 | 575.333333 | 534.333333 | 527.333333 | 552.333333 | 503.5 | 435.666667 | 331 | 391.5 | 490.333333 | 78.3584278 | 24.7791106 |
| 2 | 612.166667 | 453.666667 | 581.833333 | 525 | 504 | 411 | 485.333333 | 364.166667 | 274.166667 | 399.333333 | 461.066667 | 102.539459 | 32.4258242 |
| 3 | 597.333333 | 419 | 678.333333 | 577.666667 | 510 | 410.333333 | 450.5 | 327.5 | 269.166667 | 487.166667 | 472.7 | 124.816354 | 39.4703968 |
| 4 | 458.666667 | 377.5 | 573.333333 | 636.166667 | 565.666667 | 425.333333 | 536.333333 | 337 | 259.833333 | 495.166667 | 466.5 | 117.799762 | 37.2515556 |
| 5 | 512.166667 | 421 | 548.5 | 489.666667 | 527.333333 | 454.5 | 520.5 | 266.5 | 363.833333 | 414 | 451.8 | 87.7577966 | 27.751452 |
| 10 | 501.466667 | 289.566667 | 363.733333 | 300.066667 | 445.233333 | 340 | 336.666667 | 274.5 | 296.966667 | 352.733333 | 350.093333 | 72.3752683 | 22.8870694 |
| 15 | 439.7 | 251.166667 | 238.1 | 285.466667 | 237.4 | 250.866667 | 249.233333 | 305.2 | 263.6 | 248.5 | 276.923333 | 61.0397281 | 19.3024569 |

Het

| Time (min) | B | D | E | H | 155 | 157 | 190 | 196 | 199 | 201 | 205 | 207 | 208 | 210 | Average | SD | SE |
|---|---|---|---|---|---|---|---|---|---|---|---|---|---|---|---|---|---|
| 0 | 299.866667 | 242.911111 | 259.966667 | 258.177778 | 266 | 264.522222 | 278.555556 | 260.344444 | 244.955556 | 254.422222 | 294.133333 | 268.244444 | 268.244444 | 265.455556 | 266.128571 | 16.0479698 | 4.28900035 |
| 1 | 439.6 | 424.833333 | 362.2 | 491.833333 | 367.666667 | 376 | 392.833333 | 433 | 369.5 | 349 | 356 | 395.666667 | 374 | 470.5 | 400.188095 | 44.6702823 | 11.9386351 |
| 2 | 509.833333 | 471.5 | 368.166667 | 507 | 283.166667 | 294.166667 | 515.833333 | 531.333333 | 331.333333 | 303.5 | 332.166667 | 463.5 | 389.666667 | 484.833333 | 413.285714 | 93.0871393 | 24.8785845 |
| 3 | 583.666667 | 604.833333 | 326.666667 | 511.666667 | 259 | 272.166667 | 551 | 586.666667 | 354.333333 | 496.833333 | 328 | 582 | 299.666667 | 551.666667 | 450.583333 | 134.285411 | 35.8892857 |
| 4 | 602.833333 | 560.666667 | 435.166667 | 526.166667 | 261.5 | 515.333333 | 506.666667 | 575 | 458 | 517.833333 | 312.5 | 499 | 299.833333 | 465 | 466.821429 | 105.629806 | 28.230753 |
| 5 | 393 | 562.166667 | 506.666667 | 526.5 | 261 | 480.833333 | 436.333333 | 480 | 419.5 | 472.5 | 293.833333 | 503.166667 | 418.833333 | 326.666667 | 434.357143 | 89.4685605 | 23.9114786 |
| 10 | 429.5 | 354.833333 | 331.8 | 318.6 | 340.266667 | 329.733333 | 319.366667 | 341.866667 | 408.066667 | 352.9 | 363.633333 | 330.066667 | 415.633333 | 291.533333 | 351.985714 | 40.0048794 | 10.6917538 |
| 15 | 375.633333 | 211.266667 | 251.533333 | 279.3 | 287.833333 | 373.033333 | 276 | 320.5 | 278.133333 | 361.133333 | 374.333333 | 254.433333 | 236.033333 | 234.9 | 293.861905 | 57.1564233 | 15.2756967 |

Week 4

WT

| Time (min) | 83 | 86 | 87 | 88 | 81 | 82 | A | 156 | 158 | 161 | Average | SD | SE |
|---|---|---|---|---|---|---|---|---|---|---|---|---|---|
| 0 | 450.888889 | 294.944444 | 352.288889 | 381.111111 | 229.277778 | 320.133333 | 372.6 | 360.633333 | 241.577778 | 363.577778 | 336.703333 | 67.0358476 | 21.1985963 |
| 1 | 476.833333 | 542.833333 | 473.166667 | 520.333333 | 459.333333 | 495 | 550.666667 | 555 | 386.666667 | 470.5 | 493.033333 | 51.6452644 | 16.3316666 |
| 2 | 505.666667 | 457 | 540.666667 | 513.166667 | 432.166667 | 438.833333 | 533.5 | 439.666667 | 255.166667 | 399.333333 | 451.516667 | 83.9206875 | 26.5380515 |
| 3 | 532 | 502 | 576.666667 | 568.833333 | 421.666667 | 415.666667 | 619.833333 | 459 | 318.166667 | 485.333333 | 489.916667 | 90.1817866 | 28.5179849 |
| 4 | 548.166667 | 511.5 | 530.833333 | 534.166667 | 390.833333 | 434.5 | 592.5 | 478 | 352.333333 | 505.5 | 487.833333 | 74.6648892 | 23.6111111 |
| 5 | 505 | 515.666667 | 484 | 490.5 | 471.166667 | 441.5 | 550.166667 | 462.666667 | 374.666667 | 451 | 474.633333 | 47.6742936 | 15.0759354 |
| 10 | 353.766667 | 364.666667 | 379.766667 | 359.566667 | 392.6 | 339.3 | 341.933333 | 413.466667 | 335.266667 | 389.6 | 366.993333 | 26.0911025 | 8.25073106 |
| 15 | 270.766667 | 252.533333 | 266.5 | 243.566667 | 272.3 | 256.666667 | 264.366667 | 218.133333 | 263.733333 | 242.333333 | 255.09 | 16.6774339 | 5.27386767 |

Het

| Time (min) | B | E | H | 155 | 157 | 190 | 196 | 199 | 201 | 205 | 207 | 208 | 210 | Average | SD | SE |
|---|---|---|---|---|---|---|---|---|---|---|---|---|---|---|---|---|
| 0 | 278.322222 | 275.788889 | 240.277778 | 272.244444 | 224.922222 | 330.177778 | 236.855556 | 349.622222 | 349.988889 | 276.377778 | 403.411111 | 359.522222 | 356.955556 | 304.189744 | 56.7887611 | 15.7503685 |
| 1 | 421.166667 | 385.5 | 473.166667 | 379.333333 | 368 | 401.5 | 448 | 357.166667 | 345.333333 | 366.833333 | 361.333333 | 352.333333 | 389.833333 | 388.423077 | 38.5361552 | 10.6880064 |
| 2 | 500 | 357 | 552.666667 | 449.5 | 409.666667 | 416.166667 | 543.333333 | 320.166667 | 355.5 | 389.5 | 508.833333 | 346.833333 | 415.166667 | 428.025641 | 77.2766748 | 21.4326933 |
| 3 | 592.833333 | 445.166667 | 557.666667 | 496.333333 | 475.333333 | 604.5 | 621.833333 | 318.666667 | 372.666667 | 554.333333 | 565.666667 | 374.833333 | 370.166667 | 488.461538 | 103.476271 | 28.6991539 |
| 4 | 559.833333 | 442 | 586.5 | 436.666667 | 419.333333 | 500 | 622.333333 | 585.833333 | 526 | 557.833333 | 566.666667 | 403.833333 | 559 | 520.448718 | 72.4773076 | 20.1015884 |
| 5 | 530 | 290.666667 | 516 | 260.833333 | 510.5 | 357.5 | 497.666667 | 550.333333 | 533.833333 | 468.833333 | 520.833333 | 514.666667 | 395.5 | 457.474359 | 98.0256686 | 27.1874288 |
| 10 | 419.466667 | 273.4 | 353.033333 | 247.066667 | 335.033333 | 384.6 | 449.6 | 420.566667 | 465.166667 | 339.3 | 255.233333 | 345.233333 | 224.8 | 347.115385 | 79.3551016 | 22.0091452 |
| 15 | 405.066667 | 306.733333 | 211.766667 | 257.8 | 209.266667 | 231.1 | 450.433333 | 231 | 394.433333 | 421.5 | 227.9 | 272.724138 | 224.566667 | 295.714677 | 89.4869552 | 24.8192158 |

### End inspiratory pauase

Week 1

WT

| Time (min) | Animal ID | | | | | | | | | | | |
|---|---|---|---|---|---|---|---|---|---|---|---|---|
| | **83** | **86** | **87** | **88** | **81** | **82** | **A** | **156** | **158** | **161** | Average | SD | SE |
| 0 | 3.94444444 | 2.7 | 2.08888889 | 2.72222222 | 2.08888889 | 2.07777778 | 1.58888889 | 2.9 | 1.45555556 | 1.12222222 | 2.26888889 | 0.82578142 | 0.26113501 |
| 1 | 2.33333333 | 2 | 1.66666667 | 2 | 1.6 | 2.25 | 1 | 1.8 | 1 | 1 | 1.665 | 0.51234453 | 0.16201756 |
| 2 | 2 | 2 | 1.83333333 | 2 | 1.66666667 | 2.08333333 | 1 | 2 | 1 | 1 | 1.65833333 | 0.46886058 | 0.14826673 |
| 3 | 2 | 2.16666667 | 2 | 2 | 1.16666667 | 2.08333333 | 1 | 1.66666667 | 1 | 1 | 1.60833333 | 0.50621142 | 0.16007811 |
| 4 | 2.66666667 | 2.5 | 2 | 2.33333333 | 2 | 2.36363636 | 1.33333333 | 1.5 | 1 | 1 | 1.86969697 | 0.62025466 | 0.19614175 |
| 5 | 2.5 | 3 | 2.66666667 | 2 | 2 | 2.83333333 | 1 | 2.16666667 | 1 | 1 | 2.01666667 | 0.77559217 | 0.24526378 |
| 10 | 1.86666667 | 3.7 | 2.06666667 | 2.03333333 | 2.16666667 | 2.83333333 | 1.16666667 | 1.76666667 | 1.63333333 | 1.2 | 2.04333333 | 0.7557271 | 0.23898189 |
| 15 | 1.96666667 | 2.7 | 2.03333333 | 2 | 2.2 | 2.25925926 | 1.03333333 | 2.23333333 | 1 | 1.03333333 | 1.84592593 | 0.60418749 | 0.19106086 |

Het

| Time (min) | Animal ID | | | | | | | | | | | | | | | | | | | | |
|---|---|---|---|---|---|---|---|---|---|---|---|---|---|---|---|---|---|---|---|---|---|
| | **B** | **D** | **E** | **F** | **G** | **H** | **155** | **157** | **159** | **160** | **162** | **190** | **196** | **199** | **201** | **205** | **207** | **208** | **210** | Average | SD | SE |
| 0 | 2.53333333 | 1.95555556 | 2.76666667 | 4 | 1.88888889 | 1.72222222 | 3.23333333 | 4.06666667 | 1.76666667 | 4.5 | 1.62222222 | 3.41111111 | 2.2 | 2.62222222 | 4.05555556 | 1.8 | 2.04444444 | 1.14444444 | 1.84444444 | 2.58830409 | 1.001915 | 0.22985506 |
| 1 | 2.33333333 | 1.8 | 3.16666667 | 5 | 1.8 | 1.2 | 1.4 | 1.33333333 | 3 | 2.6 | 1.8 | 1.83333333 | 1.83333333 | 2 | 2.66666667 | 1.16666667 | 2 | 3 | 2.83333333 | 2.25087719 | 0.91559658 | 0.21005226 |
| 2 | 1.83333333 | 2.16666667 | 2 | 3 | 1.16666667 | 1.16666667 | 1.83333333 | 2.16666667 | 2 | 2 | 1 | 1.5 | 1.5 | 2.83333333 | 2.66666667 | 1 | 1 | 1 | 1.83333333 | 1.77192982 | 0.63150449 | 0.14487707 |
| 3 | 2 | 2.33333333 | 2.66666667 | 6.33333333 | 1 | 1.16666667 | 2.33333333 | 2.16666667 | 1 | 2 | 1 | 1.66666667 | 2 | 3 | 2.16666667 | 1 | 1 | 1 | 1.66666667 | 1.97368421 | 1.2338627 | 0.28306752 |
| 4 | 1.66666667 | 1.66666667 | 2.33333333 | 8.66666667 | 1.16666667 | 1 | 2.33333333 | 2.33333333 | 1 | 2.5 | 1 | 2.16666667 | 2 | 3 | 3.33333333 | 1.5 | 1 | 1 | 1.33333333 | 2.15789474 | 1.73158181 | 0.39725211 |
| 5 | 1.5 | 1.5 | 2.33333333 | 27.5 | 1 | 1 | 2.5 | 2.5 | 1 | 2.5 | 1 | 2 | 2.5 | 2.66666667 | 2.16666667 | 1.33333333 | 1 | 1 | 1.16666667 | 3.06140351 | 5.95383747 | 1.36590399 |
| 10 | 2.3 | 2.9 | 2.06666667 | 4.16666667 | 1.1 | 1.36666667 | 2.86666667 | 2.3 | 1.1 | 3.63333333 | 1.23333333 | 2.4 | 2.06666667 | 2.96666667 | 3.33333333 | 2.16666667 | 1.36666667 | 1.13333333 | 1.36666667 | 2.20175439 | 0.92499407 | 0.21220819 |
| 15 | 2.76666667 | 3.2 | 2.1 | 2.83333333 | 1.66666667 | 1.2 | 2.43333333 | 3.26666667 | 1.03333333 | 4.2 | 1.13333333 | 3.6 | 2.16666667 | 3.8 | 3.33333333 | 1.33333333 | 1.16666667 | 1.3 | 1.56666667 | 2.32105263 | 1.03453995 | 0.23733974 |

Week 2

WT

| Time (min) | Animal ID | | | | | | | | | | | | |
|---|---|---|---|---|---|---|---|---|---|---|---|---|---|
| | **83** | **86** | **87** | **88** | **81** | **82** | **A** | **156** | **158** | **161** | Average | SD | SE |
| 0 | 4.63333333 | 2.66666667 | 1.31111111 | 2.14444444 | 2.17777778 | 2.1 | 2.6 | 2.34444444 | 1.67777778 | 1.52222222 | 2.31777778 | 0.92597333 | 0.29281848 |
| 1 | 1.66666667 | 2.33333333 | 1.33333333 | 1.83333333 | 1 | 3 | 1.16666667 | 1 | 1.33333333 | 1 | 1.56666667 | 0.66295262 | 0.20964403 |
| 2 | 1.83333333 | 1.83333333 | 1 | 2 | 1.16666667 | 2.16666667 | 1.33333333 | 1 | 1.33333333 | 1.16666667 | 1.48333333 | 0.43354696 | 0.13709959 |
| 3 | 2 | 2.16666667 | 1 | 2.16666667 | 1.83333333 | 3.16666667 | 1.16666667 | 1 | 1 | 1.16666667 | 1.66666667 | 0.72435582 | 0.22906142 |
| 4 | 2 | 2 | 1.16666667 | 2.16666667 | 1.66666667 | 2.5 | 1 | 1 | 1.5 | 1 | 1.6 | 0.55109319 | 0.17427097 |
| 5 | 2.16666667 | 2.33333333 | 1.125 | 2 | 2.16666667 | 4.16666667 | 1 | 1 | 1 | 1 | 1.79583333 | 1.01159367 | 0.31989401 |
| 10 | 2.9 | 3.33333333 | 1.16666667 | 2.43333333 | 1.96666667 | 2.8 | 1.46666667 | 1.03333333 | 1 | 1.2 | 1.93 | 0.87692645 | 0.27730849 |
| 15 | 3 | 2.5 | 1.9 | 2.56666667 | 1.86666667 | 2.4 | 1.8 | 1.2 | 1.26666667 | 1.03333333 | 1.95333333 | 0.65737976 | 0.20788173 |

Het

| Time (min) | Animal ID | | | | | | | | | | | | | | | | | | | |
|---|---|---|---|---|---|---|---|---|---|---|---|---|---|---|---|---|---|---|---|---|
| | **B** | **D** | **E** | **G** | **H** | **155** | **157** | **160** | **162** | **190** | **196** | **199** | **201** | **205** | **207** | **208** | **210** | Average | SD | SE |
| 0 | 2.58888889 | 1.5 | 1.65555556 | 1.24444444 | 1.33333333 | 1.96666667 | 3.62222222 | 5.51111111 | 4.42222222 | 2.02222222 | 1.81111111 | 2.96666667 | 2.7 | 1.7 | 1.8 | 1.92222222 | 1.3 | 2.35686275 | 1.1853327 | 0.28748541 |
| 1 | 1.5 | 1.16666667 | 1.4 | 1.66666667 | 0.83333333 | 1 | 2.16666667 | 2.5 | 1.66666667 | 1.33333333 | 1.5 | 2 | 1.83333333 | 1 | 1.16666667 | 2.66666667 | 1 | 1.55294118 | 0.53787891 | 0.1304548 |
| 2 | 1.16666667 | 2 | 1 | 2.16666667 | 1 | 1 | 1.83333333 | 2.5 | 1.33333333 | 1.83333333 | 1.66666667 | 2.5 | 2.5 | 1 | 1.16666667 | 1.33333333 | 1.16666667 | 1.59803922 | 0.56843726 | 0.13786629 |
| 3 | 1 | 1.33333333 | 1 | 2 | 1 | 1 | 2 | 2.16666667 | 1.16666667 | 1.5 | 1.66666667 | 2.16666667 | 1.66666667 | 1 | 1 | 1.5 | 1 | 1.42156863 | 0.44921864 | 0.10895152 |
| 4 | 1.33333333 | 1.66666667 | 1 | 1.83333333 | 1.16666667 | 1 | 2 | 2.66666667 | 1 | 2.16666667 | 2.33333333 | 2.5 | 1.83333333 | 1 | 1.66666667 | 2.5 | 1 | 1.68627451 | 0.603469 | 0.14636273 |
| 5 | 2.16666667 | 1.66666667 | 1 | 1.5 | 1 | 1 | 2 | 3 | 1.33333333 | 2.66666667 | 3.5 | 3.33333333 | 3.16666667 | 1 | 1 | 1.33333333 | 1 | 1.8627451 | 0.92840083 | 0.22517028 |
| 10 | 2.33333333 | 2.03333333 | 1.16666667 | 1.63333333 | 1 | 1 | 2.23333333 | 4.26666667 | 1.6 | 3.06666667 | 2.46666667 | 4.1 | 3.1 | 1.03333333 | 1.66666667 | 1.5 | 1.13333333 | 2.07843137 | 1.03715297 | 0.25154654 |
| 15 | 2.43333333 | 2.6 | 1.23333333 | 1.06666667 | 1.06666667 | 1.06666667 | 2.53333333 | 4.43333333 | 1.43333333 | 3.16666667 | 2.36666667 | 4.36666667 | 3.53333333 | 1 | 1.5 | 2.33333333 | 1.1 | 2.19019608 | 1.15324209 | 0.27970229 |

Week 3

WT

| Time (min) | Animal ID | | | | | | | | | | | | |
|---|---|---|---|---|---|---|---|---|---|---|---|---|---|
| | **83** | **86** | **87** | **88** | **81** | **82** | **A** | **156** | **158** | **161** | Average | SD | SE |
| 0 | 4.86666667 | 3.2 | 2.43333333 | 2.7 | 1.92222222 | 2.53333333 | 2.15555556 | 2.21111111 | 1.02222222 | 1.35555556 | 2.44 | 1.06093108 | 0.33549587 |
| 1 | 1.66666667 | 1.66666667 | 1.33333333 | 1.33333333 | 1 | 1.5 | 1.16666667 | 1.5 | 1 | 1 | 1.31666667 | 0.26585525 | 0.08407081 |
| 2 | 2.16666667 | 1.66666667 | 1.5 | 1.33333333 | 1 | 2 | 1.66666667 | 1.66666667 | 1 | 1 | 1.5 | 0.41573971 | 0.13146844 |
| 3 | 1.66666667 | 2 | 1.16666667 | 1.16666667 | 1.16666667 | 1.83333333 | 1.16666667 | 1.83333333 | 1 | 1.16666667 | 1.41666667 | 0.37060178 | 0.11719457 |
| 4 | 2 | 2.33333333 | 1.66666667 | 1 | 1 | 2 | 1.5 | 1.5 | 1 | 1 | 1.5 | 0.49690399 | 0.15713484 |
| 5 | 2.66666667 | 3 | 1.83333333 | 1.66666667 | 1 | 2.33333333 | 2 | 2.16666667 | 1 | 1.33333333 | 1.9 | 0.67219926 | 0.21256807 |
| 10 | 2.4 | 2.2 | 2.06666667 | 2.1 | 1.96666667 | 2.9 | 1.86666667 | 1.73333333 | 1 | 1.3 | 1.95333333 | 0.53568924 | 0.16939981 |
| 15 | 3.46666667 | 1.9 | 2 | 2.36666667 | 1.96666667 | 2.46666667 | 1.43333333 | 2.2 | 1 | 1.03333333 | 1.98333333 | 0.73084595 | 0.23111378 |

Het

| Time (min) | Animal ID | | | | | | | | | | | | | | | | |
|---|---|---|---|---|---|---|---|---|---|---|---|---|---|---|---|---|---|
| | **B** | **D** | **E** | **H** | **155** | **157** | **190** | **196** | **199** | **201** | **205** | **207** | **208** | **210** | Average | SD | SE |
| 0 | 1.6 | 2.37777778 | 1.17777778 | 1.18888889 | 1.96666667 | 2.33333333 | 1.73333333 | 1.94444444 | 3.46666667 | 2 | 1.16666667 | 1.16666667 | 1.05555556 | 1.17777778 | 1.73968254 | 0.678369 | 0.18130174 |
| 1 | 1.2 | 2 | 1.2 | 1 | 1 | 1.33333333 | 1 | 1.16666667 | 2 | 6.5 | 1 | 1 | 1 | 1 | 1.6 | 1.45255448 | 0.38821152 |
| 2 | 1.5 | 2 | 1 | 1 | 1 | 2 | 1 | 1.33333333 | 2 | 1.33333333 | 1 | 1 | 1 | 1 | 1.29761905 | 0.41437097 | 0.1107453 |
| 3 | 1 | 1.16666667 | 1 | 1 | 1.5 | 2 | 1 | 1 | 2.66666667 | 1.83333333 | 1 | 1 | 1 | 1 | 1.29761905 | 0.51960937 | 0.13887144 |
| 4 | 1 | 1.66666667 | 1 | 1.33333333 | 1.5 | 2 | 1.66666667 | 1.33333333 | 2.33333333 | 1.83333333 | 1 | 1 | 1 | 1 | 1.4047619 | 0.4419957 | 0.11812832 |
| 5 | 1.83333333 | 1.83333333 | 1 | 1.5 | 1.5 | 2.5 | 2 | 2.66666667 | 2.83333333 | 2.16666667 | 1 | 1 | 1 | 1.33333333 | 1.72619048 | 0.64253485 | 0.17172466 |
| 10 | 1.5 | 2.66666667 | 1.13333333 | 1.5 | 2.06666667 | 3 | 2.23333333 | 2.56666667 | 3.36666667 | 2.23333333 | 1 | 1.1 | 1.03333333 | 1 | 1.88571429 | 0.81142634 | 0.21686281 |
| 15 | 2.03333333 | 2.46666667 | 1.2 | 1.66666667 | 2.23333333 | 2.53333333 | 2.33333333 | 2.56666667 | 4.23333333 | 2.06666667 | 1.2 | 1.06666667 | 1 | 1 | 1.97142857 | 0.8872176 | 0.23711888 |

Week 4

WT

| Time (min) | Animal ID | | | | | | | | | | | | |
|---|---|---|---|---|---|---|---|---|---|---|---|---|---|
| | **83** | **86** | **87** | **88** | **81** | **82** | **A** | **156** | **158** | **161** | Average | SD | SE |
| 0 | 3.9 | 2.56666667 | 2.14444444 | 2.55555556 | 1.55555556 | 2.75555556 | 2.05555556 | 2.38888889 | 1.58888889 | 1.64444444 | 2.31555556 | 0.70577084 | 0.22318434 |
| 1 | 2 | 1.5 | 1.33333333 | 1.16666667 | 1 | 1.83333333 | 1.5 | 1.33333333 | 1 | 1 | 1.36666667 | 0.34960295 | 0.11055416 |
| 2 | 1.83333333 | 1.83333333 | 1.33333333 | 1 | 1 | 1.66666667 | 1.83333333 | 1.33333333 | 1 | 1 | 1.38333333 | 0.37720545 | 0.11928284 |
| 3 | 2 | 1.83333333 | 1.16666667 | 1.16666667 | 1 | 1.83333333 | 1.5 | 1.83333333 | 1 | 1 | 1.43333333 | 0.40975753 | 0.12957671 |
| 4 | 2.16666667 | 1.33333333 | 1.5 | 1.5 | 1 | 1.5 | 1.83333333 | 2.16666667 | 1.16666667 | 1.33333333 | 1.55 | 0.39322974 | 0.12435016 |
| 5 | 2.33333333 | 1.5 | 1.66666667 | 1.33333333 | 1.16666667 | 1.83333333 | 1.83333333 | 2.5 | 1.5 | 1 | 1.66666667 | 0.47790696 | 0.15112745 |
| 10 | 2.8 | 1.96666667 | 1.86666667 | 2.03333333 | 1.36666667 | 2 | 1.66666667 | 2.8 | 1.26666667 | 1.46666667 | 1.92333333 | 0.53496279 | 0.16917009 |
| 15 | 3.1 | 2 | 1.83333333 | 1.63333333 | 1.53333333 | 2 | 1.56666667 | 2 | 1.13333333 | 1.03333333 | 1.78333333 | 0.57676193 | 0.18238814 |

Het

| Time (min) | Animal ID | | | | | | | | | | | | | | | |
|---|---|---|---|---|---|---|---|---|---|---|---|---|---|---|---|---|
| | **B** | **E** | **H** | **155** | **157** | **190** | **196** | **199** | **201** | **205** | **207** | **208** | **210** | Average | SD | SE |
| 0 | 1.84444444 | 1.18888889 | 1.26666667 | 1.83333333 | 2.24444444 | 2.95555556 | 1.95555556 | 2.51111111 | 2.55555556 | 1.01111111 | 1.52222222 | 1.63333333 | 1.32222222 | 1.83418803 | 0.5940629 | 0.1647634 |
| 1 | 1.5 | 1 | 1.33333333 | 1 | 1 | 1.16666667 | 1.33333333 | 1.33333333 | 3.16666667 | 1 | 1 | 1 | 1.33333333 | 1.32051282 | 0.58318069 | 0.16174522 |
| 2 | 1.33333333 | 1 | 1.5 | 1.33333333 | 1.5 | 1.16666667 | 1 | 1.83333333 | 1.16666667 | 1 | 1 | 1 | 1 | 1.21794872 | 0.26688026 | 0.07401927 |
| 3 | 1.16666667 | 1 | 1.16666667 | 1.66666667 | 1.5 | 1.16666667 | 1 | 2.5 | 1.33333333 | 1 | 1 | 1 | 1 | 1.26923077 | 0.4278416 | 0.11866191 |
| 4 | 1.16666667 | 1 | 1.66666667 | 2.16666667 | 1.66666667 | 1.33333333 | 1 | 1.33333333 | 1.33333333 | 1 | 1 | 1 | 1 | 1.28205128 | 0.36250553 | 0.10054094 |
| 5 | 1.16666667 | 1 | 1.16666667 | 2.16666667 | 1.33333333 | 1.83333333 | 1.66666667 | 2.66666667 | 1.5 | 1 | 1.16666667 | 1 | 1 | 1.43589744 | 0.52058755 | 0.14438501 |
| 10 | 1.36666667 | 1.06666667 | 2 | 2.06666667 | 1.56666667 | 2.2 | 2.13333333 | 3.46666667 | 2.2 | 1 | 1.03333333 | 1.13333333 | 1 | 1.71025641 | 0.72500123 | 0.20107916 |
| 15 | 1.63333333 | 1.2 | 1.5 | 2 | 2.03333333 | 2.26666667 | 2.4 | 4 | 2.83333333 | 1.46666667 | 1 | 1.4137931 | 1 | 1.90362511 | 0.83956579 | 0.23285365 |

### End expiratory pauase

Week 1

WT

| Time (min) | Animal ID | | | | | | | | | | | | |
|---|---|---|---|---|---|---|---|---|---|---|---|---|---|
| | **83** | **86** | **87** | **88** | **81** | **82** | **A** | **156** | **158** | **161** | Average | SD | SE |
| 0 | 9.57188889 | 14.2588889 | 17.151 | 11.9413333 | 16.3303333 | 15.5614444 | 13.8037778 | 14.0015556 | 19.1913333 | 16.7617778 | 14.8573333 | 2.76839951 | 0.87544479 |
| 1 | 11.2116667 | 8.06 | 10.4416667 | 8.575 | 11.602 | 8.89916667 | 18.56 | 8.298 | 14.802 | 11.874 | 11.23235 | 3.31311011 | 1.04769741 |
| 2 | 10.1416667 | 9.27 | 10.645 | 11.4733333 | 12.25 | 12.2383333 | 12.01 | 7.69 | 14.8333333 | 12.09 | 11.2641667 | 1.95324741 | 0.61767107 |
| 3 | 7.205 | 9.865 | 9.96833333 | 16.7483333 | 15.3316667 | 11.3175 | 10.7733333 | 9.87 | 12.785 | 12.5316667 | 11.6395833 | 2.81724652 | 0.89089157 |
| 4 | 8.395 | 9.64833333 | 7.16833333 | 10 | 12.805 | 7.75818182 | 11.2933333 | 6.30333333 | 9.71 | 10.54 | 9.36215152 | 1.97925499 | 0.62589539 |
| 5 | 8.99166667 | 8.79833333 | 11.4616667 | 14.8433333 | 17.9866667 | 9.605 | 13.8266667 | 10.1483333 | 11.525 | 10.5 | 11.7686667 | 2.94311142 | 0.93069355 |
| 10 | 16.0713333 | 12.2176667 | 18.336 | 18.5 | 15.76 | 15.243 | 15.608 | 13.987 | 11.8033333 | 14.0943333 | 15.1620667 | 2.23797442 | 0.70770965 |
| 15 | 17.07 | 12.4323333 | 15.988 | 18.485 | 16.4273333 | 17.9851852 | 23.011 | 13.2723333 | 22.4103333 | 16.6393333 | 17.3720852 | 3.3902035 | 1.07207648 |

Het

| Time (min) | Animal ID | | | | | | | | | | | | | | | | | | | | |
|---|---|---|---|---|---|---|---|---|---|---|---|---|---|---|---|---|---|---|---|---|---|
| | **B** | **D** | **E** | **F** | **G** | **H** | **155** | **157** | **159** | **160** | **162** | **190** | **196** | **199** | **201** | **205** | **207** | **208** | **210** | Average | SD | SE |
| 0 | 22.2876667 | 16.4431111 | 15.9296667 | 24.7781111 | 19.7625556 | 11.5418889 | 9.86055556 | 7.89844444 | 21.4474444 | 10.3068889 | 14.2316667 | 7.19966667 | 25.0264444 | 23.7248889 | 14.7087778 | 8.94311111 | 17.987 | 16.8431111 | 27.4802222 | 16.6526959 | 6.3148313 | 1.44872166 |
| 1 | 13.8283333 | 8.334 | 9.58833333 | 17.322 | 10.098 | 10.192 | 10.266 | 10.0783333 | 14.2933333 | 14.342 | 11.672 | 13.3816667 | 10.2316667 | 10.5333333 | 10.2616667 | 13.6566667 | 13.3783333 | 12.2683333 | 18.68 | 12.2318947 | 2.72729081 | 0.62568342 |
| 2 | 15.2883333 | 8.90666667 | 19.7683333 | 17.2683333 | 10.02 | 8.91 | 14.105 | 16.57 | 13.4666667 | 12.0066667 | 12.8933333 | 10.355 | 10.925 | 15.1516667 | 8.87333333 | 14.44 | 11.785 | 12.1766667 | 16.74 | 13.1394737 | 3.14128016 | 0.72065909 |
| 3 | 17.25 | 7.36 | 22.58 | 19.3816667 | 11.1166667 | 10.195 | 8.76666667 | 14.7516667 | 14.535 | 18.4416667 | 18.285 | 8.65166667 | 11.0816667 | 13.3016667 | 11.3916667 | 17.0366667 | 12.4316667 | 16.7783333 | 22.4683333 | 14.5160526 | 4.57539579 | 1.04966778 |
| 4 | 15.545 | 6.29 | 22.9733333 | 20.2566667 | 13.1533333 | 11.6266667 | 7.39833333 | 18.0633333 | 15.615 | 13.135 | 13.5733333 | 7.41333333 | 9.585 | 8.94666667 | 9.76666667 | 12.1416667 | 10.505 | 13.3983333 | 19.6583333 | 13.1076316 | 4.65909622 | 1.06886998 |
| 5 | 20.5333333 | 6.63333333 | 25.7783333 | 29.73 | 11.39 | 12.4633333 | 10.1633333 | 19.37 | 12.9483333 | 10.0383333 | 17.1033333 | 7.13833333 | 9.00833333 | 8.00666667 | 6.94666667 | 10.5083333 | 11.7233333 | 12.2116667 | 18.7283333 | 13.7064912 | 6.50315427 | 1.49192591 |
| 10 | 14.6886667 | 7.105 | 19.301 | 19.9886667 | 11.2833333 | 13.2263333 | 11.1093333 | 16.825 | 14.627 | 9.53233333 | 11.5383333 | 7.68333333 | 19.5746667 | 6.051 | 9.06866667 | 11.0616667 | 14.79 | 12.3293333 | 21.344 | 13.2172456 | 4.56759253 | 1.04787759 |
| 15 | 18.5486667 | 8.47133333 | 18.135 | 20.5343333 | 12.0423333 | 15.248 | 19.266 | 16.047 | 17.832 | 13.894 | 18.911 | 10.263 | 22.1913333 | 8.231 | 16.028 | 21.084 | 19.2866667 | 15.58 | 22.4473333 | 16.5284737 | 4.3214527 | 0.99140924 |

Week 2

WT

| Time (min) | Animal ID | | | | | | | | | | | | |
|---|---|---|---|---|---|---|---|---|---|---|---|---|---|
| | **83** | **86** | **87** | **88** | **81** | **82** | **A** | **156** | **158** | **161** | Average | SD | SE |
| 0 | 7.52644444 | 11.0421111 | 13.6523333 | 10.3076667 | 12.227 | 16.7321111 | 12.8531111 | 16.75 | 10.5135556 | 12.7102222 | 12.4314556 | 2.85017631 | 0.90130489 |
| 1 | 7.97333333 | 7.205 | 7.36666667 | 7.12666667 | 7.475 | 6.67 | 8.98666667 | 13.1433333 | 10.3766667 | 9.27333333 | 8.55966667 | 1.98343492 | 0.62721719 |
| 2 | 7.41333333 | 7.28 | 8.62333333 | 7.685 | 8.33 | 6.63333333 | 10.425 | 8.93666667 | 10.7416667 | 9.505 | 8.55733333 | 1.36465507 | 0.43154182 |
| 3 | 8.355 | 7.13333333 | 8.68666667 | 7.96833333 | 9.19166667 | 7.79 | 10.5366667 | 10.3583333 | 11.0916667 | 9.51 | 9.06216667 | 1.3066818 | 0.41320907 |
| 4 | 10.1666667 | 7.245 | 8.24 | 8.73 | 8.97166667 | 8.49 | 9.81833333 | 11.2333333 | 10.245 | 7.13333333 | 9.02733333 | 1.33487346 | 0.42212405 |
| 5 | 7.78833333 | 7.53666667 | 11.5933333 | 8.83833333 | 9.48666667 | 7.39666667 | 13.375 | 14.1783333 | 12.31 | 8.54333333 | 10.1046667 | 2.54047322 | 0.80336817 |
| 10 | 11.6033333 | 9.175 | 16.4016667 | 10.105 | 17.6856667 | 12.775 | 18.2236667 | 15.587 | 15.8553333 | 13.5603333 | 14.0972 | 3.1437318 | 0.99413529 |
| 15 | 10.6363333 | 12.525 | 17.4003333 | 13.7436667 | 18.798 | 13.9596667 | 16.373 | 15.9766667 | 11.7423333 | 17.0236667 | 14.8178667 | 2.69215285 | 0.85133348 |

Het

| Time (min) | Animal ID | | | | | | | | | | | | | | | | | | | |
|---|---|---|---|---|---|---|---|---|---|---|---|---|---|---|---|---|---|---|---|---|
| | **B** | **D** | **E** | **G** | **H** | **155** | **157** | **160** | **162** | **190** | **196** | **199** | **201** | **205** | **207** | **208** | **210** | Average | SD | SE |
| 0 | 11.4568889 | 16.8072222 | 16.1114444 | 13.8722222 | 11.9104444 | 17.0056667 | 10.9601111 | 11.6273333 | 13.8735556 | 16.9934444 | 20.4133333 | 25.9135556 | 20.9087778 | 9.84111111 | 17.3461111 | 8.30022222 | 15.3876667 | 15.2193595 | 4.50343205 | 1.09224271 |
| 1 | 10.135 | 9.49333333 | 8.658 | 9.70666667 | 7.30833333 | 10.6616667 | 14.7433333 | 11.3316667 | 10.8066667 | 7.985 | 9.52666667 | 12.2683333 | 11.5366667 | 10.7033333 | 8.96833333 | 12.1333333 | 9.69333333 | 10.3329216 | 1.78323739 | 0.4324986 |
| 2 | 10.5016667 | 8.72333333 | 9.24333333 | 10.2566667 | 9.68333333 | 11.3433333 | 14.93 | 9.85833333 | 9.33666667 | 7.08166667 | 6.59666667 | 10.8033333 | 7.21833333 | 9.49 | 8.02666667 | 10.9516667 | 8.17333333 | 9.5422549 | 1.9765285 | 0.47937858 |
| 3 | 13.1466667 | 9.19 | 9.38666667 | 9.67166667 | 8.955 | 12.0066667 | 15.4283333 | 10.49 | 9.39833333 | 5.69 | 5.59666667 | 10.3283333 | 6.18166667 | 8.03833333 | 7.365 | 13.385 | 9.10666667 | 9.60970588 | 2.73107327 | 0.66238256 |
| 4 | 8.24333333 | 9.07333333 | 9.46833333 | 9.92166667 | 9.91833333 | 13.0516667 | 18.49 | 9.185 | 11.7333333 | 6.035 | 7.37833333 | 7.19833333 | 6.16333333 | 8.88166667 | 7.47666667 | 8.51666667 | 7.53666667 | 9.31009804 | 2.98181971 | 0.72319751 |
| 5 | 7.62333333 | 8.08166667 | 10.675 | 9.51333333 | 12.7166667 | 12.5366667 | 14.69 | 10.31 | 15.1666667 | 6.65833333 | 7.565 | 8.99 | 8.16666667 | 9.43 | 6.975 | 7.83166667 | 10.1116667 | 9.82598039 | 2.59536419 | 0.62946828 |
| 10 | 17.761 | 9.72933333 | 12.0763333 | 10.4883333 | 14.312 | 13.3823333 | 11.994 | 16.7243333 | 17.69 | 11.7033333 | 17.025 | 11.7356667 | 18.3076667 | 11.4316667 | 14.1313333 | 9.24266667 | 11.1913333 | 13.4662549 | 3.00371524 | 0.72850795 |
| 15 | 13.633 | 12.578 | 14.7683333 | 12.2786667 | 14.9926667 | 12.1503333 | 12.8823333 | 29.6533333 | 30.2483333 | 12.899 | 10.6003333 | 12.998 | 13.485 | 12.9833333 | 20.4323333 | 11.45 | 15.227 | 15.4858824 | 5.84873618 | 1.41852688 |

Week 3

WT

| Time (min) | Animal ID | | | | | | | | | | | | |
|---|---|---|---|---|---|---|---|---|---|---|---|---|---|
| | **83** | **86** | **87** | **88** | **81** | **82** | **A** | **156** | **158** | **161** | Average | SD | SE |
| 0 | 7.47433333 | 9.76255556 | 15.4585556 | 13.5355556 | 20.7591111 | 17.6588889 | 11.1244444 | 16.7842222 | 17.8638889 | 13.8433333 | 14.4264889 | 4.09759241 | 1.29577249 |
| 1 | 6.75333333 | 6.38833333 | 5.98166667 | 6.825 | 6.96166667 | 6.46666667 | 6.75166667 | 8.28333333 | 10.9433333 | 10.2683333 | 7.56233333 | 1.71810066 | 0.54331113 |
| 2 | 6.035 | 8.08166667 | 6.36166667 | 7.375 | 7.72833333 | 9.935 | 8.12666667 | 10.8733333 | 11.7016667 | 10.7066667 | 8.6925 | 1.97949461 | 0.62597116 |
| 3 | 5.88166667 | 9.185 | 5.12666667 | 6.50666667 | 7.82333333 | 10.3733333 | 8.12 | 12.4216667 | 12.97 | 9.44833333 | 8.78566667 | 2.62959559 | 0.83155114 |
| 4 | 8.36166667 | 9.67 | 7.13833333 | 5.73666667 | 6.65166667 | 10.2283333 | 7.145 | 14.2166667 | 13.755 | 9.41166667 | 9.2315 | 2.88469519 | 0.91222071 |
| 5 | 7.005 | 8.545 | 7.475 | 8.03 | 6.96166667 | 9.015 | 8.04 | 16.7966667 | 12.5666667 | 11.7266667 | 9.61616667 | 3.15660546 | 0.99820629 |
| 10 | 8.24133333 | 15.243 | 16.4553333 | 17.347 | 10.7636667 | 12.8403333 | 16.3273333 | 16.962 | 15.4003333 | 14.7443333 | 14.4324667 | 2.95506453 | 0.93447346 |
| 15 | 8.917 | 19.5406667 | 26.9926667 | 19.712 | 23.142 | 16.458 | 21.348 | 13.7206667 | 17.4143333 | 18.7796667 | 18.6025 | 4.99259551 | 1.57879732 |

Het

| Time (min) | Animal ID | | | | | | | | | | | | | | | | |
|---|---|---|---|---|---|---|---|---|---|---|---|---|---|---|---|---|---|
| | **B** | **D** | **E** | **H** | **155** | **157** | **190** | **196** | **199** | **201** | **205** | **207** | **208** | **210** | Average | SD | SE |
| 0 | 16.0368889 | 18.7427778 | 16.6051111 | 18.5902222 | 19.1223333 | 18.5621111 | 15.9795556 | 18.699 | 23.8407778 | 20.1587778 | 14.247 | 15.0491111 | 15.8218889 | 15.5115556 | 17.6405079 | 2.53500114 | 0.67750755 |
| 1 | 7.792 | 8.475 | 9.602 | 8.27666667 | 10.845 | 9.91166667 | 9.23833333 | 8.15833333 | 9.525 | 11.22 | 10.8333333 | 9.65833333 | 10.9583333 | 8.43333333 | 9.49480952 | 1.15518496 | 0.30873617 |
| 2 | 6.995 | 7.85833333 | 10.8266667 | 8.34 | 13.6133333 | 14.7433333 | 7.2 | 6.215 | 10.5983333 | 12.5333333 | 11.7633333 | 8.235 | 9.57833333 | 7.94166667 | 9.74583333 | 2.64993225 | 0.70822418 |
| 3 | 6.39666667 | 5.8 | 11.2466667 | 8.68833333 | 17.485 | 16.2583333 | 5.985 | 5.69 | 10.0133333 | 6.75 | 10.9116667 | 7.14166667 | 11.0716667 | 7.08833333 | 9.32333333 | 3.78831007 | 1.01246845 |
| 4 | 6.515 | 6.24 | 9.97666667 | 8.40333333 | 14.47 | 7.29833333 | 6.25833333 | 5.95666667 | 8.19666667 | 6.47833333 | 10.5133333 | 8.36166667 | 10.4566667 | 8.15833333 | 8.37738095 | 2.35016873 | 0.62810901 |
| 5 | 11.1316667 | 6.395 | 10.3133333 | 10.7183333 | 15.255 | 7.025 | 8.50666667 | 9.22833333 | 9.785 | 7.335 | 10.865 | 7.555 | 8.32166667 | 13.1383333 | 9.68380952 | 2.47017607 | 0.66018233 |
| 10 | 9.947 | 12.708 | 17.18 | 19.6416667 | 11.5626667 | 11.914 | 13.606 | 14.8346667 | 9.907 | 12.6926667 | 9.83533333 | 12.938 | 8.73133333 | 16.5886667 | 13.0062143 | 3.14701733 | 0.84107576 |
| 15 | 13.6543333 | 23.9723333 | 24.9756667 | 20.0043333 | 18.747 | 11.6263333 | 18.7866667 | 16.0046667 | 17.9233333 | 10.498 | 11.1993333 | 15.0976667 | 16.2836667 | 20.4633333 | 17.0883333 | 4.48372347 | 1.1983255 |

Week 4

WT

| Time (min) | Animal ID | | | | | | | | | | | | |
|---|---|---|---|---|---|---|---|---|---|---|---|---|---|
| | **83** | **86** | **87** | **88** | **81** | **82** | **A** | **156** | **158** | **161** | Average | SD | SE |
| 0 | 7.95911111 | 19.5455556 | 15.2096667 | 12.4216667 | 22.4704444 | 13.9331111 | 14.893 | 14.1685556 | 19.7656667 | 14.3488889 | 15.4715667 | 4.15649784 | 1.31440003 |
| 1 | 7.35 | 6.68833333 | 7.38333333 | 6.84833333 | 8.29 | 6.52166667 | 6.05666667 | 6.125 | 9.51833333 | 8.41666667 | 7.31983333 | 1.11872956 | 0.35377335 |
| 2 | 7.86166667 | 9.665 | 7.475 | 8.625 | 9.57666667 | 9.33166667 | 7.005 | 10.025 | 13.9766667 | 9.945 | 9.34866667 | 1.94713541 | 0.61573828 |
| 3 | 7.485 | 9.92666667 | 6.93333333 | 7.45 | 10.4683333 | 9.76666667 | 5.82333333 | 8.69166667 | 12.845 | 8.91 | 8.83 | 2.03471902 | 0.64343465 |
| 4 | 7.27 | 9.355 | 7.725 | 8.53 | 11.1833333 | 10.3533333 | 5.91666667 | 8.09833333 | 11.995 | 7.82 | 8.82466667 | 1.88381587 | 0.59571488 |
| 5 | 8.29666667 | 8.47333333 | 8.94166667 | 9.69166667 | 10.4266667 | 10.9 | 7.03166667 | 9.43166667 | 12.955 | 11.1533333 | 9.73016667 | 1.69664201 | 0.53652531 |
| 10 | 13.9456667 | 14.906 | 14.5116667 | 13.8026667 | 14.5 | 14.4393333 | 15.3263333 | 12.1546667 | 16.5246667 | 12.992 | 14.3103 | 1.2091827 | 0.38237714 |
| 15 | 18.176 | 20.9643333 | 18.574 | 20.138 | 19.6113333 | 18.4996667 | 19.3586667 | 22.417 | 18.231 | 21.204 | 19.7174 | 1.44318489 | 0.45637513 |

Het

| Time (min) | Animal ID | | | | | | | | | | | | | | | |
|---|---|---|---|---|---|---|---|---|---|---|---|---|---|---|---|---|
| | **B** | **E** | **H** | **155** | **157** | **190** | **196** | **199** | **201** | **205** | **207** | **208** | **210** | Average | SD | SE |
| 0 | 17.4577778 | 15.07 | 19.7595556 | 17.0881111 | 22.8403333 | 14.476 | 20.2795556 | 14.5634444 | 13.9576667 | 15.1532222 | 11.1728889 | 13.3544444 | 13.8441111 | 16.0782393 | 3.26031295 | 0.90424812 |
| 1 | 8.885 | 9.65166667 | 7.775 | 9.46 | 11.6366667 | 9.38166667 | 8.01166667 | 9.985 | 15.1266667 | 10.2566667 | 10.315 | 11.5783333 | 9.78333333 | 10.1420513 | 1.87849824 | 0.52100167 |
| 2 | 7.81166667 | 11.2 | 8.375 | 9.73166667 | 10.3 | 8.84333333 | 7.265 | 12.035 | 10.9416667 | 9.6 | 7.86333333 | 10.4816667 | 9.10666667 | 9.50423077 | 1.45595482 | 0.40380921 |
| 3 | 6.53 | 12.7766667 | 8.57 | 7.40333333 | 8.38166667 | 5.83666667 | 6.08833333 | 10.825 | 10.5316667 | 7.14833333 | 7.31333333 | 9.90833333 | 9.24333333 | 8.50435897 | 2.07376498 | 0.57515892 |
| 4 | 6.87 | 11.5166667 | 9.18833333 | 8.58333333 | 8.93 | 7.135 | 6.06833333 | 6.05333333 | 6.695 | 7.52 | 7.05666667 | 9.68333333 | 7.04666667 | 7.87282051 | 1.60430979 | 0.44495548 |
| 5 | 7.53333333 | 15.0483333 | 11.97 | 13.8466667 | 7.2 | 9.92833333 | 7.40333333 | 6.45 | 6.635 | 8.46833333 | 7.65333333 | 7.51166667 | 9.92833333 | 9.19820513 | 2.80555711 | 0.77812154 |
| 10 | 10.6763333 | 16.125 | 17.8386667 | 18.9963333 | 13.834 | 11.5813333 | 9.64333333 | 9.652 | 8.746 | 11.1176667 | 15.1296667 | 11.489 | 17.0016667 | 13.2177692 | 3.45896629 | 0.95934464 |
| 15 | 9.63566667 | 14.7323333 | 22.924 | 17.4316667 | 23.3876667 | 20.2456667 | 9.09266667 | 21.9706667 | 9.98233333 | 9.575 | 17.0923333 | 14.967931 | 19.238 | 16.1750716 | 5.30647458 | 1.47175125 |

### Enhanced pause

#### Week 1

WT

| Time (min) | 83 | 86 | 87 | 88 | 81 | 82 | A | 156 | 158 | 161 | Average | SD | SE |
|---|---|---|---|---|---|---|---|---|---|---|---|---|---|
| 0 | 0.26233333 | 0.328 | 0.42188889 | 0.30266667 | 0.38233333 | 0.41188889 | 0.31611111 | 0.34944444 | 0.35833333 | 0.34633333 | 0.34793333 | 0.04903799 | 0.01550718 |
| 1 | 0.43666667 | 0.32166667 | 0.36 | 0.30833333 | 0.382 | 0.37416667 | 0.49 | 0.36 | 0.376 | 0.44 | 0.38488333 | 0.05584259 | 0.01765898 |
| 2 | 0.35833333 | 0.28833333 | 0.33 | 0.34333333 | 0.37666667 | 0.4325 | 0.795 | 0.34166667 | 0.34666667 | 0.44333333 | 0.40558333 | 0.14443592 | 0.04567465 |
| 3 | 0.25833333 | 0.27833333 | 0.305 | 0.50833333 | 0.44833333 | 0.38166667 | 0.78333333 | 0.34666667 | 0.33833333 | 0.50166667 | 0.415 | 0.15634127 | 0.04943945 |
| 4 | 0.27166667 | 0.28666667 | 0.26 | 0.31 | 0.295 | 0.34818182 | 0.69 | 0.335 | 0.28166667 | 0.41 | 0.34881818 | 0.1277828 | 0.04040847 |
| 5 | 0.29333333 | 0.28333333 | 0.28 | 0.44 | 0.41333333 | 0.27666667 | 0.73166667 | 0.395 | 0.29666667 | 0.31833333 | 0.37283333 | 0.13998909 | 0.04426844 |
| 10 | 0.391 | 0.26 | 0.39866667 | 0.46666667 | 0.33666667 | 0.30766667 | 0.63166667 | 0.53033333 | 0.30233333 | 0.37766667 | 0.40026667 | 0.11419128 | 0.03611045 |
| 15 | 0.46466667 | 0.351 | 0.42933333 | 0.45933333 | 0.357 | 0.39074074 | 0.539 | 0.49233333 | 0.43066667 | 0.40033333 | 0.43144074 | 0.05944203 | 0.01879722 |

Het

| Time (min) | B | D | E | F | G | H | 155 | 157 | 159 | 160 | 162 | 190 | 196 | 199 | 201 | 205 | 207 | 208 | 210 | Average | SD | SE |
|---|---|---|---|---|---|---|---|---|---|---|---|---|---|---|---|---|---|---|---|---|---|---|
| 0 | 0.538 | 0.45222222 | 0.382 | 0.94088889 | 0.31488889 | 0.25488889 | 0.27855556 | 0.25522222 | 0.31077778 | 0.28111111 | 0.33666667 | 0.265 | 0.52933333 | 0.457 | 0.31688889 | 0.30788889 | 0.33944444 | 0.496 | 0.53411111 | 0.39952047 | 0.16486192 | 0.03782192 |
| 1 | 0.69166667 | 0.362 | 0.46666667 | 1.286 | 0.53 | 0.372 | 0.492 | 0.49166667 | 0.74 | 0.778 | 0.596 | 0.62833333 | 0.5 | 0.325 | 0.36833333 | 0.65 | 0.97833333 | 1.02333333 | 0.64333333 | 0.62750877 | 0.25137941 | 0.05767039 |
| 2 | 0.65333333 | 0.35833333 | 0.73666667 | 0.74166667 | 0.54666667 | 0.35333333 | 0.41 | 0.465 | 0.66166667 | 0.58 | 0.45166667 | 0.53166667 | 0.45833333 | 0.30666667 | 0.33833333 | 0.465 | 0.58166667 | 0.67 | 0.68833333 | 0.52622807 | 0.13970741 | 0.03205108 |
| 3 | 0.735 | 0.285 | 0.73666667 | 1.06333333 | 0.66333333 | 0.32833333 | 0.28666667 | 0.41166667 | 0.49 | 0.78833333 | 0.38166667 | 0.42333333 | 0.33333333 | 0.25833333 | 0.38833333 | 0.61666667 | 0.55 | 0.69666667 | 0.765 | 0.53692982 | 0.22175246 | 0.0508735 |
| 4 | 0.67 | 0.335 | 0.64 | 1.225 | 0.675 | 0.395 | 0.26 | 0.365 | 0.41666667 | 0.395 | 0.27166667 | 0.33666667 | 0.27666667 | 0.30166667 | 0.33 | 0.355 | 0.47333333 | 0.60166667 | 0.69833333 | 0.47482456 | 0.2348863 | 0.05388661 |
| 5 | 0.93833333 | 0.31333333 | 0.78666667 | 1.03333333 | 0.50833333 | 0.46666667 | 0.26 | 0.42166667 | 0.34333333 | 0.34833333 | 0.39333333 | 0.32166667 | 0.28833333 | 0.30166667 | 0.32833333 | 0.33666667 | 0.475 | 0.54333333 | 0.69166667 | 0.47894737 | 0.22612081 | 0.05187567 |
| 10 | 0.47 | 0.28266667 | 0.6 | 1.17166667 | 0.356 | 0.397 | 0.265 | 0.41533333 | 0.35433333 | 0.29766667 | 0.25866667 | 0.29833333 | 0.35233333 | 0.24633333 | 0.301 | 0.274 | 0.38866667 | 0.464 | 0.55833333 | 0.40796491 | 0.21043098 | 0.04827618 |
| 15 | 0.43433333 | 0.27666667 | 0.50566667 | 1.016 | 0.37333333 | 0.327 | 0.42 | 0.37 | 0.29566667 | 0.33 | 0.23233333 | 0.322 | 0.39633333 | 0.263 | 0.37733333 | 0.46166667 | 0.38366667 | 0.39333333 | 0.47233333 | 0.40266667 | 0.16546907 | 0.03796121 |

#### Week 2

WT

| Time (min) | 83 | 86 | 87 | 88 | 81 | 82 | A | 156 | 158 | 161 | Average | SD | SE |
|---|---|---|---|---|---|---|---|---|---|---|---|---|---|
| 0 | 0.24488889 | 0.33022222 | 0.43644444 | 0.28966667 | 0.28911111 | 0.37833333 | 0.28433333 | 0.30755556 | 0.30044444 | 0.29755556 | 0.31585556 | 0.05449146 | 0.01723171 |
| 1 | 0.36 | 0.32666667 | 0.34166667 | 0.31666667 | 0.35833333 | 0.28833333 | 0.38333333 | 0.77666667 | 0.38666667 | 0.375 | 0.39133333 | 0.13893955 | 0.04393654 |
| 2 | 0.275 | 0.30333333 | 0.335 | 0.285 | 0.35333333 | 0.28666667 | 0.43 | 0.41166667 | 0.4 | 0.36666667 | 0.34466667 | 0.05673634 | 0.01794161 |
| 3 | 0.295 | 0.265 | 0.325 | 0.26 | 0.33166667 | 0.23833333 | 0.48666667 | 0.41 | 0.27833333 | 0.33166667 | 0.32216667 | 0.07585338 | 0.02398694 |
| 4 | 0.38333333 | 0.27833333 | 0.30666667 | 0.24333333 | 0.31166667 | 0.27166667 | 0.485 | 0.345 | 0.30833333 | 0.36333333 | 0.32966667 | 0.06923212 | 0.02189312 |
| 5 | 0.28666667 | 0.26833333 | 0.31166667 | 0.27 | 0.30833333 | 0.24 | 0.54166667 | 0.34833333 | 0.27833333 | 0.33 | 0.31833333 | 0.08474362 | 0.02679829 |
| 10 | 0.291 | 0.275 | 0.329 | 0.299 | 0.318 | 0.20966667 | 0.77066667 | 0.40333333 | 0.25366667 | 0.35 | 0.34993333 | 0.15703871 | 0.04966 |
| 15 | 0.263 | 0.23833333 | 0.386 | 0.35 | 0.34633333 | 0.234 | 0.5 | 0.342 | 0.25666667 | 0.34866667 | 0.3265 | 0.08202864 | 0.02593973 |

Het

| Time (min) | B | D | E | G | H | 155 | 157 | 160 | 162 | 190 | 196 | 199 | 201 | 205 | 207 | 208 | 210 | Average | SD | SE |
|---|---|---|---|---|---|---|---|---|---|---|---|---|---|---|---|---|---|---|---|---|
| 0 | 0.34288889 | 0.45288889 | 0.33966667 | 0.513 | 0.25888889 | 0.40433333 | 0.27111111 | 0.29544444 | 0.29466667 | 0.44433333 | 0.46077778 | 0.48488889 | 0.41433333 | 0.28066667 | 0.35922222 | 0.30066667 | 0.31244444 | 0.36648366 | 0.08232896 | 0.0199677 |
| 1 | 0.42666667 | 0.40833333 | 0.488 | 0.35 | 0.35333333 | 0.42333333 | 0.625 | 0.39333333 | 0.42833333 | 0.35666667 | 0.33 | 0.345 | 0.36333333 | 0.525 | 0.38 | 0.64 | 0.58833333 | 0.4367451 | 0.10078046 | 0.02444285 |
| 2 | 0.42166667 | 0.34166667 | 0.35333333 | 0.39 | 0.33 | 0.31833333 | 0.515 | 0.36 | 0.465 | 0.26833333 | 0.27666667 | 0.28 | 0.27666667 | 0.41833333 | 0.38666667 | 0.565 | 0.47 | 0.37862745 | 0.08796352 | 0.02133429 |
| 3 | 0.49 | 0.38833333 | 0.345 | 0.26833333 | 0.41333333 | 0.315 | 0.33333333 | 0.415 | 0.38833333 | 0.29333333 | 0.255 | 0.295 | 0.305 | 0.29333333 | 0.455 | 0.61833333 | 0.43166667 | 0.37078431 | 0.09414845 | 0.02283435 |
| 4 | 0.31833333 | 0.35833333 | 0.40166667 | 0.31833333 | 0.34666667 | 0.34166667 | 0.32333333 | 0.27333333 | 0.33833333 | 0.26 | 0.26 | 0.28833333 | 0.28666667 | 0.275 | 0.35166667 | 0.30666667 | 0.38166667 | 0.31941176 | 0.04195708 | 0.01017609 |
| 5 | 0.32333333 | 0.345 | 0.32166667 | 0.30833333 | 0.335 | 0.34 | 0.245 | 0.29833333 | 0.465 | 0.24833333 | 0.22833333 | 0.27833333 | 0.25833333 | 0.26333333 | 0.27166667 | 0.34833333 | 0.305 | 0.30490196 | 0.05606751 | 0.01359837 |
| 10 | 0.344 | 0.35633333 | 0.34733333 | 0.31033333 | 0.35233333 | 0.315 | 0.233 | 0.329 | 0.41333333 | 0.301 | 0.28866667 | 0.27766667 | 0.28233333 | 0.33733333 | 0.288 | 0.31933333 | 0.36433333 | 0.32113725 | 0.04182571 | 0.01014422 |
| 15 | 0.329 | 0.369 | 0.28366667 | 0.42333333 | 0.323 | 0.28366667 | 0.22 | 0.38033333 | 0.26133333 | 0.27633333 | 0.41733333 | 0.27633333 | 0.27766667 | 0.267 | 0.32 | 0.33666667 | 0.41733333 | 0.32129412 | 0.0614931 | 0.01491427 |

#### Week 3

WT

| Time (min) | 83 | 86 | 87 | 88 | 81 | 82 | A | 156 | 158 | 161 | Average | SD | SE |
|---|---|---|---|---|---|---|---|---|---|---|---|---|---|
| 0 | 0.23922222 | 0.29333333 | 0.29344444 | 0.35322222 | 0.41933333 | 0.35066667 | 0.28755556 | 0.30933333 | 0.31577778 | 0.30455556 | 0.31664444 | 0.04849148 | 0.01533435 |
| 1 | 0.34 | 0.34166667 | 0.36333333 | 0.41333333 | 0.39666667 | 0.375 | 0.35166667 | 0.36166667 | 0.34666667 | 0.46 | 0.375 | 0.03824063 | 0.01209275 |
| 2 | 0.33666667 | 0.37333333 | 0.33333333 | 0.36 | 0.33833333 | 0.27333333 | 0.365 | 0.35833333 | 0.245 | 0.43 | 0.34133333 | 0.05172816 | 0.01635788 |
| 3 | 0.35166667 | 0.29833333 | 0.31833333 | 0.345 | 0.35833333 | 0.28166667 | 0.35166667 | 0.32333333 | 0.27 | 0.365 | 0.32633333 | 0.03362649 | 0.01063363 |
| 4 | 0.33833333 | 0.33833333 | 0.35166667 | 0.33333333 | 0.38166667 | 0.32 | 0.31833333 | 0.34166667 | 0.245 | 0.36833333 | 0.33366667 | 0.03684971 | 0.0116529 |
| 5 | 0.27333333 | 0.25 | 0.30833333 | 0.325 | 0.39666667 | 0.27333333 | 0.335 | 0.32 | 0.36833333 | 0.36 | 0.321 | 0.04648244 | 0.01469904 |
| 10 | 0.29366667 | 0.265 | 0.38133333 | 0.304 | 0.32766667 | 0.24633333 | 0.404 | 0.35566667 | 0.27566667 | 0.32966667 | 0.3183 | 0.0511858 | 0.01618637 |
| 15 | 0.25566667 | 0.318 | 0.50666667 | 0.416 | 0.34566667 | 0.268 | 0.43366667 | 0.28 | 0.27333333 | 0.27533333 | 0.33723333 | 0.08646716 | 0.02734332 |

Het

| Time (min) | B | D | E | H | 155 | 157 | 190 | 196 | 199 | 201 | 205 | 207 | 208 | 210 | Average | SD | SE |
|---|---|---|---|---|---|---|---|---|---|---|---|---|---|---|---|---|---|
| 0 | 0.37122222 | 0.34122222 | 0.42711111 | 0.29877778 | 0.41988889 | 0.35722222 | 0.44677778 | 0.45077778 | 0.38955556 | 0.41744444 | 0.40211111 | 0.36466667 | 0.43 | 0.55411111 | 0.40506349 | 0.0608994 | 0.01627605 |
| 1 | 0.388 | 0.36833333 | 0.592 | 0.44166667 | 0.39333333 | 0.40333333 | 0.365 | 0.34833333 | 0.32166667 | 0.39833333 | 0.39 | 0.40666667 | 0.47 | 0.38166667 | 0.40488095 | 0.06497503 | 0.01736531 |
| 2 | 0.36833333 | 0.33666667 | 0.77833333 | 0.34333333 | 0.30666667 | 0.33833333 | 0.34333333 | 0.31333333 | 0.25166667 | 0.4 | 0.40333333 | 0.40666667 | 0.4 | 0.32166667 | 0.37940476 | 0.12305514 | 0.03288787 |
| 3 | 0.37666667 | 0.34333333 | 0.56333333 | 0.41166667 | 0.325 | 0.33166667 | 0.285 | 0.295 | 0.255 | 0.285 | 0.25166667 | 0.43 | 0.21166667 | 0.31666667 | 0.33440476 | 0.0896327 | 0.02395535 |
| 4 | 0.40833333 | 0.335 | 0.65833333 | 0.47666667 | 0.25833333 | 0.29333333 | 0.27 | 0.29 | 0.28166667 | 0.275 | 0.21833333 | 0.37666667 | 0.19666667 | 0.28 | 0.32988095 | 0.12007737 | 0.03209203 |
| 5 | 0.40333333 | 0.295 | 0.615 | 0.49666667 | 0.29666667 | 0.26 | 0.25 | 0.29 | 0.28833333 | 0.245 | 0.19 | 0.27666667 | 0.25833333 | 0.35666667 | 0.32297619 | 0.11306185 | 0.03021705 |
| 10 | 0.301 | 0.31066667 | 0.415 | 0.62833333 | 0.26233333 | 0.25166667 | 0.29233333 | 0.28433333 | 0.26066667 | 0.47866667 | 0.24633333 | 0.26266667 | 0.26966667 | 0.33933333 | 0.32878571 | 0.10877324 | 0.02907087 |
| 15 | 0.28666667 | 0.32033333 | 0.381 | 0.426 | 0.30666667 | 0.279 | 0.337 | 0.296 | 0.24 | 0.276 | 0.33966667 | 0.254 | 0.24666667 | 0.41166667 | 0.31433333 | 0.0589666 | 0.01575949 |

#### Week 4

WT

| Time (min) | 83 | 86 | 87 | 88 | 81 | 82 | A | 156 | 158 | 161 | Average | SD | SE |
|---|---|---|---|---|---|---|---|---|---|---|---|---|---|
| 0 | 0.26211111 | 0.32355556 | 0.35 | 0.37622222 | 0.455 | 0.31222222 | 0.30811111 | 0.28244444 | 0.28877778 | 0.29588889 | 0.32543333 | 0.05629577 | 0.01780229 |
| 1 | 0.32166667 | 0.39 | 0.38 | 0.41 | 0.47666667 | 0.31333333 | 0.32166667 | 0.33333333 | 0.3 | 0.33666667 | 0.35833333 | 0.05513172 | 0.01743418 |
| 2 | 0.31666667 | 0.37333333 | 0.355 | 0.39666667 | 0.39333333 | 0.31666667 | 0.30166667 | 0.405 | 0.25 | 0.36333333 | 0.34716667 | 0.04978379 | 0.01574302 |
| 3 | 0.29333333 | 0.41666667 | 0.41333333 | 0.41333333 | 0.37166667 | 0.31333333 | 0.29666667 | 0.30833333 | 0.33666667 | 0.375 | 0.35383333 | 0.05025276 | 0.01589132 |
| 4 | 0.31333333 | 0.45666667 | 0.41166667 | 0.45 | 0.33666667 | 0.31166667 | 0.28333333 | 0.27333333 | 0.27166667 | 0.34333333 | 0.34516667 | 0.07027042 | 0.02222146 |
| 5 | 0.315 | 0.395 | 0.35 | 0.36666667 | 0.37 | 0.32166667 | 0.255 | 0.28666667 | 0.28166667 | 0.34166667 | 0.32833333 | 0.04448609 | 0.01406774 |
| 10 | 0.29933333 | 0.382 | 0.387 | 0.38833333 | 0.44133333 | 0.32533333 | 0.36433333 | 0.3 | 0.31666667 | 0.3 | 0.35043333 | 0.04909679 | 0.01552577 |
| 15 | 0.3 | 0.38266667 | 0.35233333 | 0.39866667 | 0.36433333 | 0.35366667 | 0.392 | 0.32566667 | 0.296 | 0.30233333 | 0.34676667 | 0.03887239 | 0.01229253 |

Het

| Time (min) | B | E | H | 155 | 157 | 190 | 196 | 199 | 201 | 205 | 207 | 208 | 210 | Average | SD | SE |
|---|---|---|---|---|---|---|---|---|---|---|---|---|---|---|---|---|
| 0 | 0.345 | 0.377 | 0.28244444 | 0.392 | 0.41 | 0.385 | 0.42544444 | 0.32766667 | 0.32311111 | 0.36811111 | 0.32177778 | 0.367 | 0.33388889 | 0.35834188 | 0.04040302 | 0.01120578 |
| 1 | 0.38166667 | 0.65666667 | 0.395 | 0.33166667 | 0.49166667 | 0.365 | 0.33833333 | 0.30833333 | 0.85333333 | 0.45 | 0.39833333 | 0.535 | 0.44333333 | 0.4575641 | 0.15214609 | 0.04219773 |
| 2 | 0.38 | 0.915 | 0.395 | 0.31333333 | 0.305 | 0.29333333 | 0.34 | 0.27666667 | 0.51666667 | 0.34166667 | 0.34166667 | 0.34333333 | 0.30666667 | 0.38987179 | 0.16915977 | 0.04691648 |
| 3 | 0.445 | 1.075 | 0.425 | 0.27333333 | 0.30166667 | 0.29833333 | 0.3 | 0.215 | 0.39666667 | 0.365 | 0.43666667 | 0.27 | 0.285 | 0.39128205 | 0.21799609 | 0.06046124 |
| 4 | 0.38166667 | 0.80166667 | 0.44333333 | 0.25833333 | 0.28666667 | 0.26166667 | 0.30166667 | 0.31166667 | 0.32166667 | 0.40166667 | 0.41833333 | 0.28333333 | 0.31333333 | 0.36807692 | 0.14359125 | 0.03982505 |
| 5 | 0.38666667 | 0.51833333 | 0.48166667 | 0.205 | 0.27666667 | 0.24333333 | 0.285 | 0.26166667 | 0.29666667 | 0.325 | 0.36166667 | 0.35333333 | 0.28 | 0.32884615 | 0.09106973 | 0.0252582 |
| 10 | 0.36966667 | 0.41766667 | 0.35166667 | 0.30366667 | 0.27 | 0.27633333 | 0.26033333 | 0.266 | 0.29766667 | 0.281 | 0.23433333 | 0.292 | 0.24433333 | 0.29728205 | 0.05269142 | 0.01461397 |
| 15 | 0.37266667 | 0.41033333 | 0.27466667 | 0.28966667 | 0.27533333 | 0.283 | 0.25766667 | 0.28333333 | 0.27733333 | 0.475 | 0.266 | 0.25137931 | 0.289 | 0.3081061 | 0.06775895 | 0.01879295 |

### Expiratory flow 50% through expiration

Week 1

WT

| Time (min) | Animal ID | | | | | | | | | | | |
|---|---|---|---|---|---|---|---|---|---|---|---|---|
| | 83 | 86 | 87 | 88 | 81 | 82 | A | 156 | 158 | 161 | Average | SD | SE |
| 0 | 1.62222222 | 1.52 | 1.40333333 | 1.72555556 | 1.27888889 | 1.31222222 | 1.09555556 | 1.64666667 | 0.70666667 | 0.75888889 | 1.307 | 0.35760446 | 0.11308446 |
| 1 | 1.86666667 | 2.48333333 | 1.95 | 3.18333333 | 1.94 | 2.075 | 0.48333333 | 2.04 | 0.94 | 1.28 | 1.82416667 | 0.7672531 | 0.24262673 |
| 2 | 1.55 | 2.23333333 | 1.75 | 1.6 | 1.6 | 1.73333333 | 0.66666667 | 1.76666667 | 0.7 | 1.01666667 | 1.46166667 | 0.50571121 | 0.15991993 |
| 3 | 1.55 | 2.03333333 | 1.83333333 | 1.36666667 | 1.63333333 | 1.83333333 | 0.9 | 1.78333333 | 0.93333333 | 1.2 | 1.50666667 | 0.39379446 | 0.12452874 |
| 4 | 1.6 | 1.86666667 | 2.13333333 | 1.76666667 | 1.33333333 | 1.5 | 0.71666667 | 2.48333333 | 1.13333333 | 1.21666667 | 1.575 | 0.5146166 | 0.16273606 |
| 5 | 1.35 | 1.96666667 | 1.73333333 | 1.65 | 1.38333333 | 1.7 | 0.63333333 | 1.83333333 | 1.1 | 1.33333333 | 1.46833333 | 0.39592445 | 0.12520231 |
| 10 | 1.11666667 | 1.54 | 1.30666667 | 1.42 | 1.36666667 | 1.39333333 | 0.56333333 | 1.67666667 | 1.15666667 | 1.03666667 | 1.25766667 | 0.31186437 | 0.09862017 |
| 15 | 1.22333333 | 1.65 | 1.43 | 1.28 | 1.48 | 1.27037037 | 0.43333333 | 1.76 | 0.69666667 | 0.83666667 | 1.20603704 | 0.42584104 | 0.13466276 |

Het

| Time (min) | Animal ID | | | | | | | | | | | | | | | | | | | | |
|---|---|---|---|---|---|---|---|---|---|---|---|---|---|---|---|---|---|---|---|---|---|
| | B | D | E | F | G | H | 155 | 157 | 159 | 160 | 162 | 190 | 196 | 199 | 201 | 205 | 207 | 208 | 210 | Average | SD | SE |
| 0 | 0.95666667 | 1.02888889 | 1.13777778 | 0.61444444 | 0.71777778 | 1.18888889 | 1.27333333 | 1.41777778 | 0.63555556 | 1.34 | 0.89666667 | 1.59222222 | 0.92222222 | 1.02222222 | 1.27444444 | 1.00333333 | 0.79111111 | 0.58444444 | 0.50777778 | 0.99502924 | 0.30593744 | 0.07018686 |
| 1 | 1.78333333 | 2.2 | 2.23333333 | 0.94 | 1.18 | 1.3 | 1.86 | 1.75 | 1.28333333 | 1.5 | 1.52 | 1.85 | 2.13333333 | 2.05 | 2 | 1.23333333 | 1.33333333 | 1.48333333 | 1.11666667 | 1.61842105 | 0.39855816 | 0.09143551 |
| 2 | 1.45 | 1.68333333 | 1.46666667 | 0.86666667 | 0.96666667 | 1.15 | 1.36666667 | 1.15 | 1.35 | 1.3 | 1.1 | 1.66666667 | 1.81666667 | 1.78333333 | 1.98333333 | 1 | 1.06666667 | 0.76666667 | 0.93333333 | 1.30877193 | 0.35370039 | 0.08114443 |
| 3 | 1.2 | 1.76666667 | 0.88333333 | 0.51666667 | 0.86666667 | 1.23333333 | 1.38333333 | 1.05 | 0.98333333 | 1.21666667 | 0.86666667 | 1.86666667 | 1.61666667 | 1.46666667 | 1.7 | 1.03333333 | 1.15 | 0.7 | 0.81666667 | 1.1745614 | 0.37779283 | 0.08667162 |
| 4 | 1.21666667 | 1.98333333 | 0.73333333 | 0.46666667 | 0.9 | 1.08333333 | 1.58333333 | 0.93333333 | 0.81666667 | 1.25 | 0.81666667 | 2.56666667 | 1.61666667 | 1.86666667 | 1.96666667 | 1.03333333 | 1.4 | 0.7 | 0.9 | 1.25438596 | 0.54836576 | 0.12580373 |
| 5 | 1.06666667 | 1.86666667 | 0.81666667 | 1.35 | 0.91666667 | 1.08333333 | 1.28333333 | 0.91666667 | 1.03333333 | 1.53333333 | 0.78333333 | 2.7 | 1.63333333 | 2.03333333 | 2.16666667 | 1.18333333 | 1.35 | 0.76666667 | 0.88333333 | 1.33508772 | 0.53520778 | 0.12278509 |
| 10 | 1.56 | 1.53 | 1.00666667 | 0.55666667 | 0.85666667 | 1.16 | 1.16666667 | 1.05666667 | 1.01 | 1.50666667 | 0.97333333 | 2.07333333 | 1.06333333 | 1.56666667 | 1.60333333 | 0.97333333 | 1.06333333 | 0.90666667 | 0.68333333 | 1.1745614 | 0.37280255 | 0.08552677 |
| 15 | 1.25333333 | 1.4 | 1.12333333 | 0.71 | 0.96 | 1.06 | 0.82666667 | 1.14666667 | 0.72666667 | 1.29666667 | 0.66666667 | 1.63666667 | 1.02666667 | 1.52333333 | 1.38333333 | 0.78 | 0.61666667 | 0.66666667 | 0.63 | 1.02280702 | 0.32612439 | 0.07481807 |

Week 2

WT

| Time (min) | Animal ID | | | | | | | | | | | |
|---|---|---|---|---|---|---|---|---|---|---|---|---|
| | 83 | 86 | 87 | 88 | 81 | 82 | A | 156 | 158 | 161 | Average | SD | SE |
| 0 | 2.09222222 | 2.36333333 | 1.70555556 | 2.50777778 | 1.97333333 | 1.32111111 | 1.81777778 | 1.33444444 | 1.35 | 1.45777778 | 1.79233333 | 0.43564419 | 0.13776279 |
| 1 | 2.61666667 | 3.63333333 | 3.2 | 4.01666667 | 3.01666667 | 3.63333333 | 2.48333333 | 2.23333333 | 1 | 1.7 | 2.75333333 | 0.94040968 | 0.29738365 |
| 2 | 1.96666667 | 3.65 | 3.05 | 4.11666667 | 2.96666667 | 3.31666667 | 1.95 | 2.38333333 | 0.98333333 | 1.55 | 2.59333333 | 0.99104634 | 0.31339637 |
| 3 | 2.11666667 | 3.18333333 | 2.98333333 | 3.55 | 2.71666667 | 2.81666667 | 1.95 | 1.98333333 | 1.15 | 1.71666667 | 2.41666667 | 0.74783639 | 0.23648663 |
| 4 | 2.28333333 | 2.98333333 | 3.08333333 | 2.98333333 | 2.86666667 | 2.68333333 | 2.18333333 | 1.71666667 | 1.25 | 2.3 | 2.43333333 | 0.60502015 | 0.19132417 |
| 5 | 2.36666667 | 2.93333333 | 2.61666667 | 2.68333333 | 2.3 | 2.36666667 | 1.71666667 | 1.35 | 1.21666667 | 1.95 | 2.15 | 0.57456371 | 0.181693 |
| 10 | 1.93 | 2.3 | 2.09666667 | 2.68 | 1.69666667 | 1.54333333 | 1.96333333 | 1.61333333 | 1.05 | 1.28 | 1.81533333 | 0.48280866 | 0.1526775 |
| 15 | 1.9 | 1.8 | 1.84333333 | 2.55666667 | 1.47666667 | 1.45 | 1.81333333 | 1.58 | 1.14666667 | 0.96 | 1.65266667 | 0.44414935 | 0.14045236 |

Het

| Time (min) | Animal ID | | | | | | | | | | | | | | | | | | | |
|---|---|---|---|---|---|---|---|---|---|---|---|---|---|---|---|---|---|---|---|---|
| | B | D | E | G | H | 155 | 157 | 160 | 162 | 190 | 196 | 199 | 201 | 205 | 207 | 208 | 210 | Average | SD | SE |
| 0 | 1.87 | 1.05444444 | 0.87 | 0.55777778 | 1.33666667 | 1.06555556 | 1.38 | 1.01555556 | 0.87333333 | 1.18111111 | 1.02111111 | 0.95333333 | 1.25888889 | 1.30333333 | 0.79555556 | 1.46 | 1.04 | 1.11980392 | 0.30167338 | 0.07316654 |
| 1 | 2.73333333 | 2.2 | 2.2 | 1.11666667 | 1.43333333 | 2.21666667 | 1.81666667 | 1.48333333 | 1.38333333 | 2.51666667 | 2.11666667 | 1.81666667 | 2.01666667 | 1.3 | 1.73333333 | 2.15 | 1.66666667 | 1.87647059 | 0.4469829 | 0.10840928 |
| 2 | 2.1 | 2.4 | 1.66666667 | 1.11666667 | 1.43333333 | 1.35 | 1.66666667 | 1.31666667 | 1.3 | 2.25 | 2.15 | 1.73333333 | 2.06666667 | 0.96666667 | 1.9 | 1.48333333 | 1.41666667 | 1.66568627 | 0.41971678 | 0.10179627 |
| 3 | 1.5 | 2.05 | 1.48333333 | 1.11666667 | 1.73333333 | 1.1 | 1.05 | 1.28333333 | 1.15 | 2.21666667 | 2.46666667 | 1.83333333 | 2.48333333 | 1.13333333 | 1.93333333 | 1.2 | 1.31666667 | 1.59117647 | 0.49160019 | 0.11923056 |
| 4 | 2.21666667 | 2.13333333 | 1.71666667 | 1.26666667 | 1.63333333 | 0.96666667 | 0.96666667 | 1.41666667 | 0.95 | 2.06666667 | 2.06666667 | 2.23333333 | 2.18333333 | 1.2 | 1.6 | 1.56666667 | 1.4 | 1.62254902 | 0.46221617 | 0.11210389 |
| 5 | 2.5 | 2.4 | 1.43333333 | 1.3 | 1.2 | 0.95 | 0.88333333 | 1.45 | 0.85 | 1.9 | 1.55 | 1.96666667 | 1.86666667 | 1.15 | 1.35 | 1.9 | 1.28333333 | 1.5254902 | 0.49420663 | 0.11986271 |
| 10 | 1.30333333 | 2.13333333 | 1.32 | 0.99333333 | 1.23333333 | 0.92666667 | 1.22666667 | 1.05333333 | 0.78 | 1.59666667 | 1.05 | 1.44 | 1.26 | 0.99333333 | 1.06666667 | 1.43333333 | 1.10666667 | 1.23039216 | 0.31248307 | 0.07578828 |
| 15 | 1.51333333 | 1.75 | 1.00333333 | 0.85333333 | 1.14 | 1.12666667 | 1.20666667 | 0.68333333 | 0.42333333 | 1.46 | 1.99333333 | 1.37333333 | 1.48333333 | 0.85333333 | 0.74333333 | 1.19666667 | 0.99 | 1.16431373 | 0.40289536 | 0.09771648 |

Week 3

WT

| Time (min) | Animal ID | | | | | | | | | | | |
|---|---|---|---|---|---|---|---|---|---|---|---|---|
| | 83 | 86 | 87 | 88 | 81 | 82 | A | 156 | 158 | 161 | Average | SD | SE |
| 0 | 2.21444444 | 2.23 | 1.78111111 | 1.88444444 | 1.39444444 | 1.34222222 | 1.95777778 | 1.80666667 | 0.81444444 | 1.31555556 | 1.67411111 | 0.4485349 | 0.14183919 |
| 1 | 3.2 | 3.71666667 | 4.2 | 4.03333333 | 3.8 | 3.01666667 | 3.4 | 2.98333333 | 1.25 | 1.81666667 | 3.14166667 | 0.95008934 | 0.30044463 |
| 2 | 2.96666667 | 3.18333333 | 3.03333333 | 3.25 | 3.1 | 1.85 | 2.35 | 1.85 | 0.75 | 1.35 | 2.36833333 | 0.88007611 | 0.2783045 |
| 3 | 3.21666667 | 2.56666667 | 3.16666667 | 3.55 | 3.3 | 1.75 | 2.78333333 | 1.63333333 | 0.75 | 1.88333333 | 2.46 | 0.91443162 | 0.28916867 |
| 4 | 2.53333333 | 2.6 | 2.86666667 | 3.38333333 | 3.88333333 | 1.86666667 | 2.65 | 1.65 | 0.75 | 1.98333333 | 2.41666667 | 0.89725232 | 0.2837361 |
| 5 | 2.08333333 | 2.08333333 | 2.16666667 | 2.65 | 3.8 | 2.01666667 | 2.41666667 | 1.48333333 | 1.05 | 1.55 | 2.13 | 0.75136912 | 0.23760378 |
| 10 | 2.42 | 1.78 | 2.06333333 | 1.48666667 | 2.11333333 | 1.73 | 1.74333333 | 1.76 | 1.14666667 | 1.34333333 | 1.75866667 | 0.37690487 | 0.11918779 |
| 15 | 2.15666667 | 1.56333333 | 1.46666667 | 1.64 | 1.37666667 | 1.43 | 1.28666667 | 1.82333333 | 1.01333333 | 0.89333333 | 1.465 | 0.36798836 | 0.11636814 |

Het

| Time (min) | Animal ID | | | | | | | | | | | | | | | | |
|---|---|---|---|---|---|---|---|---|---|---|---|---|---|---|---|---|---|
| | B | D | E | H | 155 | 157 | 190 | 196 | 199 | 201 | 205 | 207 | 208 | 210 | Average | SD | SE |
| 0 | 1.36333333 | 1.18222222 | 0.72111111 | 0.99111111 | 1.13111111 | 1.18333333 | 1.23 | 1.17666667 | 1.14666667 | 1.22555556 | 0.75888889 | 0.76777778 | 0.69111111 | 0.80222222 | 1.02650794 | 0.22990392 | 0.06144441 |
| 1 | 3.04 | 2.6 | 1.64 | 2.38333333 | 2.33333333 | 2.36666667 | 2.3 | 2.93333333 | 2.46666667 | 2.08333333 | 1.43333333 | 1.4 | 1.21666667 | 1.75 | 2.13904762 | 0.57083449 | 0.15256193 |
| 2 | 3.18333333 | 2.33333333 | 1.7 | 1.68333333 | 1.16666667 | 1.63333333 | 2.1 | 2.45 | 2.03333333 | 1.91666667 | 1.15 | 1.46666667 | 1.1 | 1.46666667 | 1.81309524 | 0.57935193 | 0.15483832 |
| 3 | 3.26666667 | 2.63333333 | 1.51666667 | 1.98333333 | 1.06666667 | 1.41666667 | 2.23333333 | 2.75 | 1.66666667 | 2.61666667 | 1.18333333 | 1.9 | 0.96666667 | 1.38333333 | 1.89880952 | 0.70932327 | 0.18957462 |
| 4 | 2.86666667 | 2.86666667 | 2.28333333 | 2.45 | 1 | 2.01666667 | 2.3 | 2.36666667 | 2.15 | 2.56666667 | 1.05 | 1.65 | 0.96666667 | 1.33333333 | 1.99047619 | 0.67306907 | 0.17988528 |
| 5 | 2.1 | 2.26666667 | 2.78333333 | 1.96666667 | 1.08333333 | 1.91666667 | 1.85 | 1.53333333 | 2.01666667 | 2.06666667 | 0.95 | 1.26666667 | 1.33333333 | 1.08333333 | 1.7297619 | 0.53281941 | 0.14240198 |
| 10 | 2.06666667 | 1.55 | 1.17 | 1.49 | 1.59333333 | 1.51 | 1.52666667 | 1.38 | 1.72333333 | 2.21 | 1.16333333 | 1.00333333 | 1.24333333 | 1.01333333 | 1.47452381 | 0.35778905 | 0.09562315 |
| 15 | 1.76666667 | 1.03333333 | 0.86333333 | 1.31333333 | 1.29666667 | 1.64666667 | 1.31333333 | 1.46333333 | 1.23 | 1.93 | 1.07 | 0.89333333 | 0.76333333 | 0.91333333 | 1.2497619 | 0.35616059 | 0.09518792 |

Week 4

WT

| Time (min) | Animal ID | | | | | | | | | | | |
|---|---|---|---|---|---|---|---|---|---|---|---|---|
| | 83 | 86 | 87 | 88 | 81 | 82 | A | 156 | 158 | 161 | Average | SD | SE |
| 0 | 2.50777778 | 1.66666667 | 2.07111111 | 2.39444444 | 1.24666667 | 1.69888889 | 2.20111111 | 2.14 | 0.91 | 1.44777778 | 1.82844444 | 0.52138209 | 0.16487549 |
| 1 | 3.35 | 3.68333333 | 4.06666667 | 4.43333333 | 3.26666667 | 2.8 | 3.38333333 | 3.48333333 | 1.6 | 2.15 | 3.22166667 | 0.84751088 | 0.26800647 |
| 2 | 2.51666667 | 3.1 | 3.8 | 3.86666667 | 2.56666667 | 2.05 | 2.6 | 2.43333333 | 0.81666667 | 1.48333333 | 2.52333333 | 0.94288106 | 0.29816517 |
| 3 | 2.7 | 3.13333333 | 4.08333333 | 3.95 | 2.35 | 1.93333333 | 2.76666667 | 2.5 | 1.1 | 2.3 | 2.68166667 | 0.89039629 | 0.28156803 |
| 4 | 2.81666667 | 3.8 | 4.03333333 | 4.1 | 2.66666667 | 2.01666667 | 2.63333333 | 2.81666667 | 1.2 | 2.26666667 | 2.835 | 0.9251443 | 0.29255632 |
| 5 | 2.6 | 3.5 | 3.13333333 | 3.11666667 | 3.23333333 | 2.18333333 | 2.26666667 | 2.73333333 | 1.31666667 | 1.6 | 2.56833333 | 0.72279678 | 0.22856841 |
| 10 | 1.96666667 | 2.22666667 | 2.29666667 | 2.28666667 | 2.30666667 | 1.83666667 | 1.94 | 2.39333333 | 1.27666667 | 1.54 | 2.007 | 0.37058995 | 0.11719083 |
| 15 | 1.57666667 | 1.56333333 | 1.82666667 | 1.48333333 | 1.61666667 | 1.58666667 | 1.67333333 | 1.28666667 | 1.03666667 | 0.97666667 | 1.46266667 | 0.27652071 | 0.08744353 |

Het

| Time (min) | Animal ID | | | | | | | | | | | | | | | |
|---|---|---|---|---|---|---|---|---|---|---|---|---|---|---|---|---|
| | B | E | H | 155 | 157 | 190 | 196 | 199 | 201 | 205 | 207 | 208 | 210 | Average | SD | SE |
| 0 | 1.47777778 | 0.88444444 | 0.99222222 | 1.26333333 | 1.29333333 | 1.60222222 | 1.11888889 | 1.47222222 | 1.54111111 | 0.85222222 | 1.23888889 | 1.15888889 | 1.09222222 | 1.22982906 | 0.24420858 | 0.06773127 |
| 1 | 3.66666667 | 2 | 2.3 | 2.4 | 2.23333333 | 2.63333333 | 2.63333333 | 2.3 | 2.68333333 | 1.68333333 | 1.33333333 | 1.53333333 | 1.5 | 2.22307692 | 0.63291428 | 0.17553884 |
| 2 | 3.03333333 | 1.9 | 2.23333333 | 1.61666667 | 1.51666667 | 1.81666667 | 2.41666667 | 1.63333333 | 2.4 | 1.38333333 | 1.45 | 1.13333333 | 1.2 | 1.82564103 | 0.55476118 | 0.15386307 |
| 3 | 4.48333333 | 3.18333333 | 2.38333333 | 1.53333333 | 1.81666667 | 2.35 | 2.83333333 | 1.46666667 | 2.13333333 | 1.9 | 1.58333333 | 1.25 | 1.23333333 | 2.16538462 | 0.91574351 | 0.25398155 |
| 4 | 4.1 | 2.45 | 2.11666667 | 1.5 | 1.85 | 2.2 | 2.46666667 | 2.65 | 2.96666667 | 2.1 | 1.93333333 | 1.23333333 | 1.91666667 | 2.26794872 | 0.71816579 | 0.19918335 |
| 5 | 3.1 | 1.15 | 1.9 | 0.96666667 | 2.06666667 | 2.06666667 | 2.21666667 | 2.21666667 | 2.91666667 | 1.6 | 1.31666667 | 1.56666667 | 1.33333333 | 1.87820513 | 0.64676491 | 0.17938031 |
| 10 | 2.58666667 | 1.00333333 | 1.31333333 | 1.13666667 | 1.75666667 | 1.77666667 | 1.91333333 | 1.71333333 | 2.13666667 | 1.20666667 | 0.86333333 | 0.94666667 | 0.77333333 | 1.47128205 | 0.55327734 | 0.15345152 |
| 15 | 2.77 | 1.11666667 | 0.95666667 | 1.23 | 1.09333333 | 1.25333333 | 2.03 | 1.22666667 | 1.78333333 | 2.11 | 0.75 | 0.88275862 | 0.83666667 | 1.3876481 | 0.60426032 | 0.16759166 |

### Minute ventilation

Week 1

WT

| Time (min) | 83 | 86 | 87 | 88 | 81 | 82 | A | 156 | 158 | 161 | Average | SD | SE |
|---|---|---|---|---|---|---|---|---|---|---|---|---|---|
| 0 | 51.9222222 | 45.1888889 | 39.2555556 | 56.8555556 | 36.8222222 | 35.9 | 38.9111111 | 53.4555556 | 23.3333333 | 25.0333333 | 40.6677778 | 11.3538708 | 3.59040921 |
| 1 | 49.3333333 | 70.3333333 | 54.1666667 | 83.6666667 | 52 | 61.6666667 | 17 | 58.8 | 28.2 | 35.8 | 51.0966667 | 19.8217018 | 6.26817249 |
| 2 | 46.3333333 | 62.1666667 | 50.6666667 | 44.8333333 | 45 | 46.8333333 | 25.8333333 | 52.5 | 24 | 31.1666667 | 42.9333333 | 12.2258076 | 3.86613981 |
| 3 | 47.1666667 | 57.8333333 | 52 | 35.5 | 42.6666667 | 48.5 | 35.8333333 | 54.3333333 | 30.8333333 | 35.5 | 44.0166667 | 9.30201425 | 2.94155519 |
| 4 | 48.1666667 | 54.1666667 | 64.8333333 | 53.3333333 | 37.1666667 | 50.1666667 | 31.6666667 | 70.3333333 | 37.1666667 | 38.3333333 | 48.5333333 | 12.6729517 | 4.00753919 |
| 5 | 46 | 57.3333333 | 51.6666667 | 43.8333333 | 37.1666667 | 49.3333333 | 25.3333333 | 59.5 | 34.1666667 | 41.8333333 | 44.6166667 | 10.5514758 | 3.33666963 |
| 10 | 31.6666667 | 44.6333333 | 35.0666667 | 39.2333333 | 39 | 40.5 | 20.8333333 | 49.2333333 | 35.7 | 33.1333333 | 36.9 | 7.74054581 | 2.44777551 |
| 15 | 31.2666667 | 45.7333333 | 38.1666667 | 34.8333333 | 41 | 35.037037 | 14.8666667 | 50.4666667 | 22.1 | 26.2333333 | 33.9703704 | 10.8137316 | 3.41960217 |

Het

| Time (min) | B | D | E | F | G | H | 155 | 157 | 159 | 160 | 162 |
|---|---|---|---|---|---|---|---|---|---|---|---|
| 0 | 25.2 | 29.6333333 | 35.9333333 | 14.2777778 | 22.6666667 | 39.5222222 | 37.8444444 | 42.5111111 | 21.3888889 | 42.9444444 | 31.5111111 |
| 1 | 52.5 | 60 | 52 | 20.2 | 33 | 40.6 | 48.2 | 43.6666667 | 45.1666667 | 37.8 | 42 |
| 2 | 37.1666667 | 49.3333333 | 35.1666667 | 20.8333333 | 28.6666667 | 39.3333333 | 38 | 35.1666667 | 31 | 31.5 | 32.3333333 |
| 3 | 29.8333333 | 52.5 | 23.1666667 | 12.3333333 | 25.6666667 | 39 | 44.6666667 | 30.3333333 | 26.3333333 | 25.1666667 | 26.3333333 |
| 4 | 31.8333333 | 60 | 20.5 | 10.6666667 | 25.3333333 | 31.3333333 | 48.3333333 | 25.3333333 | 24 | 31.8333333 | 26 |
| 5 | 27.1666667 | 56.1666667 | 21.5 | 22.1666667 | 27.6666667 | 32 | 40 | 23.1666667 | 29.8333333 | 41 | 24.5 |
| 10 | 37.7666667 | 47.6666667 | 26.7333333 | 11.6 | 25.9 | 36.6666667 | 37.3333333 | 28.3666667 | 28.3666667 | 41.9666667 | 33.5 |
| 15 | 31.2 | 40.2 | 29.4333333 | 15.0333333 | 29.4 | 35.4 | 22.8333333 | 31.1666667 | 22.7 | 36.4666667 | 23.1666667 |

| Time (min) | 190 | 196 | 199 | 201 | 205 | 207 | 208 | 210 | Average | SD | SE |
|---|---|---|---|---|---|---|---|---|---|---|---|
| 0 | 46.9444444 | 25.2222222 | 27.8555556 | 37.1222222 | 32.4888889 | 26.4444444 | 17.4555556 | 15.5111111 | 30.1304094 | 9.61233193 | 2.20522018 |
| 1 | 45.3333333 | 50.8333333 | 56.5 | 52.5 | 30.1666667 | 37.6666667 | 38 | 29.5 | 42.9280702 | 10.2351926 | 2.34811423 |
| 2 | 40.8333333 | 46.6666667 | 49.5 | 52.6666667 | 30.3333333 | 29.5 | 21.6666667 | 22.8333333 | 35.3947368 | 9.37346045 | 2.15041931 |
| 3 | 45.3333333 | 45.1666667 | 44 | 45.3333333 | 27.3333333 | 30.8333333 | 19 | 19.6666667 | 32.2105263 | 11.2282495 | 2.5759371 |
| 4 | 59.6666667 | 47.5 | 51.1666667 | 54 | 31.1666667 | 37 | 19.5 | 22.5 | 34.6140351 | 14.5810901 | 3.34513148 |
| 5 | 66.5 | 49.6666667 | 57.8333333 | 64.8333333 | 37.1666667 | 35.3333333 | 21.8333333 | 21.1666667 | 36.8157895 | 15.2437899 | 3.49716526 |
| 10 | 56.0666667 | 32.6333333 | 49.2333333 | 46.2333333 | 31.8333333 | 29.5333333 | 24.8666667 | 18.1666667 | 33.9175439 | 11.0445862 | 2.53380186 |
| 15 | 44.4 | 29.5 | 44.5 | 38.9666667 | 23.2 | 18.9333333 | 19.4333333 | 17.4666667 | 29.1263158 | 9.05248468 | 2.07678242 |

Week 2

WT

| Time (min) | 83 | 86 | 87 | 88 | 81 | 82 | A | 156 | 158 | 161 | Average | SD | SE |
|---|---|---|---|---|---|---|---|---|---|---|---|---|---|
| 0 | 69.0444444 | 70.9222222 | 53.6555556 | 77.5888889 | 66.5444444 | 40.5888889 | 62.7777778 | 44.9111111 | 46.1555556 | 51.3 | 58.3488889 | 12.674675 | 4.00808415 |
| 1 | 77.1666667 | 103 | 93.3333333 | 118.166667 | 89.6666667 | 95.1666667 | 72.1666667 | 60.5 | 32.3333333 | 53.1666667 | 79.4666667 | 25.6599077 | 8.11437529 |
| 2 | 60.3333333 | 103.333333 | 90.1666667 | 123.833333 | 83.8333333 | 98.3333333 | 53 | 78 | 34.1666667 | 51.8333333 | 77.6833333 | 27.648862 | 8.74333785 |
| 3 | 65.1666667 | 97 | 91.8333333 | 113 | 76 | 83.5 | 53.5 | 59.6666667 | 37 | 56.3333333 | 73.3 | 23.1953327 | 7.33500823 |
| 4 | 59.1666667 | 97.3333333 | 99 | 102.5 | 82.6666667 | 81 | 65.1666667 | 55.3333333 | 40.6666667 | 72 | 75.4833333 | 20.6704671 | 6.53657563 |
| 5 | 70 | 93.6666667 | 78.1666667 | 85.6666667 | 69.1666667 | 81.8333333 | 54 | 43.8333333 | 41.5 | 64.1666667 | 68.2 | 17.5358187 | 5.54531278 |
| 10 | 58.1 | 76.8333333 | 64.4666667 | 84.7333333 | 50.6 | 53.9 | 53.1333333 | 47.6333333 | 32.9666667 | 43.6666667 | 56.6033333 | 15.3714056 | 4.86086526 |
| 15 | 58.3666667 | 58 | 52.4333333 | 75.0333333 | 43.5333333 | 49.4 | 54.2333333 | 43.4666667 | 39.9333333 | 32.3666667 | 50.6766667 | 11.9120237 | 3.76691264 |

Het

| Time (min) | B | D | E | G | H | 155 | 157 | 160 | 162 | 190 |
|---|---|---|---|---|---|---|---|---|---|---|
| 0 | 52.4444444 | 32.2777778 | 29.0222222 | 18.3 | 46.7555556 | 30.6555556 | 40.6111111 | 32.3333333 | 30.0888889 | 33.4333333 |
| 1 | 71.3333333 | 62 | 55.8 | 33 | 44 | 59 | 50 | 41.6666667 | 43.1666667 | 71.1666667 |
| 2 | 57.5 | 67.1666667 | 51.1666667 | 35 | 47.5 | 46 | 38.6666667 | 41.3333333 | 45.3333333 | 64 |
| 3 | 42.3333333 | 62.5 | 47.3333333 | 33.8333333 | 57.6666667 | 41.8333333 | 28.6666667 | 40.5 | 41.5 | 67.5 |
| 4 | 69.5 | 65.3333333 | 56.5 | 34.5 | 52.8333333 | 36 | 26.6666667 | 43.6666667 | 33.6666667 | 64.1666667 |
| 5 | 71.8333333 | 73.1666667 | 49.8333333 | 40.1666667 | 41.6666667 | 34.5 | 26.3333333 | 42 | 26.8333333 | 56.5 |
| 10 | 37.1333333 | 64.9333333 | 45.9666667 | 33.9666667 | 38.6666667 | 32.3666667 | 33.8333333 | 31.9333333 | 25.1 | 48.3 |
| 15 | 45.6333333 | 54.8333333 | 35.3333333 | 25.6666667 | 36.0666667 | 40.2333333 | 33.6333333 | 21.6666667 | 15.3333333 | 44.4666667 |

| Time (min) | 196 | 199 | 201 | 205 | 207 | 208 | 210 | Average | SD | SE |
|---|---|---|---|---|---|---|---|---|---|---|
| 0 | 35.1111111 | 26.9333333 | 35.9777778 | 43.3444444 | 25.7555556 | 50.7333333 | 38.3 | 35.4163399 | 9.10078778 | 2.20726525 |
| 1 | 59.8333333 | 52.8333333 | 57.1666667 | 36.3333333 | 48.6666667 | 50.6666667 | 46.3333333 | 51.9392157 | 10.8901219 | 2.64124252 |
| 2 | 69.1666667 | 49.5 | 61.1666667 | 33.5 | 55 | 40 | 53.3333333 | 50.3137255 | 10.9541155 | 2.65676326 |
| 3 | 74.5 | 52.1666667 | 74.5 | 42.3333333 | 62.8333333 | 29.8333333 | 49.5 | 49.9607843 | 14.5095191 | 3.51907528 |
| 4 | 64.8333333 | 66.6666667 | 65.3333333 | 38.3333333 | 55 | 46.6666667 | 53.5 | 51.3627451 | 13.7834486 | 3.34297733 |
| 5 | 52.1666667 | 62 | 55.5 | 35.3333333 | 46.8333333 | 58.6666667 | 42.1666667 | 47.9705882 | 13.90459 | 3.37235842 |
| 10 | 38.5666667 | 45.8666667 | 38.3666667 | 31.9 | 35.5666667 | 45.8666667 | 38.3 | 39.2137255 | 9.03859056 | 2.19218021 |
| 15 | 54.0333333 | 41.5666667 | 43.8333333 | 30.0333333 | 23.0333333 | 37.9333333 | 32.8 | 36.2411765 | 10.9452014 | 2.65460127 |

Week 3

WT

| Time (min) | 83 | 86 | 87 | 88 | 81 | 82 | A | 156 | 158 | 161 | Average | SD | SE |
|---|---|---|---|---|---|---|---|---|---|---|---|---|---|
| 0 | 80.5888889 | 73.6777778 | 61.6111111 | 63.1444444 | 42.1444444 | 39.9444444 | 68.7 | 55.8444444 | 27.5888889 | 46.3 | 55.9544444 | 16.706542 | 5.28307244 |
| 1 | 88.5 | 100 | 118.166667 | 106.5 | 107.666667 | 81.5 | 88.6666667 | 84.1666667 | 39.1666667 | 51.3333333 | 86.5666667 | 24.8311831 | 7.85230956 |
| 2 | 92.6666667 | 83.8333333 | 97 | 95.3333333 | 94.5 | 55.1666667 | 68.3333333 | 50.1666667 | 26.3333333 | 41.5 | 70.4833333 | 25.8393004 | 8.17110424 |
| 3 | 101.333333 | 74.6666667 | 109 | 111 | 107 | 55.5 | 79.1666667 | 48.5 | 26.5 | 59 | 77.1666667 | 29.4971227 | 9.32780921 |
| 4 | 83.3333333 | 69 | 95.1666667 | 108.333333 | 122.666667 | 56.8333333 | 79.1666667 | 49.1666667 | 27 | 69 | 75.9666667 | 28.3642315 | 8.96955757 |
| 5 | 78.6666667 | 65.8333333 | 83.1666667 | 90.5 | 107.666667 | 65.6666667 | 83.8333333 | 45.8333333 | 36.3333333 | 55.3333333 | 71.2833334 | 21.6823377 | 6.85655721 |
| 10 | 73.5666667 | 54.2 | 61.2333333 | 51 | 78.1333333 | 57.6333333 | 55.6333333 | 51.8666667 | 37.1666667 | 46.6333333 | 56.7066667 | 12.071186 | 3.81724417 |
| 15 | 69.7333333 | 43.9 | 38.4 | 48.7 | 44.9333333 | 43.2333333 | 36.7666667 | 58.1333333 | 31.9 | 29.2666667 | 44.4966667 | 12.1573475 | 3.84449085 |

Het

| Time (min) | B | D | E | H | 155 | 157 | 190 | 196 | 199 | 201 | 205 | 207 | 208 | 210 | Average | SD | SE |
|---|---|---|---|---|---|---|---|---|---|---|---|---|---|---|---|---|---|
| 0 | 42.7444444 | 34.1111111 | 25.4 | 32.3444444 | 32.6222222 | 34.7555556 | 35.2222222 | 33.6111111 | 33.2 | 35.9666667 | 24.2555556 | 25.2777778 | 23.9777778 | 25.8444444 | 31.3809524 | 5.57299899 | 1.48944663 |
| 1 | 87 | 73.6666667 | 52.8 | 68.5 | 62.8333333 | 60.3333333 | 65.1666667 | 84 | 69 | 61.1666667 | 44.5 | 46.8333333 | 37.8333333 | 52.5 | 61.8666667 | 14.2653486 | 3.81257479 |
| 2 | 93.6666667 | 70.6666667 | 53.6666667 | 56.1666667 | 39.1666667 | 46.6666667 | 64.8333333 | 82.5 | 59.5 | 50.3333333 | 33.8333333 | 50.1666667 | 42 | 49 | 56.5833333 | 16.6412306 | 4.44755595 |
| 3 | 98.6666667 | 86 | 51.5 | 67.6666667 | 33.6666667 | 40.3333333 | 72.3333333 | 89.5 | 53 | 72 | 36.5 | 60.1666667 | 34.5 | 49 | 60.3452381 | 21.3389881 | 5.70308445 |
| 4 | 99 | 80.6666667 | 74.5 | 73 | 33.5 | 63 | 73 | 80.8333333 | 63.5 | 77.6666667 | 33.8333333 | 54 | 33.3333333 | 48.8333333 | 63.4761905 | 20.2927145 | 5.42345608 |
| 5 | 65.6666667 | 70.8333333 | 84.3333333 | 63.1666667 | 32.1666667 | 60.1666667 | 58 | 55.8333333 | 59.3333333 | 64.3333333 | 31.3333333 | 44.3333333 | 46.8333333 | 35.1666667 | 55.1071429 | 15.4148623 | 4.11979524 |
| 10 | 70.2 | 49.3666667 | 44.3333333 | 45.3666667 | 50.3333333 | 47.9333333 | 44.7666667 | 46.5 | 57.2666667 | 57.8666667 | 40.1666667 | 32.6 | 45.5666667 | 33.2 | 47.5333333 | 9.75427147 | 2.60693871 |
| 15 | 58.4 | 30.6666667 | 31.5 | 38.4666667 | 41.9666667 | 51.7666667 | 38.7666667 | 50.2666667 | 41.1 | 55.7 | 40.3333333 | 26 | 24.7333333 | 27.3333333 | 39.7857143 | 11.0460707 | 2.95218657 |

Week 4

WT

| Time (min) | 83 | 86 | 87 | 88 | 81 | 82 | A | 156 | 158 | 161 | Average | SD | SE |
|---|---|---|---|---|---|---|---|---|---|---|---|---|---|
| 0 | 90.1555556 | 53.8111111 | 64.4111111 | 76.5777778 | 39.9555556 | 53.0555556 | 73.9 | 73.7666667 | 29.2333333 | 53.0222222 | 60.7888889 | 18.4503248 | 5.834505 |
| 1 | 96.3333333 | 104 | 115.833333 | 124 | 92.6666667 | 79.6666667 | 98.1666667 | 97.1666667 | 51.5 | 61 | 92.0333333 | 22.5421142 | 7.12844242 |
| 2 | 75.3333333 | 86.1666667 | 113.5 | 109.666667 | 79.3333333 | 64 | 84.5 | 74 | 25.5 | 46.3333333 | 75.8333333 | 26.4678934 | 8.36988281 |
| 3 | 82 | 93.3333333 | 128.333333 | 130 | 74 | 64.8333333 | 95.5 | 77.8333333 | 33.8333333 | 65 | 84.4666667 | 29.2227292 | 9.24103837 |
| 4 | 93.5 | 116.166667 | 126.5 | 126.833333 | 84.1666667 | 64.8333333 | 92 | 83.5 | 40.3333333 | 78.1666667 | 90.6 | 27.2900742 | 8.62987919 |
| 5 | 86.6666667 | 101.166667 | 99 | 104.833333 | 98 | 70 | 76.5 | 84.1666667 | 42.1666667 | 55.5 | 81.8 | 20.8779227 | 6.60217884 |
| 10 | 62.7666667 | 68.2666667 | 72.7333333 | 71 | 74.4666667 | 57 | 59.3333333 | 79.2 | 42.0333333 | 50.1666667 | 63.6966667 | 11.6873198 | 3.69585504 |
| 15 | 45.7333333 | 42.3666667 | 53.5666667 | 41.6666667 | 54.6333333 | 45.9 | 48.3 | 41.1 | 32.4666667 | 32.4333333 | 43.8166667 | 7.54751615 | 2.38673417 |

Het

| Time (min) | B | E | H | 155 | 157 | 190 | 196 | 199 | 201 | 205 | 207 | 208 | 210 | Average | SD | SE |
|---|---|---|---|---|---|---|---|---|---|---|---|---|---|---|---|---|
| 0 | 47.4222222 | 29.7 | 32.9666667 | 37.8555556 | 37.8666667 | 45.5333333 | 34.1888889 | 45.8 | 47.6888889 | 29.3555556 | 42.3444444 | 37.6888889 | 37.9222222 | 38.9487179 | 6.3968365 | 1.77416323 |
| 1 | 101.666667 | 63.6666667 | 70.3333333 | 68.1666667 | 56 | 72 | 76.6666667 | 66.5 | 71.5 | 53 | 44.3333333 | 44 | 47 | 64.2179487 | 15.8731057 | 4.40240742 |
| 2 | 90 | 61.3333333 | 69.3333333 | 52.6666667 | 45.8333333 | 56.1666667 | 81.6666667 | 50.5 | 64 | 45.3333333 | 48.3333333 | 38.6666667 | 46.5 | 57.7179487 | 15.1509969 | 4.20213047 |
| 3 | 125.5 | 83.3333333 | 81 | 50.1666667 | 53.5 | 72.6666667 | 85.1666667 | 47.1666667 | 57.6666667 | 66.3333333 | 58.6666667 | 44.8333333 | 45.1666667 | 67.0128205 | 22.7874094 | 6.32009022 |
| 4 | 115.666667 | 70.1666667 | 71.8333333 | 52.5 | 54.8333333 | 66.3333333 | 85.6666667 | 83 | 90.3333333 | 73.5 | 66.1666667 | 42.8333333 | 65.8333333 | 72.2051282 | 18.676108 | 5.17982038 |
| 5 | 106 | 39.6666667 | 59.3333333 | 36.1666667 | 63.3333333 | 60.1666667 | 72.1666667 | 69 | 89.6666667 | 58.5 | 52.1666667 | 57 | 48 | 62.3974359 | 19.0614914 | 5.28670652 |
| 10 | 79.2666667 | 35.5333333 | 43.2333333 | 34.2666667 | 54.1 | 55.2 | 63.6333333 | 55.4333333 | 66.6 | 42.7666667 | 26.4333333 | 35.3333333 | 26.2666667 | 47.5435897 | 16.3340044 | 4.53023773 |
| 15 | 80.3 | 39 | 28.8 | 35.8333333 | 34.4666667 | 35.3666667 | 65.9 | 37.7 | 55.4333333 | 63.9 | 22.9 | 29.8965517 | 25.2666667 | 42.6740937 | 17.8514125 | 4.951091 |

### Table S2. Respiratory parameters following sham stimulation

#### Time of inspiration

Week 1

WT

| Time (min) | A | 156 | 158 | 161 | Average | SD | SEM |
|---|---|---|---|---|---|---|---|
| 0 | 53.8333333 | 50.8111111 | 57.3666667 | 50.0222222 | 53.0083333 | 3.33768081 | 1.6688404 |
| 1 | 74.8333333 | 56.6666667 | 53.1666667 | 55.6666667 | 60.0833333 | 9.94289249 | 4.97144625 |
| 2 | 57.6666667 | 54.8333333 | 48.8333333 | 51.1666667 | 53.125 | 3.90719896 | 1.95359948 |
| 3 | 59.1666667 | 50.1666667 | 52.8333333 | 51 | 53.2916667 | 4.07197968 | 2.03598984 |
| 4 | 58.5 | 61.3333333 | 53.8333333 | 59.5 | 58.2916667 | 3.19541048 | 1.59770524 |
| 5 | 70.5 | 62.6666667 | 48.8333333 | 59.5 | 60.375 | 8.97669307 | 4.48834654 |
| 10 | 63.6666667 | 60.1333333 | 58.1 | 55 | 59.225 | 3.63637742 | 1.81818871 |
| 15 | 83.3 | 79.9666667 | 76.7 | 75.4666667 | 78.8583333 | 3.51750911 | 1.75875456 |

Het

| Time (min) | B | C | D | E | F | G | H | 155 | 157 | 159 | 160 | 162 | 190 | 196 | 199 | 201 | 205 | 207 | 208 | 210 | Average | SD | SEM |
|---|---|---|---|---|---|---|---|---|---|---|---|---|---|---|---|---|---|---|---|---|---|---|---|
| 0 | 81.3111111 | 61.0111111 | 58.7333333 | 53.5333333 | 68.9777778 | 64.4 | 48.9444444 | 51.6666667 | 48.3555556 | 56.3111111 | 53.2666667 | 69.5111111 | 60.0666667 | 79.8333333 | 93.5111111 | 44.6888889 | 48.1888889 | 59.7555556 | 57.0111111 | 85.1777778 | 62.2127778 | 13.5927939 | 3.03944111 |
| 1 | 56.5 | 66.8333333 | 62.6666667 | 55.1666667 | 83.3333333 | 47.5 | 54.5 | 55.8333333 | 68.6666667 | 61 | 55.1666667 | 69.8333333 | 63.8333333 | 60.5 | 62 | 67.5 | 62 | 63.8333333 | 58 | 60 | 61.7333333 | 7.51030481 | 1.67935521 |
| 2 | 59.6666667 | 65.1666667 | 65.1666667 | 57 | 76.8333333 | 49.1666667 | 53.5 | 53.6666667 | 67.1666667 | 62.1666667 | 56.8333333 | 60.3333333 | 52.5 | 65.5 | 57.5 | 54.3333333 | 72 | 64.6666667 | 58 | 59.8333333 | 60.55 | 6.88398005 | 1.53930473 |
| 3 | 63.3333333 | 63.8333333 | 63.1666667 | 50.1666667 | 81 | 46.1666667 | 51.5 | 51.6666667 | 60.5 | 56.1666667 | 55 | 59 | 49.6666667 | 66.3333333 | 64.1666667 | 67.5 | 76.5 | 62.1666667 | 67.1666667 | 64.5 | 60.975 | 8.87803568 | 1.98518913 |
| 4 | 61.1666667 | 60.3333333 | 65 | 49.1666667 | 79.6666667 | 41.6666667 | 59.3333333 | 53 | 62.6666667 | 59.6666667 | 61.8333333 | 61.3333333 | 47.8333333 | 62.8333333 | 50.6666667 | 60.1666667 | 78 | 49.5 | 66.1666667 | 58.8333333 | 59.4416667 | 9.28815428 | 2.07689444 |
| 5 | 67.3333333 | 62.1666667 | 62.3333333 | 53.6666667 | 89.3333333 | 41.6666667 | 56.5 | 52.6666667 | 61.5 | 61.8333333 | 58.3333333 | 58.3333333 | 60.3333333 | 79.1666667 | 51.6666667 | 60.1666667 | 75.3333333 | 46.1666667 | 50.3333333 | 62.8333333 | 60.5833333 | 11.0878338 | 2.47931501 |
| 10 | 81.5 | 68.7333333 | 74.0333333 | 88.3 | 88.4666667 | 42.2333333 | 54.4 | 58.7666667 | 79.6666667 | 64.1666667 | 71.8333333 | 58.7666667 | 58.1666667 | 84.2 | 53.4 | 53.2 | 67.9333333 | 54.8666667 | 78.4 | 71.6666667 | 67.635 | 13.214459 | 2.95484286 |
| 15 | 84.4 | 81.2 | 68 | 86.7333333 | 85.0333333 | 57.7666667 | 62.5666667 | 70.7333333 | 89.6666667 | 83.5666667 | 80.7333333 | 73.9666667 | 75.1666667 | 88.2 | 83.7 | 69.5 | 56.8 | 85.1 | 94.1333333 | 84.9333333 | 78.095 | 10.7433933 | 2.40229578 |

Week 2

WT

| Time (min) | A | 156 | 158 | 161 | Average | SD | SEM |
|---|---|---|---|---|---|---|---|
| 0 | 61.5 | 81.0222222 | 50.2888889 | 48.2111111 | 60.2555556 | 15.024493 | 7.51224652 |
| 1 | 56.1666667 | 65.3333333 | 54.3333333 | 54.8333333 | 57.6666667 | 5.16935414 | 2.58467707 |
| 2 | 57.5 | 56.8333333 | 51 | 51.6666667 | 54.25 | 3.38979951 | 1.69489976 |
| 3 | 46.1666667 | 50.1666667 | 55.5 | 58 | 52.4583333 | 5.316597 | 2.6582985 |
| 4 | 48.6666667 | 64.8333333 | 55.5 | 55.1666667 | 56.0416667 | 6.65189335 | 3.32594668 |
| 5 | 54.5 | 57.6666667 | 55.8333333 | 56 | 56 | 1.29814683 | 0.64907341 |
| 10 | 53.4666667 | 64.4666667 | 62.7666667 | 60.0666667 | 60.1916667 | 4.83554547 | 2.41777274 |
| 15 | 69.1333333 | 79.6333333 | 73.8666667 | 73.1333333 | 73.9416667 | 4.32720721 | 2.1636036 |

Het

| Time (min) | B | D | E | G | H | 155 | 157 | 160 | 162 | 190 | 196 | 199 | 201 | 205 | 207 | 208 | 210 | Average | SD | SEM |
|---|---|---|---|---|---|---|---|---|---|---|---|---|---|---|---|---|---|---|---|---|
| 0 | 90.6222222 | 68.9 | 86.8666667 | 84.0888889 | 53.8111111 | 81.3111111 | 89.1111111 | 47.9444444 | 60.8777778 | 63.1888889 | 93.4777778 | 96.4555556 | 77.1 | 39.5555556 | 49.6777778 | 41.7 | 41.5222222 | 68.6006536 | 20.1567781 | 4.88873677 |
| 1 | 53.1666667 | 59.1666667 | 57.8333333 | 49.3333333 | 50.1666667 | 68.8333333 | 72.1666667 | 50.6666667 | 73.6666667 | 78.6 | 59.6 | 77.6 | 92.8 | 84.2 | 79.8 | 68.8 | 72.4 | 67.5764706 | 13.0557712 | 3.16648963 |
| 2 | 50.3333333 | 50.3333333 | 49 | 50.1666667 | 50.3333333 | 54.3333333 | 63 | 57.5 | 82.3333333 | 55.8333333 | 51 | 54.6666667 | 74.1666667 | 81.6666667 | 55.5 | 64.5 | 77.5 | 60.127451 | 11.6929064 | 2.83594637 |
| 3 | 49.1666667 | 53.3333333 | 51.5 | 50.3333333 | 50.5 | 56.3333333 | 72.1666667 | 50.8333333 | 64.5 | 58 | 58 | 56.3333333 | 61.8333333 | 63.3333333 | 64 | 78 | 60.5 | 58.745098 | 8.03467262 | 1.94869434 |
| 4 | 50 | 49.5 | 44.1666667 | 53.6666667 | 55.1666667 | 51.8333333 | 74.3333333 | 59 | 57.1666667 | 66.5 | 56.1666667 | 58.1666667 | 79.3333333 | 72.6666667 | 78.6666667 | 76.5 | 74 | 62.1666667 | 11.5569243 | 2.80296587 |
| 5 | 55.8333333 | 54.1666667 | 50 | 45.5 | 68.1666667 | 56.8333333 | 75.1666667 | 61.5 | 66.3333333 | 54.1666667 | 60.6666667 | 47.6666667 | 81 | 68 | 82.5 | 85.3333333 | 68.5 | 63.6078431 | 12.250842 | 2.97126562 |
| 10 | 55.6666667 | 54.6 | 60.7666667 | 51.8333333 | 73.2333333 | 58.7 | 75.7 | 68.2333333 | 68.5333333 | 71.6 | 53.0333333 | 54.2333333 | 74.3 | 60.1666667 | 70.8333333 | 76 | 67.1666667 | 64.3882353 | 8.65069796 | 2.09810244 |
| 15 | 51.5666667 | 86.3 | 90.8 | 55.3333333 | 67.6 | 66.1333333 | 64.0333333 | 87.4 | 114.033333 | 80.3333333 | 64.7 | 67.1333333 | 57.1 | 77.1666667 | 67.0333333 | 67.3666667 | 93.3 | 73.9607843 | 16.2096132 | 3.93140867 |

Week 3

WT

| Time (min) | 83 | 86 | 87 | 88 | 81 | 82 | A | 156 | 158 | 161 | Average | SD | SEM |
|---|---|---|---|---|---|---|---|---|---|---|---|---|---|
| 0 | 52.8666667 | 88.1777778 | 65.9888889 | 50.5888889 | 78.741573 | 88.4222222 | 50.9888889 | 66.8444444 | 51.1777778 | 67.5111111 | 66.130824 | 14.9621805 | 4.7314569 |
| 1 | 56 | 59.3333333 | 50 | 48.8333333 | 54.1666667 | 53 | 46.6666667 | 50.6 | 56.1666667 | 53.3333333 | 52.81 | 3.83669901 | 1.21327076 |
| 2 | 55 | 50.5 | 41.1666667 | 48 | 53.5 | 52.6666667 | 46.8333333 | 49.8333333 | 55.5 | 47.1666667 | 50.0166667 | 4.40920054 | 1.39431164 |
| 3 | 53 | 52.3333333 | 47.3333333 | 50.3333333 | 56.6666667 | 62.8333333 | 53.5 | 54 | 49 | 49.3333333 | 52.8333333 | 4.47489568 | 1.41508626 |
| 4 | 47.3333333 | 50.3333333 | 45.6666667 | 48 | 54.6666667 | 54 | 40.1666667 | 43.6666667 | 47.8333333 | 51 | 48.2666667 | 4.47503362 | 1.41512989 |
| 5 | 54.6666667 | 52.8333333 | 43.3333333 | 41 | 45.3333333 | 42.8333333 | 49.3333333 | 44.6666667 | 53.8333333 | 45.6666667 | 47.35 | 4.95002806 | 1.56533631 |
| 10 | 62.7333333 | 50 | 51.0666667 | 52.6666667 | 44.7 | 52.3333333 | 49.7333333 | 49.9666667 | 60.4333333 | 47.1666667 | 52.08 | 5.55389308 | 1.7562952 |
| 15 | 56.0666667 | 66.4 | 78.2666667 | 80.1 | 59.7333333 | 64.2666667 | 74.6333333 | 85.1666667 | 77.2666667 | 53.4333333 | 69.5333333 | 11.0166429 | 3.48376836 |

Het

| Time (min) | B | D | E | G | H | 155 | 157 | 190 | 196 | 199 | 201 | 205 | 207 | 208 | 210 | Average | SD | SEM |
|---|---|---|---|---|---|---|---|---|---|---|---|---|---|---|---|---|---|---|
| 0 | 88.7444444 | 64.4555556 | 77.2222222 | 95.4444444 | 55.4888889 | 48.3555556 | 90.4 | 77.8444444 | 86.5333333 | 76.2 | 83.6333333 | 80.6222222 | 79.0333333 | 90.2333333 | 94.8444444 | 79.2703704 | 13.8049014 | 3.56441022 |
| 1 | 57.6666667 | 95.5 | 55.1666667 | 84 | 52.8333333 | 67.8 | 54.5 | 70.8333333 | 63.8333333 | 79.3333333 | 80.1666667 | 85.8333333 | 83.3333333 | 78.3333333 | 70.1666667 | 71.9533333 | 13.169444 | 3.40033581 |
| 2 | 46.8333333 | 57.3333333 | 46.1666667 | 83.8333333 | 51.1666667 | 70.3333333 | 62.5 | 57.5 | 59 | 65.5 | 56.1666667 | 94.5 | 58.6666667 | 73.6666667 | 71.5 | 63.6444444 | 13.3495239 | 3.44683225 |
| 3 | 58.1666667 | 52.3333333 | 45.5 | 80.3333333 | 44.3333333 | 65.8333333 | 61.6666667 | 63.5 | 59.6666667 | 62.3333333 | 58.3333333 | 93.6666667 | 66.1666667 | 69.5 | 60 | 62.7555556 | 12.3534049 | 3.18963544 |
| 4 | 44.5 | 67.3333333 | 46.5 | 81.3333333 | 43.8333333 | 66.1666667 | 61.6666667 | 69.5 | 57.6666667 | 58 | 59.8333333 | 88.5 | 72.8333333 | 69.5 | 49.8333333 | 62.4666667 | 13.1273071 | 3.38945611 |
| 5 | 49.3333333 | 79.5 | 42.6666667 | 71.1666667 | 46.5 | 68.6666667 | 62 | 65.3333333 | 62.1666667 | 63.3333333 | 53.8333333 | 88.5 | 79 | 74.3333333 | 72.6666667 | 65.2666667 | 13.0544951 | 3.37065613 |
| 10 | 65.0333333 | 73.1666667 | 77.3 | 83.9666667 | 50.9666667 | 68.4333333 | 65.6 | 64.5 | 62.6666667 | 49 | 65.1 | 66.3333333 | 80.7333333 | 59.1666667 | 69 | 66.7311111 | 9.64697886 | 2.49083923 |
| 15 | 67.8333333 | 96.1 | 66.6333333 | 90.4 | 48.9666667 | 66.7 | 57.7333333 | 49.8 | 51.0333333 | 77.4333333 | 86.6666667 | 78.0333333 | 87.4666667 | 72.8666667 | 70.2333333 | 71.1933333 | 15.0582161 | 3.88801469 |

Week 4

WT

| Time (min) | A | 156 | 158 | 161 | Average | SD | SEM |
|---|---|---|---|---|---|---|---|
| 0 | 60.5888889 | 62.2444444 | 72.2111111 | 43.0555556 | 59.525 | 12.1203308 | 6.06016538 |
| 1 | 46.1666667 | 47.8333333 | 59.8333333 | 59.6666667 | 53.375 | 7.39290826 | 3.69645413 |
| 2 | 46 | 45.6666667 | 45.3333333 | 48.6666667 | 46.4166667 | 1.52449141 | 0.76224571 |
| 3 | 49.8333333 | 47.1666667 | 54.3333333 | 49.6666667 | 50.25 | 2.98297639 | 1.4914882 |
| 4 | 45.5 | 42.1666667 | 57 | 48.3333333 | 48.25 | 6.3544968 | 3.1772484 |
| 5 | 45 | 46 | 41.8333333 | 48.3333333 | 45.2916667 | 2.69558968 | 1.34779484 |
| 10 | 60.9333333 | 60.5333333 | 49.4666667 | 62.1333333 | 58.2666667 | 5.90592922 | 2.95296461 |
| 15 | 80.3333333 | 78.9666667 | 70.8 | 63.0333333 | 73.2833333 | 8.02563024 | 4.01281512 |

Het

| Time (min) | B | D | E | H | 155 | 157 | 190 | 196 | 199 | 201 | 205 | 207 | 208 | 210 | Average | SD | SEM |
|---|---|---|---|---|---|---|---|---|---|---|---|---|---|---|---|---|---|
| 0 | 84.2888889 | 151.833333 | 84.3 | 64.4666667 | 84.3111111 | 86.2555556 | 86.5444444 | 90.1666667 | 92.0888889 | 73.2666667 | 70.9333333 | 87.5 | 42.9444444 | 96.3777778 | 85.3769841 | 23.548015 | 6.29347174 |
| 1 | 56.6666667 | 70.5 | 64.6666667 | 47 | 79.1666667 | 92.3333333 | 81.6666667 | 72.1666667 | 62 | 69.1666667 | 73.5 | 55.6666667 | 57.1666667 | 72.3333333 | 68.1428571 | 11.9721645 | 3.19969555 |
| 2 | 53.8333333 | 83 | 45 | 49 | 82.6666667 | 92.8333333 | 73.1666667 | 65 | 69.3333333 | 51.5 | 52.1666667 | 41.5 | 46.3333333 | 70.1666667 | 62.5357143 | 16.329039 | 4.36411925 |
| 3 | 45.8333333 | 81.6666667 | 49.5 | 55.6666667 | 83 | 92.8333333 | 74.8333333 | 71.8333333 | 73.3333333 | 53.5 | 64.1666667 | 40.6666667 | 51.5 | 75.6666667 | 65.2857143 | 15.9421267 | 4.26071258 |
| 4 | 49.3333333 | 73.6666667 | 39.6666667 | 38.1666667 | 75.5 | 90.3333333 | 64.1666667 | 68.5 | 54.1666667 | 57.5 | 60 | 43 | 43.6666667 | 79.8333333 | 59.8214286 | 16.2424821 | 4.34098593 |
| 5 | 48 | 61 | 45.3333333 | 46.1666667 | 74.1666667 | 86 | 79.5 | 73.3333333 | 56.3333333 | 57 | 69.1666667 | 41 | 44.1666667 | 83.3333333 | 61.75 | 15.7038226 | 4.19702312 |
| 10 | 71.2666667 | 88.9 | 49.2333333 | 41.7 | 77.2 | 91.0333333 | 72.2 | 71.3333333 | 54.4666667 | 68.1333333 | 72.3333333 | 49.8666667 | 66.2666667 | 64.9 | 67.0595238 | 14.3212601 | 3.82751777 |
| 15 | 56.5666667 | 117.833333 | 84.7333333 | 63.7666667 | 79 | 94.6666667 | 99.9666667 | 74.6551724 | 87.5 | 78.2333333 | 74.7 | 53.6 | 85.1 | 55.0333333 | 78.9539409 | 18.2466015 | 4.87660937 |

### Time of expiration

Week 1

WT

| Time (min) | Animal ID A | 156 | 158 | 161 | Average | SD | SEM |
|---|---|---|---|---|---|---|---|
| 0 | 112.044444 | 97.9111111 | 134.677778 | 132.588889 | 119.305556 | 17.5423122 | 8.77115608 |
| 1 | 103.166667 | 96.3333333 | 107.833333 | 111.833333 | 104.791667 | 6.65884958 | 3.32942479 |
| 2 | 91.3333333 | 110.333333 | 98 | 97.3333333 | 99.25 | 7.97391581 | 3.9869579 |
| 3 | 100.666667 | 94.8333333 | 112.833333 | 94.6666667 | 100.75 | 8.52501765 | 4.26250882 |
| 4 | 103.166667 | 108.666667 | 117.166667 | 112.5 | 110.375 | 5.93073445 | 2.96536722 |
| 5 | 115.5 | 106.333333 | 115.5 | 120.166667 | 114.375 | 5.79491316 | 2.89745658 |
| 10 | 109.6 | 110.533333 | 139.966667 | 109.9 | 117.5 | 14.9828297 | 7.49141484 |
| 15 | 141 | 134.533333 | 153.166667 | 135.533333 | 141.058333 | 8.55796683 | 4.27898342 |

Het

| Time (min) | Animal ID B | C | D | E | F | G | H | 155 | 157 | 159 | 160 | 162 | 190 | 196 | 199 | 201 | 205 | 207 | 208 | 210 | Average | SD | SEM |
|---|---|---|---|---|---|---|---|---|---|---|---|---|---|---|---|---|---|---|---|---|---|---|---|
| 0 | 137.422222 | 106.311111 | 122.144444 | 118.266667 | 167.833333 | 110.966667 | 112.455556 | 90.0666667 | 84.7666667 | 115.288889 | 109.666667 | 145.088889 | 119.8 | 138.955556 | 170.966667 | 85.7333333 | 108.255556 | 133.455556 | 142.022222 | 153.022222 | 123.624444 | 24.6915105 | 5.52118959 |
| 1 | 89 | 93.8333333 | 99.3333333 | 105 | 179 | 87.3333333 | 99.8333333 | 83.1666667 | 109.5 | 112.833333 | 92.6666667 | 123 | 99 | 99.1666667 | 105.666667 | 131.666667 | 105.833333 | 117.333333 | 115.666667 | 124.166667 | 108.65 | 21.0401941 | 4.70473042 |
| 2 | 103.166667 | 94.5 | 106.833333 | 109.666667 | 197.333333 | 100.5 | 90.6666667 | 104.166667 | 166 | 110.5 | 105.666667 | 127.5 | 88.8333333 | 103.666667 | 101.5 | 130.5 | 128.333333 | 120.166667 | 118 | 117.166667 | 116.233333 | 25.7946456 | 5.76785811 |
| 3 | 112.166667 | 89 | 104.166667 | 94.8333333 | 196.166667 | 94.1666667 | 92.6666667 | 118.833333 | 129.833333 | 102.666667 | 100.333333 | 117 | 89.8333333 | 111.5 | 123 | 134.833333 | 113.5 | 126 | 129.833333 | 113.333333 | 114.683333 | 23.8215773 | 5.32666661 |
| 4 | 116.166667 | 87.6666667 | 114.333333 | 96.3333333 | 215.833333 | 84.8333333 | 106.666667 | 115.666667 | 136.166667 | 115.166667 | 121.666667 | 112.5 | 87.6666667 | 119.666667 | 96 | 115.833333 | 119.833333 | 96.1666667 | 119.333333 | 109.666667 | 114.358333 | 27.4255118 | 6.13253088 |
| 5 | 119 | 91.5 | 110.5 | 117 | 259.5 | 89.3333333 | 120.666667 | 108 | 120.5 | 119 | 110.666667 | 119.833333 | 131.5 | 131.333333 | 104.833333 | 125.833333 | 125.166667 | 104.333333 | 99.6666667 | 128.833333 | 121.85 | 34.6151312 | 7.74017865 |
| 10 | 148.766667 | 101.633333 | 171.166667 | 176.1 | 252.2 | 95.7666667 | 130.933333 | 120.133333 | 164.533333 | 138.6 | 141.9 | 129.266667 | 119.4 | 148.466667 | 128.6 | 116.966667 | 127.566667 | 143.133333 | 168.033333 | 155.7 | 143.943333 | 33.7254448 | 7.54123871 |
| 15 | 135.133333 | 117.733333 | 152.8 | 163.366667 | 214.7 | 123.966667 | 139.2 | 118 | 173.466667 | 161.833333 | 157.766667 | 168.866667 | 143.566667 | 165.166667 | 226.9 | 178.566667 | 143.7 | 174.533333 | 151.033333 | 165.6 | 158.795 | 28.0039038 | 6.26186326 |

Week 2

WT

| Time (min) | Animal ID A | 156 | 158 | 161 | Average | SD | SEM |
|---|---|---|---|---|---|---|---|
| 0 | 136.277778 | 136.922222 | 115.777778 | 108.911111 | 124.472222 | 14.2842009 | 7.14210044 |
| 1 | 94.8333333 | 95.3333333 | 100 | 96.6666667 | 96.7083333 | 2.32687599 | 1.16343799 |
| 2 | 100.833333 | 106.166667 | 98.3333333 | 107.333333 | 103.166667 | 4.28822678 | 2.14411339 |
| 3 | 86.3333333 | 99.8333333 | 110 | 117.333333 | 103.375 | 13.4373547 | 6.71867733 |
| 4 | 100.333333 | 127.333333 | 110 | 111.833333 | 112.375 | 11.1756804 | 5.58784022 |
| 5 | 120 | 125.166667 | 105.833333 | 103.166667 | 113.541667 | 10.7068471 | 5.35342353 |
| 10 | 118.266667 | 128.2 | 149.666667 | 126.666667 | 130.7 | 13.3770949 | 6.68854743 |
| 15 | 153.266667 | 147.133333 | 166 | 147.566667 | 153.491667 | 8.79475328 | 4.39737664 |

Het

| Time (min) | Animal ID B | D | E | G | H | 155 | 157 | 160 | 162 | 190 | 196 | 199 | 201 | 205 | 207 | 208 | 210 | Average | SD | SEM |
|---|---|---|---|---|---|---|---|---|---|---|---|---|---|---|---|---|---|---|---|---|
| 0 | 159.477778 | 125.955556 | 178.222222 | 136.711111 | 123.588889 | 130.766667 | 150.166667 | 100.288889 | 183.455556 | 114.055556 | 156.488889 | 171.055556 | 148.666667 | 85.5777778 | 105.811111 | 88.2666667 | 93.6444444 | 132.482353 | 31.5746316 | 7.65797302 |
| 1 | 96.5 | 83.5 | 113.333333 | 102.333333 | 100.166667 | 100.5 | 114.333333 | 88.3333333 | 122.5 | 90.2 | 99.8 | 151.4 | 120.6 | 103.4 | 103 | 95 | 83.6 | 104.029412 | 16.8016198 | 4.07499136 |
| 2 | 94.8333333 | 82 | 107.166667 | 106.833333 | 103.5 | 103.5 | 107.166667 | 110.166667 | 204.333333 | 97.1666667 | 95.5 | 102.166667 | 127.833333 | 110.166667 | 142.333333 | 157.333333 | 115 | 115.705882 | 29.0852154 | 7.0542009 |
| 3 | 103.166667 | 90.5 | 126.666667 | 109.833333 | 96 | 125.833333 | 123 | 92.6666667 | 187 | 103.5 | 111.166667 | 109.833333 | 131.5 | 118.166667 | 150.666667 | 209 | 112.666667 | 123.598039 | 32.0555648 | 7.77461643 |
| 4 | 106.666667 | 83.3333333 | 103.333333 | 106.666667 | 126.166667 | 114.166667 | 128.833333 | 109.166667 | 157.166667 | 110.5 | 129.5 | 119 | 158.333333 | 132.333333 | 147.166667 | 217.5 | 147.166667 | 129.235294 | 30.5798169 | 7.41669502 |
| 5 | 141.666667 | 96.8333333 | 156.333333 | 96.5 | 141.333333 | 141.666667 | 139 | 125.833333 | 159.166667 | 103.333333 | 156.833333 | 82.5 | 149.666667 | 133.333333 | 146.5 | 167.333333 | 140.5 | 134.019608 | 24.8147219 | 6.0184541 |
| 10 | 153.133333 | 102.033333 | 204.966667 | 117.866667 | 150.633333 | 132.1 | 146.133333 | 165.166667 | 162.933333 | 129.733333 | 130.933333 | 109.233333 | 132.7 | 133.966667 | 141.6 | 147.2 | 145 | 141.490196 | 23.6675345 | 5.74022027 |
| 15 | 151.066667 | 150.333333 | 259.766667 | 142.166667 | 156.666667 | 158.466667 | 125.4 | 217 | 270.5 | 161.533333 | 174 | 139.633333 | 116.7 | 157.033333 | 157.766667 | 138 | 179.366667 | 167.964706 | 42.8638939 | 10.3960213 |

Week 3

WT

| Time (min) | Animal ID 83 | 86 | 87 | 88 | 81 | 82 | A | 156 | 158 | 161 | Average | SD | SEM |
|---|---|---|---|---|---|---|---|---|---|---|---|---|---|
| 0 | 124.933333 | 178.211111 | 135.155556 | 109.122222 | 164.314607 | 150.088889 | 117.188889 | 146.355556 | 113.633333 | 139.655556 | 137.865905 | 22.5277633 | 7.12390425 |
| 1 | 98.8333333 | 89.6666667 | 81.1666667 | 76.6666667 | 89 | 75.5 | 76.3333333 | 87 | 97.1666667 | 89.1666667 | 86.05 | 8.41297132 | 2.66041513 |
| 2 | 126.333333 | 84.5 | 80.8333333 | 91.8333333 | 106.833333 | 91.3333333 | 91.3333333 | 87.8333333 | 104.5 | 96.6666667 | 96.2 | 13.3318981 | 4.21591634 |
| 3 | 132.5 | 93.1666667 | 93 | 103.333333 | 140.333333 | 125.833333 | 109.333333 | 108.666667 | 92.1666667 | 109.5 | 110.783333 | 17.0235947 | 5.38333333 |
| 4 | 123.833333 | 96 | 94.6666667 | 93 | 134.5 | 107.333333 | 76.1666667 | 79.5 | 101 | 117.666667 | 102.366667 | 18.7108934 | 5.91690401 |
| 5 | 136.666667 | 106.166667 | 111.833333 | 84 | 104.833333 | 78.3333333 | 111 | 87.5 | 136.5 | 97.5 | 105.433333 | 19.9773328 | 6.31738733 |
| 10 | 178.1 | 119.3 | 131.366667 | 114.3 | 99.4 | 103.333333 | 109.933333 | 110.633333 | 126.933333 | 101.866667 | 119.516667 | 23.1025585 | 7.30567047 |
| 15 | 125.7 | 167.4 | 179.833333 | 190.933333 | 164 | 126.333333 | 134.733333 | 209.666667 | 156.833333 | 142.066667 | 159.75 | 28.2196073 | 8.92382337 |

Het

| Time (min) | Animal ID B | D | E | G | H | 155 | 157 | 190 | 196 | 199 | 201 | 205 | 207 | 208 | 210 | Average | SD | SEM |
|---|---|---|---|---|---|---|---|---|---|---|---|---|---|---|---|---|---|---|
| 0 | 146.266667 | 134.466667 | 166.522222 | 239.444444 | 128.677778 | 96.2 | 152.144444 | 135.255556 | 155.388889 | 170.355556 | 158.122222 | 147.888889 | 140.088889 | 145.144444 | 143.066667 | 150.602222 | 30.2109757 | 7.80044039 |
| 1 | 79.1666667 | 110.166667 | 78.5 | 182.166667 | 103.166667 | 71.8 | 85.3333333 | 101.833333 | 112.166667 | 123.833333 | 114.5 | 102.666667 | 105.166667 | 148.833333 | 89.5 | 107.253333 | 28.5865014 | 7.38100293 |
| 2 | 86.5 | 84.5 | 80 | 228.833333 | 102.5 | 82.6666667 | 112.5 | 122.333333 | 134.166667 | 136.5 | 116 | 124.833333 | 157 | 141.166667 | 94.5 | 120.266667 | 38.3288714 | 9.89647203 |
| 3 | 107.833333 | 91.6666667 | 76 | 224.166667 | 85.1666667 | 92.3333333 | 109.833333 | 125 | 147.333333 | 116.833333 | 117.666667 | 119.333333 | 175.166667 | 134.666667 | 80.6666667 | 120.244444 | 39.1082577 | 10.0977087 |
| 4 | 82.8333333 | 133.5 | 93.1666667 | 243.833333 | 86 | 100.166667 | 110.5 | 122.166667 | 129.833333 | 113.5 | 123 | 124.166667 | 164.166667 | 152 | 98.8333333 | 125.177778 | 40.0433033 | 10.3391364 |
| 5 | 103 | 142.666667 | 79.5 | 206.666667 | 92.5 | 104.333333 | 135 | 122.5 | 160.666667 | 132 | 106.166667 | 124.5 | 152.833333 | 153.333333 | 127.166667 | 129.522222 | 31.7094865 | 8.18735421 |
| 10 | 145.133333 | 155.2 | 176.666667 | 216.6 | 115.9 | 105.866667 | 148.266667 | 114.033333 | 157.466667 | 99.6666667 | 145.566667 | 127.533333 | 149.666667 | 134.566667 | 131.366667 | 141.566667 | 29.5645138 | 7.63352464 |
| 15 | 144.2 | 175.833333 | 150.533333 | 225.466667 | 109.166667 | 116.333333 | 124.233333 | 110 | 118.8 | 210.033333 | 183.166667 | 149.133333 | 152.2 | 177.033333 | 150.266667 | 153.093333 | 35.5885531 | 9.1889249 |

Week 4

WT

| Time (min) | Animal ID A | 156 | 158 | 161 | Average | SD | SEM |
|---|---|---|---|---|---|---|---|
| 0 | 118.611111 | 130.811111 | 179.288889 | 90.8222222 | 129.883333 | 36.9441158 | 18.4720579 |
| 1 | 75.5 | 86.8333333 | 106.5 | 98.1666667 | 91.75 | 13.5027432 | 6.7513716 |
| 2 | 95 | 87 | 90 | 105 | 94.25 | 7.88986692 | 3.94493346 |
| 3 | 101 | 88.6666667 | 139.333333 | 117.166667 | 111.541667 | 21.8968013 | 10.9484007 |
| 4 | 85.3333333 | 89 | 135 | 130.333333 | 109.916667 | 26.3809345 | 13.1904672 |
| 5 | 98.8333333 | 98.6666667 | 82.8333333 | 115.666667 | 99 | 13.407433 | 6.70371649 |
| 10 | 140.633333 | 141.166667 | 112.5 | 137.2 | 132.875 | 13.6965899 | 6.84829494 |
| 15 | 165.966667 | 178.066667 | 190.666667 | 150.2 | 171.225 | 17.2673825 | 8.63369127 |

Het

| Time (min) | Animal ID B | D | E | H | 155 | 157 | 190 | 196 | 199 | 201 | 205 | 207 | 208 | 210 | Average | SD | SEM |
|---|---|---|---|---|---|---|---|---|---|---|---|---|---|---|---|---|---|
| 0 | 137.344444 | 328.511111 | 153.055556 | 149.888889 | 136.433333 | 162.266667 | 147.022222 | 164.777778 | 177.855556 | 144.577778 | 140.2 | 158.677778 | 93.1555556 | 148.033333 | 160.128571 | 52.1632847 | 13.9412243 |
| 1 | 93.3333333 | 129.833333 | 86.3333333 | 88.1666667 | 79.6666667 | 101.333333 | 89.3333333 | 123.666667 | 113.833333 | 124.333333 | 93.8333333 | 129 | 92 | 84.1666667 | 102.059524 | 18.1196884 | 4.84269042 |
| 2 | 118.333333 | 161.666667 | 84 | 93.5 | 86.5 | 126.833333 | 104.166667 | 160.5 | 157.333333 | 110.333333 | 117 | 85 | 92.6666667 | 153 | 117.916667 | 29.4743659 | 7.87735565 |
| 3 | 92.6666667 | 160 | 106.833333 | 112.666667 | 102.833333 | 110.166667 | 118.833333 | 165.833333 | 171.833333 | 105.666667 | 126.166667 | 88.1666667 | 109.833333 | 151.166667 | 123.047619 | 27.6738042 | 7.39613528 |
| 4 | 109.333333 | 153.666667 | 77.6666667 | 64.5 | 101.833333 | 107.833333 | 125 | 146.5 | 109.666667 | 110.5 | 143.666667 | 89.1666667 | 98.8333333 | 137.666667 | 112.559524 | 26.3377639 | 7.03906349 |
| 5 | 106.5 | 125.333333 | 108.666667 | 111.5 | 103 | 114 | 131.666667 | 162.666667 | 119.166667 | 122.333333 | 166.833333 | 93.5 | 92 | 129.5 | 120.47619 | 22.3017244 | 5.96038656 |
| 10 | 154.266667 | 196.6 | 133.3 | 95.7333333 | 103.733333 | 125.466667 | 140.666667 | 169.633333 | 132.733333 | 150.433333 | 164.533333 | 114.933333 | 171.9 | 149.433333 | 143.097619 | 27.9845271 | 7.47917946 |
| 15 | 149.333333 | 276.733333 | 173.3 | 196.8 | 131.233333 | 135.533333 | 172.566667 | 186.310345 | 221 | 185.666667 | 163.9 | 124.4 | 161.4 | 139.866667 | 172.717406 | 40.5285069 | 10.8316991 |

### Peak inspiratory flow

Week 1

WT — Time (min) / Animal ID

| Time (min) | A | 156 | 158 | 161 | Average | SD | SEM |
|---|---|---|---|---|---|---|---|
| 0 | 4.04144444 | 4.27955556 | 2.87088889 | 3.89133333 | 3.77080556 | 0.62087275 | 0.31043638 |
| 1 | 3.92333333 | 4.26 | 2.565 | 3.055 | 3.45083333 | 0.77875791 | 0.38937896 |
| 2 | 4.80833333 | 4.18333333 | 2.77666667 | 3.80833333 | 3.89416667 | 0.85156555 | 0.42578278 |
| 3 | 4.40333333 | 4.23833333 | 2.80166667 | 3.735 | 3.79458333 | 0.72039903 | 0.36019952 |
| 4 | 4.085 | 3.935 | 2.725 | 3.20666667 | 3.48791667 | 0.63705976 | 0.31852988 |
| 5 | 3.56666667 | 3.53666667 | 3.005 | 3.26 | 3.34208333 | 0.26373168 | 0.13186584 |
| 10 | 3.93966667 | 3.697 | 2.776 | 3.251 | 3.41591667 | 0.51316428 | 0.25658214 |
| 15 | 2.4 | 2.60766667 | 1.84633333 | 1.98733333 | 2.21033333 | 0.35405848 | 0.17702924 |

Het — Time (min) / Animal ID

| Time (min) | B | C | D | E | F | G | H | 155 | 157 | 159 | 160 | 162 | 190 | 196 | 199 | 201 | 205 | 207 | 208 | 210 | Average | SD | SEM |
|---|---|---|---|---|---|---|---|---|---|---|---|---|---|---|---|---|---|---|---|---|---|---|---|
| 0 | 2.296 | 3.24622222 | 2.72955556 | 3.36688889 | 1.83255556 | 2.174 | 3.67844444 | 3.19633333 | 3.08533333 | 2.78166667 | 3.69166667 | 1.791 | 4.23466667 | 2.38422222 | 2.00144444 | 3.25344444 | 2.59533333 | 2.436 | 2.42877778 | 1.29866667 | 2.72511111 | 0.74195313 | 0.16590576 |
| 1 | 4.12666667 | 2.565 | 2.91666667 | 4.255 | 1.58 | 3.44166667 | 4.12166667 | 2.93333333 | 2.35833333 | 2.72166667 | 3.37333333 | 2.05833333 | 3.56833333 | 3.21666667 | 3.36833333 | 2.655 | 1.97666667 | 2.44 | 2.36833333 | 2.00666667 | 2.90258333 | 0.76815939 | 0.17176566 |
| 2 | 3.91166667 | 3.305 | 2.765 | 3.78833333 | 1.52333333 | 3.32833333 | 4.12666667 | 3.30666667 | 2.625 | 2.70666667 | 3.45333333 | 2.385 | 4.405 | 3.41166667 | 3.545 | 2.88 | 1.91166667 | 2.55833333 | 2.42 | 2.28 | 3.03183333 | 0.75156054 | 0.16805405 |
| 3 | 3.49666667 | 3.25166667 | 2.735 | 3.87333333 | 1.49166667 | 3.43333333 | 4.32166667 | 3.65666667 | 2.89 | 2.92833333 | 3.545 | 2.48166667 | 4.74333333 | 3.36833333 | 3.42833333 | 2.98666667 | 1.63333333 | 2.66166667 | 2.21666667 | 2.16166667 | 3.06525 | 0.8252202 | 0.18452485 |
| 4 | 3.495 | 3.32833333 | 2.75666667 | 4.025 | 1.42333333 | 3.34333333 | 3.43833333 | 3.42 | 2.78833333 | 2.90666667 | 3.32833333 | 2.42833333 | 4.50333333 | 3.53666667 | 3.55333333 | 3.31 | 1.53833333 | 2.91666667 | 2.28666667 | 2.44 | 3.03833333 | 0.75428057 | 0.16866226 |
| 5 | 3.06333333 | 3.27833333 | 2.83 | 3.77666667 | 1.37833333 | 3.34166667 | 3.52833333 | 3.14333333 | 2.69 | 2.63666667 | 3.21666667 | 2.50833333 | 3.565 | 2.55333333 | 3.475 | 3.06 | 1.65666667 | 2.95166667 | 2.58333333 | 2.27833333 | 2.87575 | 0.61631746 | 0.13781277 |
| 10 | 2.58233333 | 2.81266667 | 2.40566667 | 2.229 | 1.35333333 | 3.14266667 | 3.41633333 | 3.04366667 | 2.34566667 | 2.52933333 | 3.02333333 | 2.7 | 3.97633333 | 2.45633333 | 3.47266667 | 3.092 | 1.98766667 | 2.57533333 | 1.65066667 | 1.97266667 | 2.63838333 | 0.63871424 | 0.14282084 |
| 15 | 2.11233333 | 2.08766667 | 2.61266667 | 2.104 | 1.43 | 2.54366667 | 2.906 | 2.12466667 | 1.905 | 1.48966667 | 2.53933333 | 1.665 | 3.11333333 | 2.47966667 | 2.77066667 | 2.77866667 | 2.53 | 1.43733333 | 1.092 | 1.34466667 | 2.15331667 | 0.58880084 | 0.13165987 |

Week 2

WT — Time (min) / Animal ID

| Time (min) | A | 156 | 158 | 161 | Average | SD | SEM |
|---|---|---|---|---|---|---|---|
| 0 | 4.56344444 | 3.01333333 | 3.70611111 | 4.56088889 | 3.96094444 | 0.74963245 | 0.37481623 |
| 1 | 5 | 4.54833333 | 3.255 | 4.02333333 | 4.20666667 | 0.74953195 | 0.37476598 |
| 2 | 5.13 | 4.47166667 | 3.78333333 | 3.86833333 | 4.31333333 | 0.62475032 | 0.31237516 |
| 3 | 5.39 | 4.91 | 3.71 | 3.48666667 | 4.37416667 | 0.92155816 | 0.46077908 |
| 4 | 4.955 | 4.45666667 | 3.81166667 | 3.81166667 | 4.25875 | 0.554888 | 0.277444 |
| 5 | 4.31833333 | 4.60333333 | 3.42666667 | 3.50666667 | 3.96375 | 0.58656624 | 0.29328312 |
| 10 | 4.48433333 | 4.40733333 | 3.186 | 3.69433333 | 3.943 | 0.61739543 | 0.30869771 |
| 15 | 3.98966667 | 3.41 | 2.49266667 | 2.89166667 | 3.196 | 0.64884496 | 0.32442248 |

Het — Time (min) / Animal ID

| Time (min) | B | D | E | G | H | 155 | 157 | 160 | 162 | 190 | 196 | 199 | 201 | 205 | 207 | 208 | 210 | Average | SD | SEM |
|---|---|---|---|---|---|---|---|---|---|---|---|---|---|---|---|---|---|---|---|---|
| 0 | 2.40922222 | 3.23811111 | 1.62322222 | 1.39233333 | 4.44044444 | 2.24366667 | 2.40177778 | 3.23944444 | 3.63622222 | 3.44911111 | 2.34511111 | 2.34411111 | 3.12122222 | 3.88 | 3.10322222 | 3.71388889 | 4.26 | 2.99065359 | 0.87710459 | 0.21272911 |
| 1 | 4.99333333 | 4.25 | 3.06166667 | 3.125 | 4.42333333 | 3.14833333 | 2.54833333 | 3.49 | 2.505 | 2.242 | 4.38 | 2.768 | 2.692 | 1.646 | 1.906 | 2.21 | 1.824 | 3.01252941 | 0.9985779 | 0.24219071 |
| 2 | 5.49666667 | 5.095 | 3.91666667 | 3.245 | 4.59333333 | 3.79333333 | 3.41 | 3.39833333 | 2.11666667 | 4.21833333 | 5.23666667 | 3.80333333 | 2.635 | 1.555 | 2.72833333 | 2.25333333 | 1.63166667 | 3.47803922 | 1.21886734 | 0.29561875 |
| 3 | 5.27333333 | 4.91666667 | 3.45333333 | 3.32333333 | 4.60833333 | 3.76166667 | 2.985 | 3.43333333 | 2.37333333 | 3.74333333 | 4.79333333 | 4.05333333 | 3.02833333 | 2.335 | 2.51666667 | 1.67666667 | 2.755 | 3.47235294 | 1.01465136 | 0.2460891 |
| 4 | 5.06166667 | 4.96833333 | 3.84 | 3.01166667 | 4.23166667 | 3.85333333 | 2.96833333 | 3.29 | 2.65333333 | 2.92666667 | 5.11 | 4.11666667 | 2.74666667 | 1.88833333 | 1.95833333 | 1.64 | 2.27333333 | 3.32578431 | 1.11646253 | 0.27078194 |
| 5 | 4.81333333 | 4.43333333 | 3.44 | 3.425 | 3.33666667 | 3.755 | 2.89 | 3.18166667 | 2.505 | 4.02666667 | 4.50833333 | 4.42833333 | 2.82 | 2.165 | 1.82166667 | 1.385 | 2.655 | 3.27 | 0.98949605 | 0.23998804 |
| 10 | 4.67533333 | 3.99666667 | 2.99666667 | 2.91466667 | 3.23733333 | 3.57933333 | 3.088 | 2.85933333 | 2.602 | 3.00533333 | 4.76133333 | 3.63966667 | 3.25666667 | 2.56266667 | 2.42966667 | 1.93233333 | 2.872 | 3.20052941 | 0.74799211 | 0.18141473 |
| 15 | 4.78733333 | 2.728 | 1.495 | 2.62133333 | 3.49866667 | 3.38633333 | 3.79133333 | 2.42733333 | 2.04466667 | 2.90166667 | 4.15233333 | 3.44566667 | 4.16833333 | 2.103 | 2.67833333 | 2.50566667 | 1.87566667 | 2.97709804 | 0.91003135 | 0.22071502 |

Week 3

WT — Time (min) / Animal ID

| Time (min) | 83 | 86 | 87 | 88 | 81 | 82 | A | 156 | 158 | 161 | Average | SD | SEM |
|---|---|---|---|---|---|---|---|---|---|---|---|---|---|
| 0 | 5.976 | 3.45477778 | 4.92333333 | 6.06577778 | 3.94752809 | 2.76955556 | 5.93266667 | 4.64511111 | 4.01211111 | 3.66077778 | 4.53876392 | 1.16440228 | 0.36821633 |
| 1 | 5.86166667 | 5.72166667 | 7.935 | 7.98666667 | 7.69166667 | 6.835 | 6.31666667 | 7.348 | 3.52333333 | 7.29833333 | 6.6518 | 1.36768794 | 0.4325009 |
| 2 | 5.63333333 | 6.19833333 | 7.49166667 | 6.68 | 7.51333333 | 7.20833333 | 6.64 | 7.33 | 3.67833333 | 4.23166667 | 6.2605 | 1.35926997 | 0.42983891 |
| 3 | 5.685 | 6.42333333 | 6.51 | 6.605 | 6.92666667 | 6.185 | 5.97833333 | 6.92833333 | 4.03833333 | 4.02166667 | 5.93016667 | 1.07356837 | 0.33949213 |
| 4 | 5.71833333 | 6.075 | 6.69833333 | 6.655 | 6.445 | 5.72833333 | 6.135 | 6.72 | 4.215 | 4.17166667 | 5.85616667 | 0.95057035 | 0.30059674 |
| 5 | 4.96 | 5.585 | 6.55666667 | 7.13333333 | 6.635 | 5.44166667 | 5.48 | 6.08 | 3.63 | 4.64333333 | 5.6145 | 1.04409718 | 0.33017252 |
| 10 | 5.25733333 | 5.833 | 6.03866667 | 5.75266667 | 6.382 | 5.12866667 | 5.45933333 | 5.65766667 | 3.32633333 | 4.35233333 | 5.3188 | 0.89316755 | 0.28244438 |
| 15 | 5.46433333 | 4.44966667 | 4.20333333 | 4.17433333 | 5.71633333 | 4.29066667 | 4.112 | 4.09433333 | 2.41933333 | 4.61133333 | 4.35356667 | 0.88769868 | 0.28071497 |

Het — Time (min) / Animal ID

| Time (min) | B | D | E | G | H | 155 | 157 | 190 | 196 | 199 | 201 | 205 | 207 | 208 | 210 | Average | SD | SEM |
|---|---|---|---|---|---|---|---|---|---|---|---|---|---|---|---|---|---|---|
| 0 | 2.71444444 | 3.81966667 | 2.54222222 | 1.581 | 4.51311111 | 5.02433333 | 2.32055556 | 2.81455556 | 2.81488889 | 3.24144444 | 2.81477778 | 1.84722222 | 2.16111111 | 1.71622222 | 1.67477778 | 2.77335556 | 1.02306492 | 0.26415423 |
| 1 | 4.68833333 | 3.08333333 | 3.66166667 | 1.36666667 | 5.26166667 | 2.618 | 4.01333333 | 3.10833333 | 4.00666667 | 2.91 | 2.69 | 2.02166667 | 2.22666667 | 1.975 | 1.84333333 | 3.03164444 | 1.11191198 | 0.28709444 |
| 2 | 6.07666667 | 4.47833333 | 5.02166667 | 1.43666667 | 5.57833333 | 2.22166667 | 3.76 | 3.97 | 4.18666667 | 3.685 | 3.66666667 | 1.45666667 | 2.66833333 | 2.30833333 | 1.57333333 | 3.47255556 | 1.48642613 | 0.38379358 |
| 3 | 4.58833333 | 4.96 | 5.22666667 | 1.495 | 5.55666667 | 2.975 | 3.98166667 | 3.78 | 4.735 | 3.65833333 | 3.89833333 | 1.36833333 | 2.19833333 | 2.68166667 | 2.58833333 | 3.57944444 | 1.32251964 | 0.3414731 |
| 4 | 6.08166667 | 3.63833333 | 4.915 | 1.41 | 5.24333333 | 2.93833333 | 4.22833333 | 3.43666667 | 4.90666667 | 4.35833333 | 4.12333333 | 1.575 | 2.14833333 | 2.76833333 | 3.78333333 | 3.70366667 | 1.35155087 | 0.34896893 |
| 5 | 5.94666667 | 3.09333333 | 5.195 | 1.50833333 | 5.19 | 2.81 | 4.19833333 | 3.77 | 4.815 | 3.91333333 | 4.35666667 | 1.58833333 | 1.98333333 | 2.55166667 | 2.02666667 | 3.52977778 | 1.42752351 | 0.36858498 |
| 10 | 4.34566667 | 3.7 | 2.84133333 | 1.46366667 | 4.69033333 | 2.87266667 | 3.838 | 3.647 | 4.82133333 | 4.076 | 4.027 | 2.49966667 | 2.08466667 | 3.34666667 | 2.696 | 3.39666667 | 0.96920304 | 0.25024715 |
| 15 | 3.83133333 | 2.48733333 | 3.374 | 1.417 | 4.83633333 | 3.26166667 | 4.295 | 4.20933333 | 5.11533333 | 3.28566667 | 2.98666667 | 1.99933333 | 1.82866667 | 2.75766667 | 2.94833333 | 3.24224444 | 1.07871425 | 0.27852282 |

Week 4

WT — Time (min) / Animal ID

| Time (min) | A | 156 | 158 | 161 | Average | SD | SEM |
|---|---|---|---|---|---|---|---|
| 0 | 5.46811111 | 5.94177778 | 3.26466667 | 5.62077778 | 5.07383333 | 1.22215858 | 0.61107929 |
| 1 | 6.73 | 6.30166667 | 4.13833333 | 3.805 | 5.24375 | 1.48549367 | 0.74274683 |
| 2 | 6.15333333 | 5.92 | 4.36166667 | 3.98666667 | 5.10541667 | 1.09032732 | 0.54516366 |
| 3 | 5.98333333 | 5.89166667 | 3.82166667 | 4.375 | 5.01791667 | 1.08625109 | 0.54312555 |
| 4 | 6.74 | 6.14 | 4.23 | 4.27333333 | 5.34583333 | 1.28708219 | 0.64354109 |
| 5 | 6.025 | 5.92666667 | 4.81 | 4.21833333 | 5.245 | 0.87869959 | 0.43934979 |
| 10 | 5.10633333 | 5.72466667 | 4.356 | 3.926 | 4.77825 | 0.79748946 | 0.39874473 |
| 15 | 3.93933333 | 4.85133333 | 3.49966667 | 4.02366667 | 4.0785 | 0.5641198 | 0.2820599 |

Het — Time (min) / Animal ID

| Time (min) | B | D | E | H | 155 | 157 | 190 | 196 | 199 | 201 | 205 | 207 | 208 | 210 | Average | SD | SEM |
|---|---|---|---|---|---|---|---|---|---|---|---|---|---|---|---|---|---|
| 0 | 3.10088889 | 1.71622222 | 2.17033333 | 4.09111111 | 2.72744444 | 3.27466667 | 2.76255556 | 3.018 | 2.83622222 | 3.51611111 | 3.05122222 | 1.87255556 | 4.27177778 | 1.854 | 2.87593651 | 0.78572676 | 0.20999431 |
| 1 | 5.46666667 | 3.48666667 | 3.14666667 | 5.915 | 3.13666667 | 3.01 | 2.97333333 | 3.79166667 | 3.93833333 | 3.49666667 | 2.12666667 | 3.77833333 | 3.77 | 2.36833333 | 3.60035714 | 1.03507012 | 0.27663413 |
| 2 | 4.845 | 3.31666667 | 5.29833333 | 6.065 | 2.27666667 | 2.295 | 2.99833333 | 3.85666667 | 3.91833333 | 4.41 | 3.68 | 4.58 | 4.84166667 | 2.305 | 3.90619048 | 1.18007305 | 0.31538779 |
| 3 | 5.13833333 | 3.48666667 | 4.84166667 | 5.565 | 2.53 | 2.37833333 | 3.10166667 | 3.955 | 3.895 | 4.63833333 | 3.14666667 | 4.44166667 | 4.47166667 | 2.19 | 3.84142857 | 1.06716572 | 0.28521204 |
| 4 | 5.18166667 | 3.43166667 | 5.62333333 | 6.09666667 | 2.87166667 | 2.47333333 | 4.11333333 | 4.18333333 | 4.42333333 | 4.71833333 | 3.52 | 4.265 | 4.57166667 | 2.175 | 4.1177381 | 1.1388273 | 0.3043644 |
| 5 | 5.47833333 | 3.885 | 4.40166667 | 5.03833333 | 2.90666667 | 2.815 | 3.125 | 4.035 | 4.74 | 4.57333333 | 3.00833333 | 4.62166667 | 4.675 | 2.13666667 | 3.96 | 0.99830776 | 0.26680897 |
| 10 | 4.33533333 | 3.062 | 4.41933333 | 5.56366667 | 2.81133333 | 2.69733333 | 3.74566667 | 4.13833333 | 4.42866667 | 4.28633333 | 2.899 | 3.84633333 | 3.177 | 3.37033333 | 3.77004762 | 0.8153314 | 0.21790648 |
| 15 | 5.11933333 | 2.448 | 2.28333333 | 4.46766667 | 3.09466667 | 2.87533333 | 2.60233333 | 3.97551724 | 3.34766667 | 3.757 | 2.83166667 | 3.56166667 | 2.15233333 | 3.91 | 3.3161798 | 0.86748978 | 0.23184639 |

### Peak expiratory flow

Week 1

WT

| Time (min) | A | 156 | 158 | 161 | Average | SD | SE |
|---|---|---|---|---|---|---|---|
| 0 | 2.17588889 | 2.34844444 | 1.26433333 | 1.63944444 | 1.85702778 | 0.49725646 | 0.24862823 |
| 1 | 2.88666667 | 2.60833333 | 1.37333333 | 1.52166667 | 2.0975 | 0.76151956 | 0.38075978 |
| 2 | 2.84666667 | 2.31833333 | 1.44333333 | 2.025 | 2.15833333 | 0.58549283 | 0.29274641 |
| 3 | 2.51166667 | 2.26833333 | 1.45 | 1.94 | 2.0425 | 0.45923144 | 0.22961572 |
| 4 | 2.35666667 | 2.22 | 1.435 | 1.75 | 1.94041667 | 0.42549399 | 0.212747 |
| 5 | 2.29833333 | 2.20833333 | 1.42833333 | 1.735 | 1.9175 | 0.40914885 | 0.20457442 |
| 10 | 2.23933333 | 2.054 | 1.18166667 | 1.645 | 1.78 | 0.4698549 | 0.23492745 |
| 15 | 1.46466667 | 1.551 | 0.87333333 | 1.06533333 | 1.23858333 | 0.32256316 | 0.16128158 |

Het

| Time (min) | B | C | D | E | F | G | H | 155 | 157 | 159 | 160 | 162 | 190 | 196 | 199 | 201 | 205 | 207 | 208 | 210 | Average | SD | SE |
|---|---|---|---|---|---|---|---|---|---|---|---|---|---|---|---|---|---|---|---|---|---|---|---|
| 0 | 1.41433333 | 1.88177778 | 1.45744444 | 1.81233333 | 1.10022222 | 1.21677778 | 1.85311111 | 1.84488889 | 1.90333333 | 1.42744444 | 2.06666667 | 0.90011111 | 2.35944444 | 1.42111111 | 1.22311111 | 1.84344444 | 1.29555556 | 1.15233333 | 1.15755556 | 0.76522222 | 1.50481111 | 0.41951802 | 0.09380708 |
| 1 | 2.495 | 1.86 | 1.76333333 | 2.32833333 | 1.01166667 | 1.75166667 | 2.17833333 | 1.97 | 1.57833333 | 1.59166667 | 2.11 | 1.12833333 | 2.34166667 | 2.07833333 | 1.88 | 1.60833333 | 1.17166667 | 1.43166667 | 1.26333333 | 1.04666667 | 1.72941667 | 0.45323892 | 0.1013473 |
| 2 | 2.26666667 | 2.20833333 | 1.74166667 | 2.11166667 | 0.94166667 | 1.57833333 | 2.315 | 1.79666667 | 1.25 | 1.45333333 | 2.015 | 1.22833333 | 2.47833333 | 2.19 | 1.99 | 1.53 | 1.01166667 | 1.525 | 1.34 | 1.29166667 | 1.71316667 | 0.46279468 | 0.10348404 |
| 3 | 1.99666667 | 2.10666667 | 1.69 | 2.03 | 0.915 | 1.635 | 2.18666667 | 1.74166667 | 1.41833333 | 1.58666667 | 1.93166667 | 1.22666667 | 2.73666667 | 2.00166667 | 1.95666667 | 1.63333333 | 0.96166667 | 1.47166667 | 1.24666667 | 1.275 | 1.68741667 | 0.44765813 | 0.1000994 |
| 4 | 2.01666667 | 2.085 | 1.58333333 | 2.26 | 0.89666667 | 1.62833333 | 1.76333333 | 1.67 | 1.38166667 | 1.565 | 1.70833333 | 1.34666667 | 2.53 | 1.93 | 1.92 | 1.75666667 | 0.88 | 1.47833333 | 1.26 | 1.375 | 1.65175 | 0.41315166 | 0.09238352 |
| 5 | 1.855 | 2.06 | 1.555 | 1.88166667 | 0.89666667 | 1.74833333 | 1.61666667 | 1.68166667 | 1.40333333 | 1.49333333 | 1.81666667 | 1.19833333 | 1.81333333 | 1.535 | 1.78166667 | 1.52 | 0.90166667 | 1.46833333 | 1.29833333 | 1.16 | 1.53425 | 0.31889909 | 0.071308 |
| 10 | 1.527 | 1.76266667 | 1.21633333 | 1.322 | 0.868 | 1.49066667 | 1.61133333 | 1.578 | 1.26333333 | 1.24366667 | 1.692 | 1.25566667 | 2.02566667 | 1.49066667 | 1.64833333 | 1.58166667 | 1.02433333 | 1.14166667 | 0.80266667 | 1.044 | 1.37948333 | 0.31469411 | 0.07036774 |
| 15 | 1.379 | 1.43266667 | 1.32366667 | 1.31633333 | 0.934 | 1.236 | 1.375 | 1.371 | 1.10033333 | 0.73733333 | 1.53266667 | 0.809 | 1.698 | 1.42 | 1.299 | 1.32133333 | 1.05033333 | 0.73733333 | 0.69433333 | 0.80266667 | 1.1785 | 0.29878496 | 0.06681035 |

Week 2

WT

| Time (min) | A | 156 | 158 | 161 | Average | SD | SE |
|---|---|---|---|---|---|---|---|
| 0 | 2.23511111 | 1.67677778 | 1.68877778 | 2.22633333 | 1.95675 | 0.3164141 | 0.15820705 |
| 1 | 3.02833333 | 3.08833333 | 1.81666667 | 2.395 | 2.58208333 | 0.59896883 | 0.29948441 |
| 2 | 2.805 | 2.52833333 | 1.94333333 | 2.04833333 | 2.33125 | 0.40571011 | 0.20285506 |
| 3 | 2.765 | 2.44333333 | 1.83833333 | 2.06 | 2.27666667 | 0.41041758 | 0.20520879 |
| 4 | 2.52666667 | 2.23333333 | 1.87166667 | 1.99666667 | 2.15708333 | 0.28844531 | 0.14422265 |
| 5 | 2.23 | 2.24 | 1.87833333 | 1.97166667 | 2.08 | 0.18303511 | 0.09151756 |
| 10 | 2.17566667 | 2.24033333 | 1.31933333 | 1.66933333 | 1.85116667 | 0.43690617 | 0.21845308 |
| 15 | 1.899 | 1.73633333 | 1.03066667 | 1.293 | 1.48975 | 0.39906431 | 0.19953216 |

Het

| Time (min) | B | D | E | G | H | 155 | 157 | 160 | 162 | 190 | 196 | 199 | 201 | 205 | 207 | 208 | 210 | Average | SD | SE |
|---|---|---|---|---|---|---|---|---|---|---|---|---|---|---|---|---|---|---|---|---|
| 0 | 1.42111111 | 1.81033333 | 0.75077778 | 0.91677778 | 2.08044444 | 1.46233333 | 1.41955556 | 1.74844444 | 1.63555556 | 1.93522222 | 1.47944444 | 1.368 | 1.71 | 1.98222222 | 1.46288889 | 1.97055556 | 2.06277778 | 1.60096732 | 0.37741184 | 0.09153582 |
| 1 | 2.82 | 2.77333333 | 1.45166667 | 1.45666667 | 2.35333333 | 2.22833333 | 1.685 | 2.055 | 1.71333333 | 1.96 | 2.604 | 1.556 | 2.184 | 1.626 | 1.658 | 1.8 | 1.7 | 1.97792157 | 0.44747169 | 0.10852783 |
| 2 | 3.02 | 2.89833333 | 1.7 | 1.61333333 | 2.32 | 2.12166667 | 1.89333333 | 1.925 | 1.20333333 | 2.38666667 | 2.81 | 1.97833333 | 1.73 | 1.20333333 | 1.32333333 | 1.195 | 1.065 | 1.90509804 | 0.62231799 | 0.15093428 |
| 3 | 2.88333333 | 2.815 | 1.54833333 | 1.63833333 | 2.42 | 1.78833333 | 1.70666667 | 1.87 | 1.23833333 | 2.14 | 2.63 | 2.09 | 1.715 | 1.16333333 | 1.19166667 | 0.95166667 | 1.41833333 | 1.83578431 | 0.58745925 | 0.1424798 |
| 4 | 2.70833333 | 2.72166667 | 1.64 | 1.60166667 | 1.93166667 | 1.86166667 | 1.69166667 | 1.805 | 1.315 | 1.74333333 | 2.57833333 | 2.15 | 1.61 | 1.005 | 1.04166667 | 0.84666667 | 1.085 | 1.72568627 | 0.57600427 | 0.13970156 |
| 5 | 2.27666667 | 2.49166667 | 1.335 | 1.7 | 1.65333333 | 1.69 | 1.605 | 1.56333333 | 1.255 | 2.18166667 | 2.14833333 | 2.31666667 | 1.62333333 | 1.055 | 0.98333333 | 0.77666667 | 1.20666667 | 1.63892157 | 0.50632319 | 0.12280141 |
| 10 | 2.14666667 | 2.17666667 | 1.17666667 | 1.43166667 | 1.54366667 | 1.73533333 | 1.56566667 | 1.353 | 1.26866667 | 1.742 | 2.262 | 1.88566667 | 1.81733333 | 1.18166667 | 1.17933333 | 0.97166667 | 1.281 | 1.57168627 | 0.3916743 | 0.09499497 |
| 15 | 2.04333333 | 1.62433333 | 0.57866667 | 1.25766667 | 1.51633333 | 1.52633333 | 1.93466667 | 1.15233333 | 0.907 | 1.60933333 | 1.85666667 | 1.72266667 | 2.179 | 1.009 | 1.22633333 | 1.17233333 | 1.05333333 | 1.4334902 | 0.43861314 | 0.10637931 |

Week 3

WT

| Time (min) | 83 | 86 | 87 | 88 | 81 | 82 | A | 156 | 158 | 161 | Average | SD | SE |
|---|---|---|---|---|---|---|---|---|---|---|---|---|---|
| 0 | 2.83333333 | 1.809 | 2.53988889 | 2.97133333 | 1.95134831 | 1.58122222 | 2.78744444 | 2.16666667 | 2.02722222 | 1.97388889 | 2.26413483 | 0.48249732 | 0.15257905 |
| 1 | 3.425 | 3.61 | 4.96666667 | 5.08833333 | 4.50666667 | 4.51833333 | 3.875 | 4.064 | 2.09666667 | 2.98 | 3.91306667 | 0.92701697 | 0.29314851 |
| 2 | 3.02 | 3.47333333 | 3.885 | 3.60333333 | 3.65333333 | 4.265 | 3.525 | 3.905 | 1.995 | 2.265 | 3.359 | 0.72685319 | 0.22985116 |
| 3 | 2.77333333 | 3.43166667 | 3.27 | 3.49166667 | 3.03166667 | 3.275 | 2.94333333 | 3.42666667 | 1.955 | 2.10833333 | 2.97066667 | 0.54698399 | 0.17297152 |
| 4 | 2.495 | 3.15333333 | 3.25 | 3.53833333 | 2.83 | 3.03833333 | 3.23166667 | 3.59833333 | 2.015 | 2.065 | 2.9215 | 0.56329468 | 0.17812942 |
| 5 | 2.38666667 | 2.83 | 2.92833333 | 3.68333333 | 2.98833333 | 2.87333333 | 2.63833333 | 2.98166667 | 1.76166667 | 2.49333333 | 2.7565 | 0.49684961 | 0.15711764 |
| 10 | 2.23433333 | 2.828 | 2.77666667 | 2.798 | 3.04066667 | 2.742 | 2.57666667 | 2.74133333 | 1.56633333 | 2.22766667 | 2.55316667 | 0.43155367 | 0.13646925 |
| 15 | 2.544 | 2.04966667 | 2.15333333 | 1.99566667 | 2.461 | 2.34533333 | 2.21033333 | 1.684 | 1.088 | 2.09133333 | 2.06226667 | 0.42152096 | 0.13329663 |

Het

| Time (min) | B | D | E | G | H | 155 | 157 | 190 | 196 | 199 | 201 | 205 | 207 | 208 | 210 | Average | SD | SE |
|---|---|---|---|---|---|---|---|---|---|---|---|---|---|---|---|---|---|---|
| 0 | 1.60866667 | 1.92444444 | 1.29266667 | 0.91666667 | 2.12477778 | 2.55355556 | 1.38666667 | 1.60022222 | 1.58866667 | 1.58444444 | 1.55444444 | 0.94477778 | 1.14733333 | 1.08577778 | 1.05188889 | 1.491 | 0.45671286 | 0.11792275 |
| 1 | 3.02833333 | 2.59333333 | 2.52333333 | 0.73166667 | 2.82 | 2.378 | 2.44666667 | 2.17166667 | 2.41166667 | 1.77833333 | 1.96833333 | 1.755 | 1.98666667 | 1.245 | 1.5 | 2.0892 | 0.61625668 | 0.15911679 |
| 2 | 3.33666667 | 2.77833333 | 2.94166667 | 0.755 | 2.76 | 1.82333333 | 2.14666667 | 1.975 | 2.18666667 | 1.81666667 | 1.98 | 1.19666667 | 1.51666667 | 1.25333333 | 1.20666667 | 1.97822222 | 0.73619557 | 0.19008488 |
| 3 | 2.49 | 2.75833333 | 3.00166667 | 0.82 | 2.89166667 | 1.965 | 2.19166667 | 1.91333333 | 2.21333333 | 1.93166667 | 2.105 | 1.16 | 1.21 | 1.33166667 | 1.74333333 | 1.98177778 | 0.65191582 | 0.16832394 |
| 4 | 3.34333333 | 1.87166667 | 2.49833333 | 0.72666667 | 2.69833333 | 1.825 | 2.24166667 | 1.89 | 2.40666667 | 2.21333333 | 2.14 | 1.185 | 1.125 | 1.25 | 1.99166667 | 1.96044444 | 0.68028165 | 0.17564797 |
| 5 | 2.89 | 1.72333333 | 2.67833333 | 0.75666667 | 2.59666667 | 1.70833333 | 2.155 | 1.94 | 2.24 | 2.03666667 | 2.28333333 | 1.16166667 | 1.03166667 | 1.195 | 1.20333333 | 1.84 | 0.65583976 | 0.1693371 |
| 10 | 2.10733333 | 1.846 | 1.323 | 0.816 | 2.33266667 | 1.72733333 | 1.934 | 2.04966667 | 2.24033333 | 2.09766667 | 2.024 | 1.29166667 | 1.10533333 | 1.49333333 | 1.35266667 | 1.71606667 | 0.45699151 | 0.1179947 |
| 15 | 1.931 | 1.44733333 | 1.54433333 | 0.86666667 | 2.345 | 1.78033333 | 2.25733333 | 1.98533333 | 2.5 | 1.465 | 1.51333333 | 1.02666667 | 0.99066667 | 1.20033333 | 1.41033333 | 1.61757778 | 0.50488084 | 0.13035967 |

Week 4

WT

| Time (min) | A | 156 | 158 | 161 | Average | SD | SE |
|---|---|---|---|---|---|---|---|
| 0 | 2.846 | 2.98466667 | 1.43566667 | 3.10433333 | 2.59266667 | 0.77852287 | 0.38926143 |
| 1 | 3.92333333 | 3.615 | 2.49333333 | 2.54666667 | 3.14458333 | 0.732433 | 0.3662165 |
| 2 | 3.14 | 3.16666667 | 2.12666667 | 2.16166667 | 2.64875 | 0.5829195 | 0.29145975 |
| 3 | 3.095 | 3.06 | 1.96166667 | 2.27 | 2.59666667 | 0.56948807 | 0.28474404 |
| 4 | 3.71666667 | 3.11166667 | 2.42333333 | 2.02833333 | 2.82 | 0.74680678 | 0.37340339 |
| 5 | 2.92 | 2.915 | 2.425 | 2.06 | 2.58 | 0.41723295 | 0.20861647 |
| 10 | 2.45666667 | 2.69766667 | 2.125 | 1.78966667 | 2.26725 | 0.39558315 | 0.19779157 |
| 15 | 1.89266667 | 2.15833333 | 1.487 | 1.82 | 1.8395 | 0.27635598 | 0.13817799 |

Het

| Time (min) | B | D | E | H | 155 | 157 | 190 | 196 | 199 | 201 | 205 | 207 | 208 | 210 | Average | SD | SE |
|---|---|---|---|---|---|---|---|---|---|---|---|---|---|---|---|---|---|
| 0 | 1.90522222 | 1.06366667 | 1.18822222 | 1.88211111 | 1.66577778 | 1.75088889 | 1.60744444 | 1.63655556 | 1.49233333 | 1.81644444 | 1.46866667 | 0.96211111 | 2.05888889 | 1.17022222 | 1.54775397 | 0.33946921 | 0.09072696 |
| 1 | 3.35833333 | 2.01333333 | 2.25333333 | 3.23166667 | 2.94666667 | 2.615 | 2.59833333 | 2.43833333 | 2.28166667 | 2.16166667 | 1.80666667 | 1.67833333 | 2.25833333 | 2.08333333 | 2.40892857 | 0.49844785 | 0.13321579 |
| 2 | 2.71166667 | 1.86166667 | 2.93833333 | 3.27166667 | 2.13166667 | 1.81 | 2.03166667 | 1.96666667 | 1.93 | 2.375 | 1.785 | 2.42833333 | 2.27833333 | 1.46166667 | 2.21297619 | 0.49507022 | 0.13231308 |
| 3 | 2.72 | 1.94 | 2.37833333 | 3.00666667 | 2.015 | 1.96166667 | 1.885 | 2.22 | 1.92833333 | 2.54 | 1.64333333 | 2.09333333 | 2.05666667 | 1.33833333 | 2.12333333 | 0.42989713 | 0.11489484 |
| 4 | 2.66166667 | 1.78333333 | 2.92333333 | 3.63666667 | 2.02166667 | 2.02 | 2.12333333 | 2.14 | 2.18666667 | 2.485 | 1.655 | 2.035 | 2.06 | 1.315 | 2.21761905 | 0.57189172 | 0.15284449 |
| 5 | 2.76 | 1.93833333 | 2.18166667 | 2.50333333 | 2.01833333 | 2.005 | 1.81166667 | 2.08166667 | 2.31666667 | 2.34666667 | 1.46333333 | 2.335 | 2.21 | 1.34666667 | 2.09416667 | 0.38005496 | 0.10157396 |
| 10 | 2.17366667 | 1.656 | 1.95333333 | 2.787 | 1.97966667 | 1.89666667 | 1.95266667 | 2.083 | 2.14033333 | 2.16466667 | 1.422 | 1.70166667 | 1.26766667 | 1.562 | 1.91002381 | 0.37977473 | 0.10149907 |
| 15 | 2.28133333 | 1.25966667 | 1.07033333 | 1.85566667 | 1.808 | 1.95066667 | 1.64466667 | 1.87517241 | 1.469 | 1.75933333 | 1.36733333 | 1.525 | 1.00766667 | 1.73866667 | 1.61517898 | 0.35508056 | 0.09489927 |

### Tidal volume

Week 1

WT

| Time (min) | A | 156 | 158 | 161 | Average | SD | SE |
|---|---|---|---|---|---|---|---|
| 0 | 0.13111111 | 0.12466667 | 0.09022222 | 0.11311111 | 0.11477778 | 0.01798445 | 0.00899222 |
| 1 | 0.17833333 | 0.14833333 | 0.08333333 | 0.105 | 0.12875 | 0.04269563 | 0.02134781 |
| 2 | 0.175 | 0.13833333 | 0.08666667 | 0.125 | 0.13125 | 0.03647361 | 0.0182368 |
| 3 | 0.16333333 | 0.13166667 | 0.09166667 | 0.12 | 0.12666667 | 0.02965855 | 0.01482928 |
| 4 | 0.14833333 | 0.14333333 | 0.09333333 | 0.11833333 | 0.12583333 | 0.02533114 | 0.01266557 |
| 5 | 0.15666667 | 0.13833333 | 0.09 | 0.12333333 | 0.12708333 | 0.02823102 | 0.01411551 |
| 10 | 0.153 | 0.12866667 | 0.09433333 | 0.107 | 0.12075 | 0.02575292 | 0.01287646 |
| 15 | 0.115 | 0.12433333 | 0.086 | 0.092 | 0.10433333 | 0.01827567 | 0.00913783 |

Het

| Time (min) | B | C | D | E | F | G | H | 155 | 157 | 159 | 160 | 162 | 190 | 196 | 199 | 201 | 205 | 207 | 208 | 210 | Average | SD | SE |
|---|---|---|---|---|---|---|---|---|---|---|---|---|---|---|---|---|---|---|---|---|---|---|---|
| 0 | 0.10311111 | 0.11588889 | 0.08744444 | 0.10888889 | 0.07188889 | 0.086 | 0.10788889 | 0.09711111 | 0.08755556 | 0.08744444 | 0.11544444 | 0.07744444 | 0.14855556 | 0.10466667 | 0.10811111 | 0.08366667 | 0.07366667 | 0.08277778 | 0.07655556 | 0.068 | 0.09460556 | 0.01961347 | 0.0043857 |
| 1 | 0.14333333 | 0.13333333 | 0.105 | 0.14666667 | 0.07833333 | 0.10333333 | 0.14333333 | 0.10333333 | 0.1 | 0.10833333 | 0.11166667 | 0.09 | 0.135 | 0.12 | 0.125 | 0.11 | 0.07333333 | 0.10333333 | 0.08166667 | 0.07666667 | 0.10958333 | 0.02292165 | 0.00512544 |
| 2 | 0.14166667 | 0.135 | 0.10166667 | 0.13666667 | 0.06666667 | 0.10333333 | 0.14166667 | 0.10833333 | 0.09833333 | 0.10666667 | 0.11666667 | 0.09 | 0.14 | 0.13833333 | 0.12166667 | 0.09 | 0.08 | 0.11 | 0.08666667 | 0.08666667 | 0.11 | 0.02311932 | 0.00516964 |
| 3 | 0.13833333 | 0.12833333 | 0.09666667 | 0.12166667 | 0.07 | 0.09666667 | 0.13833333 | 0.11166667 | 0.09333333 | 0.10333333 | 0.115 | 0.095 | 0.13666667 | 0.12833333 | 0.13666667 | 0.11833333 | 0.07166667 | 0.105 | 0.09166667 | 0.08666667 | 0.10916667 | 0.02151771 | 0.00481151 |
| 4 | 0.13333333 | 0.12333333 | 0.10166667 | 0.12333333 | 0.06166667 | 0.08833333 | 0.12333333 | 0.10666667 | 0.09333333 | 0.11166667 | 0.115 | 0.09666667 | 0.12666667 | 0.12833333 | 0.10833333 | 0.11 | 0.07166667 | 0.09 | 0.08833333 | 0.09 | 0.10458333 | 0.01948815 | 0.00435768 |
| 5 | 0.12333333 | 0.12333333 | 0.09666667 | 0.12333333 | 0.06666667 | 0.08833333 | 0.11833333 | 0.09666667 | 0.095 | 0.10833333 | 0.11166667 | 0.09 | 0.125 | 0.12166667 | 0.10166667 | 0.10166667 | 0.07166667 | 0.085 | 0.07666667 | 0.085 | 0.1005 | 0.01844147 | 0.00412364 |
| 10 | 0.117 | 0.115 | 0.095 | 0.114 | 0.06666667 | 0.07933333 | 0.10866667 | 0.10266667 | 0.10366667 | 0.09633333 | 0.124 | 0.09566667 | 0.129 | 0.11966667 | 0.10366667 | 0.096 | 0.07866667 | 0.083 | 0.07366667 | 0.082 | 0.09918333 | 0.01774461 | 0.00396781 |
| 15 | 0.105 | 0.10166667 | 0.09333333 | 0.10766667 | 0.068 | 0.08866667 | 0.10566667 | 0.092 | 0.09633333 | 0.07666667 | 0.12133333 | 0.07933333 | 0.12966667 | 0.125 | 0.12733333 | 0.10566667 | 0.08166667 | 0.07433333 | 0.06533333 | 0.07166667 | 0.09581667 | 0.02020088 | 0.00451706 |

Week 2

WT

| Time (min) | A | 156 | 158 | 161 | Average | SD | SE |
|---|---|---|---|---|---|---|---|
| 0 | 0.161 | 0.14166667 | 0.10811111 | 0.13044444 | 0.13530556 | 0.02208949 | 0.01104474 |
| 1 | 0.17833333 | 0.19333333 | 0.115 | 0.13833333 | 0.15625 | 0.03598804 | 0.01799402 |
| 2 | 0.18333333 | 0.15833333 | 0.12333333 | 0.12666667 | 0.14791667 | 0.02839454 | 0.01419727 |
| 3 | 0.15666667 | 0.15333333 | 0.13333333 | 0.13166667 | 0.14375 | 0.01307917 | 0.00653959 |
| 4 | 0.14666667 | 0.175 | 0.135 | 0.13 | 0.14666667 | 0.02013841 | 0.0100692 |
| 5 | 0.14666667 | 0.16333333 | 0.11833333 | 0.12166667 | 0.1375 | 0.02136543 | 0.01068271 |
| 10 | 0.144 | 0.16933333 | 0.11233333 | 0.128 | 0.13841667 | 0.0243301 | 0.01216505 |
| 15 | 0.15266667 | 0.161 | 0.10633333 | 0.12233333 | 0.13558333 | 0.02561882 | 0.01280941 |

Het

| Time (min) | B | D | E | G | H | 155 | 157 | 160 | 162 | 190 | 196 | 199 | 201 | 205 | 207 | 208 | 210 | Average | SD | SE |
|---|---|---|---|---|---|---|---|---|---|---|---|---|---|---|---|---|---|---|---|---|
| 0 | 0.12822222 | 0.12911111 | 0.08388889 | 0.07622222 | 0.14188889 | 0.10777778 | 0.124 | 0.08922222 | 0.12422222 | 0.11888889 | 0.13166667 | 0.13422222 | 0.13422222 | 0.09355556 | 0.09122222 | 0.09244444 | 0.10811111 | 0.11228758 | 0.020826 | 0.00505105 |
| 1 | 0.15666667 | 0.15333333 | 0.10833333 | 0.09666667 | 0.14166667 | 0.135 | 0.11333333 | 0.10333333 | 0.11833333 | 0.116 | 0.162 | 0.128 | 0.164 | 0.096 | 0.102 | 0.1 | 0.086 | 0.12239216 | 0.02536518 | 0.00615196 |
| 2 | 0.16333333 | 0.15833333 | 0.11666667 | 0.1 | 0.14833333 | 0.12666667 | 0.12833333 | 0.11833333 | 0.10166667 | 0.135 | 0.165 | 0.12 | 0.11666667 | 0.08166667 | 0.09333333 | 0.085 | 0.08166667 | 0.12 | 0.02755676 | 0.0066835 |
| 3 | 0.155 | 0.16 | 0.11 | 0.105 | 0.14833333 | 0.13166667 | 0.135 | 0.10333333 | 0.09166667 | 0.12333333 | 0.17 | 0.14 | 0.11333333 | 0.08833333 | 0.1 | 0.08166667 | 0.09666667 | 0.12078431 | 0.02710651 | 0.0065743 |
| 4 | 0.15166667 | 0.15166667 | 0.10166667 | 0.10166667 | 0.14333333 | 0.12333333 | 0.13666667 | 0.11 | 0.09333333 | 0.11333333 | 0.17 | 0.14166667 | 0.13666667 | 0.08666667 | 0.09166667 | 0.075 | 0.1 | 0.11931373 | 0.02732303 | 0.00662681 |
| 5 | 0.15333333 | 0.14666667 | 0.1 | 0.09833333 | 0.13833333 | 0.12833333 | 0.13166667 | 0.10833333 | 0.09833333 | 0.125 | 0.16166667 | 0.125 | 0.14166667 | 0.08833333 | 0.09333333 | 0.075 | 0.105 | 0.11872549 | 0.02503837 | 0.0060727 |
| 10 | 0.147 | 0.12433333 | 0.09866667 | 0.092 | 0.14066667 | 0.12533333 | 0.139 | 0.10433333 | 0.10633333 | 0.123 | 0.14633333 | 0.10933333 | 0.14566667 | 0.091 | 0.101 | 0.088 | 0.11066667 | 0.11721569 | 0.0207545 | 0.00503371 |
| 15 | 0.14233333 | 0.14166667 | 0.08166667 | 0.08666667 | 0.13766667 | 0.12833333 | 0.13333333 | 0.11233333 | 0.12966667 | 0.13533333 | 0.15133333 | 0.127 | 0.141 | 0.093 | 0.10033333 | 0.09633333 | 0.10733333 | 0.12031373 | 0.02208207 | 0.00535569 |

Week 3

WT

| Time (min) | 83 | 86 | 87 | 88 | 81 | 82 | A | 156 | 158 | 161 | Average | SD | SE |
|---|---|---|---|---|---|---|---|---|---|---|---|---|---|
| 0 | 0.17933333 | 0.16622222 | 0.18033333 | 0.18155556 | 0.17123596 | 0.14744444 | 0.18066667 | 0.17322222 | 0.12122222 | 0.13655556 | 0.16377915 | 0.02130974 | 0.00673873 |
| 1 | 0.20333333 | 0.22 | 0.25333333 | 0.24666667 | 0.26666667 | 0.225 | 0.19166667 | 0.232 | 0.12666667 | 0.21166667 | 0.2177 | 0.03940842 | 0.01246204 |
| 2 | 0.18666667 | 0.195 | 0.19833333 | 0.2 | 0.24833333 | 0.235 | 0.2 | 0.22 | 0.13166667 | 0.12333333 | 0.19383333 | 0.03990769 | 0.01261992 |
| 3 | 0.17833333 | 0.20166667 | 0.18666667 | 0.20333333 | 0.23166667 | 0.22333333 | 0.2 | 0.21833333 | 0.12333333 | 0.125 | 0.18916667 | 0.03782065 | 0.01195994 |
| 4 | 0.16833333 | 0.185 | 0.18666667 | 0.19 | 0.21166667 | 0.18333333 | 0.15833333 | 0.17666667 | 0.12333333 | 0.13 | 0.17133333 | 0.02740641 | 0.00866667 |
| 5 | 0.16166667 | 0.175 | 0.17166667 | 0.185 | 0.19 | 0.145 | 0.17 | 0.16 | 0.12333333 | 0.125 | 0.16066667 | 0.02304317 | 0.00728689 |
| 10 | 0.194 | 0.174 | 0.183 | 0.17933333 | 0.177 | 0.158 | 0.165 | 0.16766667 | 0.11733333 | 0.12233333 | 0.16376667 | 0.02522983 | 0.00797837 |
| 15 | 0.18033333 | 0.16633333 | 0.18533333 | 0.18233333 | 0.194 | 0.16133333 | 0.18233333 | 0.19933333 | 0.11233333 | 0.14133333 | 0.1705 | 0.02649074 | 0.00837711 |

Het

| Time (min) | B | D | E | G | H | 155 | 157 | 190 | 196 | 199 | 201 | 205 | 207 | 208 | 210 | Average | SD | SE |
|---|---|---|---|---|---|---|---|---|---|---|---|---|---|---|---|---|---|---|
| 0 | 0.14466667 | 0.13633333 | 0.10977778 | 0.08655556 | 0.14711111 | 0.14433333 | 0.12455556 | 0.12611111 | 0.14288889 | 0.13411111 | 0.13433333 | 0.08911111 | 0.09911111 | 0.09922222 | 0.10133333 | 0.1213037 | 0.02164168 | 0.00558786 |
| 1 | 0.16666667 | 0.195 | 0.12666667 | 0.06833333 | 0.17833333 | 0.118 | 0.135 | 0.14 | 0.16 | 0.14 | 0.13333333 | 0.11666667 | 0.12666667 | 0.1 | 0.085 | 0.13264444 | 0.0338152 | 0.00873105 |
| 2 | 0.17333333 | 0.15333333 | 0.14833333 | 0.07333333 | 0.18166667 | 0.10333333 | 0.14333333 | 0.14166667 | 0.15333333 | 0.14 | 0.12333333 | 0.095 | 0.09666667 | 0.10833333 | 0.07 | 0.127 | 0.03445839 | 0.00889712 |
| 3 | 0.16166667 | 0.16333333 | 0.14666667 | 0.07333333 | 0.15833333 | 0.12166667 | 0.14833333 | 0.14833333 | 0.17 | 0.135 | 0.13833333 | 0.09 | 0.09166667 | 0.115 | 0.08833333 | 0.13 | 0.03147814 | 0.00812762 |
| 4 | 0.16333333 | 0.14833333 | 0.14166667 | 0.07 | 0.14 | 0.12333333 | 0.15833333 | 0.14833333 | 0.175 | 0.14666667 | 0.145 | 0.09166667 | 0.1 | 0.11833333 | 0.115 | 0.13233333 | 0.02883863 | 0.0074461 |
| 5 | 0.17833333 | 0.14666667 | 0.13666667 | 0.06333333 | 0.14833333 | 0.12 | 0.15666667 | 0.145 | 0.17833333 | 0.13833333 | 0.13666667 | 0.09166667 | 0.1 | 0.11666667 | 0.09 | 0.12977778 | 0.03281058 | 0.00847166 |
| 10 | 0.16566667 | 0.155 | 0.129 | 0.07433333 | 0.14266667 | 0.12233333 | 0.146 | 0.14166667 | 0.178 | 0.11766667 | 0.14933333 | 0.1 | 0.10566667 | 0.11366667 | 0.105 | 0.12973333 | 0.02780151 | 0.00717832 |
| 15 | 0.14933333 | 0.14333333 | 0.122 | 0.07766667 | 0.141 | 0.12966667 | 0.142 | 0.12266667 | 0.15666667 | 0.13733333 | 0.155 | 0.09966667 | 0.10333333 | 0.10966667 | 0.112 | 0.12675556 | 0.02279392 | 0.00588537 |

Week 4

WT

| Time (min) | A | 156 | 158 | 161 | Average | SD | SE |
|---|---|---|---|---|---|---|---|
| 0 | 0.18922222 | 0.20122222 | 0.13188889 | 0.15 | 0.16808333 | 0.0325683 | 0.01628415 |
| 1 | 0.19833333 | 0.18833333 | 0.165 | 0.14666667 | 0.17458333 | 0.02326876 | 0.01163438 |
| 2 | 0.17333333 | 0.16333333 | 0.12333333 | 0.11666667 | 0.14416667 | 0.02833333 | 0.01416667 |
| 3 | 0.18333333 | 0.165 | 0.13333333 | 0.13 | 0.15291667 | 0.02568956 | 0.01284478 |
| 4 | 0.18833333 | 0.15833333 | 0.15333333 | 0.12833333 | 0.15708333 | 0.02462214 | 0.01231107 |
| 5 | 0.16666667 | 0.16333333 | 0.12833333 | 0.12666667 | 0.14625 | 0.02170403 | 0.01085201 |
| 10 | 0.17 | 0.19966667 | 0.13133333 | 0.14566667 | 0.16166667 | 0.02994192 | 0.01497096 |
| 15 | 0.17766667 | 0.21866667 | 0.14766667 | 0.15033333 | 0.17358333 | 0.03297179 | 0.01648589 |

Het

| Time (min) | B | D | E | H | 155 | 157 | 190 | 196 | 199 | 201 | 205 | 207 | 208 | 210 | Average | SD | SE |
|---|---|---|---|---|---|---|---|---|---|---|---|---|---|---|---|---|---|
| 0 | 0.16044444 | 0.14922222 | 0.11455556 | 0.14477778 | 0.141 | 0.16055556 | 0.14011111 | 0.16 | 0.15033333 | 0.14288889 | 0.11433333 | 0.10277778 | 0.10933333 | 0.11577778 | 0.13615079 | 0.0205238 | 0.00548522 |
| 1 | 0.19333333 | 0.15166667 | 0.125 | 0.17833333 | 0.16333333 | 0.18 | 0.15833333 | 0.17833333 | 0.15166667 | 0.15333333 | 0.10666667 | 0.13 | 0.135 | 0.11666667 | 0.15154762 | 0.02609536 | 0.00697428 |
| 2 | 0.16 | 0.17 | 0.14833333 | 0.185 | 0.125 | 0.13833333 | 0.135 | 0.15666667 | 0.165 | 0.13666667 | 0.11833333 | 0.11833333 | 0.145 | 0.10333333 | 0.14321429 | 0.02274406 | 0.00607861 |
| 3 | 0.14833333 | 0.17333333 | 0.15166667 | 0.19166667 | 0.135 | 0.14666667 | 0.14833333 | 0.17166667 | 0.17 | 0.14666667 | 0.125 | 0.11 | 0.13833333 | 0.10833333 | 0.1475 | 0.02388317 | 0.00638305 |
| 4 | 0.15833333 | 0.15 | 0.13833333 | 0.14666667 | 0.14 | 0.15 | 0.155 | 0.175 | 0.13666667 | 0.15833333 | 0.13 | 0.10833333 | 0.125 | 0.11666667 | 0.14202381 | 0.01796457 | 0.00480123 |
| 5 | 0.165 | 0.13833333 | 0.12166667 | 0.14333333 | 0.13666667 | 0.155 | 0.15833333 | 0.17666667 | 0.15333333 | 0.15 | 0.12833333 | 0.12 | 0.12666667 | 0.12166667 | 0.1425 | 0.01782812 | 0.00476477 |
| 10 | 0.18266667 | 0.15366667 | 0.12766667 | 0.14266667 | 0.13933333 | 0.16033333 | 0.158 | 0.177 | 0.13966667 | 0.169 | 0.12866667 | 0.11466667 | 0.122 | 0.12833333 | 0.14597619 | 0.02120372 | 0.00566693 |
| 15 | 0.17 | 0.156 | 0.12433333 | 0.157 | 0.15066667 | 0.177 | 0.16433333 | 0.17482759 | 0.167 | 0.17366667 | 0.12566667 | 0.11566667 | 0.11066667 | 0.12633333 | 0.14951149 | 0.0238891 | 0.00638463 |

### Expiratory volume

Week 1

WT

| Time (min) | Animal ID: A | 156 | 158 | 161 | Average | SD | SE |
|---|---|---|---|---|---|---|---|
| 0 | 0.13677778 | 0.128 | 0.09111111 | 0.116 | 0.11797222 | 0.01982939 | 0.0099147 |
| 1 | 0.18333333 | 0.14666667 | 0.08833333 | 0.105 | 0.13083333 | 0.04274169 | 0.02137084 |
| 2 | 0.17166667 | 0.13833333 | 0.085 | 0.11666667 | 0.12791667 | 0.03647361 | 0.0182368 |
| 3 | 0.16 | 0.13 | 0.09333333 | 0.12 | 0.12583333 | 0.02753785 | 0.01376893 |
| 4 | 0.15333333 | 0.14333333 | 0.095 | 0.12666667 | 0.12958333 | 0.02554499 | 0.01277249 |
| 5 | 0.165 | 0.14333333 | 0.09166667 | 0.12666667 | 0.13166667 | 0.030942 | 0.015471 |
| 10 | 0.15666667 | 0.13233333 | 0.09633333 | 0.11366667 | 0.12475 | 0.02586199 | 0.012931 |
| 15 | 0.11466667 | 0.12433333 | 0.088 | 0.09233333 | 0.10483333 | 0.01747909 | 0.00873954 |

Het

| Time (min) | Animal ID: B | C | D | E | F | G | H | 155 | 157 | 159 | 160 | 162 | 190 | 196 | 199 | 201 | 205 | 207 | 208 | 210 | Average | SD | SE |
|---|---|---|---|---|---|---|---|---|---|---|---|---|---|---|---|---|---|---|---|---|---|---|---|
| 0 | 0.10488889 | 0.121 | 0.09333333 | 0.11566667 | 0.07644444 | 0.08866667 | 0.11733333 | 0.10022222 | 0.09088889 | 0.09066667 | 0.118 | 0.07888889 | 0.14633333 | 0.10722222 | 0.10822222 | 0.085 | 0.07844444 | 0.08522222 | 0.08366667 | 0.07022222 | 0.09801667 | 0.01909503 | 0.00426978 |
| 1 | 0.14666667 | 0.13166667 | 0.10333333 | 0.14833333 | 0.06166667 | 0.10333333 | 0.14166667 | 0.10333333 | 0.10166667 | 0.11166667 | 0.11166667 | 0.09 | 0.13833333 | 0.12833333 | 0.125 | 0.11166667 | 0.075 | 0.10333333 | 0.08833333 | 0.07666667 | 0.11008333 | 0.02454543 | 0.00548852 |
| 2 | 0.145 | 0.13666667 | 0.10333333 | 0.14166667 | 0.06666667 | 0.09833333 | 0.13833333 | 0.105 | 0.095 | 0.10666667 | 0.11833333 | 0.09333333 | 0.13333333 | 0.13833333 | 0.12333333 | 0.09 | 0.08 | 0.10833333 | 0.09166667 | 0.09166667 | 0.11025 | 0.02283474 | 0.005106 |
| 3 | 0.13 | 0.12833333 | 0.09833333 | 0.12 | 0.06666667 | 0.09666667 | 0.135 | 0.11 | 0.09166667 | 0.105 | 0.115 | 0.09166667 | 0.135 | 0.13 | 0.13166667 | 0.11166667 | 0.07666667 | 0.10666667 | 0.09 | 0.08333333 | 0.10766667 | 0.02051529 | 0.00458736 |
| 4 | 0.13833333 | 0.125 | 0.10333333 | 0.12833333 | 0.065 | 0.09 | 0.12166667 | 0.105 | 0.09166667 | 0.11166667 | 0.115 | 0.09666667 | 0.12166667 | 0.13166667 | 0.11166667 | 0.11333333 | 0.07166667 | 0.09166667 | 0.08666667 | 0.09333333 | 0.10566667 | 0.01974546 | 0.00441522 |
| 5 | 0.12833333 | 0.125 | 0.09666667 | 0.13 | 0.06666667 | 0.09 | 0.11833333 | 0.1 | 0.09666667 | 0.10833333 | 0.11666667 | 0.09333333 | 0.13 | 0.12166667 | 0.105 | 0.105 | 0.075 | 0.08833333 | 0.085 | 0.08833333 | 0.10341667 | 0.0186337 | 0.00416662 |
| 10 | 0.11933333 | 0.11533333 | 0.09566667 | 0.11466667 | 0.068 | 0.083 | 0.11733333 | 0.10233333 | 0.10333333 | 0.1 | 0.12433333 | 0.09966667 | 0.13033333 | 0.11966667 | 0.10733333 | 0.10133333 | 0.08166667 | 0.086 | 0.076 | 0.084 | 0.10146667 | 0.01739409 | 0.00388944 |
| 15 | 0.10466667 | 0.101 | 0.09733333 | 0.108 | 0.072 | 0.09433333 | 0.11066667 | 0.09166667 | 0.09633333 | 0.07966667 | 0.12166667 | 0.07966667 | 0.133 | 0.12566667 | 0.12766667 | 0.11 | 0.08666667 | 0.07633333 | 0.06366667 | 0.07333333 | 0.09766667 | 0.02010033 | 0.00449457 |

Week 2

WT

| Time (min) | Animal ID: A | 156 | 158 | 161 | Average | SD | SE |
|---|---|---|---|---|---|---|---|
| 0 | 0.16377778 | 0.14255556 | 0.11311111 | 0.13366667 | 0.13827778 | 0.02100137 | 0.01050069 |
| 1 | 0.18 | 0.19666667 | 0.11166667 | 0.13333333 | 0.15541667 | 0.03961329 | 0.01980665 |
| 2 | 0.17166667 | 0.165 | 0.12833333 | 0.12833333 | 0.14833333 | 0.02325383 | 0.01162692 |
| 3 | 0.15333333 | 0.15166667 | 0.12833333 | 0.13333333 | 0.14166667 | 0.01269296 | 0.00634648 |
| 4 | 0.15166667 | 0.175 | 0.12833333 | 0.13 | 0.14625 | 0.0219163 | 0.01095815 |
| 5 | 0.15 | 0.16666667 | 0.12 | 0.125 | 0.14041667 | 0.02187401 | 0.010937 |
| 10 | 0.14766667 | 0.17166667 | 0.114 | 0.13133333 | 0.14116667 | 0.02454399 | 0.01227199 |
| 15 | 0.15466667 | 0.16233333 | 0.10833333 | 0.123 | 0.13708333 | 0.02563617 | 0.01281808 |

Het

| Time (min) | Animal ID: B | D | E | G | H | 155 | 157 | 160 | 162 | 190 | 196 | 199 | 201 | 205 | 207 | 208 | 210 | Average | SD | SE |
|---|---|---|---|---|---|---|---|---|---|---|---|---|---|---|---|---|---|---|---|---|
| 0 | 0.12844444 | 0.13433333 | 0.08477778 | 0.07666667 | 0.14733333 | 0.10711111 | 0.12466667 | 0.09066667 | 0.135 | 0.12088889 | 0.13188889 | 0.135 | 0.13833333 | 0.098 | 0.095 | 0.09977778 | 0.10988889 | 0.1151634 | 0.0214204 | 0.00519521 |
| 1 | 0.16166667 | 0.155 | 0.11 | 0.095 | 0.14166667 | 0.13666667 | 0.115 | 0.10666667 | 0.12 | 0.114 | 0.16 | 0.132 | 0.166 | 0.098 | 0.102 | 0.1 | 0.09 | 0.1237451 | 0.02549715 | 0.00618397 |
| 2 | 0.16333333 | 0.15333333 | 0.115 | 0.1 | 0.145 | 0.12333333 | 0.12666667 | 0.12333333 | 0.105 | 0.13333333 | 0.17 | 0.12 | 0.11666667 | 0.08333333 | 0.09333333 | 0.08833333 | 0.08166667 | 0.12009804 | 0.02686431 | 0.00651555 |
| 3 | 0.16 | 0.16 | 0.115 | 0.105 | 0.15 | 0.12833333 | 0.13333333 | 0.10333333 | 0.095 | 0.125 | 0.17333333 | 0.135 | 0.115 | 0.08833333 | 0.1 | 0.08333333 | 0.10166667 | 0.12186275 | 0.0269766 | 0.00654279 |
| 4 | 0.155 | 0.14833333 | 0.105 | 0.09666667 | 0.14666667 | 0.12333333 | 0.13666667 | 0.11333333 | 0.095 | 0.11166667 | 0.17166667 | 0.14833333 | 0.13666667 | 0.08666667 | 0.09666667 | 0.08 | 0.09833333 | 0.12058824 | 0.02739209 | 0.00664356 |
| 5 | 0.15166667 | 0.15 | 0.10666667 | 0.095 | 0.14 | 0.125 | 0.13166667 | 0.10666667 | 0.09833333 | 0.13 | 0.165 | 0.11833333 | 0.14166667 | 0.09 | 0.09333333 | 0.07666667 | 0.105 | 0.11911765 | 0.02526332 | 0.00612725 |
| 10 | 0.14766667 | 0.127 | 0.10133333 | 0.09166667 | 0.14133333 | 0.12333333 | 0.13933333 | 0.10533333 | 0.10766667 | 0.123 | 0.149 | 0.115 | 0.14466667 | 0.09366667 | 0.10166667 | 0.08933333 | 0.10933333 | 0.1182549 | 0.02044557 | 0.00495878 |
| 15 | 0.144 | 0.14133333 | 0.08266667 | 0.08966667 | 0.14033333 | 0.12866667 | 0.13533333 | 0.11266667 | 0.133 | 0.13466667 | 0.14433333 | 0.13133333 | 0.14066667 | 0.094 | 0.102 | 0.098 | 0.10733333 | 0.12117647 | 0.0213236 | 0.00517173 |

Week 3

WT

| Time (min) | Animal ID: 83 | 86 | 87 | 88 | 81 | 82 | A | 156 | 158 | 161 | Average | SD | SE |
|---|---|---|---|---|---|---|---|---|---|---|---|---|---|
| 0 | 0.17233333 | 0.16455556 | 0.17822222 | 0.17844444 | 0.16820225 | 0.148 | 0.18511111 | 0.17566667 | 0.12277778 | 0.13722222 | 0.16305356 | 0.02039466 | 0.00644936 |
| 1 | 0.19666667 | 0.21666667 | 0.25833333 | 0.25 | 0.27333333 | 0.22333333 | 0.19166667 | 0.226 | 0.12166667 | 0.19333333 | 0.2151 | 0.0431532 | 0.01364624 |
| 2 | 0.185 | 0.18833333 | 0.19833333 | 0.21166667 | 0.25 | 0.23666667 | 0.20833333 | 0.215 | 0.125 | 0.125 | 0.19433333 | 0.0415903 | 0.01315201 |
| 3 | 0.185 | 0.19833333 | 0.18 | 0.21333333 | 0.225 | 0.225 | 0.19166667 | 0.21666667 | 0.12333333 | 0.13333333 | 0.18916667 | 0.0358258 | 0.01132911 |
| 4 | 0.16166667 | 0.17666667 | 0.185 | 0.19 | 0.20666667 | 0.18 | 0.15833333 | 0.18 | 0.12333333 | 0.13666667 | 0.16983333 | 0.02518389 | 0.00796385 |
| 5 | 0.17 | 0.17166667 | 0.18666667 | 0.19 | 0.18166667 | 0.145 | 0.165 | 0.155 | 0.12833333 | 0.13166667 | 0.1625 | 0.0219462 | 0.00694 |
| 10 | 0.19666667 | 0.172 | 0.18833333 | 0.182 | 0.17233333 | 0.16766667 | 0.16966667 | 0.169 | 0.119 | 0.12666667 | 0.16633333 | 0.02483625 | 0.00785391 |
| 15 | 0.18 | 0.17066667 | 0.19166667 | 0.18766667 | 0.182 | 0.17166667 | 0.18366667 | 0.197 | 0.113 | 0.144 | 0.17213333 | 0.02543848 | 0.00804435 |

Het

| Time (min) | Animal ID: B | D | E | G | H | 155 | 157 | 190 | 196 | 199 | 201 | 205 | 207 | 208 | 210 | Average | SD | SE |
|---|---|---|---|---|---|---|---|---|---|---|---|---|---|---|---|---|---|---|
| 0 | 0.14433333 | 0.14011111 | 0.114 | 0.09222222 | 0.15211111 | 0.14077778 | 0.12444444 | 0.12822222 | 0.14511111 | 0.13766667 | 0.13633333 | 0.09122222 | 0.1 | 0.09944444 | 0.10122222 | 0.12314815 | 0.02141171 | 0.00552848 |
| 1 | 0.16666667 | 0.19666667 | 0.12833333 | 0.06666667 | 0.17666667 | 0.12 | 0.13666667 | 0.14333333 | 0.16666667 | 0.13666667 | 0.13333333 | 0.11833333 | 0.12666667 | 0.1 | 0.08833333 | 0.13366667 | 0.03410535 | 0.00880596 |
| 2 | 0.18 | 0.15333333 | 0.145 | 0.07333333 | 0.175 | 0.10333333 | 0.14333333 | 0.14166667 | 0.15166667 | 0.14166667 | 0.12333333 | 0.09666667 | 0.10166667 | 0.10833333 | 0.07 | 0.12722222 | 0.033857 | 0.00874184 |
| 3 | 0.16166667 | 0.15833333 | 0.14333333 | 0.075 | 0.15333333 | 0.12333333 | 0.15 | 0.14833333 | 0.17 | 0.13666667 | 0.13666667 | 0.09 | 0.095 | 0.115 | 0.08833333 | 0.12966667 | 0.03024503 | 0.00780923 |
| 4 | 0.17333333 | 0.15166667 | 0.13666667 | 0.07 | 0.14666667 | 0.12333333 | 0.155 | 0.14666667 | 0.18 | 0.145 | 0.14666667 | 0.09166667 | 0.10166667 | 0.11666667 | 0.11666667 | 0.13344444 | 0.03008234 | 0.00776723 |
| 5 | 0.17166667 | 0.14833333 | 0.12666667 | 0.07 | 0.14833333 | 0.12 | 0.155 | 0.14666667 | 0.18 | 0.14 | 0.13666667 | 0.09166667 | 0.10166667 | 0.11666667 | 0.09 | 0.12955556 | 0.03129777 | 0.00808105 |
| 10 | 0.163 | 0.155 | 0.12666667 | 0.07633333 | 0.15333333 | 0.122 | 0.14533333 | 0.14266667 | 0.174 | 0.12266667 | 0.14866667 | 0.10033333 | 0.106 | 0.11366667 | 0.10366667 | 0.13022222 | 0.02720693 | 0.0070248 |
| 15 | 0.14766667 | 0.14266667 | 0.123 | 0.08333333 | 0.14966667 | 0.12966667 | 0.14066667 | 0.12666667 | 0.16166667 | 0.139 | 0.15333333 | 0.1 | 0.102 | 0.11066667 | 0.112 | 0.12813333 | 0.02262875 | 0.00584272 |

Week 4

WT

| Time (min) | Animal ID: A | 156 | 158 | 161 | Average | SD | SE |
|---|---|---|---|---|---|---|---|
| 0 | 0.19044444 | 0.195 | 0.13122222 | 0.15277778 | 0.16736111 | 0.03063463 | 0.01531731 |
| 1 | 0.19 | 0.18833333 | 0.15166667 | 0.155 | 0.17125 | 0.02074425 | 0.01037213 |
| 2 | 0.17833333 | 0.165 | 0.11 | 0.12 | 0.14333333 | 0.03341656 | 0.01670828 |
| 3 | 0.18 | 0.16666667 | 0.14333333 | 0.13166667 | 0.15541667 | 0.0219163 | 0.01095815 |
| 4 | 0.18 | 0.16333333 | 0.175 | 0.13666667 | 0.16375 | 0.01935894 | 0.00967947 |
| 5 | 0.165 | 0.16666667 | 0.125 | 0.13166667 | 0.14708333 | 0.02183164 | 0.01091582 |
| 10 | 0.168 | 0.20066667 | 0.138 | 0.14733333 | 0.1635 | 0.02776822 | 0.01388411 |
| 15 | 0.174 | 0.21466667 | 0.14733333 | 0.15066667 | 0.17166667 | 0.03102448 | 0.01551224 |

Het

| Time (min) | Animal ID: B | D | E | H | 155 | 157 | 190 | 196 | 199 | 201 | 205 | 207 | 208 | 210 | Average | SD | SE |
|---|---|---|---|---|---|---|---|---|---|---|---|---|---|---|---|---|---|
| 0 | 0.16133333 | 0.13555556 | 0.11555556 | 0.14877778 | 0.14266667 | 0.16433333 | 0.14155556 | 0.16033333 | 0.15311111 | 0.146 | 0.11755556 | 0.10355556 | 0.11577778 | 0.11644444 | 0.1373254 | 0.02008791 | 0.00536872 |
| 1 | 0.19833333 | 0.155 | 0.12833333 | 0.17166667 | 0.16333333 | 0.18166667 | 0.16166667 | 0.18333333 | 0.15666667 | 0.15666667 | 0.11166667 | 0.12666667 | 0.13666667 | 0.11833333 | 0.15357143 | 0.0260658 | 0.00696638 |
| 2 | 0.16333333 | 0.16 | 0.14833333 | 0.18166667 | 0.125 | 0.14166667 | 0.135 | 0.16 | 0.16333333 | 0.13333333 | 0.12166667 | 0.11666667 | 0.13833333 | 0.10833333 | 0.14261905 | 0.02108909 | 0.0056363 |
| 3 | 0.155 | 0.16666667 | 0.14833333 | 0.195 | 0.13833333 | 0.14666667 | 0.15 | 0.16833333 | 0.16833333 | 0.14833333 | 0.12333333 | 0.105 | 0.14 | 0.11333333 | 0.14761905 | 0.02362197 | 0.00631324 |
| 4 | 0.16666667 | 0.145 | 0.14 | 0.15166667 | 0.13833333 | 0.14833333 | 0.15833333 | 0.17333333 | 0.13666667 | 0.16 | 0.13166667 | 0.11166667 | 0.12833333 | 0.11833333 | 0.14345238 | 0.0177887 | 0.00475423 |
| 5 | 0.17333333 | 0.13666667 | 0.12333333 | 0.14166667 | 0.13666667 | 0.15666667 | 0.15833333 | 0.175 | 0.15666667 | 0.15 | 0.13 | 0.12166667 | 0.12666667 | 0.12166667 | 0.14345238 | 0.01842592 | 0.00492454 |
| 10 | 0.18066667 | 0.16133333 | 0.13166667 | 0.14466667 | 0.13933333 | 0.16133333 | 0.16033333 | 0.17533333 | 0.142 | 0.16966667 | 0.12966667 | 0.11533333 | 0.12366667 | 0.12766667 | 0.14733333 | 0.02072325 | 0.00553852 |
| 15 | 0.16966667 | 0.13466667 | 0.12533333 | 0.15933333 | 0.15133333 | 0.177 | 0.16366667 | 0.1737931 | 0.167 | 0.17133333 | 0.127 | 0.11933333 | 0.11366667 | 0.12466667 | 0.14841379 | 0.02310733 | 0.00617569 |

### Frequency

Week 1

WT

| Time (min) | A | 156 | 158 | 161 | Average | SD | SE |
|---|---|---|---|---|---|---|---|
| 0 | 406.4 | 456.8 | 367.911111 | 388.422222 | 404.883333 | 38.0155843 | 19.0077922 |
| 1 | 370.833333 | 428.833333 | 396.666667 | 389.833333 | 396.541667 | 24.1432358 | 12.0716179 |
| 2 | 424.166667 | 413.166667 | 435.333333 | 444.333333 | 429.25 | 13.5280915 | 6.76404574 |
| 3 | 404.833333 | 464.666667 | 387.333333 | 440.833333 | 424.416667 | 34.8723864 | 17.4361932 |
| 4 | 397.833333 | 385.166667 | 375.666667 | 376.333333 | 383.75 | 10.3391561 | 5.16957803 |
| 5 | 355.5 | 386.666667 | 401.833333 | 373.5 | 379.375 | 19.6814328 | 9.84071639 |
| 10 | 367.7 | 393.833333 | 339.033333 | 394.5 | 373.766667 | 26.3043159 | 13.152158 |
| 15 | 276.7 | 284.066667 | 267.666667 | 290.966667 | 279.85 | 9.99527666 | 4.99763833 |

Het

| Time (min) | B | C | D | E | F | G | H | 155 | 157 | 159 | 160 | 162 | 190 | 196 | 199 | 201 | 205 | 207 | 208 | 210 | Average | SD | SE |
|---|---|---|---|---|---|---|---|---|---|---|---|---|---|---|---|---|---|---|---|---|---|---|---|
| 0 | 286.3 | 384.233333 | 373.433333 | 396.411111 | 285.011111 | 359.266667 | 422.7 | 452.355556 | 496.811111 | 409.311111 | 435.111111 | 284.022222 | 390.055556 | 290.577778 | 233.6 | 516.1 | 426.566667 | 355.211111 | 348.666667 | 257.111111 | 370.142778 | 78.5899084 | 17.5732378 |
| 1 | 430.333333 | 281.166667 | 397.666667 | 407.5 | 260.166667 | 475.833333 | 430.333333 | 446.166667 | 360.833333 | 366.5 | 444.166667 | 325.833333 | 407.166667 | 397.833333 | 393.833333 | 342 | 373.333333 | 364.333333 | 386.666667 | 343.333333 | 381.75 | 54.1612143 | 12.1108157 |
| 2 | 388 | 396.5 | 380.666667 | 403.333333 | 248.666667 | 448.666667 | 460.833333 | 409.833333 | 295.833333 | 357.5 | 407.666667 | 345.666667 | 481.166667 | 370.666667 | 426.333333 | 376.166667 | 315 | 368.833333 | 380.833333 | 354.166667 | 380.816667 | 54.9118619 | 12.2786656 |
| 3 | 370.833333 | 402.833333 | 389.333333 | 452.166667 | 236.5 | 476.833333 | 446.166667 | 395 | 361.5 | 410 | 424 | 362.5 | 489.666667 | 355 | 367.666667 | 329.5 | 322.666667 | 361.666667 | 348.666667 | 355 | 382.875 | 58.2082588 | 13.0157623 |
| 4 | 364.666667 | 418.666667 | 374.333333 | 454.5 | 232.333333 | 505.833333 | 381.833333 | 394.833333 | 338.333333 | 372.5 | 370.833333 | 370.5 | 497.5 | 355.333333 | 454.5 | 384.333333 | 311.833333 | 454 | 369.833333 | 388.5 | 389.75 | 63.0467083 | 14.0976726 |
| 5 | 352 | 404.5 | 390.333333 | 391.5 | 200.666667 | 509.166667 | 375.166667 | 413.833333 | 367 | 358.833333 | 384.5 | 355 | 349 | 292 | 445.833333 | 352.5 | 307.833333 | 445 | 441.166667 | 333.833333 | 373.483333 | 65.3531992 | 14.6134196 |
| 10 | 274.9 | 364.366667 | 290.8 | 236.3 | 203.133333 | 494.3 | 373.933333 | 357.966667 | 265.433333 | 327.6 | 307.666667 | 343.533333 | 385.766667 | 264.433333 | 401.5 | 400.866667 | 330 | 355.1 | 260.2 | 292.133333 | 326.496667 | 68.6047318 | 15.3404844 |
| 15 | 277 | 308.8 | 319.1 | 244.566667 | 221.866667 | 354.633333 | 329.5 | 323.433333 | 232.466667 | 248.066667 | 262.8 | 248.933333 | 302.533333 | 242.666667 | 234 | 288.333333 | 341 | 237.4 | 250.433333 | 242.833333 | 275.518333 | 41.6285601 | 9.30842902 |

Week 2

WT

| Time (min) | A | 156 | 158 | 161 | Average | SD | SE |
|---|---|---|---|---|---|---|---|
| 0 | 351.1 | 280.922222 | 413.855556 | 441.255556 | 371.783333 | 71.3676008 | 35.6838004 |
| 1 | 449.666667 | 397.833333 | 426.833333 | 444.833333 | 429.791667 | 23.4617005 | 11.7308502 |
| 2 | 425.5 | 408 | 431.166667 | 426.166667 | 422.708333 | 10.1264002 | 5.06320011 |
| 3 | 507 | 445.5 | 402.333333 | 381.333333 | 434.041667 | 55.4911363 | 27.7455681 |
| 4 | 464.166667 | 332.666667 | 413.833333 | 393.333333 | 401 | 54.4149013 | 27.2074506 |
| 5 | 398.5 | 355.833333 | 411.833333 | 404 | 392.541667 | 25.0763187 | 12.5381593 |
| 10 | 397.033333 | 345.233333 | 328.833333 | 362.633333 | 358.433333 | 29.2004566 | 14.6002283 |
| 15 | 309.866667 | 270.333333 | 265.033333 | 289.9 | 283.783333 | 20.4142017 | 10.2071008 |

Het

| Time (min) | B | D | E | G | H | 155 | 157 | 160 | 162 | 190 | 196 | 199 | 201 | 205 | 207 | 208 | 210 | Average | SD | SE |
|---|---|---|---|---|---|---|---|---|---|---|---|---|---|---|---|---|---|---|---|---|
| 0 | 247.488889 | 333.911111 | 236.077778 | 282.688889 | 383.233333 | 290.733333 | 257.877778 | 457.355556 | 308.811111 | 387.1 | 247.133333 | 230.811111 | 296.822222 | 532.922222 | 423.244444 | 515.633333 | 497.822222 | 348.803922 | 103.585798 | 25.1232463 |
| 1 | 429.833333 | 452.5 | 378 | 420.833333 | 449 | 368 | 339.5 | 491 | 331.166667 | 360.6 | 408.2 | 295.2 | 282.6 | 320.8 | 339.8 | 372.6 | 393.4 | 378.413725 | 57.9997522 | 14.0670062 |
| 2 | 457.333333 | 493.5 | 421.333333 | 418.333333 | 438.5 | 417.666667 | 370.5 | 400.5 | 251.833333 | 445.333333 | 435.833333 | 434 | 342.166667 | 317.833333 | 354.333333 | 313.333333 | 319.333333 | 390.098039 | 64.4529075 | 15.6321262 |
| 3 | 447.5 | 472.333333 | 388.166667 | 411.5 | 451 | 358.833333 | 313 | 466.333333 | 296 | 411.833333 | 388 | 400.5 | 365.333333 | 351 | 306.5 | 232.5 | 377.166667 | 378.676471 | 65.3445587 | 15.8483834 |
| 4 | 444.333333 | 497.166667 | 463 | 414.5 | 363.5 | 401.5 | 301 | 403.5 | 322 | 357.166667 | 361.833333 | 385.333333 | 274.5 | 302.166667 | 275.666667 | 227 | 289.666667 | 357.872549 | 74.5976134 | 18.0925788 |
| 5 | 384.666667 | 448.166667 | 360.666667 | 456.5 | 307.166667 | 340.5 | 289.5 | 367.333333 | 307.166667 | 424.833333 | 317.333333 | 512 | 268.833333 | 309.5 | 268.5 | 245.333333 | 313.166667 | 348.303922 | 75.0247467 | 18.1961738 |
| 10 | 376.466667 | 456.5 | 324.366667 | 403 | 286.466667 | 346.533333 | 281.233333 | 320.1 | 299.533333 | 340.4 | 375.866667 | 416.533333 | 308.233333 | 333 | 308.833333 | 281.3 | 314.133333 | 339.558824 | 50.495006 | 12.2468378 |
| 15 | 370.933333 | 261.3 | 181.466667 | 353.266667 | 297.5 | 292.4 | 354.733333 | 234.366667 | 178.7 | 262.933333 | 315.466667 | 324.233333 | 396.266667 | 273.066667 | 315.766667 | 315.033333 | 228.033333 | 291.498039 | 62.307678 | 15.1118316 |

Week 3

WT

| Time (min) | 83 | 86 | 87 | 88 | 81 | 82 | A | 156 | 158 | 161 | Average | SD | SE |
|---|---|---|---|---|---|---|---|---|---|---|---|---|---|
| 0 | 414.288889 | 257.2 | 349.2 | 430.655556 | 275.58427 | 255.566667 | 417.777778 | 322.288889 | 423.355556 | 338.511111 | 348.442871 | 70.3615533 | 22.2502768 |
| 1 | 425.333333 | 433 | 497 | 519.333333 | 437.166667 | 495.333333 | 530 | 480.6 | 433.333333 | 477 | 472.81 | 38.4036137 | 12.144289 |
| 2 | 381.666667 | 487 | 530 | 478.666667 | 411.333333 | 456.833333 | 489.833333 | 491 | 411.166667 | 474.166667 | 461.166667 | 45.8437362 | 14.4970623 |
| 3 | 368.5 | 454.166667 | 467.166667 | 441.333333 | 359.333333 | 360.666667 | 418.666667 | 437.666667 | 452.166667 | 427.166667 | 418.683333 | 40.9375051 | 12.9455758 |
| 4 | 395.666667 | 457.5 | 472 | 484.333333 | 376.666667 | 417.666667 | 572.666667 | 542.333333 | 445.166667 | 404.666667 | 456.866667 | 63.4856745 | 20.075933 |
| 5 | 343.166667 | 416.5 | 443 | 522.5 | 446 | 531 | 416.666667 | 520.666667 | 353.833333 | 487 | 448.033333 | 67.5780208 | 21.3700465 |
| 10 | 295.533333 | 432 | 393.833333 | 405.433333 | 474.866667 | 426.5 | 431.233333 | 449.433333 | 361.966667 | 464.666667 | 413.546667 | 53.2545951 | 16.8405816 |
| 15 | 364.1 | 309.3 | 275.266667 | 263.4 | 357.733333 | 340.533333 | 302.633333 | 219.2 | 266.633333 | 408.133333 | 310.693333 | 57.0778606 | 18.0496044 |

Het

| Time (min) | B | D | E | G | H | 155 | 157 | 190 | 196 | 199 | 201 | 205 | 207 | 208 | 210 | Average | SD | SE |
|---|---|---|---|---|---|---|---|---|---|---|---|---|---|---|---|---|---|---|
| 0 | 261.411111 | 365.433333 | 288.488889 | 216.077778 | 374.722222 | 475.133333 | 254.866667 | 296.2 | 256.633333 | 287.055556 | 268.488889 | 269.766667 | 297.566667 | 261.211111 | 258.622222 | 295.445185 | 64.4311796 | 16.636059 |
| 1 | 458.833333 | 295.333333 | 465.166667 | 249.666667 | 425.333333 | 432.4 | 470.666667 | 360 | 366 | 321 | 333.166667 | 320.5 | 332.833333 | 290 | 386.5 | 367.16 | 69.8719057 | 18.0408485 |
| 2 | 492.166667 | 469.333333 | 504 | 214.166667 | 435.5 | 396.666667 | 378 | 351.333333 | 347.333333 | 325.333333 | 398.333333 | 280.5 | 334 | 298 | 371 | 373.044444 | 80.2271445 | 20.7145596 |
| 3 | 398.333333 | 461 | 529.833333 | 236 | 509.166667 | 391.833333 | 386.5 | 329.333333 | 336.166667 | 378 | 377.833333 | 284.5 | 283.5 | 316.166667 | 441.666667 | 377.322222 | 83.5160489 | 21.5637511 |
| 4 | 514.666667 | 323.833333 | 481.333333 | 225 | 509.666667 | 371.5 | 386 | 319.166667 | 360 | 398.833333 | 362.333333 | 287.666667 | 273.333333 | 297.166667 | 451 | 370.766667 | 87.7493228 | 22.6567777 |
| 5 | 452.833333 | 292.333333 | 547.333333 | 258 | 486.5 | 355.5 | 361.166667 | 337 | 312.833333 | 356 | 419.5 | 286.833333 | 268 | 280.666667 | 317.666667 | 355.477778 | 85.6174554 | 22.1063319 |
| 10 | 335.7 | 310.566667 | 278.433333 | 234.966667 | 422.366667 | 354.133333 | 339.366667 | 354.833333 | 319.633333 | 464.6 | 336.8 | 336.2 | 269.833333 | 364.333333 | 340.266667 | 337.468889 | 56.2493181 | 14.5235115 |
| 15 | 334.966667 | 225.833333 | 342.8 | 213.966667 | 436.633333 | 347.733333 | 408.533333 | 428.866667 | 407.433333 | 279.133333 | 236.1 | 275.5 | 255.3 | 293.733333 | 339.2 | 321.715556 | 74.857721 | 19.3281804 |

Week 4

WT

| Time (min) | A | 156 | 158 | 161 | Average | SD | SE |
|---|---|---|---|---|---|---|---|
| 0 | 383.244444 | 386.511111 | 285.355556 | 507.411111 | 390.630556 | 90.9066339 | 45.4533169 |
| 1 | 542.5 | 511.333333 | 424 | 417.833333 | 473.916667 | 62.5584911 | 31.2792456 |
| 2 | 491.833333 | 524.5 | 514.166667 | 454 | 496.125 | 31.2173904 | 15.6086952 |
| 3 | 464.333333 | 507.333333 | 361.833333 | 425.166667 | 439.666667 | 61.7940127 | 30.8970063 |
| 4 | 517.666667 | 530.666667 | 383.333333 | 407.5 | 459.791667 | 75.1733029 | 37.5866514 |
| 5 | 483.833333 | 481.666667 | 528.666667 | 416.666667 | 477.708333 | 46.101413 | 23.0507065 |
| 10 | 386.066667 | 359.066667 | 431.866667 | 347.466667 | 381.116667 | 37.4992889 | 18.7496444 |
| 15 | 268.2 | 263.5 | 258.1 | 331.3 | 280.275 | 34.2660546 | 17.1330273 |

Het

| Time (min) | B | D | E | H | 155 | 157 | 190 | 196 | 199 | 201 | 205 | 207 | 208 | 210 | Average | SD | SE |
|---|---|---|---|---|---|---|---|---|---|---|---|---|---|---|---|---|---|
| 0 | 279.977778 | 141.866667 | 264.866667 | 331.866667 | 279.533333 | 271.222222 | 271.422222 | 246.011111 | 245.444444 | 311.922222 | 326.9 | 250.211111 | 493.911111 | 251.488889 | 283.331746 | 75.9663991 | 20.3028742 |
| 1 | 432.666667 | 322.666667 | 418.333333 | 499.333333 | 379.333333 | 314.5 | 355.5 | 333.166667 | 372.333333 | 339 | 368.833333 | 369 | 435 | 395.333333 | 381.071429 | 51.2044133 | 13.6849551 |
| 2 | 410.166667 | 280.166667 | 508.666667 | 481.166667 | 356 | 281.166667 | 357.5 | 318.666667 | 297.166667 | 428.5 | 392.666667 | 523 | 467.666667 | 308.5 | 386.5 | 85.1730516 | 22.7634555 |
| 3 | 478 | 292.666667 | 436 | 415.333333 | 326 | 298 | 319.666667 | 285 | 276.166667 | 421 | 337.833333 | 524.166667 | 419.333333 | 278 | 364.797619 | 82.1165312 | 21.9465661 |
| 4 | 424.833333 | 320.333333 | 558.166667 | 612.666667 | 345.5 | 306 | 347.666667 | 314 | 437.666667 | 398.333333 | 326.666667 | 510.666667 | 458.833333 | 283.666667 | 403.214286 | 101.921883 | 27.239769 |
| 5 | 447.666667 | 378 | 453.833333 | 452 | 349 | 306.5 | 290.833333 | 281 | 396.333333 | 382.666667 | 279.166667 | 495.833333 | 491.166667 | 286.666667 | 377.904762 | 80.4790336 | 21.5089265 |
| 10 | 307.766667 | 268.866667 | 423.3 | 509.5 | 336.033333 | 280.7 | 317.266667 | 280.3 | 404.166667 | 317.7 | 278.566667 | 412.4 | 287.133333 | 319.7 | 338.814286 | 71.3598438 | 19.0717205 |
| 15 | 359.733333 | 198.4 | 239.166667 | 315.633333 | 294.366667 | 266.066667 | 231 | 257.793103 | 227.533333 | 256.133333 | 279 | 370.666667 | 250.3 | 371.8 | 279.828079 | 55.5396426 | 14.8435939 |

### End inspiratory pause

Week 1

WT

| Time (min) | A | 156 | 158 | 161 | Average | SD | SE |
|---|---|---|---|---|---|---|---|
| 0 | 2.36666667 | 3.76666667 | 1.8 | 2.35555556 | 2.57222222 | 0.83909122 | 0.41954561 |
| 1 | 1.83333333 | 3 | 1.33333333 | 1.83333333 | 2 | 0.70710678 | 0.35355339 |
| 2 | 1.83333333 | 3.66666667 | 1 | 1.5 | 2 | 1.16269164 | 0.58134582 |
| 3 | 2 | 2.16666667 | 1.16666667 | 1.33333333 | 1.66666667 | 0.49065338 | 0.24532669 |
| 4 | 1.66666667 | 3.33333333 | 1.16666667 | 2 | 2.04166667 | 0.92671263 | 0.46335631 |
| 5 | 1.66666667 | 2.83333333 | 1.5 | 1.33333333 | 1.83333333 | 0.68041382 | 0.34020691 |
| 10 | 2.2 | 2.63333333 | 1.16666667 | 2.03333333 | 2.00833333 | 0.61546429 | 0.30773215 |
| 15 | 2 | 1.96666667 | 1.1 | 1.13333333 | 1.55 | 0.50074019 | 0.2503701 |

Het

| Time (min) | B | C | D | E | F | G | H | 155 | 157 | 159 | 160 | 162 | 190 | 196 | 199 | 201 | 205 | 207 | 208 | 210 | Average | SD | SE |
|---|---|---|---|---|---|---|---|---|---|---|---|---|---|---|---|---|---|---|---|---|---|---|---|
| 0 | 2.25555556 | 2.78888889 | 5.23333333 | 2.28888889 | 4.36666667 | 1.33333333 | 1.68888889 | 2.53333333 | 3.55555556 | 1.4 | 3.75555556 | 1.45555556 | 3.54444444 | 2.32222222 | 2.56666667 | 3.52222222 | 1.83333333 | 1.76666667 | 3.32222222 | 1.46666667 | 2.65 | 1.09069846 | 0.24388759 |
| 1 | 1.83333333 | 1.16666667 | 4.16666667 | 2.5 | 4.16666667 | 1.83333333 | 1.33333333 | 1.83333333 | 2.33333333 | 1 | 3 | 1.16666667 | 3 | 2 | 2.83333333 | 2.66666667 | 1 | 1.16666667 | 2.33333333 | 1 | 2.11666667 | 0.98244831 | 0.21968212 |
| 2 | 2.5 | 2 | 4.66666667 | 1.5 | 3.33333333 | 1.83333333 | 1 | 2 | 4.66666667 | 1 | 4 | 1 | 3.83333333 | 2.66666667 | 3.16666667 | 4.33333333 | 1 | 2.66666667 | 1.66666667 | 1 | 2.49166667 | 1.29492374 | 0.28955375 |
| 3 | 2.5 | 2.16666667 | 4.16666667 | 2 | 3.33333333 | 1.33333333 | 1.16666667 | 2.66666667 | 5 | 1.16666667 | 3.83333333 | 1.5 | 3.83333333 | 2.83333333 | 3.33333333 | 4 | 1 | 2.33333333 | 2.33333333 | 2.16666667 | 2.63333333 | 1.14017543 | 0.25495098 |
| 4 | 2 | 2.16666667 | 4.83333333 | 2 | 3.33333333 | 1.16666667 | 1.16666667 | 2.83333333 | 6 | 1.5 | 5.33333333 | 1.66666667 | 4.16666667 | 3 | 3.5 | 3.33333333 | 1 | 1.83333333 | 3 | 1.5 | 2.76666667 | 1.44428699 | 0.32295239 |
| 5 | 2.83333333 | 2.16666667 | 5.16666667 | 2.16666667 | 4 | 1 | 1.83333333 | 2.16666667 | 4.16666667 | 1 | 3.83333333 | 1 | 4 | 2 | 5 | 5.16666667 | 1 | 1.16666667 | 2.66666667 | 1.16666667 | 2.675 | 1.49802502 | 0.33496858 |
| 10 | 2.3 | 2.2 | 5.16666667 | 2.1 | 3.43333333 | 2.03333333 | 1.53333333 | 3.3 | 3.6 | 1.26666667 | 3 | 1.56666667 | 4 | 2.33333333 | 4.2 | 3.6 | 1.66666667 | 2.46666667 | 1.96666667 | 1.53333333 | 2.66333333 | 1.06808568 | 0.23883122 |
| 15 | 2.03333333 | 2.03333333 | 5.63333333 | 2 | 3.2 | 1.83333333 | 1.66666667 | 2.06666667 | 3.3 | 1 | 2.5 | 1.13333333 | 3.2 | 2.16666667 | 4 | 3.83333333 | 1.76666667 | 1.3 | 1.16666667 | 1 | 2.34166667 | 1.19672433 | 0.2675957 |

Week 2

WT

| Time (min) | A | 156 | 158 | 161 | Average | SD | SE |
|---|---|---|---|---|---|---|---|
| 0 | 2.36666667 | 2.07777778 | 2.02222222 | 1.8 | 2.06666667 | 0.23324513 | 0.11662257 |
| 1 | 2 | 1.33333333 | 1.16666667 | 2 | 1.625 | 0.43832594 | 0.21916297 |
| 2 | 2.66666667 | 1.83333333 | 1 | 1.5 | 1.75 | 0.7005289 | 0.35026445 |
| 3 | 1.83333333 | 1.66666667 | 1.66666667 | 1 | 1.54166667 | 0.3695593 | 0.18477965 |
| 4 | 2.16666667 | 2.5 | 2.33333333 | 1.33333333 | 2.08333333 | 0.51818773 | 0.25909386 |
| 5 | 2.33333333 | 2.33333333 | 2.16666667 | 1.66666667 | 2.125 | 0.31549491 | 0.15774745 |
| 10 | 2.3 | 2.36666667 | 1.16666667 | 1.13333333 | 1.74166667 | 0.68387512 | 0.34193756 |
| 15 | 2.3 | 2 | 1.23333333 | 1.06666667 | 1.65 | 0.59410686 | 0.29705343 |

Het

| Time (min) | B | D | E | G | H | 155 | 157 | 160 | 162 | 190 | 196 | 199 | 201 | 205 | 207 | 208 | 210 | Average | SD | SE |
|---|---|---|---|---|---|---|---|---|---|---|---|---|---|---|---|---|---|---|---|---|
| 0 | 1.87777778 | 2.91111111 | 2 | 1.34444444 | 1.44444444 | 1.96666667 | 2.54444444 | 3.75555556 | 4.5 | 2.78888889 | 1.61111111 | 2.2 | 2.94444444 | 1.38888889 | 2.03333333 | 2.07777778 | 1.32222222 | 2.27712418 | 0.88221864 | 0.21396945 |
| 1 | 2.66666667 | 1.83333333 | 1.5 | 1.16666667 | 1.16666667 | 1.66666667 | 2.66666667 | 3.83333333 | 1 | 2 | 2.2 | 2 | 1.4 | 1.2 | 1 | 1 | 1.2 | 1.73529412 | 0.77122473 | 0.18704947 |
| 2 | 3.83333333 | 2.83333333 | 1.66666667 | 1.5 | 1 | 2 | 3.33333333 | 3.16666667 | 1.33333333 | 3 | 2 | 3.33333333 | 2.16666667 | 1.16666667 | 1.16666667 | 1 | 1.66666667 | 2.12745098 | 0.93639641 | 0.22710949 |
| 3 | 3.5 | 2.83333333 | 1.83333333 | 1.16666667 | 1 | 2 | 2.66666667 | 3.33333333 | 1 | 3.16666667 | 2.16666667 | 3.5 | 2.83333333 | 1 | 1 | 1 | 1.33333333 | 2.07843137 | 0.98622042 | 0.23919359 |
| 4 | 3 | 2.66666667 | 1.83333333 | 1.5 | 1 | 1.66666667 | 3 | 4.16666667 | 1.16666667 | 2.66666667 | 3.5 | 3.33333333 | 2 | 1 | 1 | 1.33333333 | 1 | 2.10784314 | 1.02899151 | 0.2495671 |
| 5 | 3.33333333 | 2.66666667 | 1.66666667 | 1.66666667 | 1 | 2 | 2.83333333 | 5.66666667 | 1.16666667 | 3.83333333 | 3 | 3.5 | 2.16666667 | 1 | 1 | 1 | 1 | 2.26470588 | 1.316375 | 0.31926783 |
| 10 | 3.16666667 | 2.76666667 | 1.46666667 | 1.9 | 1.2 | 1.96666667 | 3.03333333 | 4.73333333 | 1.66666667 | 2.53333333 | 3.9 | 4.06666667 | 2.5 | 1 | 1.76666667 | 1 | 1.06666667 | 2.3372549 | 1.14405968 | 0.27747523 |
| 15 | 3.83333333 | 2.6 | 2.06666667 | 1.4 | 1.23333333 | 2 | 4.86666667 | 4.26666667 | 1.2 | 2.66666667 | 2.73333333 | 4.5 | 3.33333333 | 1.03333333 | 1.7 | 1.4 | 1.03333333 | 2.4627451 | 1.2834893 | 0.31129188 |

Week 3

WT

| Time (min) | 83 | 86 | 87 | 88 | 81 | 82 | A | 156 | 158 | 161 | Average | SD | SE |
|---|---|---|---|---|---|---|---|---|---|---|---|---|---|
| 0 | 4.63333333 | 2.54444444 | 2.51111111 | 2.51111111 | 1.83146067 | 2.02222222 | 2.26666667 | 2.98888889 | 1.58888889 | 1.27777778 | 2.41759051 | 0.92963816 | 0.2939774 |
| 1 | 2 | 1.83333333 | 1.5 | 1.83333333 | 1.16666667 | 2.33333333 | 1.5 | 2.2 | 1.83333333 | 2.33333333 | 1.85333333 | 0.38367939 | 0.12133008 |
| 2 | 3.5 | 2 | 1.66666667 | 1.5 | 1.83333333 | 3 | 1.5 | 3 | 1 | 1 | 2 | 0.87488976 | 0.27666444 |
| 3 | 2.33333333 | 2.66666667 | 2.16666667 | 1.66666667 | 2 | 4.16666667 | 2.83333333 | 4.33333333 | 1 | 1 | 2.41666667 | 1.14463321 | 0.3619648 |
| 4 | 2.33333333 | 3 | 2 | 1.66666667 | 1.83333333 | 3.66666667 | 1.66666667 | 2.5 | 1 | 1 | 2.06666667 | 0.83960602 | 0.26550674 |
| 5 | 2.66666667 | 3.33333333 | 2.33333333 | 1.66666667 | 2.16666667 | 2 | 1.83333333 | 2.83333333 | 1 | 1 | 2.08333333 | 0.75461543 | 0.23863035 |
| 10 | 2.3 | 2.46666667 | 2.46666667 | 2.56666667 | 1.8 | 3.1 | 2.1 | 2.7 | 1.03333333 | 1.33333333 | 2.18666667 | 0.63483253 | 0.20075167 |
| 15 | 2.96666667 | 2.93333333 | 2.16666667 | 2 | 1.86666667 | 2.9 | 1.96666667 | 2.4 | 1.1 | 1.1 | 2.14 | 0.68687879 | 0.21721014 |

Het

| Time (min) | B | D | E | G | H | 155 | 157 | 190 | 196 | 199 | 201 | 205 | 207 | 208 | 210 | Average | SD | SE |
|---|---|---|---|---|---|---|---|---|---|---|---|---|---|---|---|---|---|---|
| 0 | 1.62222222 | 2.87777778 | 1.3 | 5.51111111 | 1.56666667 | 2.54444444 | 2.16666667 | 2.06666667 | 1.96666667 | 3.28888889 | 2.21111111 | 1.1 | 1.18888889 | 1.04444444 | 1.34444444 | 2.12 | 1.15106116 | 0.29720271 |
| 1 | 1.66666667 | 1.5 | 1.16666667 | 2.83333333 | 1.16666667 | 1.2 | 1.66666667 | 1.5 | 1.5 | 2.16666667 | 1.83333333 | 1 | 1 | 1 | 1.5 | 1.51333333 | 0.4963038 | 0.12814509 |
| 2 | 2.5 | 1.83333333 | 1 | 1.66666667 | 1.5 | 1.5 | 2.33333333 | 2.16666667 | 2.33333333 | 3.16666667 | 3.5 | 1 | 1.16666667 | 1 | 1.33333333 | 1.86666667 | 0.78477577 | 0.20262823 |
| 3 | 2.16666667 | 2 | 1.33333333 | 2.66666667 | 1.33333333 | 1.83333333 | 2.33333333 | 2 | 2.33333333 | 3 | 2.66666667 | 1.5 | 1 | 1 | 1 | 1.87777778 | 0.65908386 | 0.17017472 |
| 4 | 2.33333333 | 2.33333333 | 1.16666667 | 1.16666667 | 1.16666667 | 2 | 2.16666667 | 2 | 2.16666667 | 4.33333333 | 3.33333333 | 1.16666667 | 1.33333333 | 1 | 1 | 1.91111111 | 0.94882272 | 0.24498497 |
| 5 | 2.83333333 | 2.16666667 | 1 | 1.16666667 | 1.66666667 | 2 | 3.16666667 | 2.66666667 | 2.83333333 | 4.33333333 | 3.66666667 | 1.16666667 | 1 | 1 | 1 | 2.11111111 | 1.09411606 | 0.28249955 |
| 10 | 2.2 | 2.6 | 1 | 3.76666667 | 1.43333333 | 1.96666667 | 3.1 | 2.26666667 | 3.03333333 | 3.43333333 | 3.6 | 1 | 1.1 | 1.9 | 1.03333333 | 2.22888889 | 0.99194107 | 0.25611808 |
| 15 | 1.96666667 | 2 | 1.13333333 | 3.63333333 | 1.46666667 | 2.23333333 | 3.4 | 2.5 | 3.1 | 4.4 | 2.36666667 | 1.1 | 1.03333333 | 1.46666667 | 1.13333333 | 2.19555556 | 1.04470446 | 0.26974153 |

Week 4

WT

| Time (min) | A | 156 | 158 | 161 | Average | SD | SE |
|---|---|---|---|---|---|---|---|
| 0 | 2.46666667 | 2.9 | 1.9 | 1.26666667 | 2.13333333 | 0.70815357 | 0.35407679 |
| 1 | 1.5 | 2.66666667 | 1.16666667 | 1.16666667 | 1.625 | 0.71200031 | 0.35600016 |
| 2 | 2.5 | 2.66666667 | 1 | 1.16666667 | 1.83333333 | 0.87135484 | 0.43567742 |
| 3 | 2.5 | 3.16666667 | 1.16666667 | 2 | 2.20833333 | 0.84299949 | 0.42149975 |
| 4 | 2.33333333 | 2.33333333 | 1.33333333 | 1 | 1.75 | 0.68718427 | 0.34359214 |
| 5 | 2.33333333 | 2.33333333 | 1 | 1.33333333 | 1.75 | 0.68718427 | 0.34359214 |
| 10 | 2.03333333 | 3.63333333 | 1.23333333 | 1.13333333 | 2.00833333 | 1.15578256 | 0.57789128 |
| 15 | 2 | 2.36666667 | 1.43333333 | 1.03333333 | 1.70833333 | 0.59152972 | 0.29576486 |

Het

| Time (min) | B | D | E | H | 155 | 157 | 190 | 196 | 199 | 201 | 205 | 207 | 208 | 210 | Average | SD | SE |
|---|---|---|---|---|---|---|---|---|---|---|---|---|---|---|---|---|---|
| 0 | 1.01111111 | 2.07777778 | 1.37777778 | 1.44444444 | 1.75555556 | 2.27777778 | 1.96666667 | 2.01111111 | 2.41111111 | 2.62222222 | 1.13333333 | 1.23333333 | 1.4 | 1.2 | 1.70873016 | 0.52074128 | 0.13917396 |
| 1 | 1.66666667 | 3.16666667 | 1.33333333 | 1.33333333 | 1.16666667 | 1.66666667 | 1.83333333 | 1.5 | 2.33333333 | 2 | 1 | 1.66666667 | 1.33333333 | 1 | 1.64285714 | 0.57682132 | 0.15416198 |
| 2 | 2 | 2.83333333 | 1.16666667 | 1.33333333 | 2 | 2 | 2 | 1.66666667 | 2.83333333 | 2.66666667 | 1 | 1.16666667 | 1 | 1 | 1.76190476 | 0.67846699 | 0.18132793 |
| 3 | 1.66666667 | 2.83333333 | 1.16666667 | 2.5 | 1.33333333 | 2 | 2 | 2.33333333 | 3.66666667 | 3.83333333 | 1 | 1.16666667 | 1.16666667 | 1 | 1.97619048 | 0.95375359 | 0.25490137 |
| 4 | 2 | 3.5 | 1 | 1 | 1.83333333 | 1.83333333 | 2.83333333 | 2 | 3.5 | 3.33333333 | 1 | 1.33333333 | 1.5 | 1 | 1.97619048 | 0.94926229 | 0.25370102 |
| 5 | 1.5 | 3 | 1.16666667 | 1.33333333 | 2 | 1.5 | 2 | 2.5 | 4.16666667 | 4.16666667 | 1 | 1.33333333 | 1 | 1 | 1.97619048 | 1.09945041 | 0.29384048 |
| 10 | 1.83333333 | 5.1 | 1.03333333 | 1.13333333 | 1.83333333 | 1.8 | 2.5 | 2.26666667 | 3.8 | 3.46666667 | 1 | 1.73333333 | 1.26666667 | 1.03333333 | 2.12857143 | 1.22208013 | 0.32661465 |
| 15 | 2.33333333 | 3 | 1.13333333 | 1.1 | 1.86666667 | 2 | 2 | 2.5862069 | 3.33333333 | 2.83333333 | 1.36666667 | 1.73333333 | 1.03333333 | 1.16666667 | 1.96330049 | 0.76400342 | 0.2041885 |

### End expiratory pause

Week 1

WT

| Time (min) | A | 156 | 158 | 161 | Average | SD | SE |
|---|---|---|---|---|---|---|---|
| 0 | 10.2968889 | 8.31855556 | 14.9114444 | 11.295 | 11.2054722 | 2.76297285 | 1.38148642 |
| 1 | 10.5083333 | 8.38833333 | 9.94833333 | 9.115 | 9.49 | 0.93119398 | 0.46559699 |
| 2 | 8.135 | 8.76166667 | 9.18333333 | 8.78166667 | 8.71541667 | 0.4329579 | 0.21647895 |
| 3 | 9.04 | 8.17 | 12.355 | 9.34 | 9.72625 | 1.82138855 | 0.91069427 |
| 4 | 10.91 | 9.27666667 | 10.2566667 | 10.5483333 | 10.2479167 | 0.70047785 | 0.35023892 |
| 5 | 13.2566667 | 10.4133333 | 10.4033333 | 10.8816667 | 11.23875 | 1.36366294 | 0.68183147 |
| 10 | 11.24 | 11.749 | 15.0826667 | 11.1406667 | 12.3030833 | 1.87211561 | 0.9360578 |
| 15 | 18.854 | 17.2593333 | 18.7473333 | 18.0173333 | 18.2195 | 0.74026789 | 0.37013395 |

Het

| Time (min) | B | C | D | E | F | G | H | 155 | 157 | 159 | 160 | 162 | 190 | 196 | 199 | 201 | 205 | 207 | 208 | 210 | Average | SD | SE |
|---|---|---|---|---|---|---|---|---|---|---|---|---|---|---|---|---|---|---|---|---|---|---|---|
| 0 | 21.6756667 | 10.5912222 | 10.3801111 | 12.7948889 | 18.1494444 | 11.9815556 | 9.13122222 | 10.0273333 | 7.27177778 | 14.0454444 | 9.80488889 | 20.3114444 | 11.2724444 | 18.9358889 | 24.5125556 | 7.54877778 | 10.4478889 | 14.683 | 13.2142222 | 21.7675556 | 13.9273667 | 5.16282467 | 1.15444269 |
| 1 | 8.01333333 | 9.23666667 | 8.55333333 | 8.94666667 | 18.4966667 | 8.27833333 | 8.59166667 | 8.58666667 | 10.6066667 | 10.0266667 | 8.13666667 | 13.8183333 | 8.555 | 9.39666667 | 10.5716667 | 11.205 | 11.535 | 9.80333333 | 9.45 | 9.96166667 | 10.0885 | 2.42859933 | 0.54305132 |
| 2 | 9.265 | 9.195 | 9.62 | 9.76333333 | 22.1083333 | 8.01166667 | 8.555 | 11.075 | 12.3533333 | 11.69 | 8.61666667 | 11.145 | 6.73666667 | 10.31 | 9.44833333 | 8.99833333 | 16.2566667 | 9.94166667 | 9.615 | 12.5516667 | 10.7628333 | 3.3485249 | 0.74875293 |
| 3 | 10.7783333 | 9.40333333 | 10.7866667 | 8.665 | 21.675 | 7.97666667 | 8.28833333 | 9.86 | 11.2166667 | 10.7733333 | 9.48166667 | 10.06 | 6.55333333 | 11.3883333 | 12.715 | 16.6416667 | 13.95 | 10.4316667 | 11.4883333 | 11.675 | 11.1904167 | 3.29928288 | 0.73774208 |
| 4 | 11.35 | 8.76833333 | 11.0316667 | 9.76166667 | 22.495 | 8.22 | 10.76 | 10.555 | 11.4416667 | 10.3416667 | 10.8616667 | 11.27 | 6.08 | 11.56 | 7.96166667 | 12.7166667 | 14.42 | 8.76666667 | 11.01 | 10.9816667 | 11.0176667 | 3.25134508 | 0.72702286 |
| 5 | 13.1583333 | 9.25 | 10.9033333 | 13.69 | 26.9666667 | 7.87666667 | 11.21 | 11.16 | 11.19 | 11.6116667 | 11.07 | 13.885 | 12.985 | 15.9033333 | 8.46833333 | 11.3133333 | 14.9816667 | 9.23333333 | 10.9333333 | 15.9183333 | 12.5854167 | 4.07723325 | 0.91169707 |
| 10 | 24.513 | 11.4673333 | 17.686 | 26.844 | 25.0443333 | 8.57933333 | 11.237 | 13.4793333 | 23.4196667 | 16.4156667 | 20.241 | 15.9696667 | 12.4356667 | 22.254 | 11.9853333 | 11.1836667 | 13.9173333 | 14.832 | 20.868 | 20.8383333 | 17.1605333 | 5.47516911 | 1.22428503 |
| 15 | 21.0576667 | 16.8773333 | 14.68 | 24.6423333 | 24.687 | 12.381 | 13.658 | 16.1593333 | 23.827 | 19.8003333 | 25.718 | 26.0706667 | 19.3866667 | 22.548 | 32.1036667 | 20.295 | 14.932 | 23.485 | 19.8106667 | 23.1966667 | 20.7658167 | 4.97232137 | 1.11184486 |

Week 2

WT

| Time (min) | A | 156 | 158 | 161 | Average | SD | SE |
|---|---|---|---|---|---|---|---|
| 0 | 14.4 | 16.1612222 | 11.0577778 | 9.97922222 | 12.8995556 | 2.87577787 | 1.43788894 |
| 1 | 7.79166667 | 8.69166667 | 8.44833333 | 8.38333333 | 8.32875 | 0.38185797 | 0.19092899 |
| 2 | 8.165 | 8.72833333 | 8.77 | 10.705 | 9.09208333 | 1.11011042 | 0.55505521 |
| 3 | 8.07333333 | 7.55666667 | 10.5083333 | 11.6416667 | 9.445 | 1.94964052 | 0.97482026 |
| 4 | 9.57833333 | 9.78 | 9.87166667 | 10.9433333 | 10.0433333 | 0.61238302 | 0.30619151 |
| 5 | 12.8916667 | 9.16333333 | 10.93 | 10.815 | 10.95 | 1.52543679 | 0.7627184 |
| 10 | 12.351 | 10.823 | 17.9223333 | 13.8166667 | 13.72825 | 3.05152681 | 1.52576341 |
| 15 | 21.1886667 | 14.537 | 19.92 | 17.6526667 | 18.3245833 | 2.91808083 | 1.45904042 |

Het

| Time (min) | B | D | E | G | H | 155 | 157 | 160 | 162 | 190 | 196 | 199 | 201 | 205 | 207 | 208 | 210 | Average | SD | SE |
|---|---|---|---|---|---|---|---|---|---|---|---|---|---|---|---|---|---|---|---|---|
| 0 | 20.5096667 | 13.5897778 | 19.846 | 14.4478889 | 10.2637778 | 17.3214444 | 19.4258889 | 8.80422222 | 11.9522222 | 12.1343333 | 19.8967778 | 22.3172222 | 18.0407778 | 7.73655556 | 10.5106667 | 8.303 | 8.41555556 | 14.3244575 | 4.99722817 | 1.21200586 |
| 1 | 8.06833333 | 8.47166667 | 11.4283333 | 8.72666667 | 7.79833333 | 10.03 | 10.26 | 7.56333333 | 11.31 | 12.048 | 8.994 | 15.322 | 16.164 | 15.872 | 14.022 | 13.29 | 10.056 | 11.1426275 | 2.88435218 | 0.69955816 |
| 2 | 6.89 | 6.605 | 9.50833333 | 8.625 | 8.63166667 | 8.14833333 | 8.965 | 9.32666667 | 12.3716667 | 7.955 | 7.53 | 9.71 | 10.995 | 16.6983333 | 9.39833333 | 10.2383333 | 11.8733333 | 9.61588235 | 2.41642184 | 0.58606838 |
| 3 | 7.49333333 | 7.54 | 9.49666667 | 8.31833333 | 8.59 | 10.8266667 | 10.6266667 | 7.23 | 12.5266667 | 9.06833333 | 8.83166667 | 10.2466667 | 9.41 | 10.9383333 | 11.7266667 | 11.7016667 | 10.9333333 | 9.73558824 | 1.61282953 | 0.39116862 |
| 4 | 8.54166667 | 6.96166667 | 8.87666667 | 9.705 | 10.485 | 8.95166667 | 10.7666667 | 9.935 | 9.62166667 | 11.4716667 | 8.79833333 | 9.31166667 | 13.3033333 | 12.995 | 17.8283333 | 12.365 | 16.2983333 | 10.9539216 | 2.84843951 | 0.69084806 |
| 5 | 10.7283333 | 8.54166667 | 15.6083333 | 8.08333333 | 12.7083333 | 10.11 | 11.8883333 | 11.105 | 11.31 | 8.1 | 10.0933333 | 6.94 | 16.4566667 | 15.0916667 | 16.1383333 | 14.0033333 | 16.7533333 | 11.98 | 3.2103034 | 0.77861294 |
| 10 | 13.128 | 11.3686667 | 22.191 | 10.0096667 | 14.2296667 | 10.3443333 | 14.0606667 | 17.7883333 | 11.736 | 16.0626667 | 9.34766667 | 10.8963333 | 13.9203333 | 15.1803333 | 14.1696667 | 14.196 | 17.4703333 | 13.8882157 | 3.28277798 | 0.79619061 |
| 15 | 13.4883333 | 20.7603333 | 30.8173333 | 11.2336667 | 13.7856667 | 14.9723333 | 10.557 | 24.426 | 24.1256667 | 22.986 | 13.6066667 | 16.0783333 | 9.85966667 | 19.05 | 15.1123333 | 13.9333333 | 26.485 | 17.7222157 | 6.17437497 | 1.49750589 |

Week 3

WT

| Time (min) | 83 | 86 | 87 | 88 | 81 | 82 | A | 156 | 158 | 161 | Average | SD | SE |
|---|---|---|---|---|---|---|---|---|---|---|---|---|---|
| 0 | 8.81677778 | 19.7367778 | 13.6202222 | 9.71677778 | 21.8046067 | 17.2117778 | 10.2585556 | 15.4773333 | 10.9751111 | 14.7781111 | 14.2396051 | 4.40777422 | 1.3938606 |
| 1 | 10.1583333 | 7.70833333 | 6.89 | 6.65833333 | 7.73666667 | 6.26333333 | 6.37 | 6.916 | 8.30166667 | 8.63666667 | 7.56393333 | 1.21507929 | 0.38424181 |
| 2 | 11.6966667 | 6.63333333 | 7.135 | 8.08166667 | 8.665 | 7.09 | 7.41833333 | 6.72833333 | 9.58833333 | 9.80666667 | 8.28433333 | 1.64965339 | 0.52166621 |
| 3 | 10.535 | 7.23 | 8.75166667 | 9.02833333 | 10.7883333 | 8.87666667 | 9.48166667 | 8.035 | 9.38166667 | 10.2833333 | 9.23916667 | 1.11299292 | 0.35195926 |
| 4 | 10.0616667 | 7.86666667 | 8.545 | 8.21333333 | 9.73 | 8.725 | 6.41666667 | 6.88333333 | 10.3366667 | 11.5033333 | 8.82816667 | 1.59017653 | 0.50285797 |
| 5 | 13.2416667 | 9.36666667 | 7.99666667 | 7.65833333 | 8.95333333 | 6.975 | 10.6266667 | 7.34166667 | 12.5583333 | 10.6566667 | 9.5375 | 2.18451781 | 0.69080519 |
| 10 | 13.2246667 | 9.05166667 | 11.308 | 11.1236667 | 8.89533333 | 9.57 | 10.1093333 | 9.91566667 | 13.1753333 | 10.754 | 10.7127667 | 1.53718541 | 0.48610071 |
| 15 | 11.7376667 | 16.1773333 | 22.4546667 | 24.4516667 | 19.169 | 13.7686667 | 16.617 | 20.537 | 16.4493333 | 12.912 | 17.4274333 | 4.17187377 | 1.31926232 |

Het

| Time (min) | B | D | E | G | H | 155 | 157 | 190 | 196 | 199 | 201 | 205 | 207 | 208 | 210 | Average | SD | SE |
|---|---|---|---|---|---|---|---|---|---|---|---|---|---|---|---|---|---|---|
| 0 | 16.6923333 | 13.5627778 | 18.3403333 | 19.3104444 | 12.2995556 | 7.99077778 | 19.2483333 | 16.9542222 | 19.0557778 | 21.4497778 | 20.3895556 | 16.822 | 14.6691111 | 15.942 | 16.743 | 16.6313333 | 3.45882317 | 0.8930643 |
| 1 | 8.04666667 | 12.81 | 9.12833333 | 14.6366667 | 8.465 | 8.414 | 7.21333333 | 10.4933333 | 9.65 | 11.7366667 | 12.6833333 | 12.29 | 13.1216667 | 12.6016667 | 10.375 | 10.7777111 | 2.23630865 | 0.57741241 |
| 2 | 7.06666667 | 7.685 | 7.99 | 15.3 | 8.14333333 | 9.57166667 | 8.75166667 | 8.77166667 | 9.82 | 10.215 | 8.21 | 17.1666667 | 8.55666667 | 13.6933333 | 12.19 | 10.2087778 | 3.01545572 | 0.77858732 |
| 3 | 9.295 | 7.30333333 | 8.025 | 14.9883333 | 8.14833333 | 9.31333333 | 8.80833333 | 9.725 | 10.1316667 | 11.5533333 | 8.48833333 | 15.8983333 | 11.725 | 13.3666667 | 10.2566667 | 10.4684444 | 2.57097817 | 0.66382371 |
| 4 | 6.52333333 | 11.385 | 9.62333333 | 20.15 | 8.29 | 9.50833333 | 9.08666667 | 11.9633333 | 11.5216667 | 11.095 | 9.145 | 15.52 | 11.6983333 | 12.9116667 | 8.87833333 | 11.1533333 | 3.29943754 | 0.85191111 |
| 5 | 7.61166667 | 14.3183333 | 7.98666667 | 16.06 | 9.175 | 10.3766667 | 9.315 | 11.1116667 | 10.0066667 | 10.8316667 | 8.91166667 | 14.385 | 13.3066667 | 16.6233333 | 14.19 | 11.614 | 2.95254053 | 0.76234269 |
| 10 | 14.044 | 15.628 | 19.362 | 16.1583333 | 10.6866667 | 10.675 | 10.144 | 10.2486667 | 10.9493333 | 8.09766667 | 12.128 | 11.2923333 | 15.8006667 | 11.4926667 | 14.47 | 12.7451556 | 3.02545461 | 0.78116902 |
| 15 | 13.0036667 | 21.4523333 | 16.761 | 19.067 | 10.3196667 | 10.912 | 9.11433333 | 10.0916667 | 10.22 | 23.6713333 | 21.567 | 13.5726667 | 16.0536667 | 15.2323333 | 15.3156667 | 15.0902889 | 4.68244733 | 1.2090027 |

Week 4

WT

| Time (min) | A | 156 | 158 | 161 | Average | SD | SE |
|---|---|---|---|---|---|---|---|
| 0 | 12.8831111 | 12.7738889 | 17.232 | 8.26911111 | 12.7895278 | 3.65963244 | 1.82981622 |
| 1 | 6.455 | 7.90833333 | 10.0366667 | 9.21 | 8.4025 | 1.56626029 | 0.78313015 |
| 2 | 8.23333333 | 8.46166667 | 8.23166667 | 10.11 | 8.75916667 | 0.90701229 | 0.45350615 |
| 3 | 8.37666667 | 7.445 | 12.3233333 | 11.3583333 | 9.87583333 | 2.33413181 | 1.1670659 |
| 4 | 7.69666667 | 7.49 | 11.3166667 | 10.8233333 | 9.33166667 | 2.01909679 | 1.0095484 |
| 5 | 8.935 | 9.095 | 7.88833333 | 10.2733333 | 9.04791667 | 0.97660066 | 0.48830033 |
| 10 | 14.2556667 | 13.426 | 10.857 | 11.895 | 12.6084167 | 1.52296846 | 0.76148423 |
| 15 | 20.5786667 | 15.9486667 | 14.42 | 11.8616667 | 15.70225 | 3.66216721 | 1.83108361 |

Het

| Time (min) | B | D | E | H | 155 | 157 | 190 | 196 | 199 | 201 | 205 | 207 | 208 | 210 | Average | SD |
|---|---|---|---|---|---|---|---|---|---|---|---|---|---|---|---|---|
| 0 | 15.5391111 | 31.9524444 | 15.884 | 15.0017778 | 15.7041111 | 19.8854444 | 17.0596667 | 19.0845556 | 21.7445556 | 17.2015556 | 14.9416667 | 16.7426667 | 9.18255556 | 16.6243333 | 17.6106032 | 5.02448186 |
| 1 | 8.55833333 | 11.735 | 9.40333333 | 8.91333333 | 10.19 | 12.0133333 | 10.3783333 | 10.6783333 | 9.83833333 | 10.5083333 | 9.93833333 | 10.7166667 | 8.68166667 | 9.53 | 10.077381 | 1.03757828 |
| 2 | 9.02833333 | 14.17 | 8.72333333 | 9.48 | 10.5616667 | 11.915 | 10.64 | 13.525 | 13.3216667 | 7.34166667 | 8.78166667 | 8.01 | 7.58666667 | 9.29333333 | 10.169881 | 2.26198342 |
| 3 | 9.15333333 | 14.0083333 | 8.67166667 | 10.7366667 | 13.195 | 11.7733333 | 10.8933333 | 10.5516667 | 14.6566667 | 7.615 | 9.61833333 | 7.93666667 | 8.565 | 10.0133333 | 10.5277381 | 2.20980204 |
| 4 | 8.80833333 | 12.88 | 7.82333333 | 7.425 | 10.65 | 12.2266667 | 11.9316667 | 12.02 | 9.93333333 | 9.24166667 | 9.76166667 | 8.92833333 | 8.185 | 11.285 | 10.0785714 | 1.77415333 |
| 5 | 9.27833333 | 10.2416667 | 9.15333333 | 11.745 | 10.6583333 | 12.14 | 12.9316667 | 11.2666667 | 9.26 | 8.93666667 | 11.5866667 | 8.92166667 | 7.97666667 | 13.02 | 10.5083333 | 1.6292618 |
| 10 | 13.3896667 | 17.908 | 11.5653333 | 9.81933333 | 11.335 | 13.0506667 | 14.4873333 | 12.349 | 11.8893333 | 13.1006667 | 11.8573333 | 10.6186667 | 13.8823333 | 12.1456667 | 12.6713095 | 1.96198467 |
| 15 | 12.6323333 | 25.3443333 | 18.3856667 | 16.1306667 | 13.981 | 14.1833333 | 22.9286667 | 13.8427586 | 20.9343333 | 17.7633333 | 12.5543333 | 11.5353333 | 17.56 | 10.4896667 | 16.304697 | 4.43034502 |

### Enhanced pause

Week 1

WT

| Time (min) | A | | 158 | 161 | Average | SD | SE |
|---|---|---|---|---|---|---|---|
| 0 | 0.29277778 | 0.26577778 | 0.30944444 | 0.24844444 | 0.27911111 | 0.02723424 | 0.01361712 |
| 1 | 0.46833333 | 0.29666667 | 0.26333333 | 0.25166667 | 0.32 | 0.10071044 | 0.05035522 |
| 2 | 0.305 | 0.27666667 | 0.32 | 0.35333333 | 0.31375 | 0.03192511 | 0.01596256 |
| 3 | 0.34 | 0.28333333 | 0.31833333 | 0.33333333 | 0.31875 | 0.02528998 | 0.01264499 |
| 4 | 0.36833333 | 0.295 | 0.29 | 0.32166667 | 0.31875 | 0.03585916 | 0.01792958 |
| 5 | 0.45666667 | 0.35666667 | 0.25333333 | 0.37666667 | 0.36083333 | 0.0836826 | 0.0418413 |
| 10 | 0.34266667 | 0.32333333 | 0.248 | 0.31966667 | 0.30841667 | 0.04152231 | 0.02076115 |
| 15 | 0.48633333 | 0.40833333 | 0.33233333 | 0.43533333 | 0.41558333 | 0.06423589 | 0.03211795 |

Het

| Time (min) | B | C | D | E | F | G | H | 155 | 157 | 159 | 160 | 162 | 190 | 196 | 199 | 201 | 205 | 207 | 208 | 210 | Average | SD | SE |
|---|---|---|---|---|---|---|---|---|---|---|---|---|---|---|---|---|---|---|---|---|---|---|---|
| 0 | 0.53588889 | 0.30177778 | 0.26 | 0.33833333 | 0.50811111 | 0.39377778 | 0.25022222 | 0.35511111 | 0.28344444 | 0.38155556 | 0.29388889 | 0.47066667 | 0.35577778 | 0.50488889 | 0.53466667 | 0.28233333 | 0.31522222 | 0.32233333 | 0.28611111 | 0.58366667 | 0.37788889 | 0.10586818 | 0.02367284 |
| 1 | 0.28166667 | 0.38833333 | 0.275 | 0.27666667 | 0.92833333 | 0.30666667 | 0.31833333 | 0.365 | 0.36833333 | 0.375 | 0.29833333 | 0.36 | 0.32666667 | 0.34 | 0.30666667 | 0.31833333 | 0.43333333 | 0.33333333 | 0.29166667 | 0.31 | 0.36008333 | 0.13995559 | 0.03129502 |
| 2 | 0.26166667 | 0.37833333 | 0.31 | 0.32333333 | 0.77833333 | 0.30166667 | 0.38166667 | 0.32166667 | 0.20333333 | 0.33666667 | 0.28833333 | 0.32 | 0.25833333 | 0.34166667 | 0.26666667 | 0.27333333 | 0.37 | 0.34666667 | 0.30833333 | 0.38666667 | 0.33783333 | 0.11395867 | 0.02548193 |
| 3 | 0.36 | 0.36 | 0.31666667 | 0.32166667 | 0.595 | 0.305 | 0.29833333 | 0.22833333 | 0.24 | 0.36666667 | 0.28166667 | 0.26333333 | 0.25666667 | 0.31666667 | 0.31833333 | 0.38 | 0.38833333 | 0.32333333 | 0.35666667 | 0.38333333 | 0.333 | 0.07795935 | 0.01743224 |
| 4 | 0.33833333 | 0.33666667 | 0.315 | 0.32 | 0.63666667 | 0.29833333 | 0.29 | 0.24 | 0.265 | 0.34833333 | 0.27 | 0.38333333 | 0.24666667 | 0.27166667 | 0.25 | 0.32666667 | 0.39666667 | 0.28 | 0.33833333 | 0.375 | 0.32633333 | 0.0863652 | 0.01931185 |
| 5 | 0.41 | 0.33166667 | 0.285 | 0.33166667 | 0.72833333 | 0.33 | 0.24666667 | 0.29 | 0.27333333 | 0.42333333 | 0.32 | 0.315 | 0.28833333 | 0.37666667 | 0.24166667 | 0.25666667 | 0.36 | 0.26166667 | 0.315 | 0.42166667 | 0.34033333 | 0.10688109 | 0.02389934 |
| 10 | 0.58166667 | 0.36566667 | 0.306 | 0.60066667 | 0.693 | 0.28633333 | 0.22366667 | 0.33433333 | 0.45933333 | 0.36233333 | 0.472 | 0.36466667 | 0.28566667 | 0.49133333 | 0.267 | 0.27566667 | 0.33766667 | 0.28333333 | 0.43033333 | 0.49766667 | 0.39591667 | 0.12794214 | 0.02860873 |
| 15 | 0.53 | 0.52433333 | 0.28433333 | 0.61933333 | 0.727 | 0.336 | 0.26433333 | 0.50033333 | 0.478 | 0.36833333 | 0.62533333 | 0.52266667 | 0.427 | 0.44933333 | 0.43133333 | 0.345 | 0.253 | 0.44933333 | 0.62166667 | 0.62433333 | 0.46905 | 0.13407907 | 0.02998099 |

Week 2

WT

| Time (min) | A | 156 | 158 | 161 | Average | SD | SE |
|---|---|---|---|---|---|---|---|
| 0 | 0.32533333 | 0.35711111 | 0.30177778 | 0.299 | 0.32080556 | 0.02693283 | 0.01346642 |
| 1 | 0.33166667 | 0.35666667 | 0.33 | 0.40833333 | 0.35666667 | 0.03654019 | 0.01827009 |
| 2 | 0.3 | 0.27 | 0.29333333 | 0.36166667 | 0.30625 | 0.03911936 | 0.01955968 |
| 3 | 0.29666667 | 0.21833333 | 0.37333333 | 0.49166667 | 0.345 | 0.11646808 | 0.05823404 |
| 4 | 0.315 | 0.2 | 0.42666667 | 0.385 | 0.33166667 | 0.09913515 | 0.04956757 |
| 5 | 0.365 | 0.19333333 | 0.44833333 | 0.42 | 0.35666667 | 0.11425281 | 0.0571264 |
| 10 | 0.32633333 | 0.25033333 | 0.27833333 | 0.29233333 | 0.28683333 | 0.03159641 | 0.01579821 |
| 15 | 0.397 | 0.26233333 | 0.277 | 0.32133333 | 0.31441667 | 0.06049939 | 0.03024969 |

Het

| Time (min) | B | D | E | G | H | 155 | 157 | 160 | 162 | 190 | 196 | 199 | 201 | 205 | 207 | 208 | 210 | Average | SD | SE |
|---|---|---|---|---|---|---|---|---|---|---|---|---|---|---|---|---|---|---|---|---|
| 0 | 0.43855556 | 0.36022222 | 0.326 | 0.44177778 | 0.26133333 | 0.49522222 | 0.40866667 | 0.28566667 | 0.22 | 0.36366667 | 0.492 | 0.42422222 | 0.39377778 | 0.31444444 | 0.28222222 | 0.29322222 | 0.29266667 | 0.35845098 | 0.08265459 | 0.02004668 |
| 1 | 0.27333333 | 0.355 | 0.31 | 0.27833333 | 0.305 | 0.39166667 | 0.285 | 0.31166667 | 0.415 | 0.592 | 0.34 | 0.308 | 0.614 | 0.906 | 0.724 | 0.748 | 0.656 | 0.45958824 | 0.20207247 | 0.04900977 |
| 2 | 0.25 | 0.31166667 | 0.275 | 0.29 | 0.27666667 | 0.28166667 | 0.22833333 | 0.30333333 | 0.23833333 | 0.30166667 | 0.22333333 | 0.28166667 | 0.335 | 0.66666667 | 0.195 | 0.34833333 | 0.38666667 | 0.3054902 | 0.10471317 | 0.02539668 |
| 3 | 0.22666667 | 0.315 | 0.26833333 | 0.27 | 0.33833333 | 0.25166667 | 0.22833333 | 0.25166667 | 0.35666667 | 0.25333333 | 0.23833333 | 0.26166667 | 0.22166667 | 0.315 | 0.21 | 0.18333333 | 0.335 | 0.26617647 | 0.04955624 | 0.01201915 |
| 4 | 0.26333333 | 0.31833333 | 0.24166667 | 0.34333333 | 0.21666667 | 0.23166667 | 0.22 | 0.28833333 | 0.23666667 | 0.31166667 | 0.18833333 | 0.24166667 | 0.24333333 | 0.325 | 0.38666667 | 0.16 | 0.31 | 0.26627451 | 0.05960381 | 0.01445605 |
| 5 | 0.22833333 | 0.34 | 0.255 | 0.275 | 0.24333333 | 0.215 | 0.23333333 | 0.28666667 | 0.27666667 | 0.25 | 0.15833333 | 0.265 | 0.315 | 0.37166667 | 0.36 | 0.28333333 | 0.33833333 | 0.27617647 | 0.05581394 | 0.01353687 |
| 10 | 0.24233333 | 0.31433333 | 0.28833333 | 0.29366667 | 0.25033333 | 0.235 | 0.24933333 | 0.30066667 | 0.30333333 | 0.40033333 | 0.20733333 | 0.287 | 0.30266667 | 0.34766667 | 0.31366667 | 0.29833333 | 0.34733333 | 0.29303922 | 0.04722683 | 0.01145419 |
| 15 | 0.253 | 0.466 | 0.27466667 | 0.316 | 0.231 | 0.239 | 0.223 | 0.337 | 0.23566667 | 0.46966667 | 0.265 | 0.30733333 | 0.25166667 | 0.398 | 0.25766667 | 0.30366667 | 0.581 | 0.31819608 | 0.10238564 | 0.02483217 |

Week 3

WT

| Time (min) | 83 | 86 | 87 | 88 | 81 | 82 | A | 156 | 158 | 161 | Average | SD | SE |
|---|---|---|---|---|---|---|---|---|---|---|---|---|---|
| 0 | 0.24777778 | 0.32122222 | 0.31211111 | 0.306 | 0.40213483 | 0.35677778 | 0.27122222 | 0.28011111 | 0.35122222 | 0.31244444 | 0.31610237 | 0.04524613 | 0.01430808 |
| 1 | 0.38166667 | 0.31333333 | 0.33 | 0.335 | 0.29 | 0.30333333 | 0.325 | 0.288 | 0.395 | 0.41666667 | 0.3378 | 0.04496918 | 0.0142205 |
| 2 | 0.335 | 0.27166667 | 0.26166667 | 0.30333333 | 0.23666667 | 0.28333333 | 0.27 | 0.26 | 0.34 | 0.41333333 | 0.2975 | 0.05236535 | 0.01655938 |
| 3 | 0.24166667 | 0.245 | 0.31666667 | 0.32166667 | 0.22 | 0.22333333 | 0.30833333 | 0.25 | 0.315 | 0.37 | 0.28116667 | 0.05120794 | 0.01619337 |
| 4 | 0.23166667 | 0.24333333 | 0.28333333 | 0.33166667 | 0.21666667 | 0.26333333 | 0.28333333 | 0.28166667 | 0.36166667 | 0.43 | 0.29266667 | 0.06517403 | 0.02060984 |
| 5 | 0.31833333 | 0.285 | 0.20666667 | 0.30166667 | 0.27666667 | 0.26666667 | 0.31 | 0.24666667 | 0.42166667 | 0.46833333 | 0.31016667 | 0.07886717 | 0.02493999 |
| 10 | 0.26066667 | 0.24966667 | 0.28433333 | 0.31733333 | 0.28566667 | 0.29166667 | 0.276 | 0.269 | 0.301 | 0.37833333 | 0.29136667 | 0.03625542 | 0.01146497 |
| 15 | 0.296 | 0.25933333 | 0.41133333 | 0.422 | 0.32033333 | 0.34033333 | 0.37266667 | 0.24633333 | 0.25033333 | 0.27366667 | 0.31923333 | 0.06549867 | 0.0207125 |

Het

| Time (min) | B | D | E | G | H | 155 | 157 | 190 | 196 | 199 | 201 | 205 | 207 | 208 | 210 | Average | SD | SE |
|---|---|---|---|---|---|---|---|---|---|---|---|---|---|---|---|---|---|---|
| 0 | 0.41911111 | 0.31377778 | 0.38333333 | 0.50477778 | 0.29777778 | 0.26855556 | 0.40755556 | 0.40922222 | 0.39955556 | 0.34622222 | 0.42822222 | 0.38422222 | 0.38188889 | 0.47322222 | 0.45622222 | 0.39157778 | 0.06472411 | 0.01671169 |
| 1 | 0.36333333 | 0.53333333 | 0.475 | 0.32833333 | 0.32333333 | 0.508 | 0.30333333 | 0.43666667 | 0.325 | 0.325 | 0.44666667 | 0.59833333 | 0.74 | 0.36666667 | 0.53166667 | 0.44031111 | 0.12551332 | 0.0324074 |
| 2 | 0.26 | 0.37666667 | 0.40833333 | 0.375 | 0.29166667 | 0.47833333 | 0.245 | 0.19833333 | 0.26666667 | 0.19833333 | 0.21833333 | 0.59 | 0.19166667 | 0.34333333 | 0.57333333 | 0.33433333 | 0.13172814 | 0.03401206 |
| 3 | 0.26 | 0.27333333 | 0.44 | 0.455 | 0.345 | 0.34833333 | 0.25 | 0.20333333 | 0.20333333 | 0.26666667 | 0.20833333 | 0.64 | 0.22333333 | 0.34 | 0.53 | 0.33244444 | 0.13125443 | 0.03388975 |
| 4 | 0.25166667 | 0.24166667 | 0.40833333 | 0.575 | 0.335 | 0.31 | 0.265 | 0.275 | 0.27 | 0.25666667 | 0.2 | 0.58 | 0.21833333 | 0.26 | 0.29666667 | 0.31622222 | 0.1170427 | 0.0302203 |
| 5 | 0.26333333 | 0.30166667 | 0.41833333 | 0.39666667 | 0.38833333 | 0.32 | 0.21666667 | 0.255 | 0.17666667 | 0.23833333 | 0.22833333 | 0.53666667 | 0.26 | 0.33833333 | 0.44166667 | 0.31866667 | 0.09997857 | 0.02581436 |
| 10 | 0.29 | 0.27866667 | 0.35966667 | 0.56066667 | 0.31566667 | 0.319 | 0.219 | 0.27566667 | 0.199 | 0.244 | 0.227 | 0.29766667 | 0.33266667 | 0.26733333 | 0.37033333 | 0.30375556 | 0.08675202 | 0.02239927 |
| 15 | 0.30766667 | 0.41066667 | 0.33066667 | 0.49666667 | 0.311 | 0.28066667 | 0.23033333 | 0.237 | 0.255 | 0.29533333 | 0.33333333 | 0.27 | 0.343 | 0.253 | 0.304 | 0.31055556 | 0.06954029 | 0.01795523 |

Week 4

WT

| Time (min) | A | 156 | 158 | 161 | Average | SD | SE |
|---|---|---|---|---|---|---|---|
| 0 | 0.347 | 0.29877778 | 0.25544444 | 0.33566667 | 0.30922222 | 0.04134249 | 0.02067125 |
| 1 | 0.34166667 | 0.305 | 0.58166667 | 0.375 | 0.40083333 | 0.12389885 | 0.06194943 |
| 2 | 0.25333333 | 0.315 | 0.45333333 | 0.41166667 | 0.35833333 | 0.09087028 | 0.04543514 |
| 3 | 0.29 | 0.26666667 | 0.35166667 | 0.345 | 0.31333333 | 0.04161107 | 0.02080554 |
| 4 | 0.29166667 | 0.24666667 | 0.40833333 | 0.34166667 | 0.32208333 | 0.06936718 | 0.03468359 |
| 5 | 0.27666667 | 0.265 | 0.31666667 | 0.335 | 0.29833333 | 0.03297024 | 0.01648512 |
| 10 | 0.28566667 | 0.28 | 0.336 | 0.24566667 | 0.28683333 | 0.03723847 | 0.01861924 |
| 15 | 0.35033333 | 0.24066667 | 0.174 | 0.23033333 | 0.24883333 | 0.07373652 | 0.03686826 |

Het

| Time (min) | B | D | E | H | 155 | 157 | 190 | 196 | 199 | 201 | 205 | 207 | 208 | 210 | Average | SD | SEM |
|---|---|---|---|---|---|---|---|---|---|---|---|---|---|---|---|---|---|
| 0 | 0.43633333 | 0.69022222 | 0.37577778 | 0.32177778 | 0.40766667 | 0.35744444 | 0.38433333 | 0.36244444 | 0.356 | 0.34888889 | 0.32777778 | 0.33811111 | 0.29944444 | 0.43544444 | 0.38869048 | 0.09565635 | 0.02556523 |
| 1 | 0.33833333 | 0.38166667 | 0.47666667 | 0.425 | 0.575 | 0.56 | 0.51666667 | 0.33166667 | 0.27333333 | 0.31 | 0.49166667 | 0.22833333 | 0.39333333 | 0.58166667 | 0.4202381 | 0.11585036 | 0.03096231 |
| 2 | 0.25833333 | 0.46333333 | 0.415 | 0.41666667 | 0.55166667 | 0.35 | 0.36166667 | 0.27166667 | 0.22333333 | 0.215 | 0.23833333 | 0.375 | 0.27333333 | 0.26833333 | 0.33440476 | 0.10128577 | 0.02706976 |
| 3 | 0.28 | 0.39833333 | 0.34833333 | 0.43333333 | 0.50166667 | 0.45833333 | 0.29166667 | 0.21 | 0.23333333 | 0.20333333 | 0.23 | 0.30833333 | 0.26 | 0.20333333 | 0.31142857 | 0.10075327 | 0.02692744 |
| 4 | 0.26 | 0.335 | 0.39166667 | 0.41666667 | 0.37 | 0.49166667 | 0.27 | 0.21833333 | 0.24166667 | 0.22833333 | 0.19333333 | 0.30833333 | 0.265 | 0.255 | 0.30321429 | 0.08644069 | 0.02310225 |
| 5 | 0.28166667 | 0.30166667 | 0.39333333 | 0.47 | 0.38 | 0.41666667 | 0.29833333 | 0.2 | 0.22166667 | 0.205 | 0.2 | 0.31666667 | 0.29666667 | 0.375 | 0.31119048 | 0.08650732 | 0.02312005 |
| 10 | 0.259 | 0.414 | 0.32866667 | 0.37133333 | 0.39233333 | 0.38866667 | 0.29666667 | 0.20733333 | 0.266 | 0.244 | 0.21 | 0.30033333 | 0.231 | 0.23166667 | 0.29578571 | 0.07186878 | 0.01920774 |
| 15 | 0.242 | 0.721 | 0.294 | 0.24966667 | 0.32966667 | 0.38266667 | 0.483 | 0.20137931 | 0.22366667 | 0.275 | 0.23633333 | 0.277 | 0.313 | 0.22733333 | 0.31826519 | 0.1375378 | 0.03675852 |

### Expiratory flow 50% through expiration

Week 1

WT

| Time (min) | Animal ID: A | 156 | 158 | 161 | Average | SD | SE |
|---|---|---|---|---|---|---|---|
| 0 | 1.61555556 | 1.78111111 | 0.96111111 | 1.18333333 | 1.38527778 | 0.37877469 | 0.18938735 |
| 1 | 2.36666667 | 2.03333333 | 1.03333333 | 1.18333333 | 1.65416667 | 0.64769892 | 0.32384946 |
| 2 | 2.38333333 | 1.76666667 | 1.03333333 | 1.53333333 | 1.67916667 | 0.56031985 | 0.28015992 |
| 3 | 2.06666667 | 1.78333333 | 1.1 | 1.53333333 | 1.62083333 | 0.40991756 | 0.20495878 |
| 4 | 1.76666667 | 1.76666667 | 1 | 1.38333333 | 1.47916667 | 0.36701372 | 0.18350686 |
| 5 | 1.7 | 1.78333333 | 1.06666667 | 1.25 | 1.45 | 0.34667735 | 0.17333868 |
| 10 | 1.81666667 | 1.64666667 | 0.90666667 | 1.26666667 | 1.40916667 | 0.40631474 | 0.20315737 |
| 15 | 1.18333333 | 1.28666667 | 0.70666667 | 0.84666667 | 1.00583333 | 0.27398804 | 0.13699402 |

Het

| Time (min) | Animal ID: B | C | D | E | F | G | H | 155 | 157 | 159 | 160 | 162 | 190 | 196 | 199 | 201 | 205 | 207 | 208 | 210 | Average | SD | SE |
|---|---|---|---|---|---|---|---|---|---|---|---|---|---|---|---|---|---|---|---|---|---|---|---|
| 0 | 1.15555556 | 1.46888889 | 1.04 | 1.38 | 0.76333333 | 0.97888889 | 1.36888889 | 1.48222222 | 1.43 | 1.08333333 | 1.58 | 0.69555556 | 1.68 | 1.08777778 | 0.95333333 | 1.34111111 | 0.93888889 | 0.86111111 | 0.80111111 | 0.57777778 | 1.13338889 | 0.31703395 | 0.07089095 |
| 1 | 2.13333333 | 1.46666667 | 1.4 | 1.88333333 | 0.55 | 1.46666667 | 1.75 | 1.78333333 | 1.33333333 | 1.16666667 | 1.75 | 0.93333333 | 1.93333333 | 1.65 | 1.58333333 | 1.23333333 | 0.91666667 | 1.08333333 | 0.96666667 | 0.73333333 | 1.38583333 | 0.43175404 | 0.09654314 |
| 2 | 1.91666667 | 1.88333333 | 1.46666667 | 1.63333333 | 0.61666667 | 1.21666667 | 1.91666667 | 1.5 | 0.95 | 1.15 | 1.63333333 | 0.88333333 | 1.88333333 | 1.85 | 1.65 | 1.11666667 | 0.86666667 | 1.2 | 1.01666667 | 0.98333333 | 1.36666667 | 0.41496142 | 0.09278819 |
| 3 | 1.48333333 | 1.86666667 | 1.36666667 | 1.55 | 0.63333333 | 1.23333333 | 1.71666667 | 1.38333333 | 1.01666667 | 1.26666667 | 1.53333333 | 1 | 2.16666667 | 1.63333333 | 1.61666667 | 1.25 | 0.86666667 | 1.08333333 | 0.95 | 1.05 | 1.33333333 | 0.37271722 | 0.0833421 |
| 4 | 1.56666667 | 1.75 | 1.28333333 | 1.8 | 0.65 | 1.31666667 | 1.46666667 | 1.35 | 1.01666667 | 1.15 | 1.3 | 1.01666667 | 1.75 | 1.45 | 1.48333333 | 1.4 | 0.75 | 1.13333333 | 0.95 | 1.1 | 1.28416667 | 0.31758753 | 0.07101473 |
| 5 | 1.43333333 | 1.78333333 | 1.2 | 1.48333333 | 0.61666667 | 1.36666667 | 1.26666667 | 1.35 | 1.08333333 | 1.13333333 | 1.41666667 | 1 | 1.4 | 1.23333333 | 1.33333333 | 1.18333333 | 0.78333333 | 1.08333333 | 1.03333333 | 0.95 | 1.20666667 | 0.26265625 | 0.05873172 |
| 10 | 1.23 | 1.51333333 | 0.88 | 0.99666667 | 0.56666667 | 1.16 | 1.17333333 | 1.24666667 | 0.97666667 | 0.96666667 | 1.36 | 0.98 | 1.53666667 | 1.19 | 1.19666667 | 1.21333333 | 0.83666667 | 0.84 | 0.62333333 | 0.78 | 1.06333333 | 0.26530794 | 0.05932466 |
| 15 | 1.14666667 | 1.18666667 | 0.95 | 0.99666667 | 0.61666667 | 0.92666667 | 1.07333333 | 1.12333333 | 0.86 | 0.59333333 | 1.23333333 | 0.60666667 | 1.33666667 | 1.1 | 0.94 | 0.96 | 0.83666667 | 0.58 | 0.52 | 0.60333333 | 0.9095 | 0.24936475 | 0.05575965 |

Week 2

WT

| Time (min) | Animal ID: A | 156 | 158 | 161 | Average | SD | SE |
|---|---|---|---|---|---|---|---|
| 0 | 1.67888889 | 1.32888889 | 1.27444444 | 1.69 | 1.49305556 | 0.22215809 | 0.11107905 |
| 1 | 2.46666667 | 2.65 | 1.41666667 | 1.75 | 2.07083333 | 0.58394809 | 0.29197404 |
| 2 | 2.3 | 2.03333333 | 1.55 | 1.5 | 1.84583333 | 0.38667026 | 0.19333513 |
| 3 | 2.23333333 | 1.95 | 1.35 | 1.43333333 | 1.74166667 | 0.42174681 | 0.2108734 |
| 4 | 1.86666667 | 1.75 | 1.31666667 | 1.41666667 | 1.5875 | 0.26259919 | 0.13129959 |
| 5 | 1.71666667 | 1.63333333 | 1.4 | 1.48333333 | 1.55833333 | 0.14304881 | 0.07152441 |
| 10 | 1.63 | 1.74 | 1.05333333 | 1.32666667 | 1.4375 | 0.31007018 | 0.15503509 |
| 15 | 1.46333333 | 1.42333333 | 0.84333333 | 0.97666667 | 1.17666667 | 0.3131205 | 0.15656025 |

Het

| Time (min) | Animal ID: B | D | E | G | H | 155 | 157 | 160 | 162 | 190 | 196 | 199 | 201 | 205 | 207 | 208 | 210 | Average | SD | SE |
|---|---|---|---|---|---|---|---|---|---|---|---|---|---|---|---|---|---|---|---|---|
| 0 | 1.07666667 | 1.44333333 | 0.59666667 | 0.67888889 | 1.53111111 | 1.14444444 | 1.14333333 | 1.23 | 1.02444444 | 1.50444444 | 1.11 | 1.06555556 | 1.32222222 | 1.47555556 | 1.13 | 1.47111111 | 1.54222222 | 1.20529412 | 0.28089296 | 0.06812655 |
| 1 | 2.3 | 2.51666667 | 1.1 | 0.98333333 | 1.8 | 2 | 1.36666667 | 1.55 | 1.35 | 1.78 | 2.14 | 1.2 | 1.92 | 1.34 | 1.38 | 1.48 | 1.46 | 1.62745098 | 0.43327168 | 0.10508382 |
| 2 | 2.3 | 2.46666667 | 1.3 | 1.13333333 | 1.86666667 | 1.66666667 | 1.51666667 | 1.45 | 0.68333333 | 1.91666667 | 2.36666667 | 1.56666667 | 1.3 | 1.03333333 | 0.93333333 | 0.73333333 | 0.86666667 | 1.47647059 | 0.5628258 | 0.13650531 |
| 3 | 2.18333333 | 2.38333333 | 1.15 | 1.08333333 | 1.91666667 | 1.35 | 1.43333333 | 1.46666667 | 0.7 | 1.78333333 | 2.16666667 | 1.71666667 | 1.3 | 0.9 | 0.91666667 | 0.48333333 | 1.1 | 1.41372549 | 0.54566586 | 0.13234341 |
| 4 | 2.01666667 | 2.2 | 1.23333333 | 1.15 | 1.48333333 | 1.41666667 | 1.38333333 | 1.4 | 0.83333333 | 1.5 | 1.95 | 1.73333333 | 1.25 | 0.76666667 | 0.86666667 | 0.5 | 0.83333333 | 1.3245098 | 0.47467524 | 0.11512566 |
| 5 | 1.7 | 2.01666667 | 0.95 | 1.18333333 | 1.38333333 | 1.18333333 | 1.31666667 | 1.16666667 | 0.81666667 | 1.78333333 | 1.6 | 1.85 | 1.31666667 | 0.81666667 | 0.85 | 0.51666667 | 0.95 | 1.25882353 | 0.4221768 | 0.10239291 |
| 10 | 1.52 | 1.79 | 0.79 | 0.95666667 | 1.32666667 | 1.26666667 | 1.29333333 | 0.95666667 | 0.86333333 | 1.38333333 | 1.59333333 | 1.45666667 | 1.52 | 0.88333333 | 0.95 | 0.78 | 0.99 | 1.19529412 | 0.3181065 | 0.07715216 |
| 15 | 1.43 | 1.33666667 | 0.44 | 0.8 | 1.19 | 1.09333333 | 1.55 | 0.77666667 | 0.61 | 1.26666667 | 1.23 | 1.35 | 1.69666667 | 0.74333333 | 0.95333333 | 0.91666667 | 0.79666667 | 1.06941176 | 0.35066254 | 0.08504816 |

Week 3

WT

| Time (min) | Animal ID: 83 | 86 | 87 | 88 | 81 | 82 | A | 156 | 158 | 161 | Average | SD | SE |
|---|---|---|---|---|---|---|---|---|---|---|---|---|---|
| 0 | 2.00888889 | 1.39777778 | 1.95444444 | 2.19888889 | 1.42359551 | 1.26111111 | 2.11555556 | 1.65555556 | 0.35122222 | 1.61888889 | 1.59859288 | 0.54436648 | 0.1721438 |
| 1 | 2.65 | 3.03333333 | 4 | 4.21666667 | 3.91666667 | 3.96666667 | 3.23333333 | 3.34 | 0.395 | 2.28333333 | 3.1035 | 1.14217965 | 0.36118892 |
| 2 | 2.2 | 2.86666667 | 3.1 | 2.83333333 | 3.01666667 | 3.58333333 | 2.88333333 | 3.31666667 | 0.34 | 1.61666667 | 2.57566667 | 0.96255582 | 0.30438688 |
| 3 | 1.85 | 2.73333333 | 2.51666667 | 2.71666667 | 2.31666667 | 2.63333333 | 2.25 | 2.8 | 0.315 | 1.45 | 2.15816667 | 0.7771944 | 0.24577045 |
| 4 | 1.7 | 2.48333333 | 2.38333333 | 2.76666667 | 2.1 | 2.36666667 | 2.66666667 | 2.9 | 0.36166667 | 1.33333333 | 2.10616667 | 0.77982114 | 0.2466011 |
| 5 | 1.71666667 | 2.13333333 | 2.15 | 2.88333333 | 2.23333333 | 2.25 | 1.95 | 2.25 | 0.42166667 | 1.81666667 | 1.9805 | 0.63278243 | 0.20010338 |
| 10 | 1.45666667 | 2.1 | 1.96666667 | 2.08 | 2.34666667 | 2.13666667 | 1.96 | 2.12333333 | 0.301 | 1.68666667 | 1.81576667 | 0.58819389 | 0.18600324 |
| 15 | 1.81666667 | 1.47 | 1.52666667 | 1.39 | 1.82666667 | 1.81 | 1.81666667 | 1.26333333 | 0.25033333 | 1.61666667 | 1.4787 | 0.47654193 | 0.15069579 |

Het

| Time (min) | Animal ID: B | D | E | G | H | 155 | 157 | 190 | 196 | 199 | 201 | 205 | 207 | 208 | 210 | Average | SD | SE |
|---|---|---|---|---|---|---|---|---|---|---|---|---|---|---|---|---|---|---|
| 0 | 1.21333333 | 1.46444444 | 0.91111111 | 0.53333333 | 1.60666667 | 1.95777778 | 1.12444444 | 1.24888889 | 1.20111111 | 1.22666667 | 1.18888889 | 0.72 | 0.84111111 | 0.77444444 | 0.84333333 | 1.1237037 | 0.3709777 | 0.09578603 |
| 1 | 2.68333333 | 2.28333333 | 2.1 | 0.41666667 | 2.16666667 | 2.26 | 2.05 | 1.8 | 1.91666667 | 1.46666667 | 1.66666667 | 1.4 | 1.56666667 | 0.83333333 | 1.31666667 | 1.72844444 | 0.59005093 | 0.1523505 |
| 2 | 2.76666667 | 2.31666667 | 2.3 | 0.36666667 | 2.21666667 | 1.63333333 | 1.83333333 | 1.41666667 | 1.43333333 | 1.43333333 | 1.43333333 | 1.01666667 | 1.03333333 | 0.9 | 1.03333333 | 1.54222222 | 0.64702821 | 0.16706197 |
| 3 | 2.1 | 2.31666667 | 2.38333333 | 0.38333333 | 2.31666667 | 1.71666667 | 1.86666667 | 1.53333333 | 1.58333333 | 1.58333333 | 1.65 | 0.96666667 | 0.81666667 | 1.05 | 1.5 | 1.58444444 | 0.58474206 | 0.15097975 |
| 4 | 2.71666667 | 1.53333333 | 1.8 | 0.36666667 | 2.06666667 | 1.56666667 | 1.91666667 | 1.55 | 1.78333333 | 1.88333333 | 1.68333333 | 0.95 | 0.78333333 | 0.93333333 | 1.61666667 | 1.54333333 | 0.58193029 | 0.15025375 |
| 5 | 2.23333333 | 1.4 | 1.93333333 | 0.41666667 | 1.95 | 1.46666667 | 1.73333333 | 1.58333333 | 1.68333333 | 1.6 | 1.85 | 0.9 | 0.81666667 | 0.91666667 | 0.91666667 | 1.42666667 | 0.5187286 | 0.13393515 |
| 10 | 1.61333333 | 1.42666667 | 0.94666667 | 0.45333333 | 1.72666667 | 1.48666667 | 1.52666667 | 1.65 | 1.66 | 1.63333333 | 1.56333333 | 0.94666667 | 0.93333333 | 1.16 | 1.05 | 1.31844444 | 0.37730428 | 0.09741955 |
| 15 | 1.47 | 1.1 | 1.15666667 | 0.5 | 1.71 | 1.48666667 | 1.79666667 | 1.52333333 | 1.9 | 1.07 | 1.18 | 0.81666667 | 0.85 | 0.89666667 | 1.07 | 1.23511111 | 0.4004261 | 0.10338957 |

Week 4

WT

| Time (min) | Animal ID: A | 156 | 158 | 161 | Average | SD | SE |
|---|---|---|---|---|---|---|---|
| 0 | 2.21222222 | 2.30222222 | 1.11666667 | 2.39222222 | 2.00583333 | 0.59731524 | 0.29865762 |
| 1 | 3.18333333 | 2.86666667 | 1.65 | 1.95 | 2.4125 | 0.72932935 | 0.36466467 |
| 2 | 2.51666667 | 2.46666667 | 1.53333333 | 1.4 | 1.97916667 | 0.59463263 | 0.29731631 |
| 3 | 2.45 | 2.4 | 1.26666667 | 1.58333333 | 1.925 | 0.59199912 | 0.29599956 |
| 4 | 3.08333333 | 2.38333333 | 1.73333333 | 1.4 | 2.15 | 0.74423712 | 0.37211856 |
| 5 | 2.18333333 | 2.26666667 | 1.88333333 | 1.45 | 1.94583333 | 0.36927732 | 0.18463866 |
| 10 | 1.88333333 | 2.05333333 | 1.55333333 | 1.38 | 1.7175 | 0.30612724 | 0.15306362 |
| 15 | 1.46666667 | 1.61666667 | 1.14666667 | 1.38666667 | 1.40416667 | 0.19636276 | 0.09818138 |

Het

| Time (min) | Animal ID: B | D | E | H | 155 | 157 | 190 | 196 | 199 | 201 | 205 | 207 | 208 | 210 | Average | SD | SEM |
|---|---|---|---|---|---|---|---|---|---|---|---|---|---|---|---|---|---|
| 0 | 1.38333333 | 0.54777778 | 0.86888889 | 1.34555556 | 1.29888889 | 1.37888889 | 1.26777778 | 1.29333333 | 1.17222222 | 1.41555556 | 1.10666667 | 0.74555556 | 1.62111111 | 0.90333333 | 1.16777778 | 0.29848175 | 0.0797726 |
| 1 | 2.78333333 | 1.43333333 | 1.91666667 | 2.55 | 2.8 | 2.38333333 | 2.41666667 | 1.96666667 | 1.9 | 1.76666667 | 1.58333333 | 1.31666667 | 1.8 | 1.81666667 | 2.03095238 | 0.47805952 | 0.12776678 |
| 2 | 2.05 | 1.16666667 | 2.3 | 2.51666667 | 1.96666667 | 1.51666667 | 1.78333333 | 1.38333333 | 1.5 | 1.85 | 1.28333333 | 1.86666667 | 1.7 | 0.91666667 | 1.7 | 0.44057073 | 0.11774748 |
| 3 | 2.18333333 | 1.3 | 1.65 | 2.05 | 1.78333333 | 1.75 | 1.58333333 | 1.61666667 | 1.45 | 2.11666667 | 1.3 | 1.58333333 | 1.55 | 0.88333333 | 1.62857143 | 0.34822066 | 0.09306589 |
| 4 | 1.95 | 1.16666667 | 2.2 | 2.98333333 | 1.75 | 1.81666667 | 1.73333333 | 1.68333333 | 1.8 | 2.16666667 | 1.25 | 1.53333333 | 1.53333333 | 1.03333333 | 1.75714286 | 0.49193123 | 0.13147415 |
| 5 | 2.05 | 1.31666667 | 1.51666667 | 1.76666667 | 1.75 | 1.8 | 1.53333333 | 1.56666667 | 1.81666667 | 1.88333333 | 1.06666667 | 1.73333333 | 1.65 | 1.1 | 1.61071429 | 0.28634203 | 0.07652813 |
| 10 | 1.61 | 1.07333333 | 1.32333333 | 2.08666667 | 1.72666667 | 1.68333333 | 1.58333333 | 1.58 | 1.60333333 | 1.72 | 1.07666667 | 1.26333333 | 0.92 | 1.20666667 | 1.46119048 | 0.32386187 | 0.08655572 |
| 15 | 1.69 | 0.58 | 0.87666667 | 1.28333333 | 1.47666667 | 1.71333333 | 1.33333333 | 1.37931034 | 1.12 | 1.39333333 | 1.00666667 | 1.14666667 | 0.84666667 | 1.29333333 | 1.22423645 | 0.32049317 | 0.0856554 |

### Minute ventilation

Week 1

WT

| Time (min) | A | 156 | 158 | 161 | Average | SD | SE |
|---|---|---|---|---|---|---|---|
| 0 | 52.5333333 | 55.7666667 | 33.7666667 | 43.4333333 | 46.375 | 9.89531782 | 4.94765891 |
| 1 | 64 | 63.1666667 | 33.1666667 | 39.8333333 | 50.0416667 | 15.8753099 | 7.93765493 |
| 2 | 73.1666667 | 55.8333333 | 36.5 | 53.6666667 | 54.7916667 | 14.9952925 | 7.49764624 |
| 3 | 65 | 58.6666667 | 35.5 | 52 | 52.7916667 | 12.6910401 | 6.34552007 |
| 4 | 57.3333333 | 54.3333333 | 34.6666667 | 43.5 | 47.4583333 | 10.3935299 | 5.19676493 |
| 5 | 53.3333333 | 50.8333333 | 35.6666667 | 44.5 | 46.0833333 | 7.87694714 | 3.93847357 |
| 10 | 55 | 49.0666667 | 31 | 41.1666667 | 44.0583333 | 10.3872563 | 5.19362813 |
| 15 | 32 | 35.3 | 22.7666667 | 26.6 | 29.1666667 | 5.57354864 | 2.78677432 |

Het

| Time (min) | B | C | D | E | F | G | H | 155 | 157 | 159 | 160 | 162 | 190 | 196 | 199 | 201 | 205 | 207 | 208 | 210 | Average | SD | SE |
|---|---|---|---|---|---|---|---|---|---|---|---|---|---|---|---|---|---|---|---|---|---|---|---|
| 0 | 29.2666667 | 43.0888889 | 31.2888889 | 40.8111111 | 19.4777778 | 30.4111111 | 44.9111111 | 42.4111111 | 41.7333333 | 34.9 | 48.2777778 | 21.8888889 | 54.8666667 | 30.0444444 | 25.2333333 | 42.0666667 | 30.3444444 | 28.4333333 | 25.5333333 | 17.4888889 | 34.1238889 | 10.1691256 | 2.2738856 |
| 1 | 61 | 51.6 | 41 | 57.6666667 | 20.1666667 | 47.5 | 60.1666667 | 45.8333333 | 35.8333333 | 39.1666667 | 49.8333333 | 29.8333333 | 53.5 | 47 | 47.6666667 | 36.5 | 27.3333333 | 36.1666667 | 31.6666667 | 26 | 42.2716667 | 11.7803746 | 2.63417184 |
| 2 | 54.3333333 | 52.5 | 38.1666667 | 52.6666667 | 16 | 44.6666667 | 63.6666667 | 44.3333333 | 28.6666667 | 38.5 | 47 | 30.8333333 | 63.3333333 | 50.1666667 | 48.1666667 | 33.6666667 | 24.1666667 | 38.1666667 | 32.8333333 | 30.3333333 | 41.6083333 | 12.7064559 | 2.84124991 |
| 3 | 49.1666667 | 50.6666667 | 37 | 54.1666667 | 15.8333333 | 46.1666667 | 61.5 | 44.3333333 | 33.6666667 | 40.6666667 | 47.1666667 | 33.6666667 | 67.5 | 46.1666667 | 46.5 | 37.3333333 | 23.5 | 36.8333333 | 31.3333333 | 30.3333333 | 41.675 | 12.3138152 | 2.75345278 |
| 4 | 46.6666667 | 50.8333333 | 35.8333333 | 55.1666667 | 14.5 | 43.6666667 | 46.5 | 40.8333333 | 30.8333333 | 40.5 | 41.5 | 35.1666667 | 62 | 44.6666667 | 46 | 42.3333333 | 22.1666667 | 40.1666667 | 31.8333333 | 34.1666667 | 40.2666667 | 10.7368034 | 2.40082223 |
| 5 | 42.1666667 | 49.3333333 | 36 | 46.1666667 | 13.3333333 | 44.8333333 | 43 | 38.8333333 | 32.6666667 | 36.8333333 | 41.3333333 | 31.8333333 | 41.3333333 | 34.5 | 42.3333333 | 35.5 | 22.3333333 | 36.6666667 | 32.6666667 | 28.1666667 | 36.4916667 | 8.43403627 | 1.88590784 |
| 10 | 31.8666667 | 41.4333333 | 26.8666667 | 26.6333333 | 13.1333333 | 38.3333333 | 39.9666667 | 36.2333333 | 26.9333333 | 30.7 | 37.2333333 | 32.4666667 | 48.6 | 31.2333333 | 39.5666667 | 36.5333333 | 25.2333333 | 28.4666667 | 19 | 23.6333333 | 31.7033333 | 8.35414174 | 1.86804288 |
| 15 | 28.5333333 | 31.1333333 | 29.0666667 | 26.1333333 | 14.8 | 30.5 | 34.2666667 | 29.3 | 22.4666667 | 19.1666667 | 31.4 | 19.3666667 | 37.8666667 | 30.3 | 27.7666667 | 28.5666667 | 27.0333333 | 17.6666667 | 16.0333333 | 17.4333333 | 25.94 | 6.53584449 | 1.46145926 |

Week 2

WT

| Time (min) | A | 156 | 158 | 161 | Average | SD | SE |
|---|---|---|---|---|---|---|---|
| 0 | 55.4444444 | 39.7555556 | 44.2555556 | 57.4 | 49.2138889 | 8.5611015 | 4.28055075 |
| 1 | 77 | 74.1666667 | 47.5 | 62 | 65.1666667 | 13.4556886 | 6.7278443 |
| 2 | 75.8333333 | 62.1666667 | 52.1666667 | 52.1666667 | 60.5833333 | 11.206397 | 5.60319849 |
| 3 | 77.8333333 | 65 | 51.6666667 | 48.3333333 | 60.7083333 | 13.4978565 | 6.74892824 |
| 4 | 65.6666667 | 57.1666667 | 53.3333333 | 50.5 | 56.6666667 | 6.59264461 | 3.29632231 |
| 5 | 54.8333333 | 55.8333333 | 45.5 | 48 | 51.0416667 | 5.07604213 | 2.53802106 |
| 10 | 55.6333333 | 56.4 | 36.5333333 | 45.3666667 | 48.4833333 | 9.42182258 | 4.71091129 |
| 15 | 46.0666667 | 43.7 | 27.9333333 | 35.3333333 | 38.2583333 | 8.28136574 | 4.14068287 |

Het

| Time (min) | B | D | E | G | H | 155 | 157 | 160 | 162 | 190 | 196 | 199 | 201 | 205 | 207 | 208 | 210 | Average | SD | SE |
|---|---|---|---|---|---|---|---|---|---|---|---|---|---|---|---|---|---|---|---|---|
| 0 | 30.9222222 | 41.6888889 | 19.6888889 | 21.0555556 | 54.2333333 | 31.1888889 | 31.8 | 38.7888889 | 35.4555556 | 45.3888889 | 32 | 30.5444444 | 38.3111111 | 49.6444444 | 37.4222222 | 47.1666667 | 54.4888889 | 37.6346405 | 10.2390471 | 2.48333368 |
| 1 | 67.8333333 | 69.1666667 | 41 | 40.6666667 | 62.3333333 | 51 | 37.1666667 | 49.1666667 | 39 | 41.4 | 63 | 37.4 | 46.4 | 31.4 | 34.6 | 37.8 | 34.8 | 46.1254902 | 12.2868806 | 2.98000626 |
| 2 | 73.3333333 | 76.6666667 | 48.6666667 | 41.3333333 | 62.5 | 54.5 | 46.3333333 | 45.1666667 | 24.6666667 | 60.8333333 | 70.3333333 | 49.5 | 38.3333333 | 26.8333333 | 33 | 27.8333333 | 26.3333333 | 47.4215686 | 16.9905118 | 4.1208044 |
| 3 | 68.8333333 | 73.1666667 | 40.8333333 | 43.5 | 65.1666667 | 47.8333333 | 41.1666667 | 46.1666667 | 25.6666667 | 51 | 64.1666667 | 52.8333333 | 38.6666667 | 31.3333333 | 29.8333333 | 19.5 | 36.1666667 | 45.6372549 | 15.4688571 | 3.75174892 |
| 4 | 66.1666667 | 73.8333333 | 46 | 40.3333333 | 50.6666667 | 49 | 40.6666667 | 43.1666667 | 29.5 | 41.1666667 | 60.6666667 | 52.5 | 36.1666667 | 26.1666667 | 26 | 16.8333333 | 29.3333333 | 42.8333333 | 15.1694137 | 3.67912323 |
| 5 | 58.5 | 63.5 | 34.5 | 44.5 | 42.3333333 | 44.1666667 | 38 | 38 | 29.6666667 | 51 | 49.3333333 | 61.1666667 | 37.8333333 | 27.1666667 | 25.3333333 | 18.3333333 | 33.6666667 | 41 | 12.7861687 | 3.10110143 |
| 10 | 54.0333333 | 53.7 | 30.3 | 35.9 | 40.4666667 | 42.9666667 | 38.6333333 | 30.7 | 30.7666667 | 40.2666667 | 53.2666667 | 43.2 | 44 | 29.3 | 30.6 | 24.9 | 34.8 | 38.6941176 | 9.04634084 | 2.19405993 |
| 15 | 51.3666667 | 36.1666667 | 14.6666667 | 30.1333333 | 40.7333333 | 36.9 | 45.6333333 | 24.9333333 | 22.2333333 | 34.6666667 | 45.1 | 38.8333333 | 53 | 25.3666667 | 31.3333333 | 30.5 | 24.5333333 | 34.4764706 | 10.5842898 | 2.56706733 |

Week 3

WT

| Time (min) | 83 | 86 | 87 | 88 | 81 | 82 | A | 156 | 158 | 161 | Average | SD | SE |
|---|---|---|---|---|---|---|---|---|---|---|---|---|---|
| 0 | 72.3222222 | 42.9111111 | 62.8555556 | 78 | 47.9550562 | 37.2777778 | 74.0111111 | 53.9888889 | 51.7111111 | 49.0222222 | 57.0055056 | 14.0213866 | 4.43395177 |
| 1 | 85.3333333 | 89.5 | 121.166667 | 124.166667 | 114.666667 | 109.833333 | 98.6666667 | 111 | 54.3333333 | 77.5 | 98.6166667 | 22.0216013 | 6.9638418 |
| 2 | 71.1666667 | 90.1666667 | 100.333333 | 92.1666667 | 100.666667 | 106 | 94.6666667 | 105.833333 | 53.5 | 57.1666667 | 87.1666667 | 19.5460836 | 6.18101434 |
| 3 | 66 | 89 | 84.5 | 89 | 84.3333333 | 80.8333333 | 80.8333333 | 92 | 55.3333333 | 51.6666667 | 77.35 | 14.4661056 | 4.57458424 |
| 4 | 63.8333333 | 82.6666667 | 86 | 92 | 77.6666667 | 74.6666667 | 87.8333333 | 94.5 | 55.5 | 51.5 | 76.6166667 | 15.0996791 | 4.77493779 |
| 5 | 55 | 71.1666667 | 72.6666667 | 97 | 82.5 | 73.5 | 69.5 | 80.5 | 42.5 | 59.1666667 | 70.35 | 15.3255717 | 4.8463713 |
| 10 | 55.9 | 73.4333333 | 70.0333333 | 72.6 | 83.6666667 | 65.3666667 | 68.9333333 | 70.8666667 | 41.9333333 | 54.6333333 | 65.7366667 | 11.8771744 | 3.75589233 |
| 15 | 64.9 | 48.9 | 48.7 | 46.3666667 | 69.8333333 | 53.3 | 55.2 | 43.7666667 | 30.1 | 56.8 | 51.7866667 | 11.1484174 | 3.52543912 |

Het

| Time (min) | B | D | E | G | H | 155 | 157 | 190 | 196 | 199 | 201 | 205 | 207 | 208 | 210 | Average | SD | SE |
|---|---|---|---|---|---|---|---|---|---|---|---|---|---|---|---|---|---|---|
| 0 | 37.3888889 | 47.3444444 | 31.1 | 17.2333333 | 55.1888889 | 68.8222222 | 31.5777778 | 37 | 36.2555556 | 36.8444444 | 35.0444444 | 23.8555556 | 29.4111111 | 25.5 | 25.8 | 35.8911111 | 13.0372971 | 3.36621563 |
| 1 | 75 | 58.3333333 | 59.5 | 17 | 75.5 | 51.8 | 62.8333333 | 50 | 58 | 43 | 43.8333333 | 38 | 41.5 | 29.6666667 | 33.3333333 | 49.1533333 | 16.3436887 | 4.21992226 |
| 2 | 84.8333333 | 70.6666667 | 74.1666667 | 16 | 77.6666667 | 41 | 54 | 48.8333333 | 51.5 | 45.1666667 | 47.5 | 26.5 | 33.1666667 | 33.1666667 | 27.5 | 48.7777778 | 20.4921265 | 5.29104431 |
| 3 | 64.1666667 | 74 | 78.6666667 | 16.6666667 | 77.6666667 | 48 | 57.5 | 48 | 56.3333333 | 47.6666667 | 50.8333333 | 25 | 26.5 | 37 | 42 | 50 | 18.8902427 | 4.8774397 |
| 4 | 85.1666667 | 48.6666667 | 68.8333333 | 15.3333333 | 70.8333333 | 45.6666667 | 60.6666667 | 46.8333333 | 59.6666667 | 55.5 | 51.6666667 | 26.6666667 | 27.6666667 | 35 | 53 | 50.0777778 | 18.4684272 | 4.76852741 |
| 5 | 78.1666667 | 43 | 75 | 16.5 | 69.8333333 | 43.5 | 55.5 | 48.3333333 | 55 | 47.8333333 | 55.5 | 26.8333333 | 27.5 | 32.5 | 29.6666667 | 46.9777778 | 18.407735 | 4.75285675 |
| 10 | 54.9 | 46.2666667 | 36.4333333 | 16.6666667 | 59.5333333 | 43.4666667 | 49.8333333 | 49.6666667 | 55.3333333 | 51.5333333 | 48.1 | 33.9333333 | 28.6 | 40.7 | 36.1666667 | 43.4088889 | 11.4752354 | 2.96289303 |
| 15 | 49.6 | 32.2 | 41.9333333 | 15.5 | 59.8666667 | 45.3333333 | 57 | 50.1333333 | 60.9 | 33.5333333 | 36.1666667 | 27.3 | 26.1666667 | 32.0333333 | 37.8666667 | 40.3688889 | 13.3195671 | 3.43909744 |

Week 4

WT

| Time (min) | A | 156 | 158 | 161 | Average | SD | SE |
|---|---|---|---|---|---|---|---|
| 0 | 71.1444444 | 77.7333333 | 38.5444444 | 77.2111111 | 66.1583333 | 18.6505825 | 9.32529126 |
| 1 | 105.5 | 91.6666667 | 67 | 59.6666667 | 80.9583333 | 21.3317599 | 10.66588 |
| 2 | 83 | 80.1666667 | 61 | 50.5 | 68.6666667 | 15.5617051 | 7.78085257 |
| 3 | 82 | 78.3333333 | 46.1666667 | 53.6666667 | 65.0416667 | 17.7942849 | 8.89714244 |
| 4 | 99.6666667 | 80.8333333 | 56 | 48.8333333 | 71.3333333 | 23.3408718 | 11.6704359 |
| 5 | 79 | 73 | 66.3333333 | 50.6666667 | 67.25 | 12.2061763 | 6.10308817 |
| 10 | 65.1 | 67.3333333 | 54.6 | 49.1666667 | 59.05 | 8.61585532 | 4.30792766 |
| 15 | 48.2 | 56.7 | 38.4333333 | 49.1333333 | 48.1166667 | 7.49402231 | 3.74701115 |

Het

| Time (min) | B | D | E | H | 155 | 157 | 190 | 196 | 199 | 201 | 205 | 207 | 208 | 210 | Average | SD | SEM |
|---|---|---|---|---|---|---|---|---|---|---|---|---|---|---|---|---|---|
| 0 | 43.8333333 | 20.0333333 | 29.8555556 | 47.7 | 38.7333333 | 41.0555556 | 38 | 38.8111111 | 35.2111111 | 43.2444444 | 37.6 | 25.6222222 | 53.4444444 | 28.7 | 37.2746032 | 8.91881546 | 2.3836537 |
| 1 | 82.1666667 | 46.8333333 | 53.1666667 | 85.1666667 | 62.1666667 | 56 | 57.5 | 58 | 55 | 50.8333333 | 39.3333333 | 46.6666667 | 58.8333333 | 46 | 56.9761905 | 12.8870599 | 3.44421163 |
| 2 | 63.5 | 43.3333333 | 74.6666667 | 86.3333333 | 44.5 | 38.5 | 49 | 46 | 47.6666667 | 57.6666667 | 47 | 63 | 66 | 31 | 54.1547619 | 15.0698465 | 4.0275859 |
| 3 | 67.3333333 | 45.3333333 | 65.1666667 | 77.5 | 44.3333333 | 43.3333333 | 47.5 | 48.6666667 | 45.8333333 | 59.6666667 | 43.5 | 57.5 | 58.6666667 | 30.3333333 | 52.4761905 | 12.3364267 | 3.29704872 |
| 4 | 65.1666667 | 43.3333333 | 76.5 | 89.3333333 | 47.8333333 | 45.3333333 | 53.1666667 | 52.5 | 56.5 | 61.3333333 | 42 | 55 | 56 | 32.5 | 55.4642857 | 14.5457102 | 3.88750458 |
| 5 | 71.8333333 | 50.8333333 | 54.6666667 | 62.1666667 | 47.6666667 | 47.1666667 | 46.1666667 | 49.5 | 58.8333333 | 56 | 36 | 59 | 60.8333333 | 34.3333333 | 52.5 | 10.1600441 | 2.715386 |
| 10 | 54.9666667 | 35.4666667 | 51.9333333 | 72 | 46.5 | 44.8 | 48.2 | 48.6333333 | 52.7333333 | 53.1333333 | 35.8 | 46.2666667 | 34.8 | 41.7333333 | 47.6404762 | 9.69663946 | 2.5915359 |
| 15 | 58.7 | 27.6666667 | 29.6 | 48.8 | 44.4 | 46.7333333 | 37.3666667 | 44.7931034 | 36.1 | 43.8333333 | 35.1 | 41.6 | 27.6333333 | 47.3333333 | 40.6899836 | 8.93794314 | 2.38876578 |

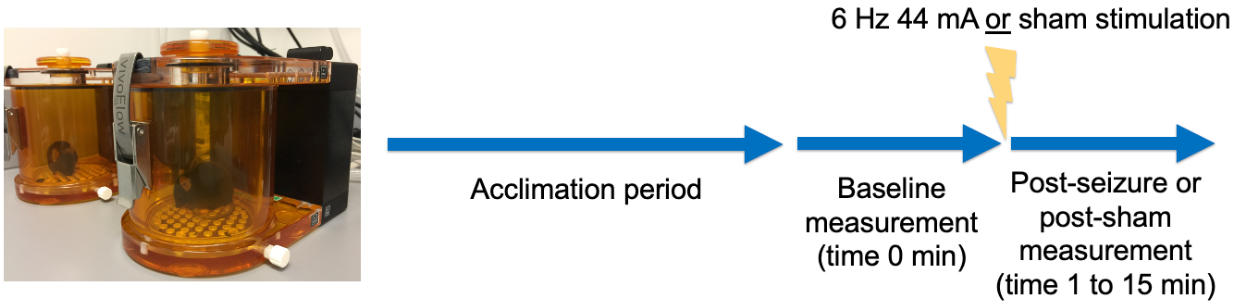


**Figure S1.** WBP experimental procedure. Animals were allowed to acclimate in the chamber for 45 minutes after which baseline measurements were taken for 15 minutes. Seizure or sham seizure was induced, then 15 minutes of post-seizure/post-sham monitoring was recorded.

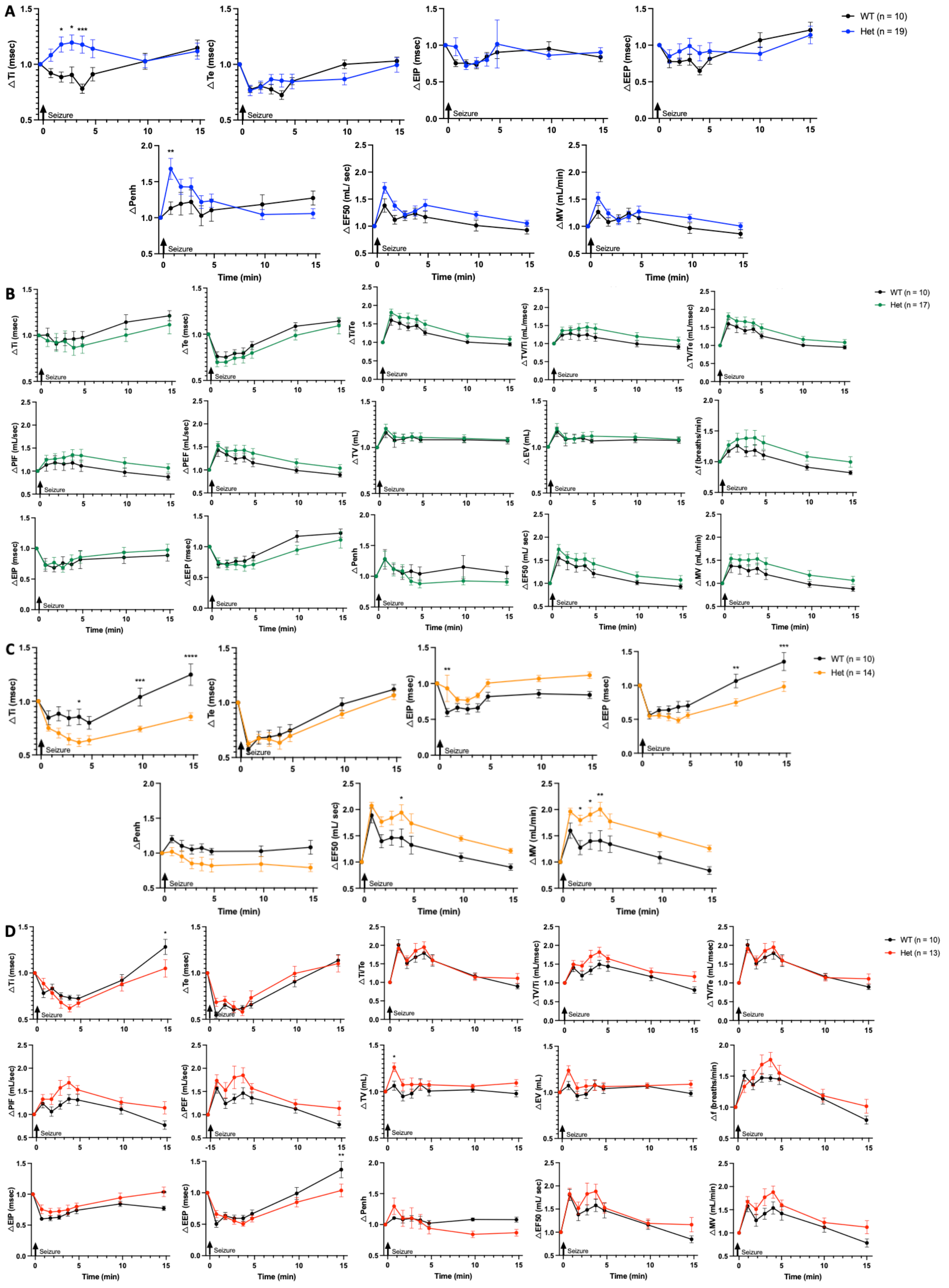
A
WT (n = 10)
Het (n = 19)
∆Ti (msec)
∆Te (msec)
∆EIP (msec)
∆EEP (msec)
∆Penh
∆EF50 (mL/ sec)
∆MV (mL/min)
Seizure
Time (min)
B
WT (n = 10)
Het (n = 17)
∆Ti/Te
∆TV/Ti (mL/msec)
∆TV/Te (mL/msec)
∆PIF (mL/sec)
∆PEF (mL/sec)
∆TV (mL)
∆EV (mL)
∆f (breaths/min)
C
WT (n = 10)
Het (n = 14)
D
WT (n = 10)
Het (n = 13)

**Figure S2.** Respiration following electrically induced 6 Hz seizure in WT and shown for week 1 (**A**), week 2 (**B**), week 3 (**C**), and week 4 (**D**). Fifteen minutes of post-seizure respiratory monitoring is normalized to baseline: time of inspiration (Ti), time of expiration (Te), time in inspiration/time of expiration (Ti/Te), inspiratory drive (TV/Ti), expiratory drive (TV/Te), peak inspiratory flow (PEF), peak expiratory flow (PEF), tidal volume (TV), expiratory volume (EV), frequency (f), end inspiratory pause (EIP), end expiratory pause (EEP), enhanced pause (Penh), expiratory flow 50% through expiration (EF50), and minute ventilation (MV) were derived from waveforms collected. Two-way ANOVA, *P<0.05, **P<0.01, ***P<0.001, ****P<0.0001.

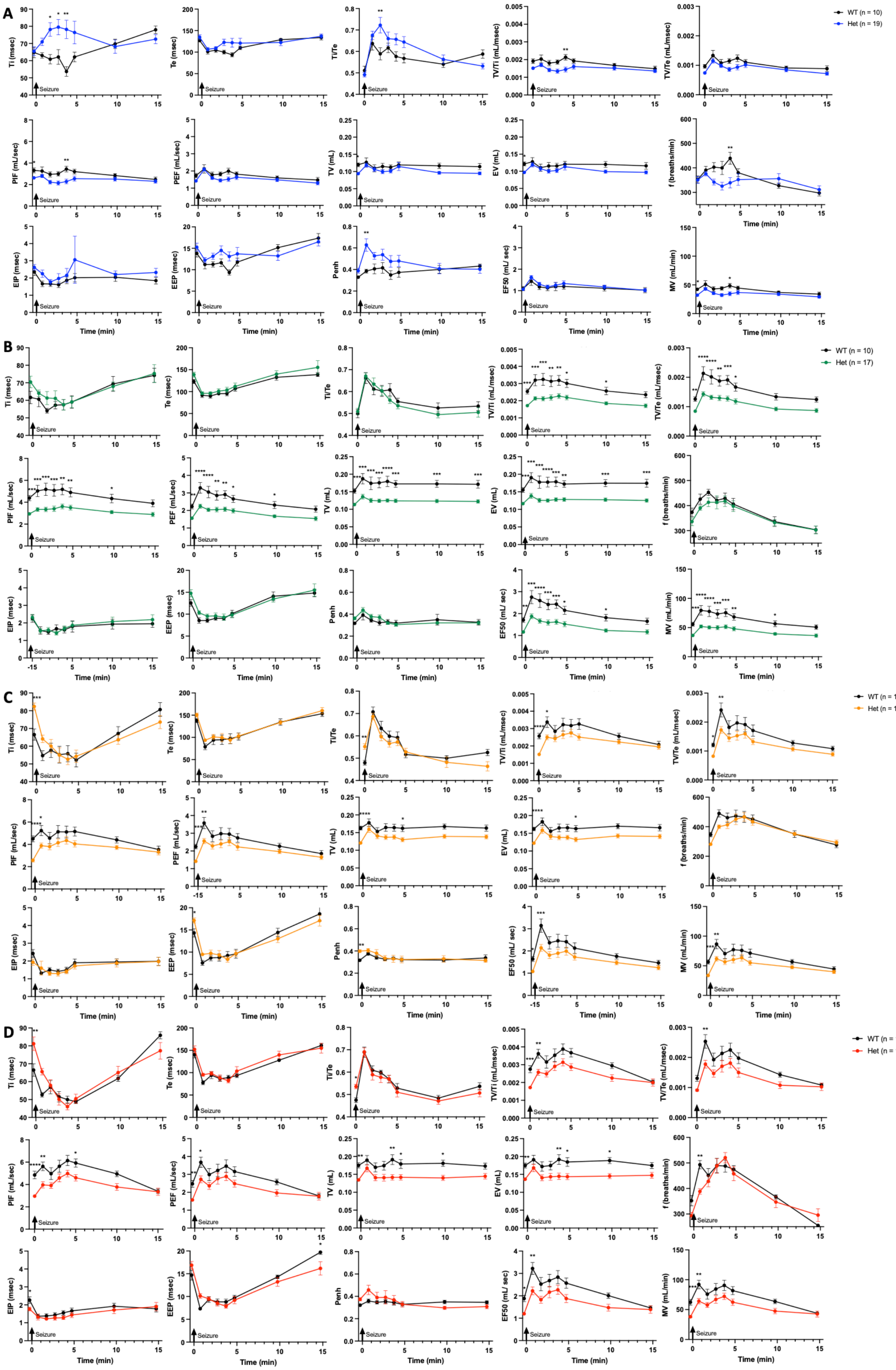
A
WT (n = 10)
Het (n = 19)
Ti (msec)
Te (msec)
Ti/Te
TV/Ti (mL/msec)
TV/Te (mL/msec)
PIF (mL/sec)
PEF (mL/sec)
TV (mL)
EV (mL)
f (breaths/min)
EIP (msec)
EEP (msec)
Penh
EF50 (mL/ sec)
MV (mL/min)
Seizure
Time (min)
B
WT (n = 10)
Het (n = 17)
C
WT (n = 10)
Het (n = 14)
D
WT (n = 10)
Het (n = 13)

**Figure S3.** Respiration following electrically induced 6 Hz seizure in WT and shown for week 1 (**A**), week 2 (**B**) week 3 (**C**), and week 4 (**D**). Fifteen minutes of post-seizure respiratory monitoring is not normalized to baseline: time of inspiration (Ti), time of expiration (Te), time in inspiration/time of expiration (Ti/Te), inspiratory drive (TV/Ti), expiratory drive (TV/Te), peak inspiratory flow (PEF), peak expiratory flow (PEF), tidal volume (TV), expiratory volume (EV), frequency (f), end inspiratory pause (EIP), end expiratory pause (EEP), enhanced pause (Penh), expiratory flow 50% through expiration (EF50), and minute ventilation (MV). Two-way ANOVA, *P<0.05, **P<0.01, ***P<0.001, ****P<0.0001.
